# Association between white-matter lesion volume and heart-rate variability: A comparison of T1w, T1w+T2w and T1w+FLAIR-based segmentations using FreeSurfer SAMSEG

**DOI:** 10.1101/2025.11.18.689161

**Authors:** Tyler D. Robinson, Stefano Cerri, Fanghan E. Zhang, Koen van Leemput, J. Jean Chen

**Author notes:** Corresponding author: Tyler D. Robinson, Ph.D., Rotman Research Institute, Baycrest, 3560 Bathurst Street, Toronto ON, Canada M6A 2E1.

## Abstract

**Background:** Heart disease is a crucial risk factor for brain degeneration, but the specific mechanisms of this association are still under investigation. Notably, the development of white-matter lesions, a hallmark of cerebral small-vessel disease with connections to progressive neurodegeneration and cognitive decline, has been linked to cardiovascular disease. As such, measures of cardiovascular function such as heart-rate variability may represent accessible indicators of white-matter lesion burden.

**Methods:** This study used data from healthy adults drawn from the Leipzig Study for Mind-Body-Emotion Interactions (LEMON) dataset to examine heart function through measures of heart-rate variability, as well as their relationship to cerebral small-vessel disease as quantified through white-matter lesion volumes, using FreeSurfer SAMSEG. White-matter lesion volumes were segmented using 3-Tesla T1w+FLAIR, T1w+T2w or T1w images alone.

**Results:** Unadjusted exploratory results suggested that lower heart-rate variability was associated with higher lesion volume, a cerebral hallmark of hypertension. After controlling for the effect of age on lesion volume, low-frequency heart rate variability specifically was found to interact with age-related lesion volume. Moreover, we showed that the relationship between white-matter lesion volume and heart-rate variability is apparent using T1w+FLAIR or T1w+T2w based lesion estimates could be observed with T1w-based lesion estimation alone, opening new avenues for studying cerebral small-vessel disease.

**Discussion:** These results suggest heart rate variability as a factor in the translation between cardiovascular risk and white matter degeneration, as well as an accessible and non-invasive indicator of white-matter lesion burden and associated cognitive impairment in later life. The unique association between reduced low-frequency band heart-rate variability and age-related lesion burden may help to establish the link between autonomic control of cardiac function and brain health in aging.

## 1 INTRODUCTION

Cardiovascular diseases serve as a crucial backdrop for brain diseases such as dementia, cerebral small vessel disease (cSVD) and cognitive impairment. cSVD, a common and underdiagnosed outcome of chronic hypertension (1), represents a critical link between cardiovascular health and brain pathology and serves as the most common cause of vascular cognitive impairment (2–4). As a hallmark of cSVD, white-matter lesions (WMLs) are commonly observed in aging, characterized by neuronal loss, demyelination and gliosis on neuropathologic examination (5,6), and are often associated with cognitive dysfunction, attention deficit, executive dysfunction, apathy and depression (7). cSVD is the narrowing of the small vessel lumen and failure of cerebral autoregulation (8), which suggests that cardiovascular health could be a target for early intervention in cSVD and its sequelae, including heightened vulnerability to the functional impacts of WM infarcts (9). While chronic hypertension has previously been associated with declines in WM microstructural integrity (10), improved cardiovascular health and physical exercise have been associated with improved WM integrity and reduced WML burden (11,12). Thus, there is growing interest not only in understanding the etiology of cSVD, but also in having accessible cardiovascular markers that predict cSVD risk.

Heart-rate variability (HRV), as a marker of cardiovascular health, is also an accessible and widely used metric of autonomic nervous (ANS) activity (13–16). There is a growing body of evidence suggesting that a higher risk for cSVD is associated with lower HRV (17,18). Indeed, low HRV has been shown to both associate cross-sectionally with WML volume, and to lead WML progression longitudinally in patients with sleep apnea (19–21). Moreover, cross-sectional mendelian randomization analysis suggests a potential causal effect of HRV on WMLs (22), and increased night-time HRV has been longitudinally associated with cSVD progression in elderly individuals (21). Conversely, decreased nocturnal HRV has been linked cross-sectionally to cSVD in patients with diabetes (23) and arteriosclerotic cSVD (22). Finally, a lower complexity of heart rate has been tied longitudinally to a more rapid decline in cognitive function with age (24), which may be the outcome of critical impacts on underlying tissue integrity. These findings highlight the complex relationship between HRV and cSVD in patient populations. Prior explorations of heart rate and pulse measures as they relate to white matter lesions have found consistent cross-sectional associations, but inconsistent longitudinal relationships across aging and significant heterogeneity between studies, in part attributed to the complexity of modeling effects of blood pressure and heart rate (25). Indeed, HRV also reflects the balance of the sympathetic and parasympathetic nervous systems, and different HRV metrics can differentially reflect stress management underlying cerebrovascular pathology (26) as an additional pathophysiological mechanism underlying cSVD.

In addition to summary parameters such as the root-mean squared successive differences (RMSSD) and the standard deviation of NN intervals (SDNN), HRV can be quantified as high-frequency (HF) and low-frequency (LF) HRV. HF-HRV has been associated with vagal and sympathetic activity, while LF-HRV has been associated with baroflex activity as well as a mixture of sympathetic and parasympathetic activity (27). The sympathetic nervous system influences both small resistance arteries and large arteries, with evidence suggesting a reciprocal relationship between endothelial function and sympathetic activity (28,29). Cardiovascular diseases often feature increased sympathetic outflow and decreased endothelial function (28). Nonetheless, the distinction between the relationships of HF- and LF-HRV with cSVD remains largely unknown. Prior investigation into the histopathology of WML formation has further suggested that the cause of WML development, and its relationship to cSVD, is regionally heterogeneous across the WM (30–32). Deep-WM regions are theoretically more sensitive to cSVD, while the periventricular WM are relatively more likely to be determined by hypertensive atherosclerosis compared to deeper WM tissue (31). However, potential diffuse deep-WM lesions may also represent a confound for the purposes of this investigation, as WM signal abnormalities are present even in healthy aging (33,34). The cumulative volume of small abnormalities may then skew analyses comparing HRV to WML volume segmentations, making it necessary to account for spatial distribution of WML tissue, particularly when comparing HRV and WML associations in relation to age.

Secondary to the presented biological questions is the accessibility of WML data. WML segmentation has relied heavily on the availability of T2-weighted fluid-attenuated inversion recovery (FLAIR) magnetic resonance imaging (MRI) data (33,35). FLAIR, while excellent for detecting larger established lesions, is limited in its ability to identify actively developing lesions in earlier stages due to lower boundary contrast and smaller size (36,37). Adding complementary T1-weighted (T1w) images to FLAIR images for lesion segmentation is advantageous (38). T1w images offer higher contrast between different brain tissues, aiding in the differentiation of lesions from surrounding tissues. The joint use of T1w and FLAIR data for WML has become a method of choice in contemporary study (39–41). Although FLAIR is routinely used to visualize white matter lesions in clinical imaging, some research and legacy datasets lack FLAIR acquisitions. We therefore evaluated the consistency of WML associations across SAMSEG input configurations, including T1w alone. In the absence of FLAIR images, T1w- and T2-weighted (T2w) images have been used jointly to segment WMLs (42), but many times even T2w images are unavailable. Accordingly, there have been other studies that define and use T1w white matter signal abnormalities detected by FreeSurfer (43) as a measure of WML in aging and Alzheimer’s disease populations (44–46). More recently, a random-forest-based WML classifier performed using T1w images alone was proposed (47) and compared successfully against the use of FLAIR data in the same classifier on multiple data sets of brain aging, including the Alzheimer’s Disease Neuroimaging Initiative (ADNI) (48), the Child and Adolescent NeuroDevelopment Initiative (CANDI) (49) and the Open Access Series of Imaging Studies (OASIS) (50) in terms of spatial overlaps and correlation coefficients (51).

In this work, we focus on the link between WMLs and HRV in a cohort of healthy young and older adults, expanding the analysis to HF- and LF-HRV. Secondarily, using the FreeSurfer Sequence Adaptive Multimodal Segmentation (SAMSEG) method for proof-of-concept (52), we demonstrate the comparability amongst WML metrics assessed based on T1w+FLAIR, T1w+T2w and T1w-alone, in terms of the observed WML-HRV relationships. In consideration of varying regional sensitivity to WM hyperintensities by FLAIR, including lower sensitivity in deep-WM regions that may skew estimation between segmentation modalities (53,54), as well as the possibility that diffuse deep WM signal abnormalities may include lacunes and complicate interpretation in relation to cSVD (55), we additionally examine these relationships in restricted analyses specific to periventricular WM tissue while excluding outlying deep-WM voxels.

## 2 MATERIALS AND METHODS

### 2.1 Participants

Participant data included in this study were drawn from the Leipzig Study for Mind-Body-Emotion Interactions (LEMON) publicly available data set (56). 227 participants aged 20 to 77 were included in the original sample, with a notable gap in representation between the ages of 35 and 59. Participants were recruited in age groups of 153 young (mean age: 25.1, 45 female) and 74 older (mean age: 67.6, 37 female) members and subsequently coded into age groups in 5-year intervals (**Supplemental Figure 1**). Recruited participants were described as healthy and without past or current cardiovascular disease, hypertension, or psychiatric disorders (56). Outliers were excluded on evidence of frequent voltage inconsistencies or poor signal quality following visual inspection of the physiological data, as were participants who did not complete one or more of the required imaging modalities. Participants were considered outliers for exclusion if they fell outside two standard deviations from the mean on one or more cardiovascular measures (92) to retain a nominally healthy sample. One additional participant was excluded due to not completing all necessary imaging modalities. The remaining sample consisted of 134 participants (104 young, 39 female; 30 older, 2 female). Participant data drawn from the LEMON set consisted of T1w, T2w, and FLAIR structural MRI images, peripheral physiological measurements, and demographic variables.

### 2.2 Heart-rate variability acquisition

Heart-rate measurements were acquired using a MR-compatible BrainAmp ExG MR amplifier with PowerPack battery, SyncBox synchronization interface, and BrainVision Recorder (Version 1.20) acquisition software (56). ECG acquisition was performed at a sampling rate of 1000 Hz with in time-varying sliding-window lengths of 300 seconds and window step/grid interval of 60 seconds, giving an 80% overlap between successive 5-minute analysis windows. ECG electrodes were taped to the lumbar region superior to the coccyx (ground), upper back at the level of the seventh cervical vertebra (reference), the left-hand side at the level of the tenth rib (recording) during concurrent resting state fMRI acquisition (56). HRV metrics analyzed consisted of root mean square of successive differences between normal heartbeats (RMSSD), as well as frequency domain low-frequency (LF) and high frequency (HF) variability bands of standard deviation of normal-to-normal intervals (SDNN), quantified as LF: 0.04-0.15 Hz (SDNN.LF) and HF: 0.15-0.40 Hz (SDNN.HF) respectively, following established standards (57), and log-transformed as applicable. All described metrics were generated using Kubios HRV software version 3.4.1 (Kubios Oy, Kuopio, Finland).

### 2.3 Imaging data acquisition

MRI images were acquired using a 3 Tesla Siemens MAGNETOM Verio system (Siemens Healthineers, Erlangen, Germany). The scans include a T1w (MP2RAGE) anatomical dataset acquired in sagittal orientation as a single 3D volume with 176 interleaved slices, voxel size= 1 mm, field of view = 256 mm, TR = 5000ms, TE = 2.92ms, TI1 = 700ms, TI2 = 2500ms, FA1=4°, FA2=5°, bandwidth=240 Hz/pixel, echo spacing=6.9ms, pre-scan normalization, and a GRAPPA acceleration factor of 3 acquired over 8 minutes and 22 seconds. T2w scans were acquired in sagittal acquisition orientation as one 3D volume with 176 slices, TR = 3200 ms, TE = 409 ms, FA = variable, with pre-scan normalization, echo spacing = 3.42 ms, bandwidth = 751 Hz/pixel, FOV = 256 mm, voxel size = 1 mm isotropic, and GRAPPA acceleration factor 2 acquired over a duration of 4 minutes and 43 seconds. FLAIR data were acquired in axial acquisition orientation over 28 slices, with a TR = 10000 ms, TE = 90 ms, TI = 2500 ms, FA = 180°, pre-scan normalization, echo spacing = 9.98 ms, bandwidth = 199 Hz/pixel, FOV = 220 mm, voxel size = 0.9×0.9×4.0 mm3, interleaved slice order, over a duration of 4 minutes and 42 seconds (56).

### 2.4 Imaging data analysis

WML volumes were segmented using FreeSurfer’s SAMSEG package (version 8.1.0) (58), a sequence-independent whole-brain Bayesian inference segmentation algorithm using a mesh-based probabilistic anatomical atlas providing spatial priors for tissue and anatomical labels, and incorporating a parametric Gaussian mixture model describing the intensity distribution of each tissue class which enables segmentation adaptive to different imaging hardware and protocols (52). Image registration in SAMSEG was performed using FreeSurfer’s rigid intermodal registration, followed by resampling into the grid of the highest-resolution image. No additional manual mask correction was performed. Three types of inputs were used with SAMSEG for each participant: (1) joint FLAIR and T1w images (T1w+FLAIR); (2) joint T1w and T2w images (T1w+T2w); (3) T1w images alone (T1w). For each input, WML volume was based on SAMSEG’s segmentation label 77 (58). In the interest of excluding deep-WM voxel outliers and limiting analysis to periventricular regions more strongly associated with hypertension, an additional set of periventricular WML measures was derived by restricting the label 77 segmentation to a region within 5 voxels of any ventricle prior to volume calculation (**Figure 1**). Periventricular WMLs were defined using a morphological dilation of 5 voxels (5 mm) from the SAMSEG-defined combined ventricular mask in each participant’s local space in fslmaths. As multiple classification criteria for “periventricular” WM exist without universal consensus, ranging from only voxels adjacent to the ventricular surface, those less than 10 mm from the ventricles, and studies which distinguish “juxtaventricular” (<3mm) and “deep-periventricular” (3-13mm) tissue (30,31), this study selected a conservative 5-voxel definition to ensure restricted analysis remained specific to tissue in close proximity to the ventricles. We directly compare WMLs based on these three approaches using pairwise correlations. All WML volume values were corrected against participant intracranial volumes from their respective scan input and demeaned prior to statistical analyses.

**Figure 1.**
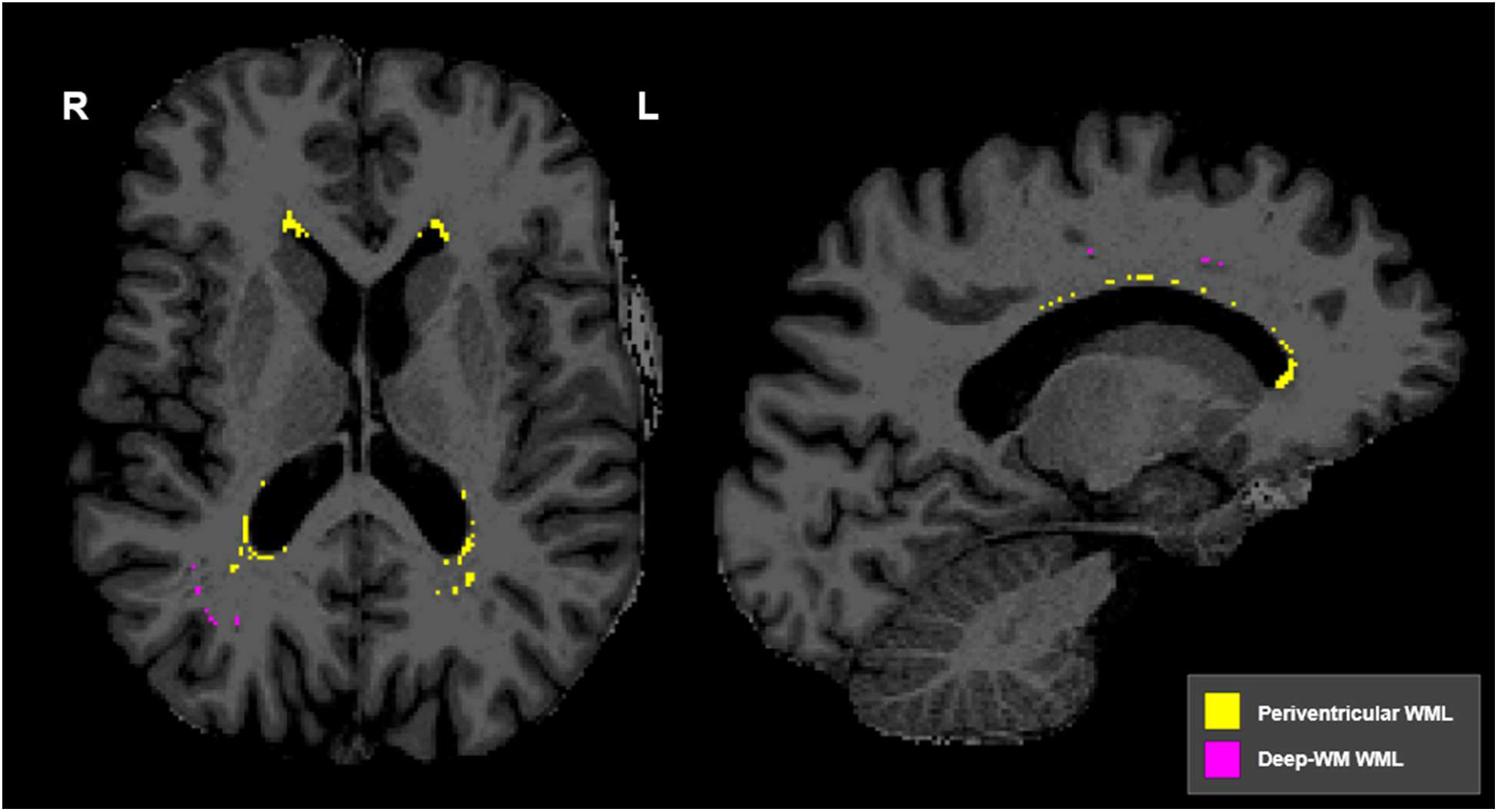
Periventricular and Deep-WM lesion voxels. Periventricular (yellow) and Deep-WM (pink) WML voxels identified by SAMSEG using T1w+FLAIR segmentation for a sample participant. Voxels included in the Periventricular analysis were restricted to the region overlapping the SAMSEG label 77 mask and a region five voxels from any ventricle defined by SAMSEG’s initial segmentation. Periventricular masking was performed per-participant and per-segmentation model (T1w+FLAIR, T1w+T2w, T1w-alone).

### 2.5 Statistical analysis

Associations between HRV and WML volume were first assessed via unadjusted exploratory linear regressions of each HRV measure (RMSSD, SDNN.LF, SDNN.HF) against each WML imaging modality (T1w+FLAIR, T1w-alone, T1w+T2w), in which HRV was treated as a cross-sectional correlate of WML. Subsequent linear regressions comprising the primary analyses examined the interaction between age and HRV by separately regressing HRV measures, along with age, sex, and the interaction effects of heart rate variability on the effects of age and sex onto volume measures from each WML modality. Sex and HRV ྾ sex interaction terms were included in the age-interaction analyses to account for potential confounds introduced by the uneven distribution of female participants between age categories noted in the sample (**Supplemental Figure 1**). Additional sex-specific exploratory sensitivity analyses were conducted for female-only and male-only subsamples by regressing HRV measures, age, and HRV ྾ age interaction coefficients onto each WML measure. Furthermore, young-only and old-only exploratory sensitivity analyses of interactions between HRV and age on WML was conducted by regression HRV measures, age, sex, HRV ྾ age, and HRV ྾ sex onto each WML measure. For all analyses following the initial direct regression of HRV onto WML, multiple comparisons correction was applied using false detection rate (FDR) correction per HRV measure, i.e. RMSSD interaction effects comprised one family, while separate corrections were performed for SDNN.LF and SDNN.HF analyses. This correction structure was repeated for exploratory age/sex restricted subsets. A separate set of analyses was conducted for periventricular WMLs, with all previous analyses repeated using WML values sampled from within 5 voxels of the ventricles. Resulting p-values from periventricular interaction models were again corrected for multiple comparisons using FDR correction.

## 3 RESULTS

Differences in estimates amongst the three approaches are illustrated in **Figure 2**, with measure agreement and biases further illustrated via Bland-Altman plots in **Figure 3**. All approaches show strong correlation in WML volume estimates across participants, indicating that participant-wise differences are detected consistently across detection approaches. Moreover, the WML values estimated by T1w+T2w and T1w were both lower than for T1w+FLAIR, with T1w+T2w more closely resembling T1w+FLAIR segmentations. Nonetheless, the Bland-Altman plots suggest proportional bias that varies depending on the pairing.

**Figure 2.**
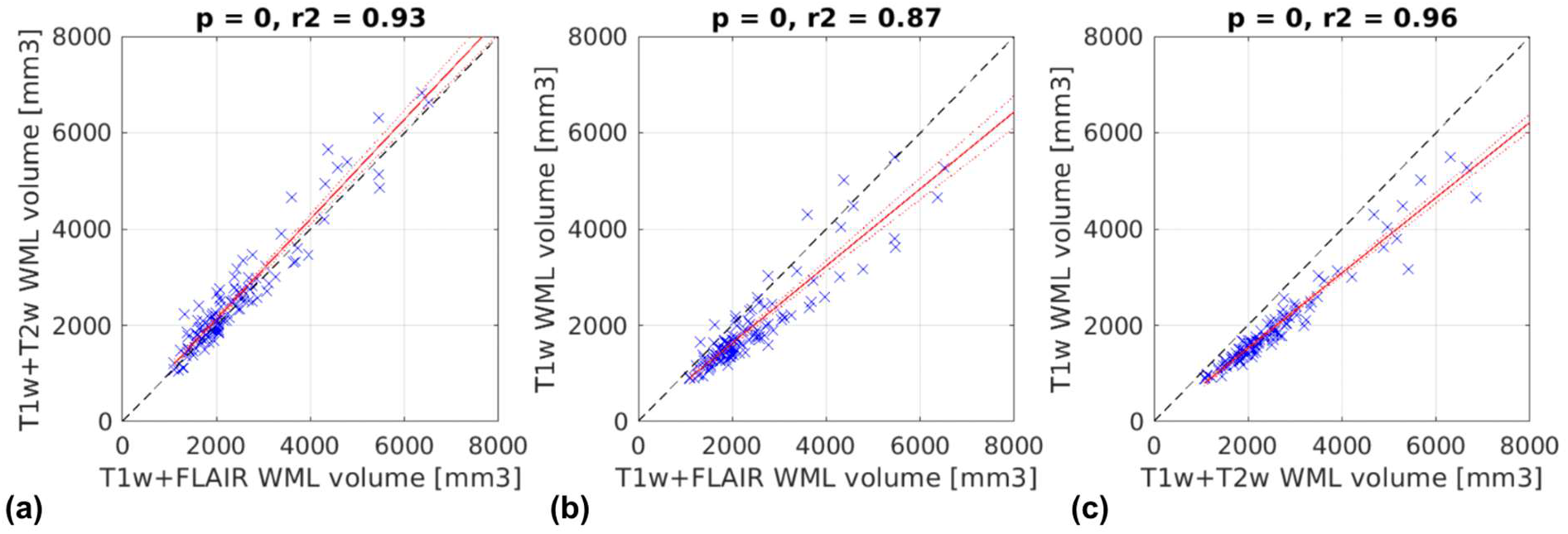
Differences in WML Estimates. Comparisons between T1w+FLAIR and T1w+T2w (a), T1w+FLAIR and T1w-alone (b), and T1w+T2w and T1w-alone (c) WML volume estimates indicate strong associations between measures, with greater agreement in volumes between T1w+FLAIR and T1w+T2w estimates than between T1w+FLAIR and T1w-alone or between T1w+T2w and T1w-alone. Dashed red lines represent the confidence interval for each regression line. The dashed black line represents the line of unity.

**Figure 3.**
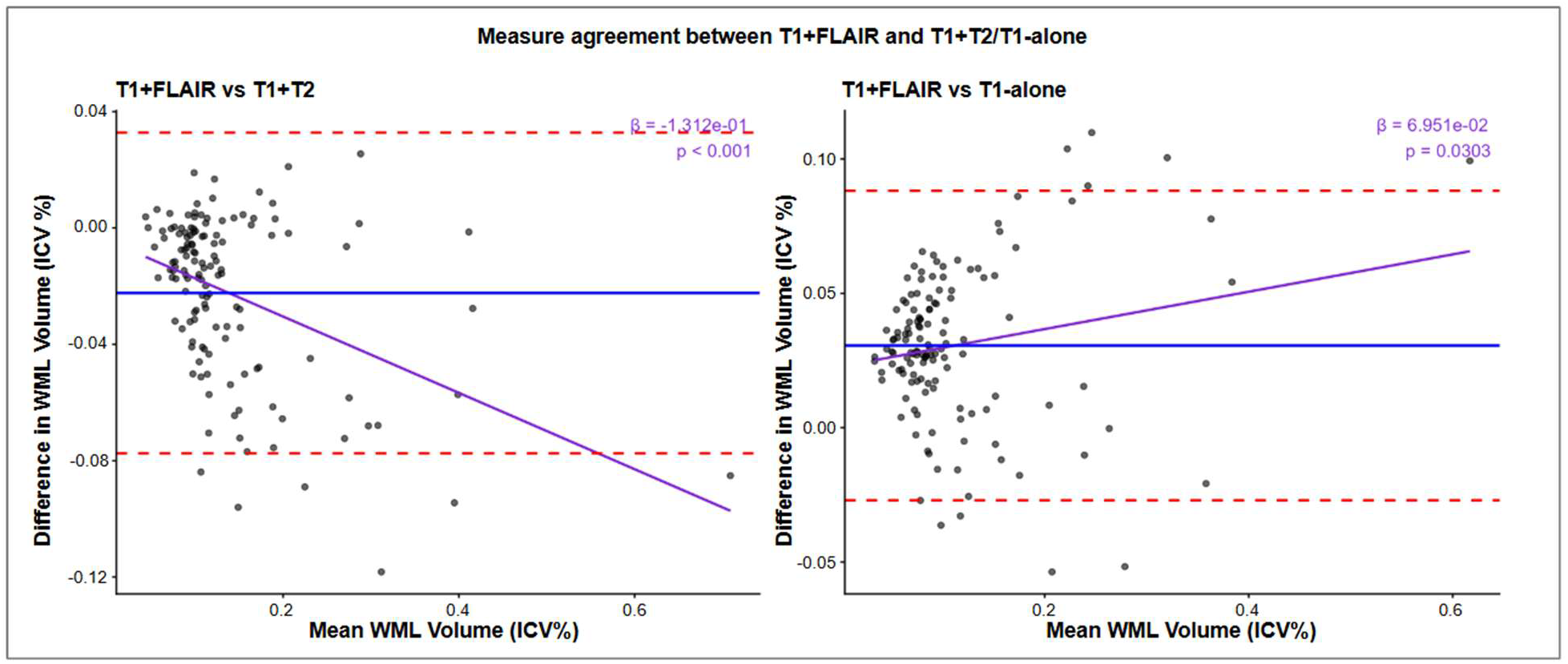
Measure agreement between WML estimates: Comparisons between T1w+FLAIR and both T1w+T2w (left) and T1w-alone WML volume estimates via Bland-Altman analysis. Blue lines denote bias between measures, while dotted red lines denote standard deviation of differences. Purple fit lines illustrate patterns in the relationship between mean WML volume estimates and the differences in WML volume estimates between measures.

All three sets of WML volume estimates were associated with HRV in exploratory analyses of unadjusted associations. Unadjusted examination of HRV as a cross-sectional correlate of WML volume found significant negative associations between all pairs of HRV-WML measures, such that in all cases, lower HRV was associated with a higher WML volume (**Supplemental Table 1**). While this relationship is reflected in both LF- and HF-HRV, HF-HRV is more strongly associated with WML volume, as reflected by the effect-size estimates. Analyses of unadjusted periventricular WML volume associations with HRV corroborated this pattern of results, with lower HRV associated with higher periventricular WML volumes in all comparisons and the strongest effects observed in HF-HRV associations (**Supplemental Table 4**).

Examination of interactions between HRV and age in prediction WML volume found significant negative interactions (i.e. a negative contribution of the age ྾ HRV term to WML, such that higher HRV was associated with a weaker relationship between age and WML volume) was found for SDNN.LF in all three sets of WML segmentations following FDR correction (**Table 1**). The pattern of interaction effects for all three sets of WML estimates is illustrated in interaction plots (**Figure 4**), showing the negative interaction of HRV on age ྾ WML associations, such that the association is more negative at higher SDNN.LF. These interactions were again identified in analyses using periventricular WML measures for all three WML segmentation approaches (**Supplemental Table 5**).

**Figure 4.**
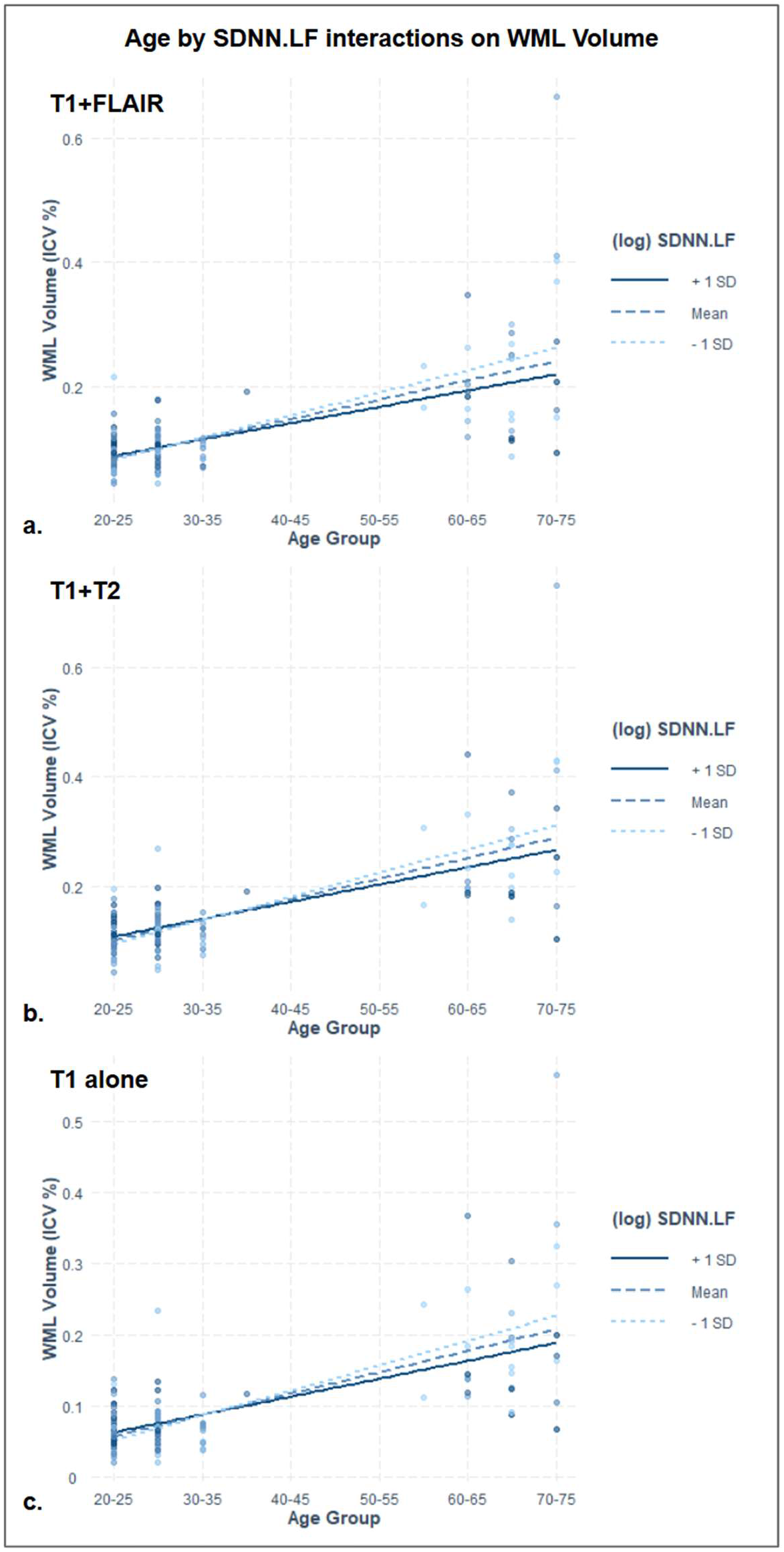
Interaction effects of HRV on age-related WML volume: Significant interaction effects are plotted for SDNN.LF interactions with age effects on T1w+FLAIR (a), T1w+T2w (b), and T1w-alone (c) WML volume estimates. Fit lines represent the age ྾ WML association at mean HRV and +/- one standard deviation from the mean.

**Table 1:**
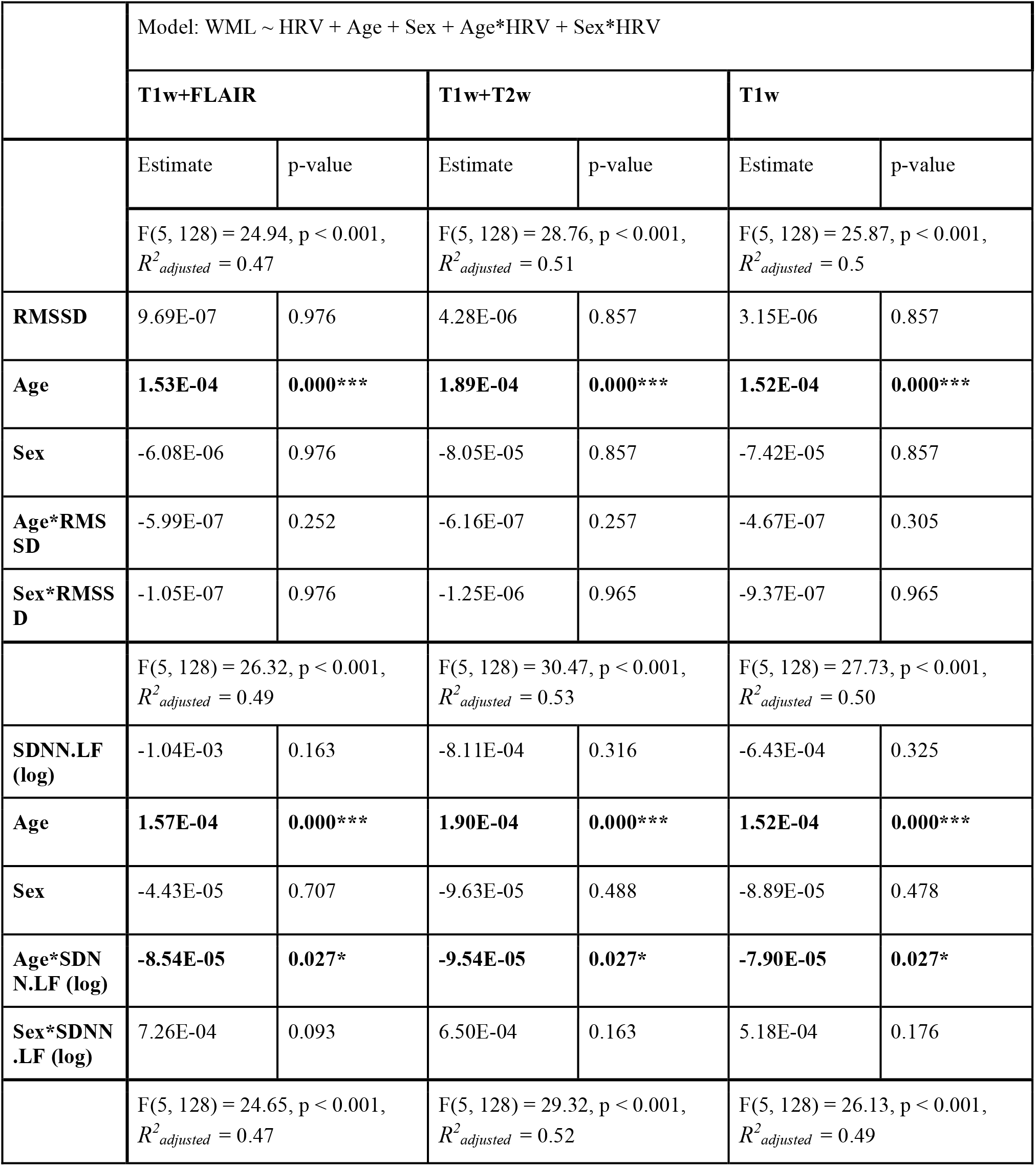
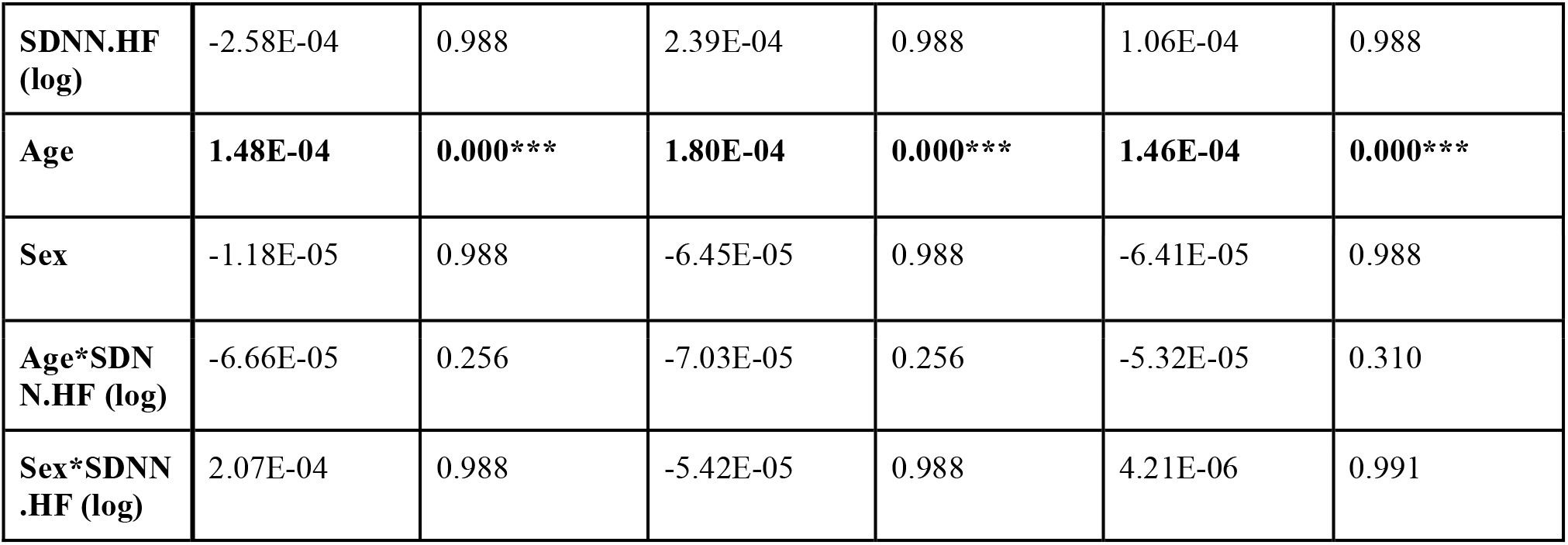
Model across all participants: Contributions of normalized and demeaned HRV, Age, Sex, and the interactions of Age*HRV and Sex*HRV to WML volume All models are linear regressions with overall F-statistics, *p*-values, and effect sizes (*R^2^_adjusted_*) listed. Slopes and FDR-corrected p-values by coefficient are noted below each regression equation and significant associations bolded with asterisks indicating significance following FDR correction by measure. (*:p<.05, **:p<.01, ***:p<.001)

|  | Model: WML ~ HRV + Age + Sex + Age*HRV + Sex*HRV |  |  |  |  |  |
|---|---|---|---|---|---|---|
|  | **T1w+FLAIR** |  | **T1w+T2w** |  | **T1w** |  |
|  | Estimate | p-value | Estimate | p-value | Estimate | p-value |
|---|---|---|---|---|---|---|
|  | F(5, 128) = 24.94, p < 0.001, *R*<sup>*2*</sup><sub>*adjusted*</sub> = 0.47 |  | F(5, 128) = 28.76, p < 0.001, *R*<sup>*2*</sup><sub>*adjusted*</sub> = 0.51 |  | F(5, 128) = 25.87, p < 0.001, *R*<sup>*2*</sup><sub>*adjusted*</sub> = 0.5 |  |
| **RMSSD** | 9.69E-07 | 0.976 | 4.28E-06 | 0.857 | 3.15E-06 | 0.857 |
| **Age** | **1.53E-04** | **0.000\*\*\*** | **1.89E-04** | **0.000\*\*\*** | **1.52E-04** | **0.000\*\*\*** |
| **Sex** | -6.08E-06 | 0.976 | -8.05E-05 | 0.857 | -7.42E-05 | 0.857 |
| **Age\*RMSSD** | -5.99E-07 | 0.252 | -6.16E-07 | 0.257 | -4.67E-07 | 0.305 |
| **Sex\*RMSSD** | -1.05E-07 | 0.976 | -1.25E-06 | 0.965 | -9.37E-07 | 0.965 |
|  | F(5, 128) = 26.32, p < 0.001, *R*<sup>*2*</sup><sub>*adjusted*</sub> = 0.49 |  | F(5, 128) = 30.47, p < 0.001, *R*<sup>*2*</sup><sub>*adjusted*</sub> = 0.53 |  | F(5, 128) = 27.73, p < 0.001, *R*<sup>*2*</sup><sub>*adjusted*</sub> = 0.50 |  |
| **SDNN.LF (log)** | -1.04E-03 | 0.163 | -8.11E-04 | 0.316 | -6.43E-04 | 0.325 |
| **Age** | **1.57E-04** | **0.000\*\*\*** | **1.90E-04** | **0.000\*\*\*** | **1.52E-04** | **0.000\*\*\*** |
| **Sex** | -4.43E-05 | 0.707 | -9.63E-05 | 0.488 | -8.89E-05 | 0.478 |
| **Age\*SDNN.LF (log)** | **-8.54E-05** | **0.027\*** | **-9.54E-05** | **0.027\*** | **-7.90E-05** | **0.027\*** |
| **Sex\*SDNN.LF (log)** | 7.26E-04 | 0.093 | 6.50E-04 | 0.163 | 5.18E-04 | 0.176 |
|  | F(5, 128) = 24.65, p < 0.001, *R*<sup>*2*</sup><sub>*adjusted*</sub> = 0.47 |  | F(5, 128) = 29.32, p < 0.001, *R*<sup>*2*</sup><sub>*adjusted*</sub> = 0.52 |  | F(5, 128) = 26.13, p < 0.001, *R*<sup>*2*</sup><sub>*adjusted*</sub> = 0.49 |  |
| **SDNN.HF (log)** | -2.58E-04 | 0.988 | 2.39E-04 | 0.988 | 1.06E-04 | 0.988 |
| **Age** | **1.48E-04** | **0.000\*\*\*** | **1.80E-04** | **0.000\*\*\*** | **1.46E-04** | **0.000\*\*\*** |
| **Sex** | -1.18E-05 | 0.988 | -6.45E-05 | 0.988 | -6.41E-05 | 0.988 |
| **Age\*SDNN.HF (log)** | -6.66E-05 | 0.256 | -7.03E-05 | 0.256 | -5.32E-05 | 0.310 |
| **Sex\*SDNN.HF (log)** | 2.07E-04 | 0.988 | -5.42E-05 | 0.988 | 4.21E-06 | 0.991 |

As age was strongly associated with WML volume, to reduce the contribution of age, we examined the contribution of HRV to WML in young adults alone (**Table 2**), as well as older adults alone (**Table 3**). The exploratory results showed no significant HRV association with WML after multiple comparisons correction for either full WML volumes or periventricular WML volumes in either exploratory subsample (**Supplemental Table 6, Supplemental Table 7**). Notably, the older subsample contained only two female participants, severely limiting the interpretability of results from this subsample in isolation.

**Table 2:** Exploratory model in young participants alone: Contributions of normalized and demeaned HRV, Age, Sex, and the interactions of Age*HRV and Sex*HRV to WML volume All models are linear regressions with overall F-statistics, *p*-values, and effect sizes (*R^2^_adjusted_*) listed. Slopes and FDR-corrected p-values by coefficient are noted below each regression equation and significant associations bolded with asterisks indicating significance following FDR correction by measure. (*:p<.05, **:p<.01, ***:p<.001)

|  | Model: WML ~ HRV + Age + Sex + Age*HRV + Sex*HRV |  |  |  |  |  |
|---|---|---|---|---|---|---|
|  | **T1w+FLAIR** |  | **T1w+T2w** |  | **T1w** |  |
|  | Estimate | p-value | Estimate | p-value | Estimate | p-value |
|  | F(5, 98) = 1.14, p = 0.346, *R<sup>2</sup><sub>adjusted</sub>* = 0.01 |  | F(5, 98) = 1.27, p = 0.281, *R<sup>2</sup><sub>adjusted</sub>* = 0.01 |  | F(5, 98) = 0.78, p = 0.568, *R<sup>2</sup><sub>adjusted</sub>* = -0.01 |  |
| **RMSSD** | -1.71E-06 | 0.731 | 1.58E-06 | 0.731 | 1.01E-06 | 0.765 |
| **Age** | 6.50E-05 | 0.668 | 7.58E-05 | 0.668 | 3.99E-05 | 0.668 |
| **Sex** | -2.70E-05 | 0.731 | -1.03E-04 | 0.668 | -8.25E-05 | 0.668 |
| **Age\*RMSSD** | -1.35E-06 | 0.668 | -1.93E-06 | 0.668 | -1.42E-06 | 0.668 |
| **Sex\*RMSSD** | 1.86E-06 | 0.668 | 1.12E-06 | 0.731 | 8.58E-07 | 0.731 |
|  | F(5, 98) = 2.57, p = 0.031, *R<sup>2</sup><sub>adjusted</sub>* = 0.07 |  | F(5, 98) = 1.37, p = 0.243, *R<sup>2</sup><sub>adjusted</sub>* = 0.02 |  | F(5, 98) = 0.82, p = 0.541, *R<sup>2</sup><sub>adjusted</sub>* = -0.01 |  |
| **SDNN.LF (log)** | -9.27E-04 | 0.073 | -5.20E-04 | 0.443 | -3.46E-04 | 0.519 |
| **Age** | 8.06E-05 | 0.258 | 9.40E-05 | 0.258 | 5.33E-05 | 0.443 |
| **Sex** | -4.38E-05 | 0.561 | -1.03E-04 | 0.434 | -8.40E-05 | 0.439 |
| **Age\*SDNN.LF (log)** | 1.22E-05 | 0.938 | 1.11E-04 | 0.646 | 6.67E-05 | 0.740 |
| **Sex\*SDNN.LF (log)** | 5.35E-04 | 0.073 | 2.59E-04 | 0.443 | 1.80E-04 | 0.519 |
|  | F(5, 98) = 2.89, p = 0.018, *R<sup>2</sup><sub>adjusted</sub>* = 0.08 |  | F(5, 98) = 1.52, p = 0.191, *R<sup>2</sup><sub>adjusted</sub>* = 0.02 |  | F(5, 98) = 0.75, p = 0.590, *R<sup>2</sup><sub>adjusted</sub>* = -0.01 |  |
| **SDNN.HF (log)** | -8.59E-05 | 0.804 | 3.51E-04 | 0.674 | 1.42E-04 | 0.798 |
| **Age** | 3.73E-05 | 0.674 | 4.90E-05 | 0.674 | 2.73E-05 | 0.798 |
| **Sex** | -1.90E-05 | 0.798 | -8.94E-05 | 0.674 | -7.57E-05 | 0.674 |
| **Age\*SDNN.HF (log)** | -4.79E-04 | 0.114 | -3.84E-04 | 0.379 | -2.03E-04 | 0.674 |
| **Sex\*SDNN.HF (log)** | 3.96E-04 | 0.312 | 8.90E-05 | 0.798 | 7.11E-05 | 0.798 |

**Table 3:**
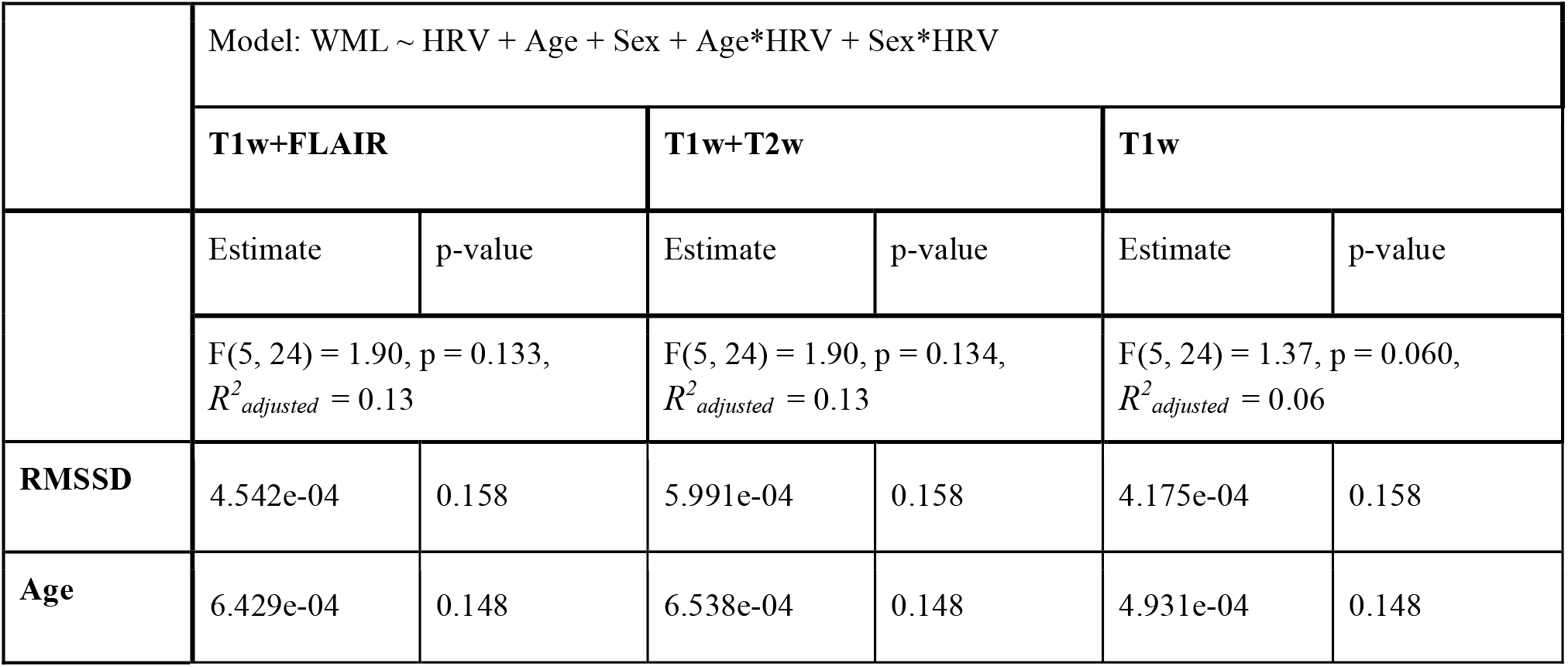
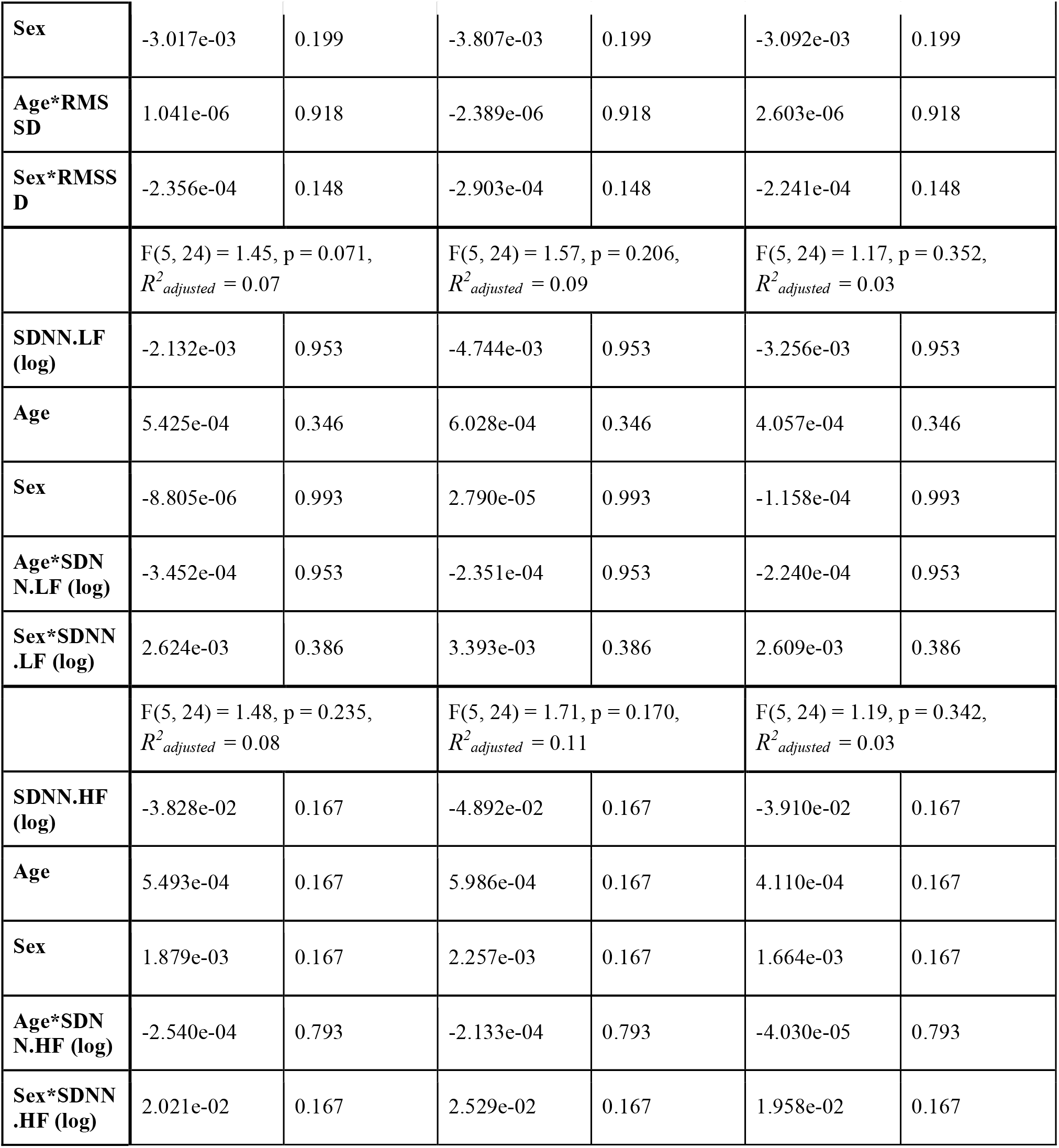
Exploratory model in older participants alone: Contributions of normalized and demeaned HRV, Age, Sex, and the interactions of Age*HRV and Sex*HRV to WML volume All models are linear regressions with overall F-statistics, *p*-values, and effect sizes (*R^2^_adjusted_*) listed. Slopes and FDR-corrected p-values by coefficient are noted below each regression equation and significant associations bolded with asterisks indicating significance following FDR correction by measure. (*:p<.05, **:p<.01, ***:p<.001)

|  | Model: WML ~ HRV + Age + Sex + Age*HRV + Sex*HRV |  |  |  |  |  |
|---|---|---|---|---|---|---|
|  | **T1w+FLAIR** |  | **T1w+T2w** |  | **T1w** |  |
|  | Estimate | p-value | Estimate | p-value | Estimate | p-value |
|  | F(5, 24) = 1.90, p = 0.133, *R<sup>2</sup><sub>adjusted</sub>* = 0.13 |  | F(5, 24) = 1.90, p = 0.134, *R<sup>2</sup><sub>adjusted</sub>* = 0.13 |  | F(5, 24) = 1.37, p = 0.060, *R<sup>2</sup><sub>adjusted</sub>* = 0.06 |  |
| **RMSSD** | 4.542e-04 | 0.158 | 5.991e-04 | 0.158 | 4.175e-04 | 0.158 |
| **Age** | 6.429e-04 | 0.148 | 6.538e-04 | 0.148 | 4.931e-04 | 0.148 |
| **Sex** | -3.017e-03 | 0.199 | -3.807e-03 | 0.199 | -3.092e-03 | 0.199 |
| **Age\*RMSSD** | 1.041e-06 | 0.918 | -2.389e-06 | 0.918 | 2.603e-06 | 0.918 |
| **Sex\*RMSSD** | -2.356e-04 | 0.148 | -2.903e-04 | 0.148 | -2.241e-04 | 0.148 |
|  | F(5, 24) = 1.45, p = 0.071, *R<sup>2</sup><sub>adjusted</sub>* = 0.07 |  | F(5, 24) = 1.57, p = 0.206, *R<sup>2</sup><sub>adjusted</sub>* = 0.09 |  | F(5, 24) = 1.17, p = 0.352, *R<sup>2</sup><sub>adjusted</sub>* = 0.03 |  |
| **SDNN.LF (log)** | -2.132e-03 | 0.953 | -4.744e-03 | 0.953 | -3.256e-03 | 0.953 |
| **Age** | 5.425e-04 | 0.346 | 6.028e-04 | 0.346 | 4.057e-04 | 0.346 |
| **Sex** | -8.805e-06 | 0.993 | 2.790e-05 | 0.993 | -1.158e-04 | 0.993 |
| **Age\*SDNN.LF (log)** | -3.452e-04 | 0.953 | -2.351e-04 | 0.953 | -2.240e-04 | 0.953 |
| **Sex\*SDNN.LF (log)** | 2.624e-03 | 0.386 | 3.393e-03 | 0.386 | 2.609e-03 | 0.386 |
|  | F(5, 24) = 1.48, p = 0.235, *R<sup>2</sup><sub>adjusted</sub>* = 0.08 |  | F(5, 24) = 1.71, p = 0.170, *R<sup>2</sup><sub>adjusted</sub>* = 0.11 |  | F(5, 24) = 1.19, p = 0.342, *R<sup>2</sup><sub>adjusted</sub>* = 0.03 |  |
| **SDNN.HF (log)** | -3.828e-02 | 0.167 | -4.892e-02 | 0.167 | -3.910e-02 | 0.167 |
| **Age** | 5.493e-04 | 0.167 | 5.986e-04 | 0.167 | 4.110e-04 | 0.167 |
| **Sex** | 1.879e-03 | 0.167 | 2.257e-03 | 0.167 | 1.664e-03 | 0.167 |
| **Age\*SDNN.HF (log)** | -2.540e-04 | 0.793 | -2.133e-04 | 0.793 | -4.030e-05 | 0.793 |
| **Sex\*SDNN.HF (log)** | 2.021e-02 | 0.167 | 2.529e-02 | 0.167 | 1.958e-02 | 0.167 |

To examine sex differences in the contribution of HRV to WML, we divided the sample into male and female groups for exploratory analyses, which resulted in significant positive RMSSD ྾ age interactions on WML volumes in female participants for T1w+FLAIR, T1w+T2w, and T1w-alone-based WML estimations, such that lower RMSSD was associated with weaker age effects on WML volume (**Supplemental Table 2)**. Conversely, lower SDNN.LF was associated with stronger age effects on WML volume in all three modalities. Notably, these effects are no longer present when excluding the two female participants representing all females over 55 years of age, suggesting that these results are strongly influenced by the few older female samples. Analysis of periventricular WML volumes similarly found positive age ྾ RMSSD interaction effects on WML volumes for all three sets of WML segmentations, as well as negative age ྾ SDNN.LF interaction effects on T1w+T2W and T1w-alone-based periventricular WML volumes. Unlike the full-WML segmentations, negative age ྾ SDNN.HF interaction effects on WML volumes were also observed for all three sets of periventricular WML segmentations (**Supplemental Table 8**). No significant HRV ྾ age interactions were identified in the male sample for either the full WML volume or periventricular WML volume analyses (**Supplemental Table 3, Supplemental Table 9**). Model diagnostics, including standardized beta coefficients, 90% confidence intervals, Breusch-Pagan heteroscedasticity tests, Normal Q-Q plots, and residuals vs. leverage plots were generated for all models and have been indexed and made available for review in **Supplemental File 1**.

## 4 DISCUSSION

HRV is a surrogate marker of the autonomic nervous system that is intrinsically connected to blood pressure. In this work, we identified a HRV domain specific LF-HRV interaction with WML volume in aging. Importantly, we additionally demonstrated that this association is detectable not only based on WML estimated by conventional T1w+FLAIR data, but also with volumes estimated from T1w+T2w, or even from T1w data alone. Thus, this proof-of-principle study not only suggests the potential of LF-HRV as a distinct marker of cSVD, but also supports the feasibility of WML quantification using widely available T1w data in this context, with the potential to open new avenues of research when other T2w and FLAIR data are not available.

### 4.1 WML vs. HRV

HF-HRV is most commonly associated with parasympathetic reactivity, broadly including stress response, anxiety levels, and sleep quality (59,60). Existing evidence indicates that lowered parasympathetic activity in aging represents the greatest contributor to declining overall HRV in older participants (61,62). LF-HRV, in contrast, is complexly associated with both parasympathetic and sympathetic reactivity, and has been suggested to reflect interactions between the two systems (63–65). While associations have been reported between HRV measures and both WML burden as well as the development of cognitive impairment (17,19,24,66), the relative predictive power of HF-as opposed to LF-HRV remains inconclusive in existing literature. Both measures have been associated with poor cardiovascular health outcomes separately (67–69), and autonomic dysregulation in the context of stroke is negatively associated with LF-HRV and LF/HF ratios (70). However, data regarding the contributions of each component of HRV to associations with WML burden separately are sparse, as HRV is more commonly represented with RMSSD (17,71,72).

Our exploratory investigation suggested that both SDNN.LF and SDNN.HF are more strongly associated with WML volume than RMSSD, and that this was the case for all three WML estimation approaches. Moreover, the exploratory unadjusted HRV-WML associations were strongest with HF-HRV (**Supplemental Table 1**). However, neither RMSSD nor SDNN.HF associations with WML volume are present in primary models controlling for the effects of age, indicating that these associations are confounded by age effects on both HRV and WML. Conversely, the only measure which demonstrated significant HRV ྾ age interactions in the HRV-WML association was LF-HRV, again regardless of segmentation approach; this is in contrast to an absence of such interaction effects by other HRV measures. This may suggest that LF-HRV is sensitive to either age-related associations in WML burden or in HRV profiles between young and old cohorts. Trends in the exploratory findings are in agreement with some previous studies that have identified HF-rather than LF-HRV as being differentially linked to WMLs (71). However, LF-HRV may offer a specific sensitivity to age-related variability in fitness and vascular health, topics of growing interest to the study of WM health in aging (73,74) (**Table 1**). Considering existing literature on relationships between HRV frequency bands and autonomic response, this suggests that sympathetic autonomic response contributes to interactions between HRV and estimated WML burden in aging in ways that are not captured by the primarily parasympathetic HF-HRV band (63–65). As the LF-HRV band is theorized to relate to the interaction of sympathetic and parasympathetic autonomic systems, the specific association between LF-HRV and WML volume observed here may capture the relationship between vagal tone and general cerebrovascular health given the established role of the vagus nerve in mediating autonomic balance and cardiovascular regulation (75,76).

Restricting our analyses to periventricular WM tissue did not alter these trends. While multiple vascular differences may contribute to the development of WML across the WM identified in previous research (31), the relative sparsity of deep-WM lesions suggest that these findings indicate the relationship between WML and HRV is driven primarily by periventricular tissue with negligible deep WM contributions. This consistency is particularly valuable in the case of T1w-alone volume estimations due to the greater susceptibility to aging-related confounds assumed to segmentation generated in the absence of T2 data. Our exploratory sensitivity analysis restricting effects to young participants findings further suggest that HRV-WML associations are negligible in younger participants (**Table 2**). While this is consistent with the understanding that cardiovascular risk rises rapidly beyond the age of 55 (77), lower general variability in WML volume in a younger cohort makes it unclear whether this represents a weaker association or limited ability to detect associations in a cohort with less age-related WM degeneration (**Supplemental Figure 4**). Examination of the mid-life age ranges not included in the LEMON sample may be necessary to determine whether there is a notable inflection point in this association.

Finally, exploratory sensitivity analyses of differential sex associations between HRV and WML (**Supplemental Tables 2 and 3**) were broadly inconclusive. The presence of effects in the female exploratory subsample analysis could suggest a trend in support of more prominent associations in female participants, consistent with past literature demonstrating stronger HRV changes in women with advancing age (78). Prior research has demonstrated sex differences in rates of both hypertension and cSVD in aging (79–81). While males have higher rates of hypertension at younger ages, this ratio is inverted by the mid 60s, with older females showing significantly higher rates of both conditions from the 60s onwards (81). However, given that these associations are driven by older age, and that the pool of older female participants available to the LEMON dataset within exclusion criteria was extremely small, our ability to analyse sex-specific associations is limited. We were similarly unable to clarify whether significant sex differences were present in the HRV-WML relationship in the older adults alone. Examination of model diagnostics in the older subsample further indicates that the two older female participants exert a disproportionate degree of leverage in these models, limiting our ability to draw conclusions from the older subgroup alone (**Supplemental File 1**). As a result, the generalizability of these findings between sexes is limited in the absence of a more robust sample of older female participants.

### 4.2 T1w+FLAIR, T1w+T2w, and T1w-alone

All three WML estimation approaches identified associations between WML volume and HRV in exploratory comparisons. With side-by-side comparisons across T1w+FLAIR, T1w+T2w, and T1w-alone quantifications, we demonstrate that, T1w+T2w and T1w-alone are effective for modeling WML variance, despite the generally higher sensitivity assumed from the inclusion of FLAIR (**Supplemental Table 1**) (33), and supported by comparisons in WML volume estimates by all three approaches used in this study (**Figure 2**). This is of particular interest in the case of T1w-alone WML estimation, which showed both a significant near-unity correlation with T1w+FLAIR as well as a spatial mismatch only slightly higher than for T1w+T2w. This finding suggests that, despite the acknowledged vascular specificity of FLAIR imaging, WML measurement based entirely on T1w images is broadly feasible, making WML volume estimation and analysis available to a greater number of studies. This finding is consistent with findings by Dadar et al. using a random-forest-based segmentation strategy (51).

Notably, direct comparisons between pairs of methods showed stronger agreement in WML values based on T1w+FLAIR and T1w+T2w as well as higher WML volume estimates than using T1w-alone (**Figure 2**, **Figure 3**). This is consistent with the improved sensitivity afforded by the presence of both T2w and T1w contrast and supports the use of T1w+FLAIR or T1w+T2w when available. While the incorporation of FLAIR data into WML volume estimations results in the inclusion of a greater volume of voxels in WML estimations, both T1w and T1w+T2w estimates are sensitive enough to variation in WML volume across HRV to identify consistent trends to those found using the T1w+FLAIR approach. The observation of proportional bias in T1w-based compared to T1w-FLAIR-based WML measurements (**Figure 3**) suggests that despite the strong agreement, measurements made using different scans are not necessarily exchangeable. Lastly, although all three approaches appear to reliably identify the anticipated relationship between HRV and WML, our approach of associating variability in LF-HRV (i.e. SDNN.LF) to WML estimates may lead to an accessible proxy measure for estimating age-related WML burden. However, this suggestion is predicated on the association of WML and HRV in the absence of additional modelled covariates.

### 4.3 Limitations and future directions

In its current form, the study relies on cross-sectional data. As the age component of lifespan HRV-WML associations is of interest to this study, the inclusion of longitudinal data would benefit our ability to interpret the discussed age interactions. Additionally, after rigorous quality control, the data size became more modest, and the sex distribution is notably skewed towards male participants, especially in the oldest age group. Expanded analyses incorporating additional physiological measures, such as blood pressure and mean heart-rate, could further control for potential confound or interactions that may influence HRV-WML associations. While no main effect or interaction effect by sex was observed in the full sample, the age-skewed female sample limits our ability to assess age effects specifically within female participants. Finally, the LEMON dataset categorizes participants into quinquennial groups, but does not specify exact age, and these age groups are further categorized into overarching older and younger groups, with the 35-59 age range omitted. In the absence of the mid-life age-range, this study is limited in its ability to capture non-linear patterns that may be present between the sampled age ranges. Future examination would benefit from using exact age across a more complete age distribution to validate the effects found in this study.

### 4.4 Conclusions

While exploratory analysis of unadjusted WML-HRV associations across measures indicate robust associations between reduced HRV and greater estimated WML burden, these effects do not survive controls for the confounding effect of age. However, unique age ྾ SDNN.LF interactions suggest stronger associations between HRV and WML in advancing age, specific to the low-frequency band, highlighting SDNN.LF (LF-HRV) as a target for further examination of potential early predictors for WML development. These findings additionally illustrate the feasibility of T1w estimations of WML volume even in the absence of T2w or FLAIR data. Future research would benefit from further examination of differences in contribution by low and high frequency HRV, as well as the inclusion of longitudinal data to clarify the lifetime trajectories of HRV and WML development within participants.

## 4.5 Acknowledgments

This study was supported through grant funding from the Canadian Institutes of Health Research (CIHR) (#PJT169688), as well as the Canada Research Chairs (CRC) program (JJC). Special thanks are given to Ahmad Hussein for assistance with the heart-rate variability analysis.

## SUPPLEMENTAL TABLES

**Supplemental Table 1:** Exploratory comparison of contributions of normalized and demeaned HRV to WML volume across all participants All models are linear regressions with overall F-statistics, *p*-values, and effect sizes (*R^2^_adjusted_*) listed. Slopes are noted below each regression equation and significant associations bolded with asterisks indicating significance. (*:p<.05, **:p<.01, ***:p<.001)

|  | Model: WML ~ HRV |  |  |  |  |  |
|---|---|---|---|---|---|---|
|  | **T1w+FLAIR** |  | **T1w+T2w** |  | **T1w** |  |
|  | *Estimate* | *p-value* | *Estimate* | *p-value* | *Estimate* | *p-value* |
|  | F(1, 132) = 8.12, p = 0.005, *R<sup>2</sup><sub>adjusted</sub>* = 0.05 |  | F(1, 132) = 5.04, p = 0.026, *R<sup>2</sup><sub>adjusted</sub>* = 0.03 |  | F(1, 132) = 4.94, p = 0.028, *R<sup>2</sup><sub>adjusted</sub>* = 0.03 |  |
| **RMSSD** | **-4.81E-06** | **0.005\*\*** | **-4.36E-06** | **0.026\*** | **-3.55E-06** | **0.028\*** |
|  | F(1, 132) = 7.46, p = 0.007, *R<sup>2</sup><sub>adjusted</sub>* = 0.05 |  | F(1, 132) = 6.29, p = 0.013, *R<sup>2</sup><sub>adjusted</sub>* = 0.04 |  | F(1, 132) = 6.26, p = 0.013, *R<sup>2</sup><sub>adjusted</sub>* = 0.04 |  |
| **SDNN.LF (log)** | **-4.85E-04** | **0.007\*\*** | **-5.09E-04** | **0.013\*\*** | **-4.21E-04** | **0.013\*\*** |
|  | F(1, 132) = 15.30, p < 0.001, *R<sup>2</sup><sub>adjusted</sub>* = 0.10 |  | F(1, 132) = 15.03, p < 0.001, *R<sup>2</sup><sub>adjusted</sub>* = 0.10 |  | F(1, 132) = 13.62, p < 0.001, *R<sup>2</sup><sub>adjusted</sub>* = 0.09 |  |
| **SDNN.HF (log)** | **-7.00E-04** | **<0.001\*\*\*** | **-7.90E-04** | **<0.001\*\*\*** | **-6.21E-04** | **<0.001\*\*\*** |

**Supplemental Table 2:**
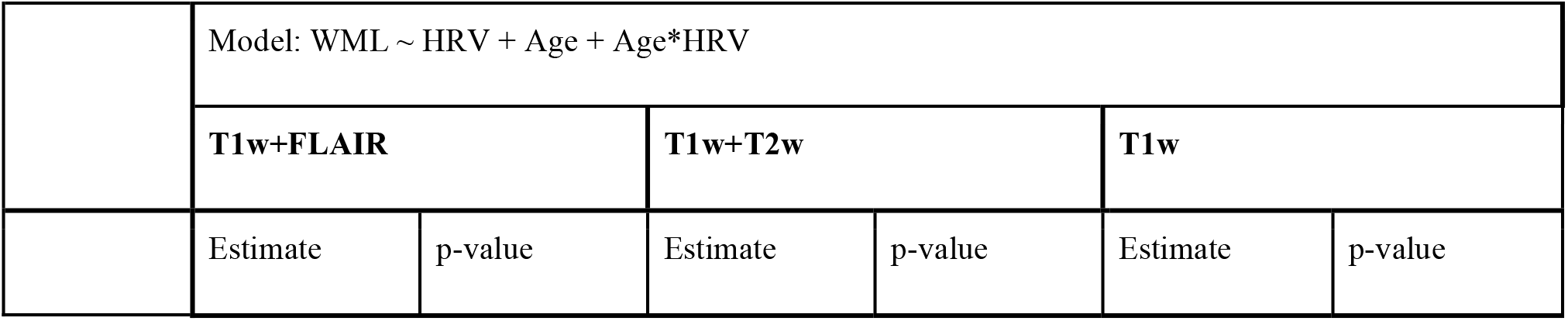
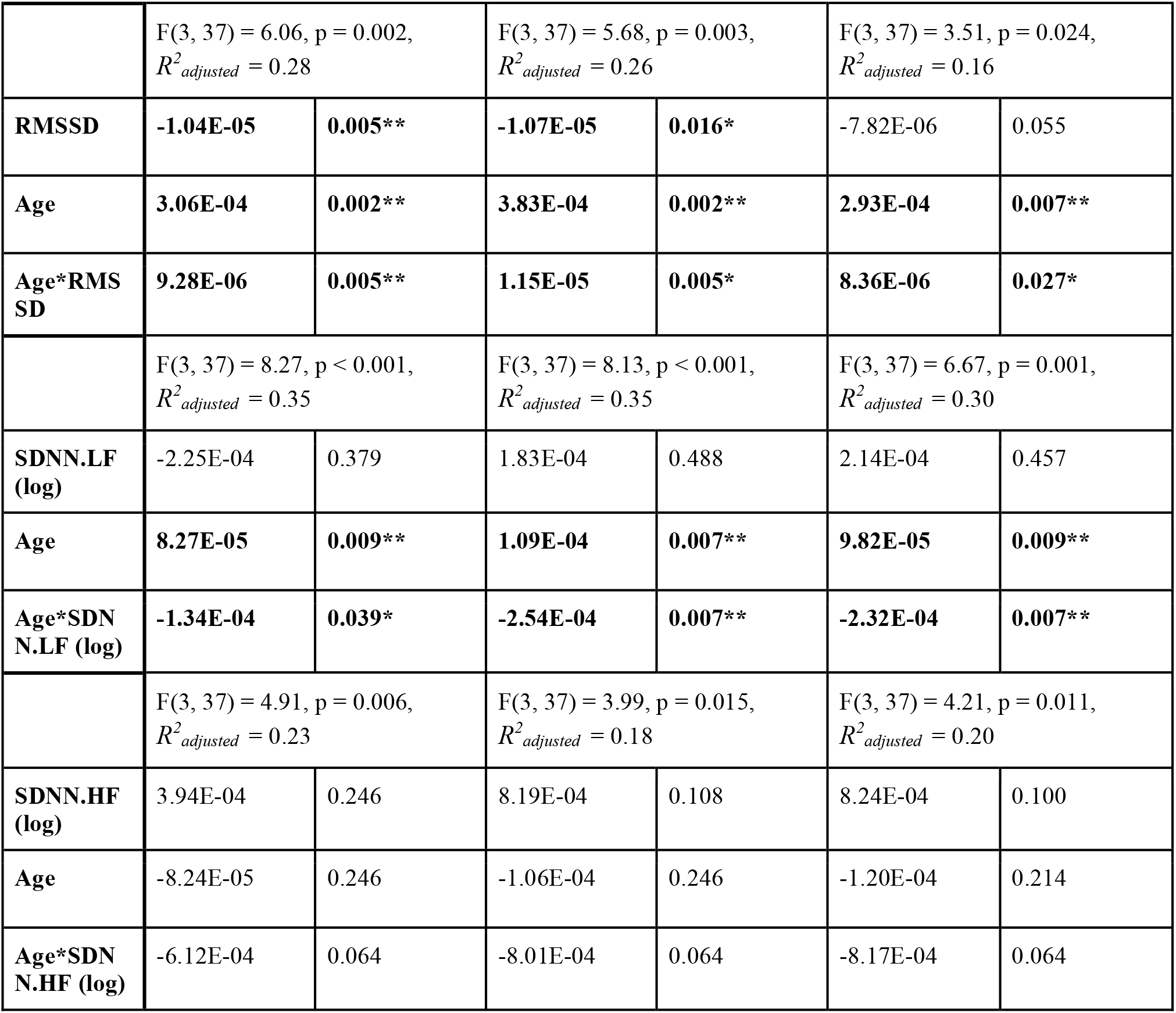
Exploratory model in female participants alone: Contributions of normalized and demeaned HRV, Age, and the interaction of Age*HRV to WML volume All models are linear regressions with overall F-statistics, *p*-values, and effect sizes (*R^2^_adjusted_*) listed. Slopes and FDR-corrected p-values by coefficient are noted below each regression equation and significant associations bolded with asterisks indicating significance following FDR correction by measure. (*:p<.05, **:p<.01, ***:p<.001)

|  | Model: WML ~ HRV + Age + Age*HRV |  |  |  |  |  |
|---|---|---|---|---|---|---|
|  | **T1w+FLAIR** |  | **T1w+T2w** |  | **T1w** |  |
|  | Estimate | p-value | Estimate | p-value | Estimate | p-value |
|  | F(3, 37) = 6.06, p = 0.002, *R<sup>2</sup><sub>adjusted</sub>* = 0.28 |  | F(3, 37) = 5.68, p = 0.003, *R<sup>2</sup><sub>adjusted</sub>* = 0.26 |  | F(3, 37) = 3.51, p = 0.024, *R<sup>2</sup><sub>adjusted</sub>* = 0.16 |  |
| **RMSSD** | **-1.04E-05** | **0.005\*\*** | **-1.07E-05** | **0.016\*** | -7.82E-06 | 0.055 |
| **Age** | **3.06E-04** | **0.002\*\*** | **3.83E-04** | **0.002\*\*** | **2.93E-04** | **0.007\*\*** |
| **Age\*RMSSD** | **9.28E-06** | **0.005\*\*** | **1.15E-05** | **0.005\*** | **8.36E-06** | **0.027\*** |
|  | F(3, 37) = 8.27, p < 0.001, *R<sup>2</sup><sub>adjusted</sub>* = 0.35 |  | F(3, 37) = 8.13, p < 0.001, *R<sup>2</sup><sub>adjusted</sub>* = 0.35 |  | F(3, 37) = 6.67, p = 0.001, *R<sup>2</sup><sub>adjusted</sub>* = 0.30 |  |
| **SDNN.LF (log)** | -2.25E-04 | 0.379 | 1.83E-04 | 0.488 | 2.14E-04 | 0.457 |
| **Age** | **8.27E-05** | **0.009\*\*** | **1.09E-04** | **0.007\*\*** | **9.82E-05** | **0.009\*\*** |
| **Age\*SDNN.LF (log)** | **-1.34E-04** | **0.039\*** | **-2.54E-04** | **0.007\*\*** | **-2.32E-04** | **0.007\*\*** |
|  | F(3, 37) = 4.91, p = 0.006, *R<sup>2</sup><sub>adjusted</sub>* = 0.23 |  | F(3, 37) = 3.99, p = 0.015, *R<sup>2</sup><sub>adjusted</sub>* = 0.18 |  | F(3, 37) = 4.21, p = 0.011, *R<sup>2</sup><sub>adjusted</sub>* = 0.20 |  |
| **SDNN.HF (log)** | 3.94E-04 | 0.246 | 8.19E-04 | 0.108 | 8.24E-04 | 0.100 |
| **Age** | -8.24E-05 | 0.246 | -1.06E-04 | 0.246 | -1.20E-04 | 0.214 |
| **Age\*SDNN.HF (log)** | -6.12E-04 | 0.064 | -8.01E-04 | 0.064 | -8.17E-04 | 0.064 |

**Supplemental Table 3:**
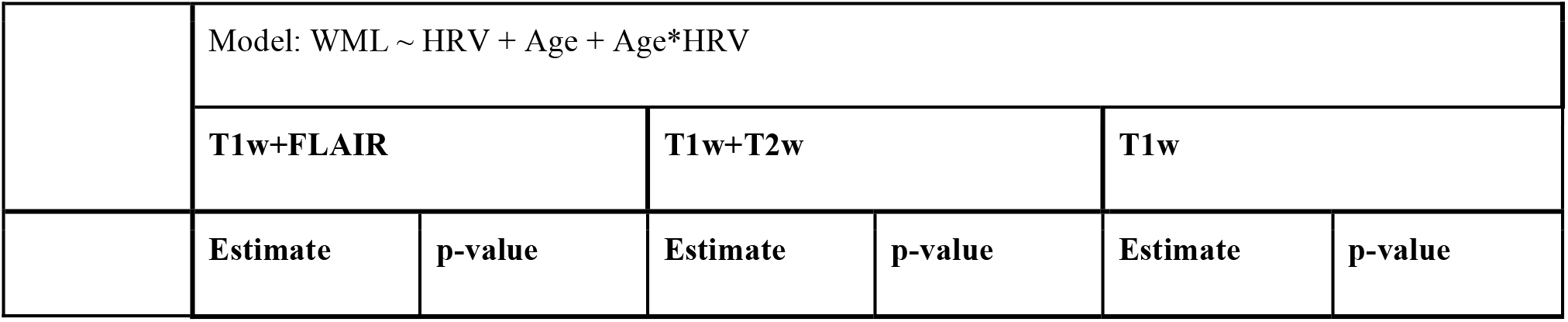
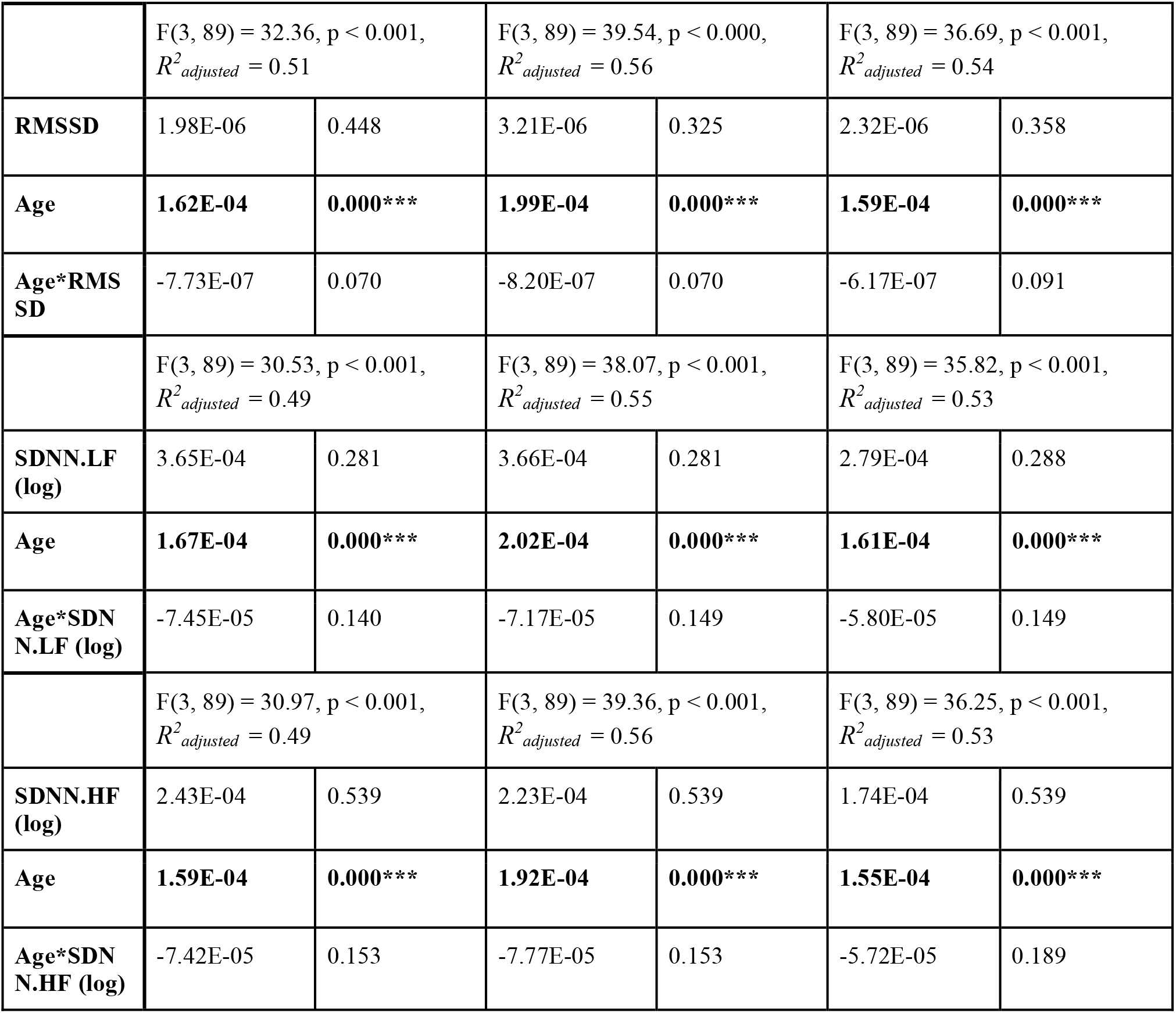
Exploratory model in male participants alone: Contributions of normalized and demeaned HRV, Age, and the interaction of Age*HRV to WML volume All models are linear regressions with overall F-statistics, *p*-values, and effect sizes (*R^2^_adjusted_*) listed. Slopes and FDR-corrected p-values by coefficient are noted below each regression equation and significant associations bolded with asterisks indicating significance following FDR correction by measure. (*:p<.05, **:p<.01, ***:p<.001)

|  | Model: WML ~ HRV + Age + Age*HRV |  |  |  |  |  |
|---|---|---|---|---|---|---|
|  | **T1w+FLAIR** |  | **T1w+T2w** |  | **T1w** |  |
|  | **Estimate** | **p-value** | **Estimate** | **p-value** | **Estimate** | **p-value** |
|  | F(3, 89) = 32.36, p < 0.001,<br>*R*<sup>*2*</sup><sub>*adjusted*</sub> = 0.51 |  | F(3, 89) = 39.54, p < 0.000,<br>*R*<sup>*2*</sup><sub>*adjusted*</sub> = 0.56 |  | F(3, 89) = 36.69, p < 0.001,<br>*R*<sup>*2*</sup><sub>*adjusted*</sub> = 0.54 |  |
| **RMSSD** | 1.98E-06 | 0.448 | 3.21E-06 | 0.325 | 2.32E-06 | 0.358 |
| **Age** | **1.62E-04** | **0.000\*\*\*** | **1.99E-04** | **0.000\*\*\*** | **1.59E-04** | **0.000\*\*\*** |
| **Age\*RMSSD** | -7.73E-07 | 0.070 | -8.20E-07 | 0.070 | -6.17E-07 | 0.091 |
|  | F(3, 89) = 30.53, p < 0.001,<br>*R*<sup>*2*</sup><sub>*adjusted*</sub> = 0.49 |  | F(3, 89) = 38.07, p < 0.001,<br>*R*<sup>*2*</sup><sub>*adjusted*</sub> = 0.55 |  | F(3, 89) = 35.82, p < 0.001,<br>*R*<sup>*2*</sup><sub>*adjusted*</sub> = 0.53 |  |
| **SDNN.LF (log)** | 3.65E-04 | 0.281 | 3.66E-04 | 0.281 | 2.79E-04 | 0.288 |
| **Age** | **1.67E-04** | **0.000\*\*\*** | **2.02E-04** | **0.000\*\*\*** | **1.61E-04** | **0.000\*\*\*** |
| **Age\*SDNN.LF (log)** | -7.45E-05 | 0.140 | -7.17E-05 | 0.149 | -5.80E-05 | 0.149 |
|  | F(3, 89) = 30.97, p < 0.001,<br>*R*<sup>*2*</sup><sub>*adjusted*</sub> = 0.49 |  | F(3, 89) = 39.36, p < 0.001,<br>*R*<sup>*2*</sup><sub>*adjusted*</sub> = 0.56 |  | F(3, 89) = 36.25, p < 0.001,<br>*R*<sup>*2*</sup><sub>*adjusted*</sub> = 0.53 |  |
| **SDNN.HF (log)** | 2.43E-04 | 0.539 | 2.23E-04 | 0.539 | 1.74E-04 | 0.539 |
| **Age** | **1.59E-04** | **0.000\*\*\*** | **1.92E-04** | **0.000\*\*\*** | **1.55E-04** | **0.000\*\*\*** |
| **Age\*SDNN.HF (log)** | -7.42E-05 | 0.153 | -7.77E-05 | 0.153 | -5.72E-05 | 0.189 |

**Supplemental Table 4:** Exploratory comparisons of contributions of normalized and demeaned HRV to periventricular WML volume across all participants All models are linear regressions with overall F-statistics, *p*-values, and effect sizes (*R^2^_adjusted_*) listed. Slopes are noted below each regression equation and significant associations bolded with asterisks indicating significance. (*:p<.05, **:p<.01, ***:p<.001)

|  | Model: WML ~ HRV |  |  |  |  |  |
|---|---|---|---|---|---|---|
|  | **T1w+FLAIR** |  | **T1w+T2w** |  | **T1w** |  |
|  | *Estimate* | *p-value* | *Estimate* | *p-value* | *Estimate* | *p-value* |
|  | F(1, 132) = 7.55, p = 0.007 , *R<sup>2</sup><sub>adjusted</sub>* = .05 |  | F(1, 132) = 4.15, p = 0.044, *R<sup>2</sup><sub>adjusted</sub>* = .02 |  | F(1, 132) = 4.41, p = 0.038, *R<sup>2</sup><sub>adjusted</sub>* = .03 |  |
| **RMSSD** | **-4.196e-06** | **0.007\*\*** | **-3.537e-06** | **0.044\*** | **-3.143e-06** | **0.038\*** |
|  | F(1, 132) = 6.48, p = 0.012, *R<sup>2</sup><sub>adjusted</sub>* = .04 |  | F(1, 132) = 5.53, p = 0.020, *R<sup>2</sup><sub>adjusted</sub>* = .03 |  | F(1, 132) = 5.87, p = 0.017, *R<sup>2</sup><sub>adjusted</sub>* = .04 |  |
| **SDNN.LF (log)** | **-4.100e-04** | **0.012\*** | **-4.268e-04** | **0.020\*** | **5.873** | **0.017\*** |
|  | F(1, 132) = 14.41, p < 0.001, *R<sup>2</sup><sub>adjusted</sub>* = .09 |  | F(1, 132) = 13.68, p < 0.001, *R<sup>2</sup><sub>adjusted</sub>* = .09 |  | F(1, 132) = 13.41, p < 0.001, *R<sup>2</sup><sub>adjusted</sub>* = .09 |  |
| **SDNN.HF (log)** | **-6.152e-04** | **<0.001\*\*\*** | **-6.749e-04** | **<0.001\*\*\*** | **-5.768e-04** | **<0.001\*\*\*** |

**Supplemental Table 5:**
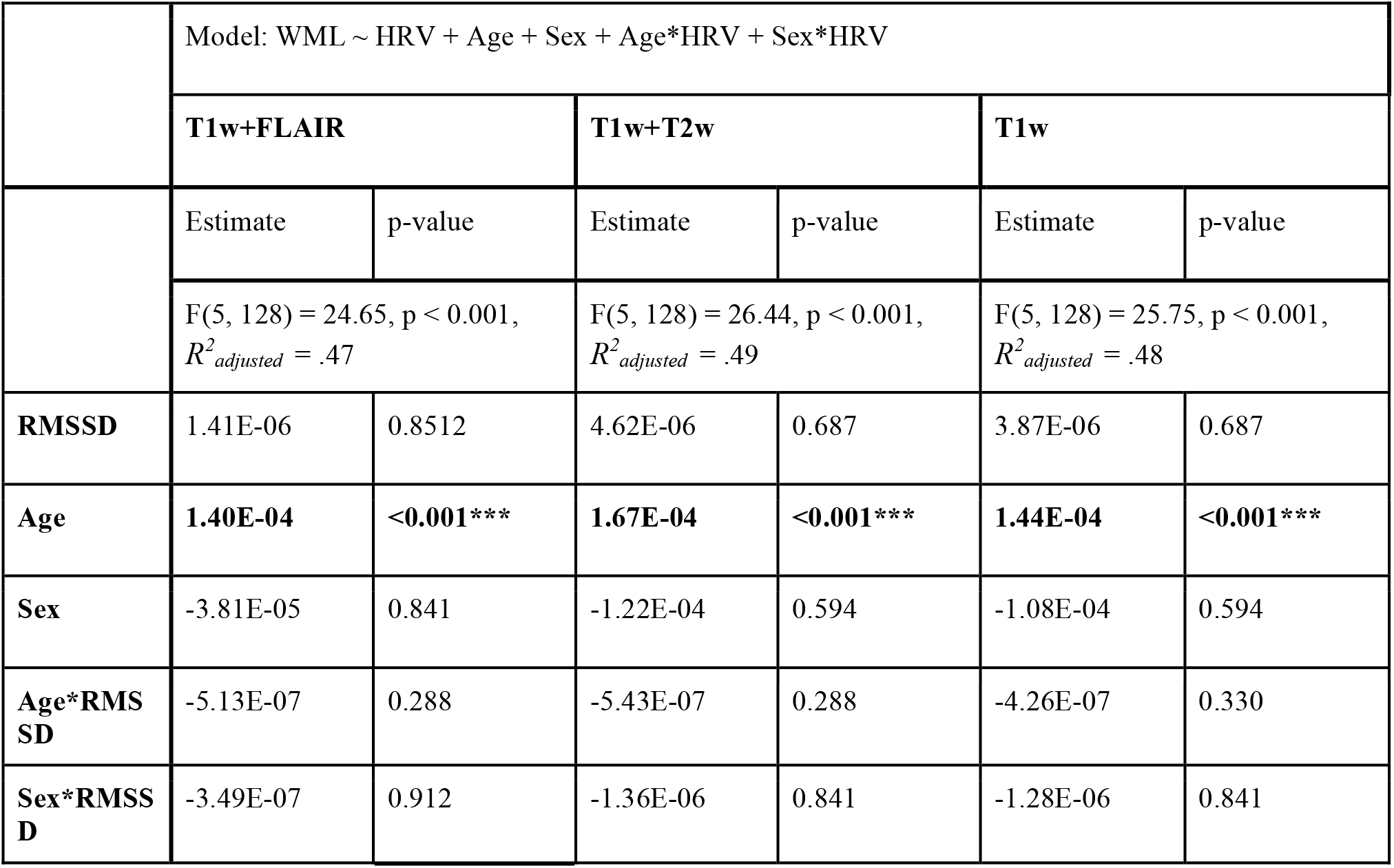
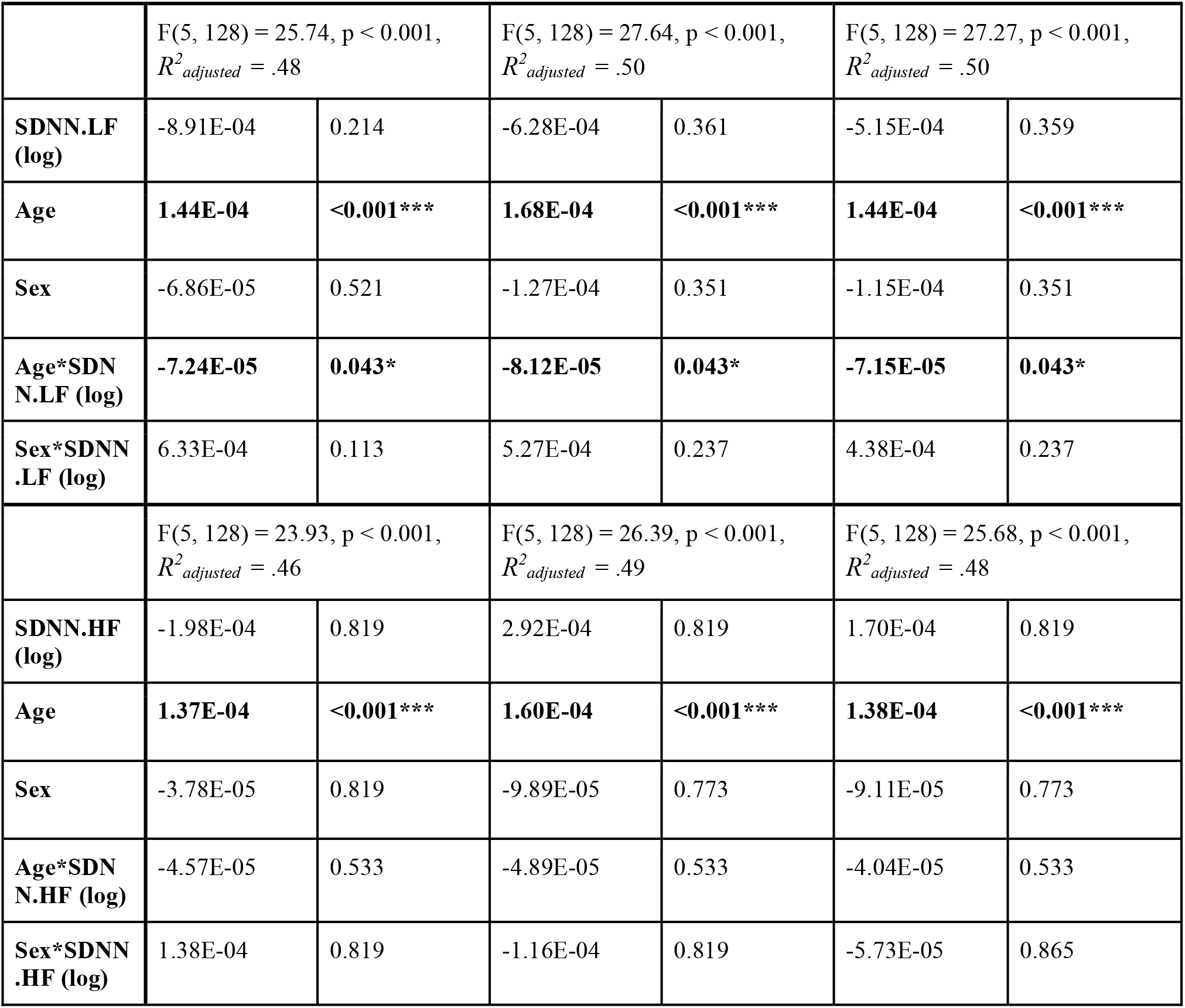
Model across all participants: Contributions of normalized and demeaned HRV, Age, Sex, and the interactions of Age*HRV and Sex*HRV to periventricular WML volume All models are linear regressions with overall F-statistics, *p*-values, and effect sizes (*R^2^_adjusted_*) listed. Slopes and FDR-corrected p-values by coefficient are noted below each regression equation and significant associations bolded with asterisks indicating significance following FDR correction by measure. (*:p<.05, **:p<.01, ***:p<.001)

|  | Model: WML ~ HRV + Age + Sex + Age*HRV + Sex*HRV |  |  |  |  |  |
|---|---|---|---|---|---|---|
|  | **T1w+FLAIR** |  | **T1w+T2w** |  | **T1w** |  |
|  | Estimate | p-value | Estimate | p-value | Estimate | p-value |
|  | F(5, 128) = 24.65, p < 0.001, *R*<sup>2</sup><sub>*adjusted*</sub> = .47 |  | F(5, 128) = 26.44, p < 0.001, *R*<sup>2</sup><sub>*adjusted*</sub> = .49 |  | F(5, 128) = 25.75, p < 0.001, *R*<sup>2</sup><sub>*adjusted*</sub> = .48 |  |
| **RMSSD** | 1.41E-06 | 0.8512 | 4.62E-06 | 0.687 | 3.87E-06 | 0.687 |
| **Age** | **1.40E-04** | **<0.001\*\*\*** | **1.67E-04** | **<0.001\*\*\*** | **1.44E-04** | **<0.001\*\*\*** |
| **Sex** | -3.81E-05 | 0.841 | -1.22E-04 | 0.594 | -1.08E-04 | 0.594 |
| **Age\*RMSSD** | -5.13E-07 | 0.288 | -5.43E-07 | 0.288 | -4.26E-07 | 0.330 |
| **Sex\*RMSSD** | -3.49E-07 | 0.912 | -1.36E-06 | 0.841 | -1.28E-06 | 0.841 |
|  | F(5, 128) = 25.74, p < 0.001, *R*<sup>2</sup><sub>*adjusted*</sub> = .48 |  | F(5, 128) = 27.64, p < 0.001, *R*<sup>2</sup><sub>*adjusted*</sub> = .50 |  | F(5, 128) = 27.27, p < 0.001, *R*<sup>2</sup><sub>*adjusted*</sub> = .50 |  |
| **SDNN.LF (log)** | -8.91E-04 | 0.214 | -6.28E-04 | 0.361 | -5.15E-04 | 0.359 |
| **Age** | **1.44E-04** | **<0.001\*\*\*** | **1.68E-04** | **<0.001\*\*\*** | **1.44E-04** | **<0.001\*\*\*** |
| **Sex** | -6.86E-05 | 0.521 | -1.27E-04 | 0.351 | -1.15E-04 | 0.351 |
| **Age\*SDNN.LF (log)** | **-7.24E-05** | **0.043\*** | **-8.12E-05** | **0.043\*** | **-7.15E-05** | **0.043\*** |
| **Sex\*SDNN.LF (log)** | 6.33E-04 | 0.113 | 5.27E-04 | 0.237 | 4.38E-04 | 0.237 |
|  | F(5, 128) = 23.93, p < 0.001, *R*<sup>2</sup><sub>*adjusted*</sub> = .46 |  | F(5, 128) = 26.39, p < 0.001, *R*<sup>2</sup><sub>*adjusted*</sub> = .49 |  | F(5, 128) = 25.68, p < 0.001, *R*<sup>2</sup><sub>*adjusted*</sub> = .48 |  |
| **SDNN.HF (log)** | -1.98E-04 | 0.819 | 2.92E-04 | 0.819 | 1.70E-04 | 0.819 |
| **Age** | **1.37E-04** | **<0.001\*\*\*** | **1.60E-04** | **<0.001\*\*\*** | **1.38E-04** | **<0.001\*\*\*** |
| **Sex** | -3.78E-05 | 0.819 | -9.89E-05 | 0.773 | -9.11E-05 | 0.773 |
| **Age\*SDNN.HF (log)** | -4.57E-05 | 0.533 | -4.89E-05 | 0.533 | -4.04E-05 | 0.533 |
| **Sex\*SDNN.HF (log)** | 1.38E-04 | 0.819 | -1.16E-04 | 0.819 | -5.73E-05 | 0.865 |

**Supplemental Table 6:**
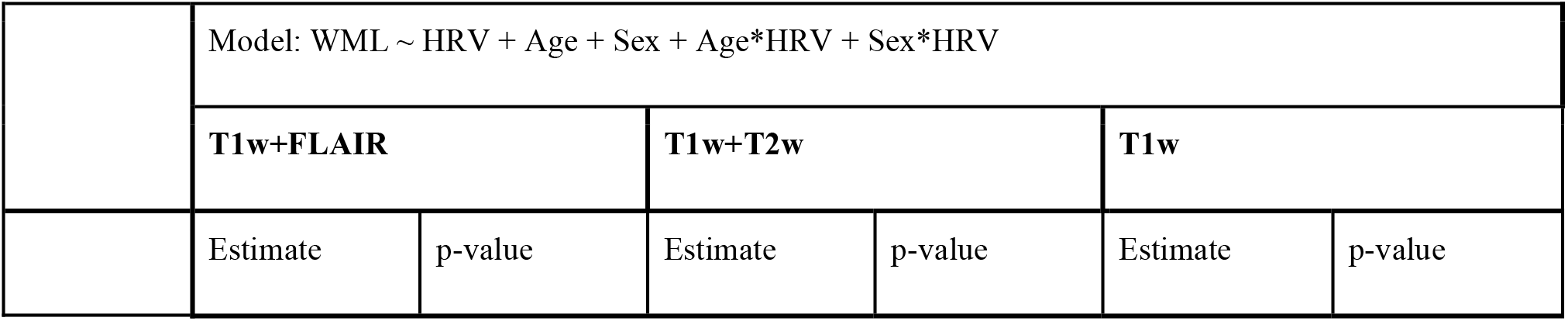
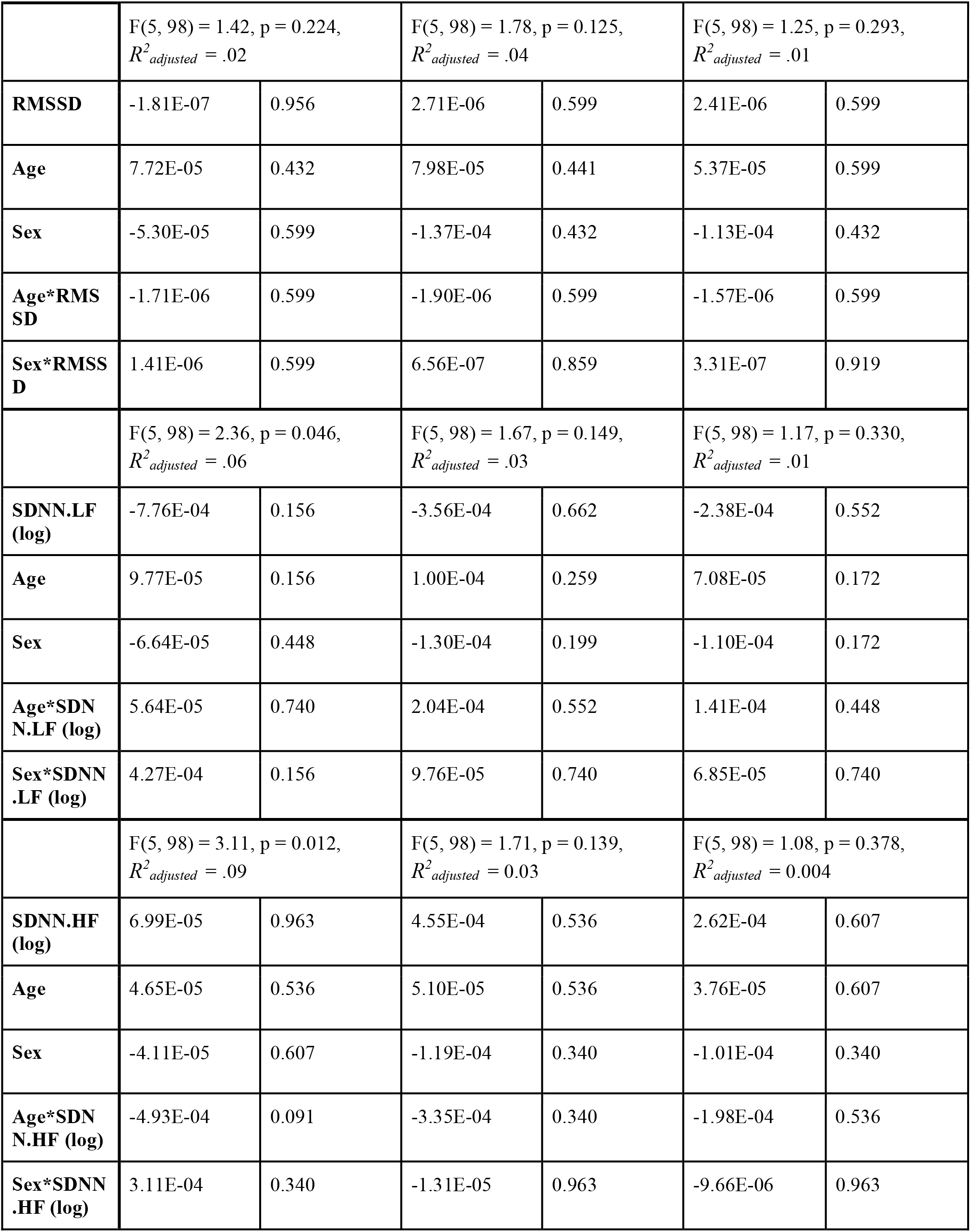
Exploratory model in young participants alone: Contributions of normalized and demeaned HRV, Age, Sex, and the interactions of Age*HRV and Sex*HRV to periventricular WML volume All models are linear regressions with overall F-statistics, *p*-values, and effect sizes (*R^2^_adjusted_*) listed. Slopes and FDR-corrected p-values by coefficient are noted below each regression equation and significant associations bolded with asterisks indicating significance following FDR correction by measure. (*:p<.05, **:p<.01, ***:p<.001)

| Model: WML ~ HRV + Age + Sex + Age*HRV + Sex*HRV |  |  |  |  |  |  |
|---|---|---|---|---|---|---|
|  | **T1w+FLAIR** |  | **T1w+T2w** |  | **T1w** |  |
|  | Estimate | p-value | Estimate | p-value | Estimate | p-value |
|  | F(5, 98) = 1.42, p = 0.224,<br>*R*<sup>*2*</sup><sub>*adjusted*</sub> = .02 |  | F(5, 98) = 1.78, p = 0.125,<br>*R*<sup>*2*</sup><sub>*adjusted*</sub> = .04 |  | F(5, 98) = 1.25, p = 0.293,<br>*R*<sup>*2*</sup><sub>*adjusted*</sub> = .01 |  |
| **RMSSD** | -1.81E-07 | 0.956 | 2.71E-06 | 0.599 | 2.41E-06 | 0.599 |
| **Age** | 7.72E-05 | 0.432 | 7.98E-05 | 0.441 | 5.37E-05 | 0.599 |
| **Sex** | -5.30E-05 | 0.599 | -1.37E-04 | 0.432 | -1.13E-04 | 0.432 |
| **Age\*RMSSD** | -1.71E-06 | 0.599 | -1.90E-06 | 0.599 | -1.57E-06 | 0.599 |
| **Sex\*RMSSD** | 1.41E-06 | 0.599 | 6.56E-07 | 0.859 | 3.31E-07 | 0.919 |
|  | F(5, 98) = 2.36, p = 0.046,<br>*R*<sup>*2*</sup><sub>*adjusted*</sub> = .06 |  | F(5, 98) = 1.67, p = 0.149,<br>*R*<sup>*2*</sup><sub>*adjusted*</sub> = .03 |  | F(5, 98) = 1.17, p = 0.330,<br>*R*<sup>*2*</sup><sub>*adjusted*</sub> = .01 |  |
| **SDNN.LF (log)** | -7.76E-04 | 0.156 | -3.56E-04 | 0.662 | -2.38E-04 | 0.552 |
| **Age** | 9.77E-05 | 0.156 | 1.00E-04 | 0.259 | 7.08E-05 | 0.172 |
| **Sex** | -6.64E-05 | 0.448 | -1.30E-04 | 0.199 | -1.10E-04 | 0.172 |
| **Age\*SDNN.LF (log)** | 5.64E-05 | 0.740 | 2.04E-04 | 0.552 | 1.41E-04 | 0.448 |
| **Sex\*SDNN.LF (log)** | 4.27E-04 | 0.156 | 9.76E-05 | 0.740 | 6.85E-05 | 0.740 |
|  | F(5, 98) = 3.11, p = 0.012,<br>*R*<sup>*2*</sup><sub>*adjusted*</sub> = .09 |  | F(5, 98) = 1.71, p = 0.139,<br>*R*<sup>*2*</sup><sub>*adjusted*</sub> = 0.03 |  | F(5, 98) = 1.08, p = 0.378,<br>*R*<sup>*2*</sup><sub>*adjusted*</sub> = 0.004 |  |
| **SDNN.HF (log)** | 6.99E-05 | 0.963 | 4.55E-04 | 0.536 | 2.62E-04 | 0.607 |
| **Age** | 4.65E-05 | 0.536 | 5.10E-05 | 0.536 | 3.76E-05 | 0.607 |
| **Sex** | -4.11E-05 | 0.607 | -1.19E-04 | 0.340 | -1.01E-04 | 0.340 |
| **Age\*SDNN.HF (log)** | -4.93E-04 | 0.091 | -3.35E-04 | 0.340 | -1.98E-04 | 0.536 |
| **Sex\*SDNN.HF (log)** | 3.11E-04 | 0.340 | -1.31E-05 | 0.963 | -9.66E-06 | 0.963 |

**Supplemental Table 7:** Exploratory model in older participants alone: Contributions of normalized and demeaned HRV, Age, Sex, and the interactions of Age*HRV and Sex*HRV to periventricular WML volume All models are linear regressions with overall F-statistics, *p*-values, and effect sizes (*R^2^_adjusted_*) listed. Slopes and FDR-corrected p-values by coefficient are noted below each regression equation and significant associations bolded with asterisks indicating significance following FDR correction by measure. (*:p<.05, **:p<.01, ***:p<.001)

|  | Model: WML ~ HRV + Age + Sex + Age*HRV + Sex*HRV |  |  |  |  |  |
|---|---|---|---|---|---|---|
|  | **T1w+FLAIR** |  | **T1w+T2w** |  | **T1w** |  |
|  | Estimate | p-value | Estimate | p-value | Estimate | p-value |
|  | F(5, 24) = 1.42, p = 0.231, *R<sup>2</sup><sub>adjusted</sub>* = .08 |  | F(5, 24) = 1.38, p = 0.267, *R<sup>2</sup><sub>adjusted</sub>* = .06 |  | F(5, 24) = 1.20, p = 0.340, *R<sup>2</sup><sub>adjusted</sub>* = .03 |  |
| **RMSSD** | 3.579e-04 | 0.214 | 4.931e-04 | 0.214 | 3.669e-04 | 0.214 |
| **Age** | 5.188e-04 | 0.181 | 5.078e-04 | 0.181 | 4.179e-04 | 0.181 |
| **Sex** | -2.609e-03 | 0.223 | -3.426e-03 | 0.223 | -2.917e-03 | 0.223 |
| **Age\*RMSSD** | 2.902e-06 | 0.804 | 1.093e-07 | 0.804 | 3.567e-06 | 0.804 |
| **Sex\*RMSSD** | -1.964e-04 | 0.181 | -2.494e-04 | 0.181 | -2.034e-04 | 0.181 |
|  | F(5, 24) = 1.09, p = 0.392, *R<sup>2</sup><sub>adjusted</sub>* = .01 |  | F(5, 24) = 1.16, p = 0.360, *R<sup>2</sup><sub>adjusted</sub>* = .03 |  | F(5, 24) = 1.01, p = 0.436, *R<sup>2</sup><sub>adjusted</sub>* = .001 |  |
| **SDNN.LF (log)** | -1.891e-03 | 0.951 | -4.745e-03 | 0.951 | -3.392e-03 | 0.951 |
| **Age** | 4.162e-04 | 0.440 | 4.504e-04 | 0.440 | 3.315e-04 | 0.440 |
| **Sex** | -7.510e-05 | 0.951 | -5.925e-05 | 0.951 | -1.700e-04 | 0.951 |
| **Age\*SDNN.LF (log)** | -2.805e-04 | 0.951 | -1.408e-04 | 0.951 | -1.624e-04 | 0.951 |
| **Sex\*SDNN.LF (log)** | 2.217e-03 | 0.455 | 2.952e-03 | 0.455 | 2.386e-03 | 0.455 |
|  | F(5, 24) = 1.06, p = 0.404, *R<sup>2</sup><sub>adjusted</sub>* = .01 |  | F(5, 24) = 1.22, p = 0.330, *R<sup>2</sup><sub>adjusted</sub>* = 0.04 |  | F(5, 24) = 1.02, p = 0.428, *R<sup>2</sup><sub>adjusted</sub>* = 0.003 |  |
| **SDNN.HF (log)** | -3.283e-02 | 0.214 | -4.369e-02 | 0.214 | -3.636e-02 | 0.214 |
| **Age** | 4.255e-04 | 0.214 | 4.487e-04 | 0.214 | 3.366e-04 | 0.214 |
| **Sex** | 1.509e-03 | 0.215 | 1.839e-03 | 0.215 | 1.430e-03 | 0.215 |
| **Age\*SDNN.HF (log)** | -1.284e-04 | 0.949 | -2.318e-05 | 0.949 | 4.590e-05 | 0.949 |
| **Sex\*SDNN.HF (log)** | 1.691e-02 | 0.214 | 2.179e-02 | 0.214 | 1.781e-02 | 0.214 |

**Supplemental Table 8:**
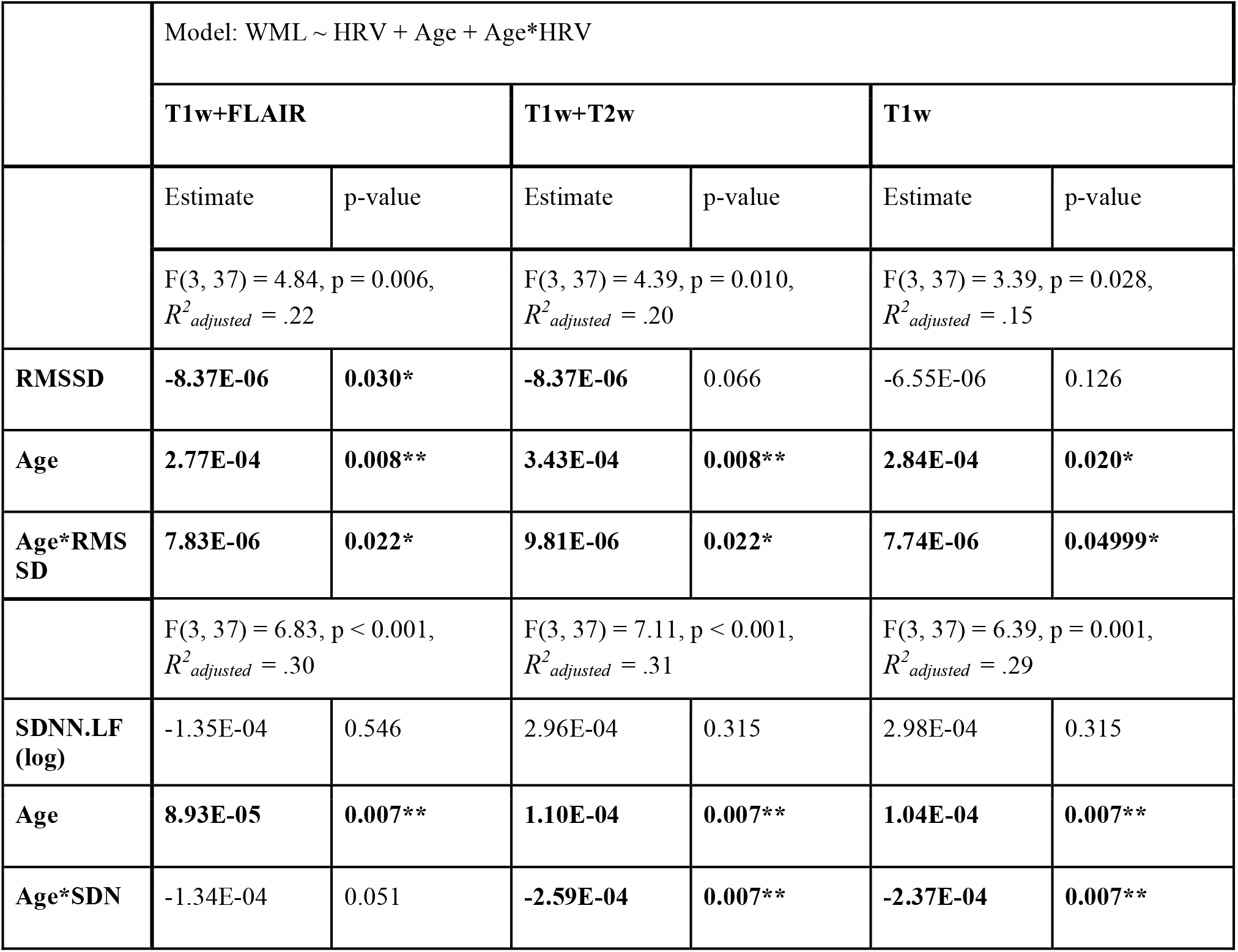
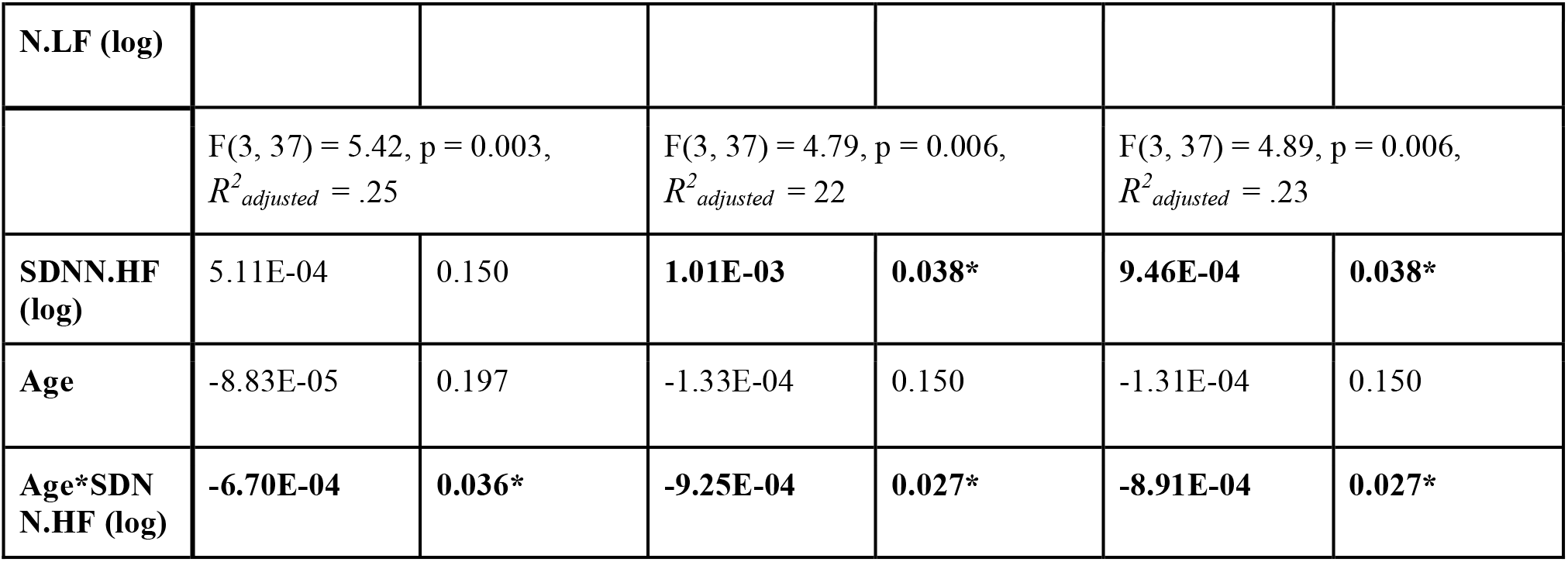
Exploratory model in female participants alone: Contributions of normalized and demeaned HRV, Age, and the interaction of Age*HRV to periventricular WML volume All models are linear regressions with overall F-statistics, *p*-values, and effect sizes (*R^2^_adjusted_*) listed. Slopes and FDR-corrected p-values by coefficient are noted below each regression equation and significant associations bolded with asterisks indicating significance following FDR correction by measure. (*:p<.05, **:p<.01, ***:p<.001)

|  | Model: WML ~ HRV + Age + Age*HRV |  |  |  |  |  |
|---|---|---|---|---|---|---|
|  | **T1w+FLAIR** |  | **T1w+T2w** |  | **T1w** |  |
|  | Estimate | p-value | Estimate | p-value | Estimate | p-value |
|  | F(3, 37) = 4.84, p = 0.006, *R<sup>2</sup><sub>adjusted</sub>* = .22 |  | F(3, 37) = 4.39, p = 0.010, *R<sup>2</sup><sub>adjusted</sub>* = .20 |  | F(3, 37) = 3.39, p = 0.028, *R<sup>2</sup><sub>adjusted</sub>* = .15 |  |
| **RMSSD** | **-8.37E-06** | **0.030\*** | **-8.37E-06** | 0.066 | -6.55E-06 | 0.126 |
| **Age** | **2.77E-04** | **0.008\*\*** | **3.43E-04** | **0.008\*\*** | **2.84E-04** | **0.020\*** |
| **Age\*RMSSD** | **7.83E-06** | **0.022\*** | **9.81E-06** | **0.022\*** | **7.74E-06** | **0.04999\*** |
|  | F(3, 37) = 6.83, p < 0.001, *R<sup>2</sup><sub>adjusted</sub>* = .30 |  | F(3, 37) = 7.11, p < 0.001, *R<sup>2</sup><sub>adjusted</sub>* = .31 |  | F(3, 37) = 6.39, p = 0.001, *R<sup>2</sup><sub>adjusted</sub>* = .29 |  |
| **SDNN.LF (log)** | -1.35E-04 | 0.546 | 2.96E-04 | 0.315 | 2.98E-04 | 0.315 |
| **Age** | **8.93E-05** | **0.007\*\*** | **1.10E-04** | **0.007\*\*** | **1.04E-04** | **0.007\*\*** |
| **Age\*SDNN.LF (log)** | -1.34E-04 | 0.051 | **-2.59E-04** | **0.007\*\*** | **-2.37E-04** | **0.007\*\*** |
|  | F(3, 37) = 5.42, p = 0.003, *R<sup>2</sup><sub>adjusted</sub>* = .25 |  | F(3, 37) = 4.79, p = 0.006, *R<sup>2</sup><sub>adjusted</sub>* = 22 |  | F(3, 37) = 4.89, p = 0.006, *R<sup>2</sup><sub>adjusted</sub>* = .23 |  |
| **SDNN.HF (log)** | 5.11E-04 | 0.150 | **1.01E-03** | **0.038\*** | **9.46E-04** | **0.038\*** |
| **Age** | -8.83E-05 | 0.197 | -1.33E-04 | 0.150 | -1.31E-04 | 0.150 |
| **Age\*SDNN.HF (log)** | **-6.70E-04** | **0.036\*** | **-9.25E-04** | **0.027\*** | **-8.91E-04** | **0.027\*** |

**Supplemental Table 9:** Exploratory model in male participants alone: Contributions of normalized and demeaned HRV, Age, and the interaction of Age*HRV to periventricular WML volume All models are linear regressions with overall F-statistics, *p*-values, and effect sizes (*R^2^_adjusted_*) listed. Slopes and FDR-corrected p-values by coefficient are noted below each regression equation and significant associations bolded with asterisks indicating significance following FDR correction by measure. (*:p<.05, **:p<.01, ***:p<.001)

|  | Model: WML ~ HRV + Age + Age*HRV |  |  |  |  |  |
|---|---|---|---|---|---|---|
|  | **T1w+FLAIR** |  | **T1w+T2w** |  | **T1w** |  |
|  | **Estimate** | **p-value** | **Estimate** | **p-value** | **Estimate** | **p-value** |
|  | F(3, 89) = 32.43, p < 0.001, *R<sup>2</sup><sub>adjusted</sub>* = .51 |  | F(3, 89) = 37.41, p < 0.001, *R<sup>2</sup><sub>adjusted</sub>* = .54 |  | F(3, 89) = 37.50, p < 0.001, *R<sup>2</sup><sub>adjusted</sub>* = .54 |  |
| **RMSSD** | 1.67E-06 | 0.477 | 3.03E-06 | 0.297 | 2.18E-06 | 0.349 |
| **Age** | **1.47E-04** | **<0.001\*\*\*** | **1.75E-04** | **<0.001\*\*\*** | **1.50E-04** | **<0.001\*\*\*** |
| **Age\*RMSSD** | -6.49E-07 | 0.090 | -7.06E-07 | 0.090 | -5.52E-07 | 0.103 |
|  | F(3, 89) = 30.67, p < 0.001, *R<sup>2</sup><sub>adjusted</sub>* = .49 |  | F(3, 89) = 35.96, p < 0.001, *R<sup>2</sup><sub>adjusted</sub>* = .53 |  | F(3, 89) = 36.61, p < 0.001, *R<sup>2</sup><sub>adjusted</sub>* = .54 |  |
| **SDNN.LF (log)** | 3.21E-04 | 0.294 | 2.97E-04 | 0.318 | 2.42E-04 | 0.318 |
| **Age** | **1.51E-04** | **<0.001\*\*\*** | **1.77E-04** | **<0.001\*\*\*** | **1.51E-04** | **<0.001\*\*\*** |
| **Age\*SDNN.LF (log)** | -6.15E-05 | 0.192 | -5.71E-05 | 0.203 | -5.01E-05 | 0.203 |
|  | F(3, 89) = 30.45, p < 0.001, *R<sup>2</sup><sub>adjusted</sub>* = .49 |  | F(3, 89) = 36.47, p < 0.001, *R<sup>2</sup><sub>adjusted</sub>* = .54 |  | F(3, 89) = 36.68, p < 0.001, *R<sup>2</sup><sub>adjusted</sub>* = .54 |  |
| **SDNN.HF (log)** | 1.37E-04 | 0.727 | 1.14E-04 | 0.727 | 9.14E-05 | 0.727 |
| **Age** | **1.45E-04** | **<0.001\*\*\*** | **1.69E-04** | **<0.001\*\*\*** | **1.45E-04** | **<0.001\*\*\*** |
| **Age\*SDNN.HF (log)** | -4.99E-05 | 0.339 | -5.22E-05 | 0.339 | -4.18E-05 | 0.339 |

## SUPPLEMENTAL FIGURES

**Supplemental Figure 1.**
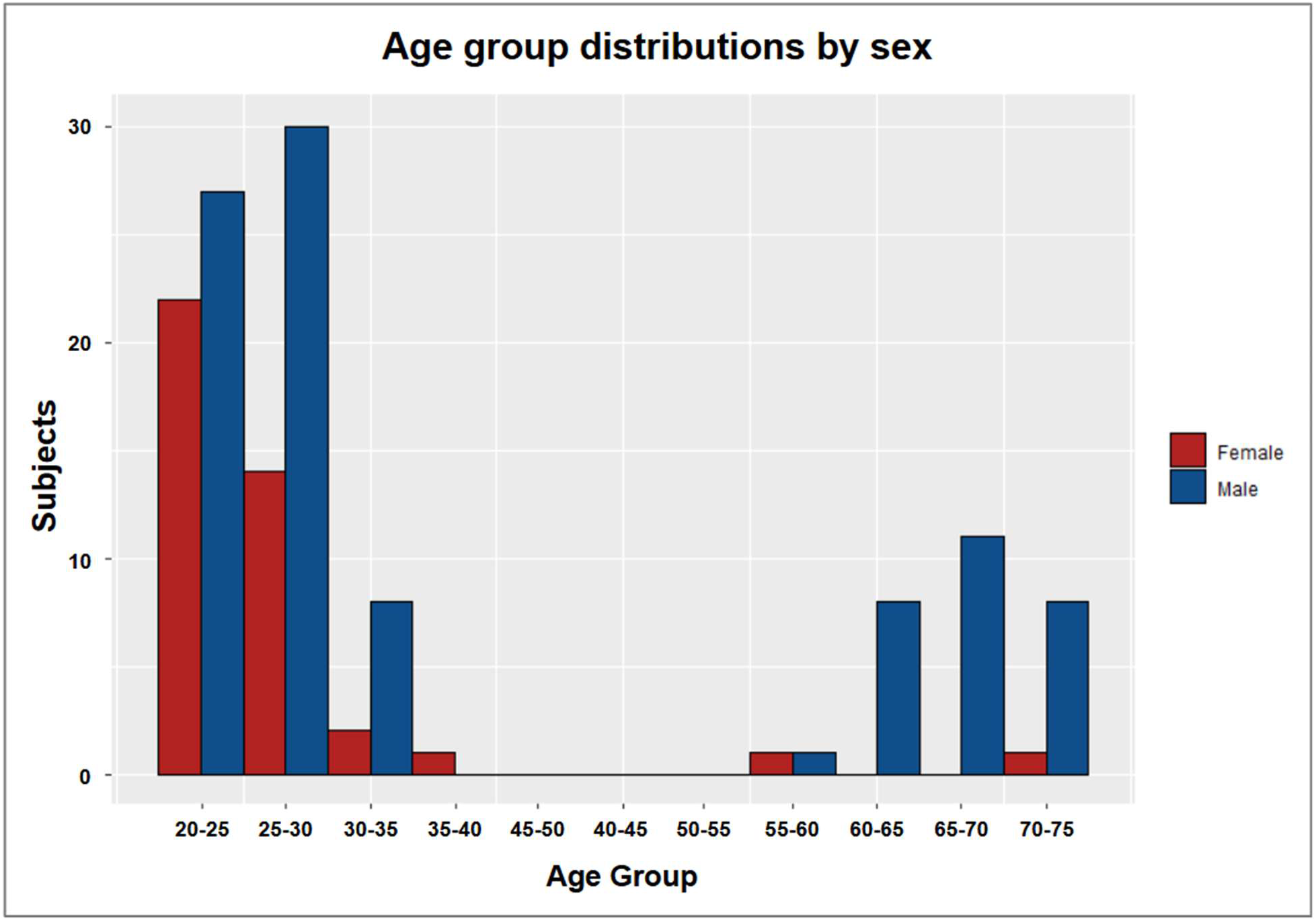
Participant Distributions by Age and Sex. Age group distributions in female (red) and male (blue) subsamples of the Leipzig Study for Mind-Body-Emotion Interactions (LEMON) dataset.

**Supplemental figure 2.**
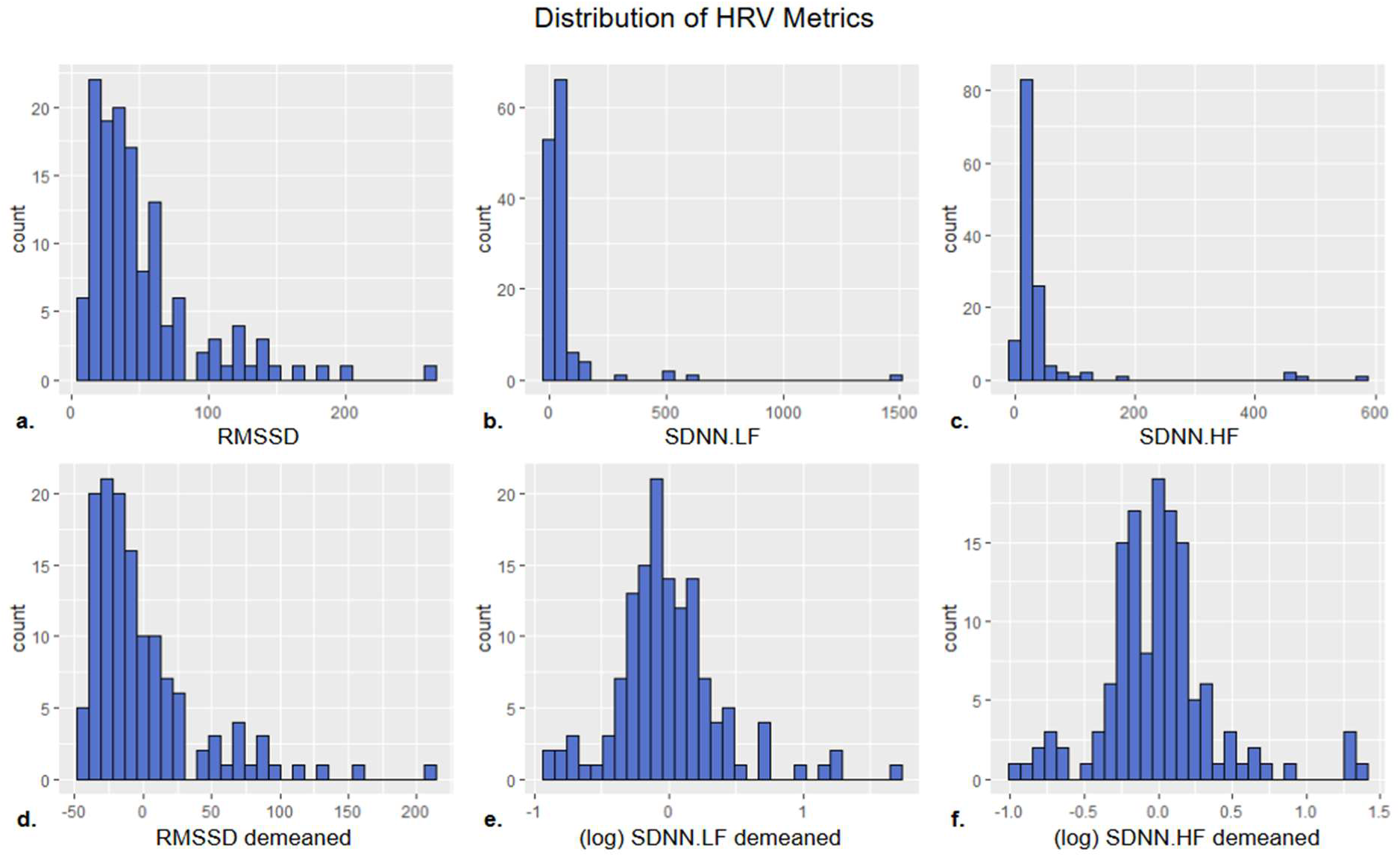
Distribution of HRV Metrics. Distributions of RMSSD before (a) and after (d) demeaning, SDNN.LF before (b) and after (e) log transformation and demeaning, and SDNN.HF before (c) and after (f) log transformation and demeaning, across all participants.

**Supplemental figure 3.**
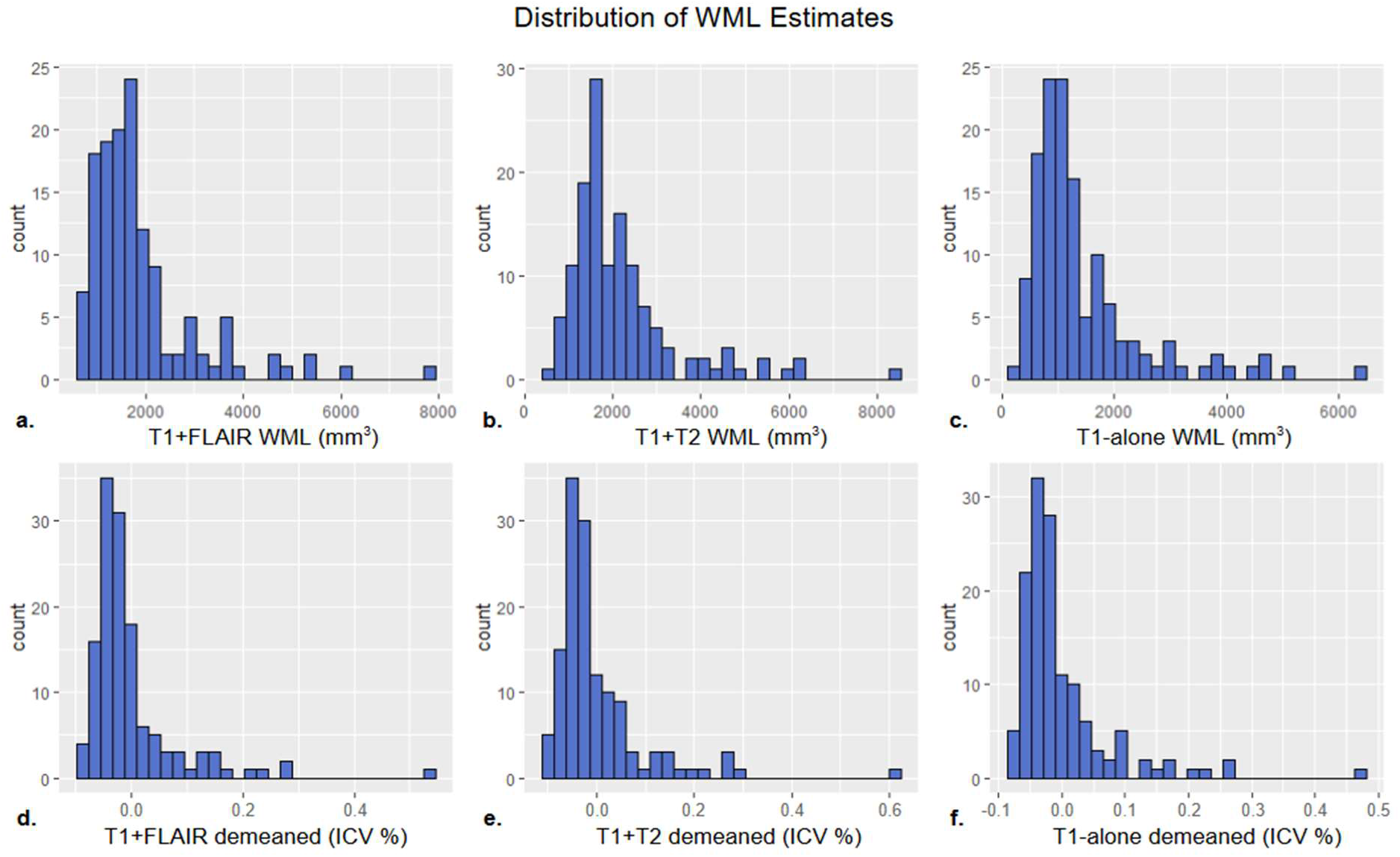
Distribution of WML Estimates. Distributions of white-matter lesion volumes estimated using T1w+FLAIR before (a) and after (d) normalization by intracranial volume (ICV %) and demeaning, T1w+T2w before (b) and after (e) normalization and demeaning and T1w-alone before (c) and after (f) normalization and demeaning, across all participants.

**Supplemental figure 4.**
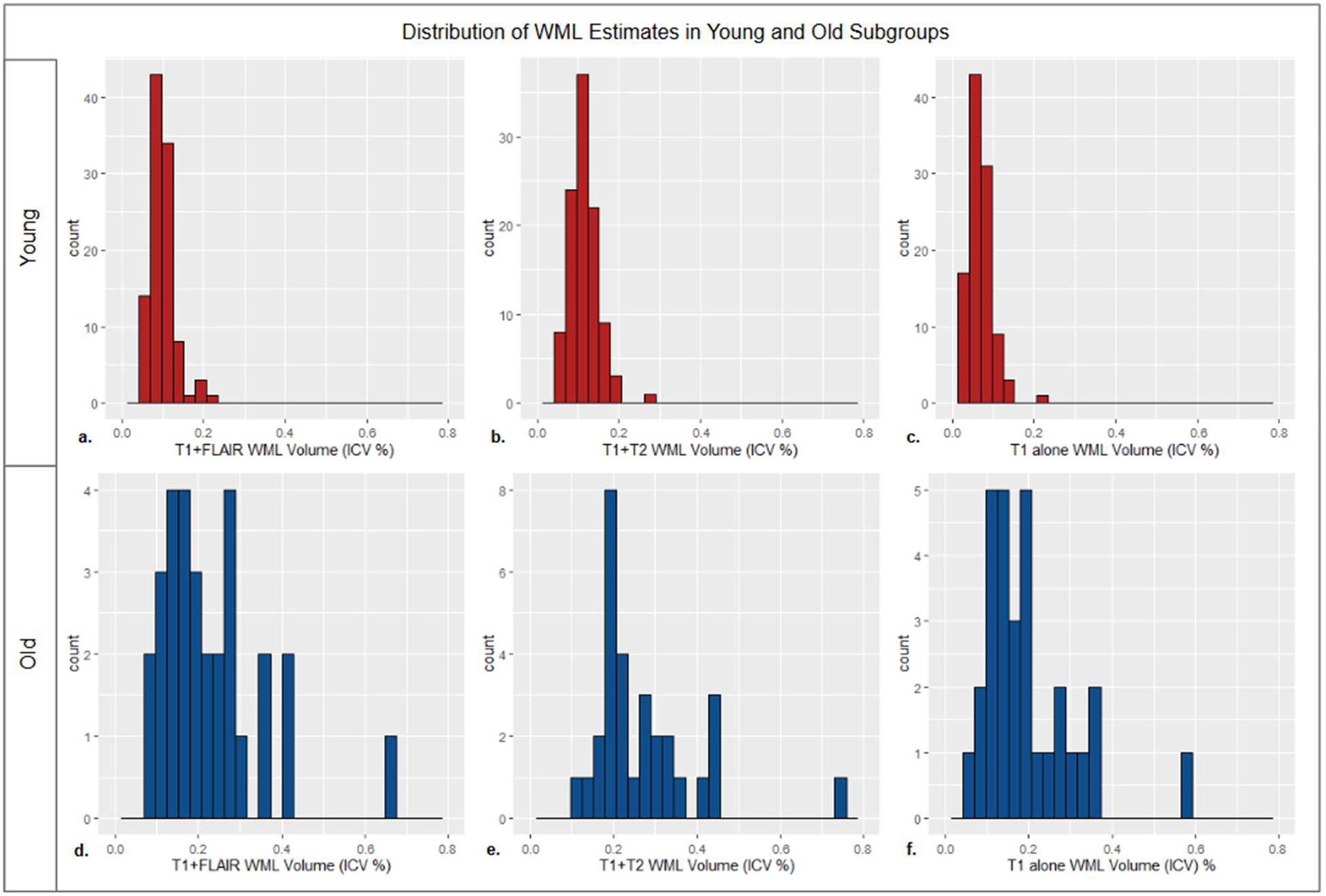
Distribution of WML Estimates by age group. Distributions of intracranial volume normalized white-matter lesion volumes (ICV %) estimated using T1w+FLAIR in young (a) and old (d) subgroups, T1w+T2w in young (b) and old (e) subgroups, and T1w-alone in young (c) and old (f) subgroups.

## SUPPLEMENTAL FILES

***Supplemental file 1:*** Additional model diagnostic tables and figures are available for download from the Harvard Dataverse free data repository: https://dataverse.harvard.edu/previewurl.xhtml?token=afe96855-5ac9-4109-a875-55e7e0be2452

