## Supplemental File 1 for "Association between white-matter lesion volume and heart-rate variability: A comparison of T1w, T1w+T2w and T1w+FLAIR-based segmentations using FreeSurfer SAMSEG"

### 1. Global | All subjects | T1w+FLAIR WML vol | RMSSD | Exploratory unadjusted model

Model: T1w+FLAIR WML vol ~ RMSSD

#### 1.1. Standardized coefficients

| **Term** | **Estimate** | **Std. Error** | **t** | **p** | **β** |
| --- | --- | --- | --- | --- | --- |
| (Intercept) | -1.527e-20 | 7.003e-05 | -2.180e-16 | 1.000 |  |
| RMSSD | -4.809e-06 | 1.688e-06 | -2.85 | .005 | -0.241 |

***Supplemental Table 10.*** *Standardized coefficients values for T1w+FLAIR WML segmentation.*

#### 1.2. 95% confidence intervals

| **Term** | **Lower 95%** | **Upper 95%** |
| --- | --- | --- |
| (Intercept) | -1.385e-04 | 1.385e-04 |
| RMSSD | -8.148e-06 | -1.470e-06 |

***Supplemental Table 11.*** *95% confidence intervals values for T1w+FLAIR WML segmentation.*

#### 1.3. Breusch-Pagan test

| **Test** | **Statistic** | **df** | **p** |
| --- | --- | --- | --- |
| Breusch-Pagan | 2.228 | 1 | .135 |

***Supplemental Table 12.*** *Breusch-Pagan test values for T1w+FLAIR WML segmentation.*

#### 1.4. Normal Q-Q plot

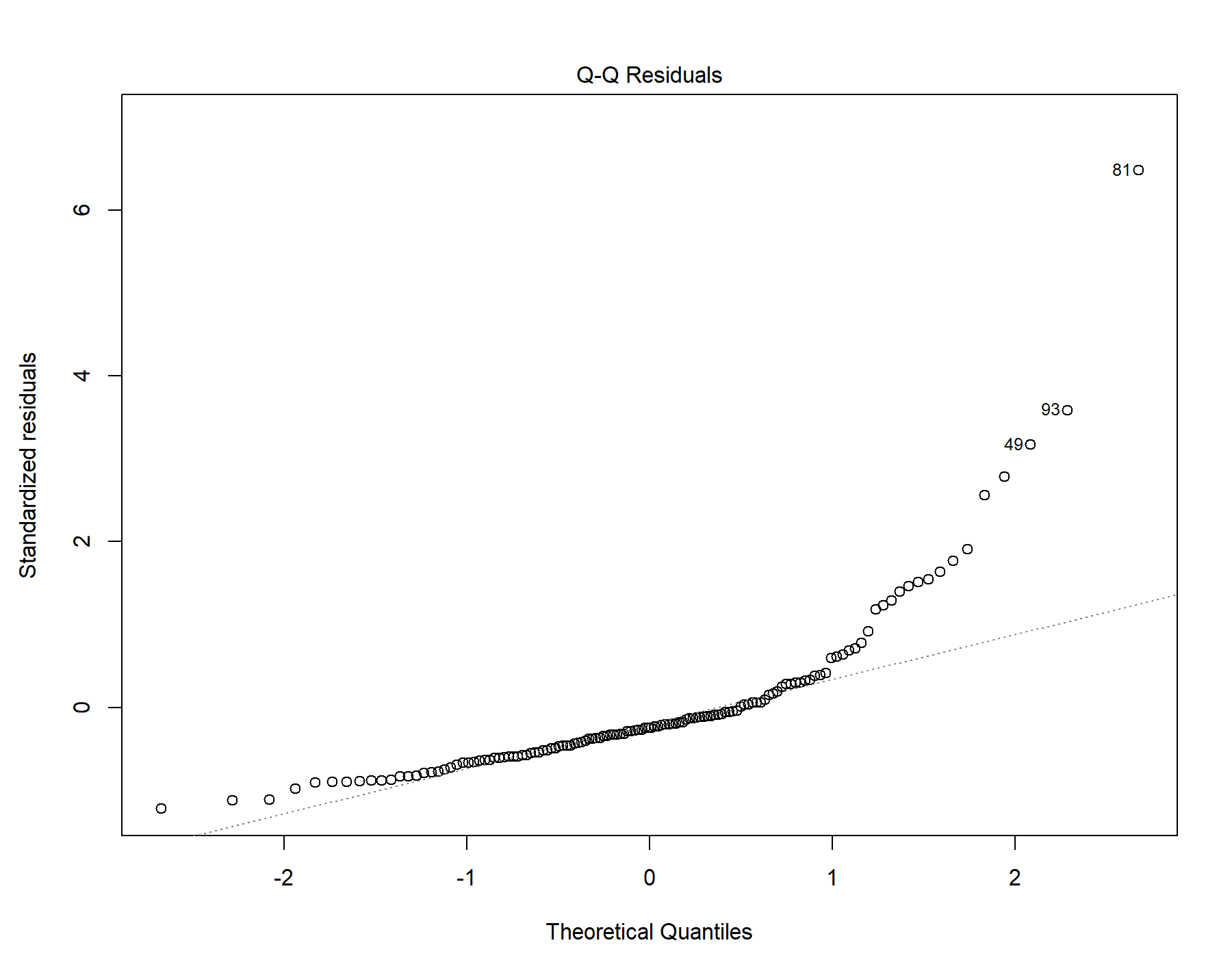

Supplemental Figure 5. Normal distribution quantile-quantile plot for T1w+FLAIR WML segmentation.

#### 1.5. Residuals vs leverage

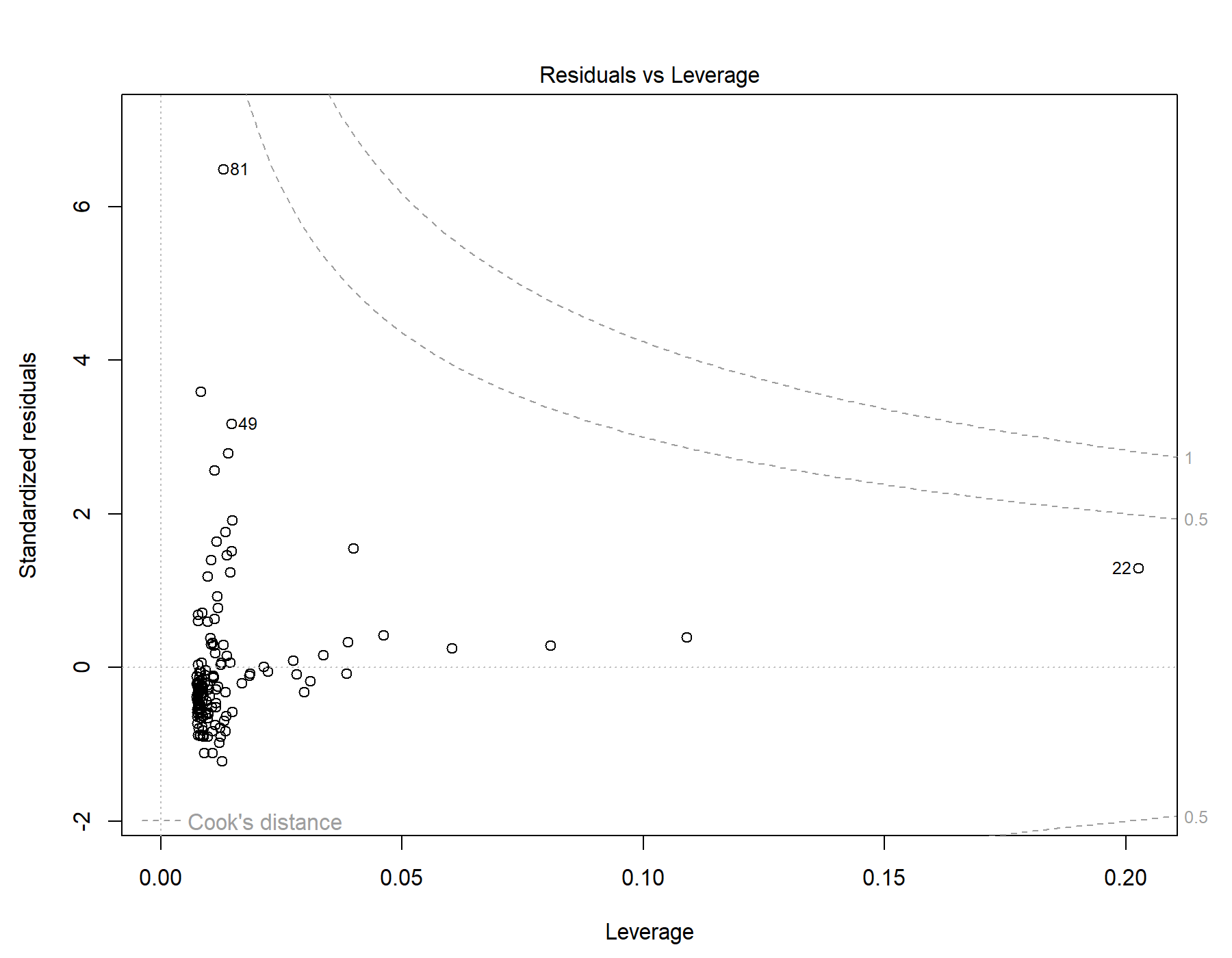

Supplemental Figure 6. Residuals vs leverage diagnostics for T1w+FLAIR WML segmentation.

### 2. Global | All subjects | T1w+T2w WML vol | RMSSD | Exploratory unadjusted model

Model: T1w+T2w WML vol ~ RMSSD

#### 2.1. Standardized coefficients

| **Term** | **Estimate** | **Std. Error** | **t** | **p** | **β** |
| --- | --- | --- | --- | --- | --- |
| (Intercept) | 6.958e-20 | 8.054e-05 | 8.640e-16 | 1.000 |  |
| RMSSD | -4.357e-06 | 1.941e-06 | -2.24 | .026 | -0.192 |

***Supplemental Table 13.*** *Standardized coefficients values for T1w+T2w WML segmentation.*

#### 2.2. 95% confidence intervals

| **Term** | **Lower 95%** | **Upper 95%** |
| --- | --- | --- |
| (Intercept) | -1.593e-04 | 1.593e-04 |
| RMSSD | -8.197e-06 | -5.175e-07 |

***Supplemental Table 14.*** *95% confidence intervals values for T1w+T2w WML segmentation.*

#### 2.3. Breusch-Pagan test

| **Test** | **Statistic** | **df** | **p** |
| --- | --- | --- | --- |
| Breusch-Pagan | 2.109 | 1 | .146 |

***Supplemental Table 15.*** *Breusch-Pagan test values for T1w+T2w WML segmentation.*

#### 2.4. Normal Q-Q plot

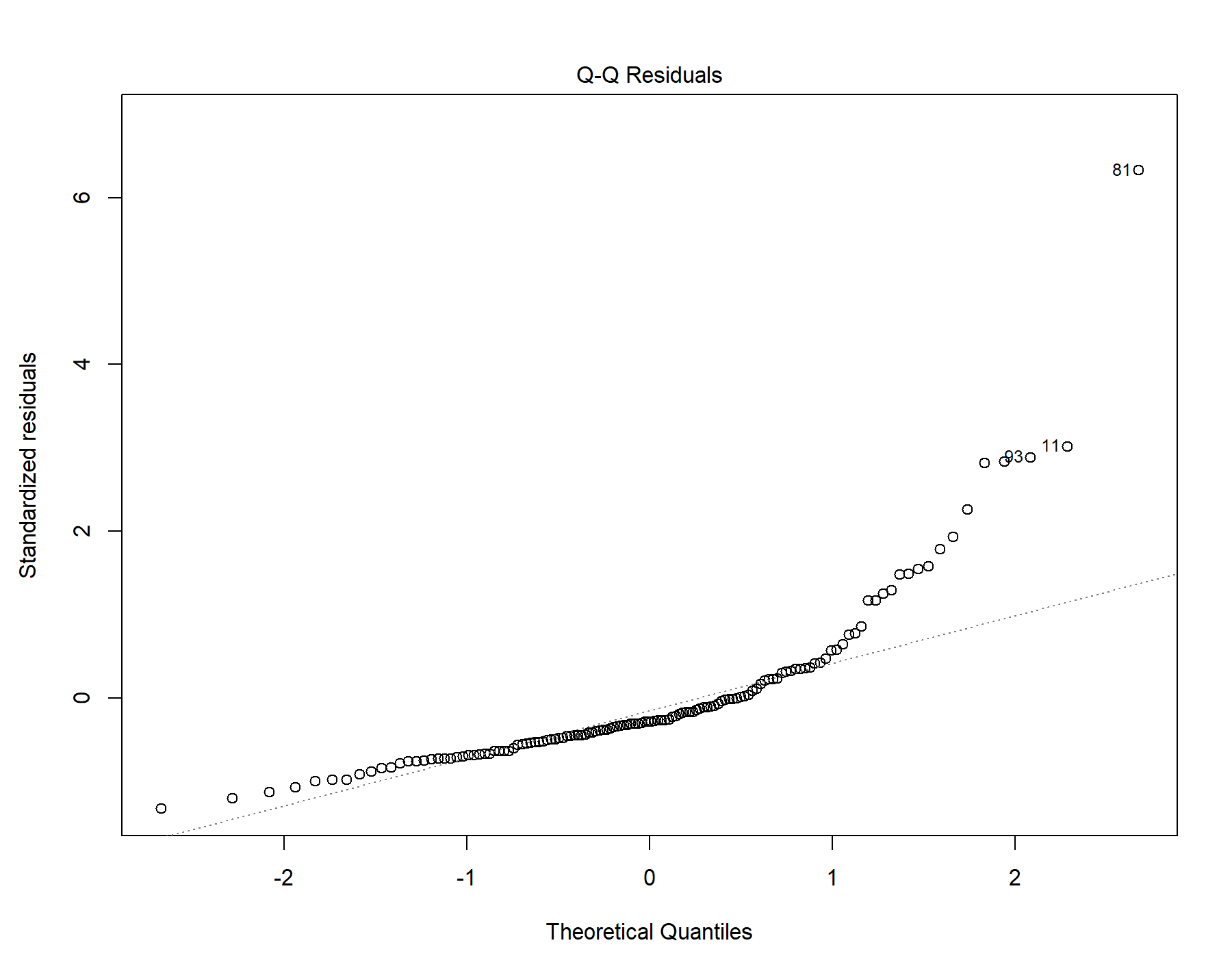

Supplemental Figure 7. Normal distribution quantile-quantile plot for T1w+T2w WML segmentation.

#### 2.5. Residuals vs leverage

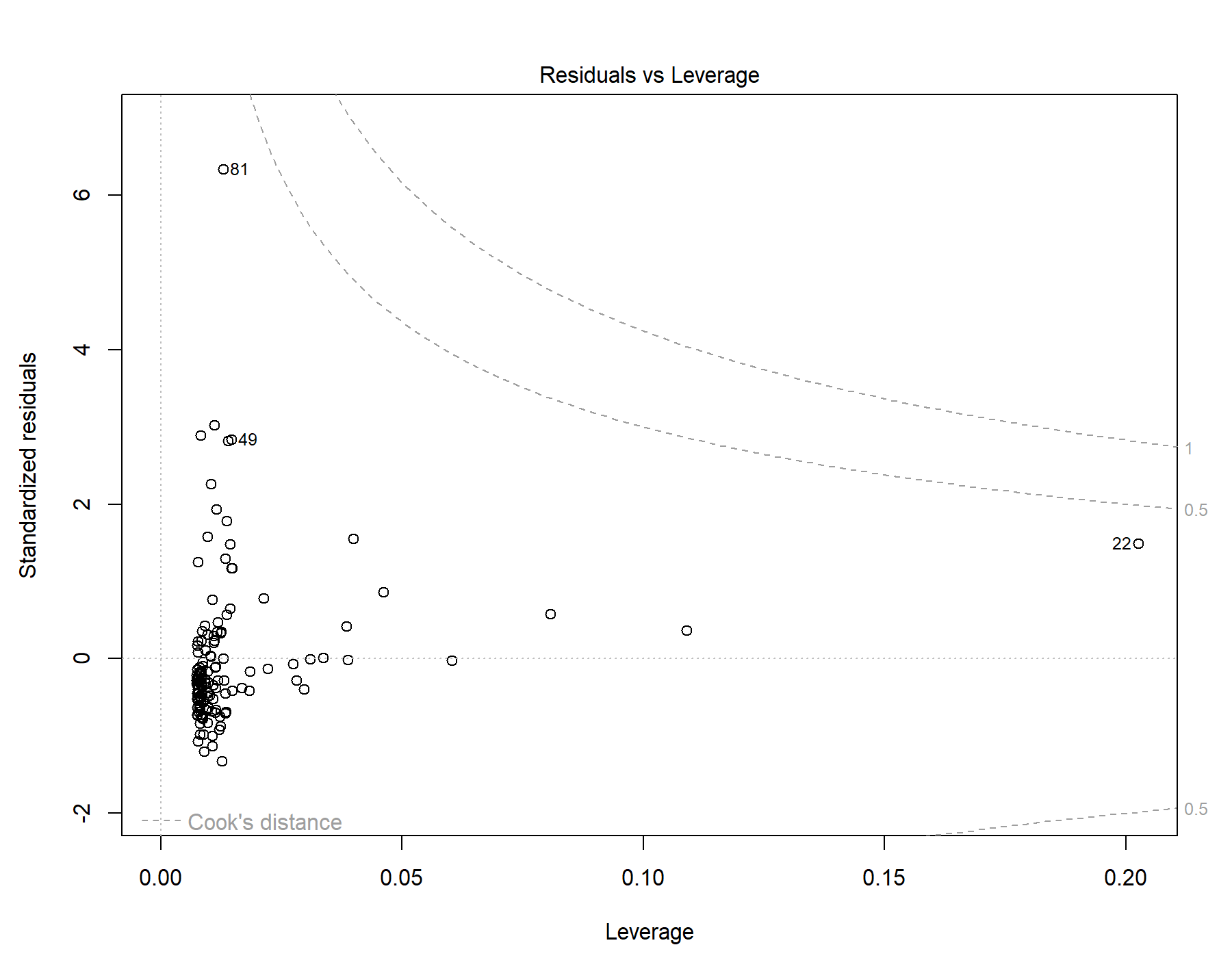

Supplemental Figure 8. Residuals vs leverage diagnostics for T1w+T2w WML segmentation.

### 3. Global | All subjects | T1w-alone WML vol | RMSSD | Exploratory unadjusted model

Model: T1w-alone WML vol ~ RMSSD

#### 3.1. Standardized coefficients

| **Term** | **Estimate** | **Std. Error** | **t** | **p** | **β** |
| --- | --- | --- | --- | --- | --- |
| (Intercept) | 4.247e-20 | 6.623e-05 | 6.413e-16 | 1.000 |  |
| RMSSD | -3.548e-06 | 1.596e-06 | -2.22 | .028 | -0.190 |

***Supplemental Table 16.*** *Standardized coefficients values for T1w-alone WML segmentation.*

#### 3.2. 95% confidence intervals

| **Term** | **Lower 95%** | **Upper 95%** |
| --- | --- | --- |
| (Intercept) | -1.310e-04 | 1.310e-04 |
| RMSSD | -6.706e-06 | -3.904e-07 |

***Supplemental Table 17.*** *95% confidence intervals values for T1w-alone WML segmentation.*

#### 3.3. Breusch-Pagan test

| **Test** | **Statistic** | **df** | **p** |
| --- | --- | --- | --- |
| Breusch-Pagan | 1.762 | 1 | .184 |

***Supplemental Table 18.*** *Breusch-Pagan test values for T1w-alone WML segmentation.*

#### 3.4. Normal Q-Q plot

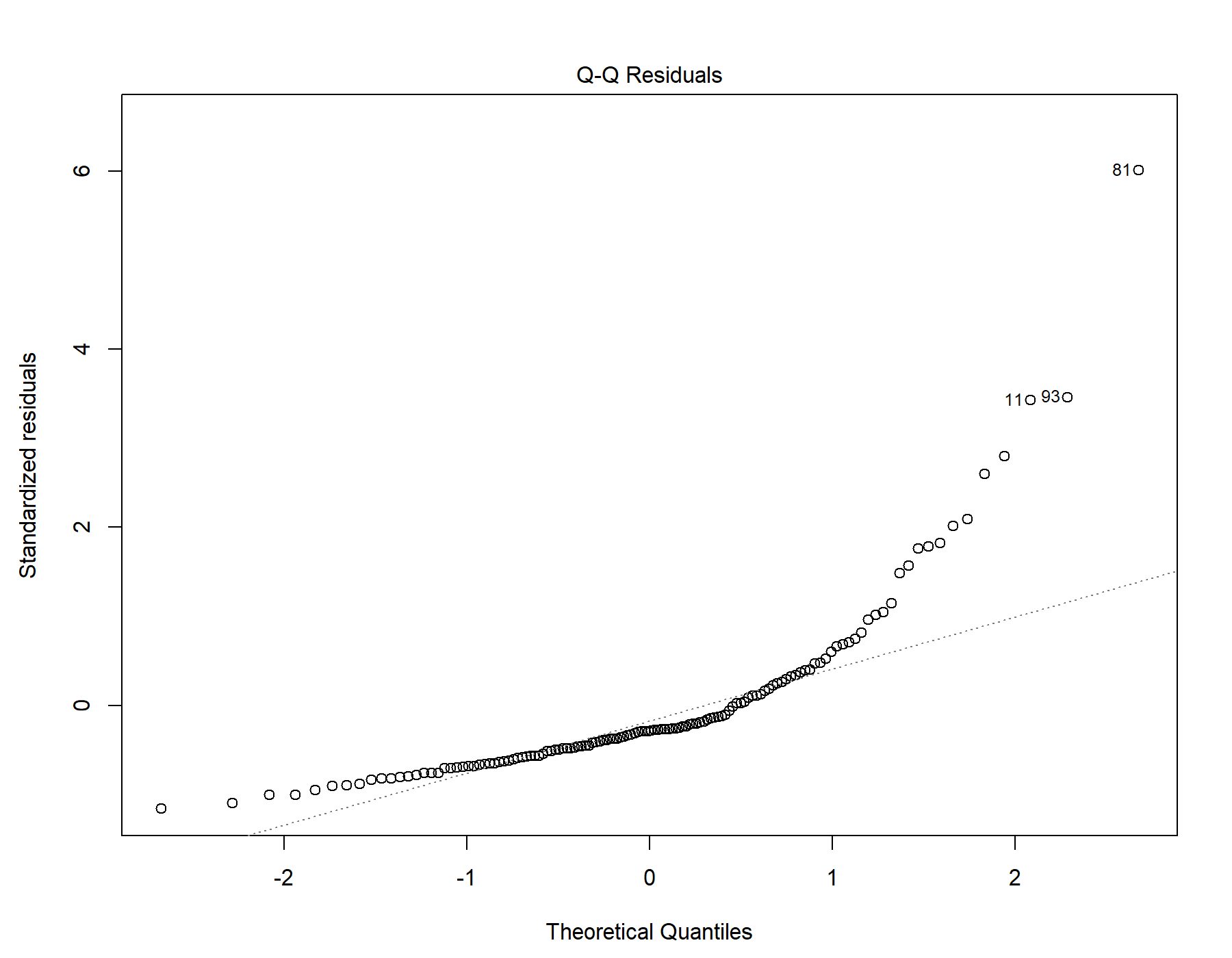

Supplemental Figure 9. Normal distribution quantile-quantile plot for T1w-alone WML segmentation.

#### 3.5. Residuals vs leverage

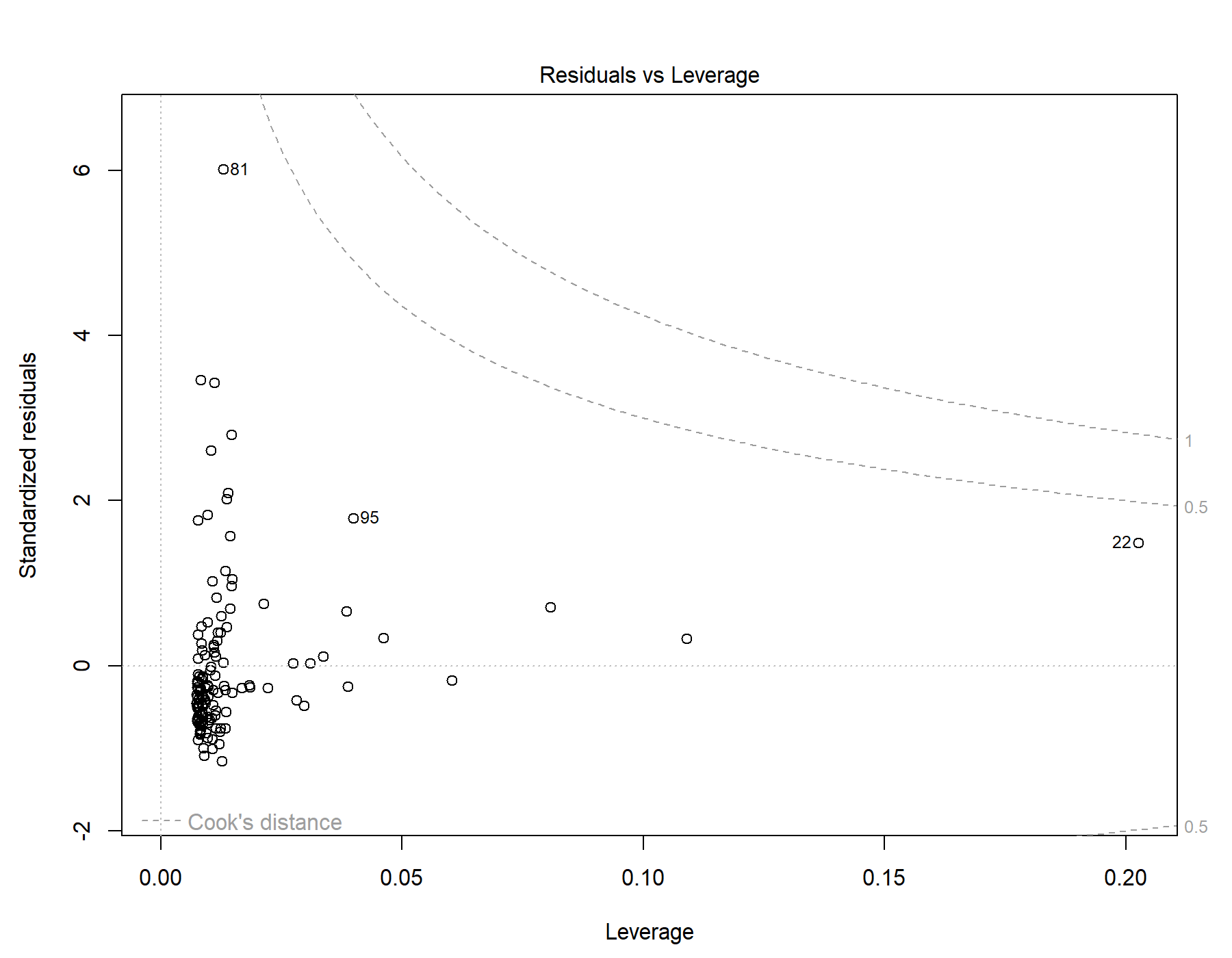

Supplemental Figure 10. Residuals vs leverage diagnostics for T1w-alone WML segmentation.

### 4. Global | All subjects | T1w+FLAIR WML vol | SDNN.LF._log | Exploratory unadjusted model

Model: T1w+FLAIR WML vol ~ SDNN.LF._log

#### 4.1. Standardized coefficients

| **Term** | **Estimate** | **Std. Error** | **t** | **p** | **β** |
| --- | --- | --- | --- | --- | --- |
| (Intercept) | -1.693e-20 | 7.020e-05 | -2.412e-16 | 1.000 |  |
| SDNN.LF._log | -4.852e-04 | 1.777e-04 | -2.73 | .007 | -0.231 |

***Supplemental Table 19.*** *Standardized coefficients values for T1w+FLAIR WML segmentation.*

#### 4.2. 95% confidence intervals

| **Term** | **Lower 95%** | **Upper 95%** |
| --- | --- | --- |
| (Intercept) | -1.389e-04 | 1.389e-04 |
| SDNN.LF._log | -8.367e-04 | -1.337e-04 |

***Supplemental Table 20.*** *95% confidence intervals values for T1w+FLAIR WML segmentation.*

#### 4.3. Breusch-Pagan test

| **Test** | **Statistic** | **df** | **p** |
| --- | --- | --- | --- |
| Breusch-Pagan | 1.618 | 1 | .203 |

***Supplemental Table 21.*** *Breusch-Pagan test values for T1w+FLAIR WML segmentation.*

#### 4.4. Normal Q-Q plot

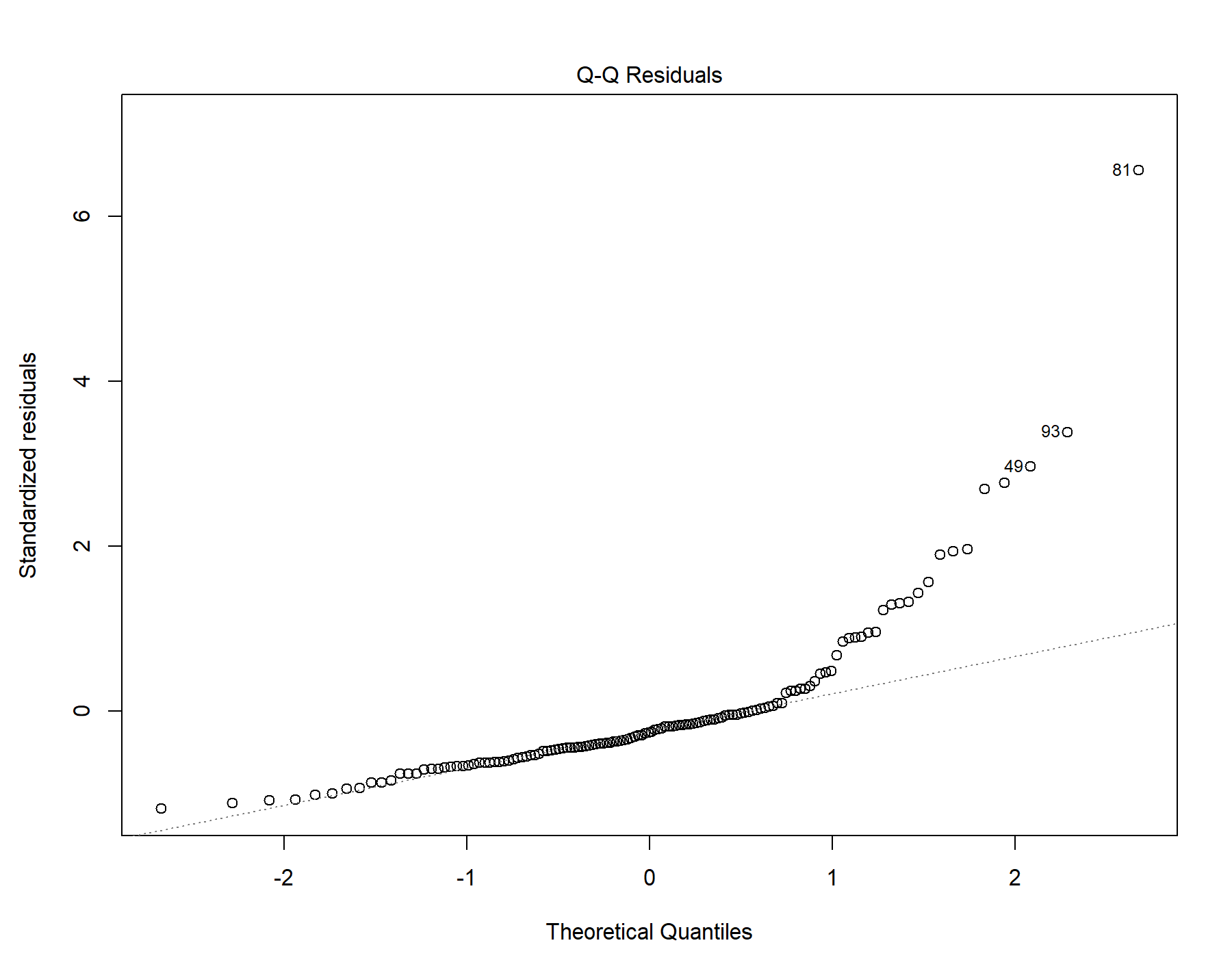

Supplemental Figure 11. Normal distribution quantile-quantile plot for T1w+FLAIR WML segmentation.

#### 4.5. Residuals vs leverage

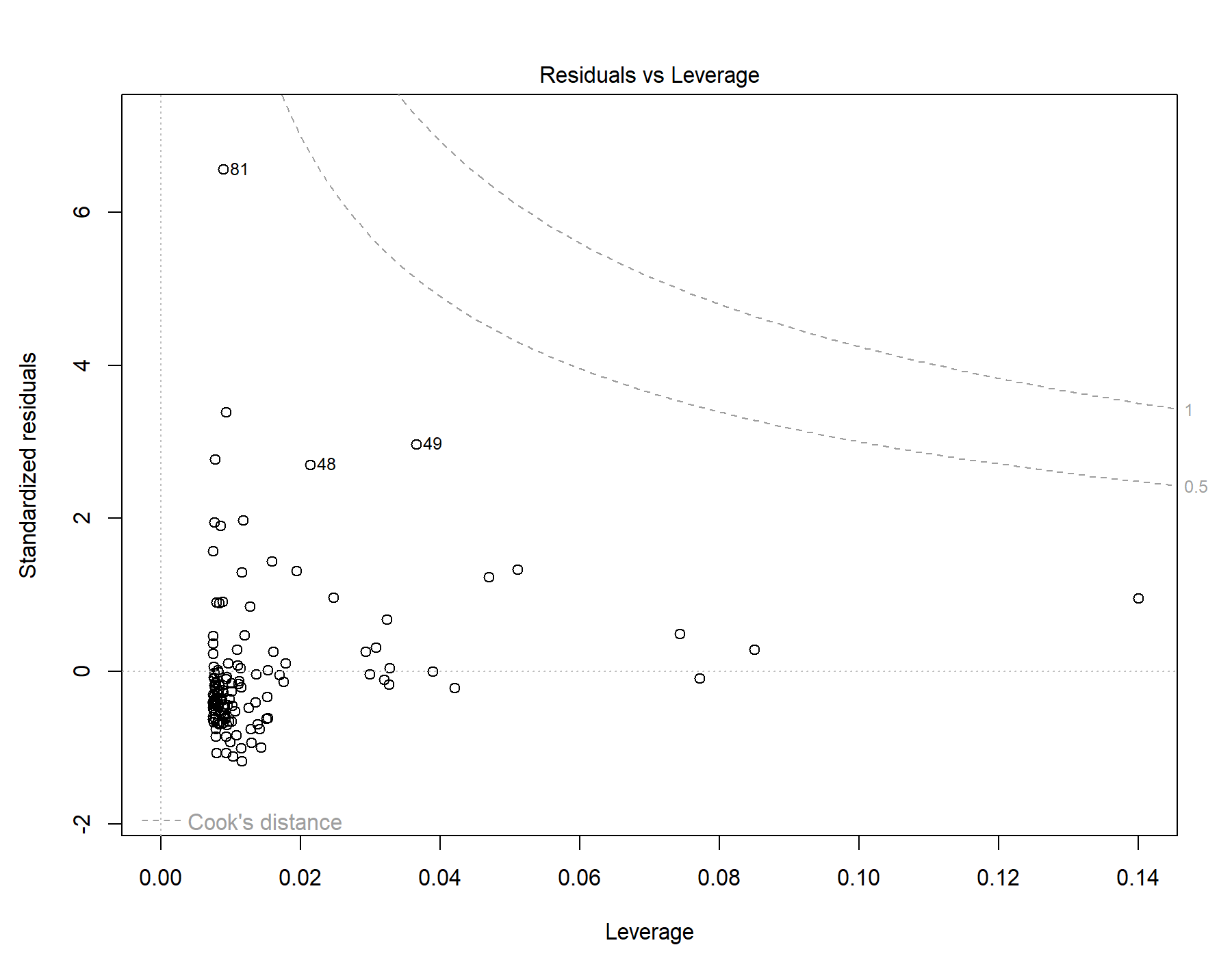

Supplemental Figure 12. Residuals vs leverage diagnostics for T1w+FLAIR WML segmentation.

### 5. Global | All subjects | T1w+T2w WML vol | SDNN.LF._log | Exploratory unadjusted model

Model: T1w+T2w WML vol ~ SDNN.LF._log

#### 5.1. Standardized coefficients

| **Term** | **Estimate** | **Std. Error** | **t** | **p** | **β** |
| --- | --- | --- | --- | --- | --- |
| (Intercept) | 6.700e-20 | 8.017e-05 | 8.357e-16 | 1.000 |  |
| SDNN.LF._log | -5.088e-04 | 2.029e-04 | -2.51 | .013 | -0.213 |

***Supplemental Table 22.*** *Standardized coefficients values for T1w+T2w WML segmentation.*

#### 5.2. 95% confidence intervals

| **Term** | **Lower 95%** | **Upper 95%** |
| --- | --- | --- |
| (Intercept) | -1.586e-04 | 1.586e-04 |
| SDNN.LF._log | -9.102e-04 | -1.073e-04 |

***Supplemental Table 23.*** *95% confidence intervals values for T1w+T2w WML segmentation.*

#### 5.3. Breusch-Pagan test

| **Test** | **Statistic** | **df** | **p** |
| --- | --- | --- | --- |
| Breusch-Pagan | 1.277 | 1 | .258 |

***Supplemental Table 24.*** *Breusch-Pagan test values for T1w+T2w WML segmentation.*

#### 5.4. Normal Q-Q plot

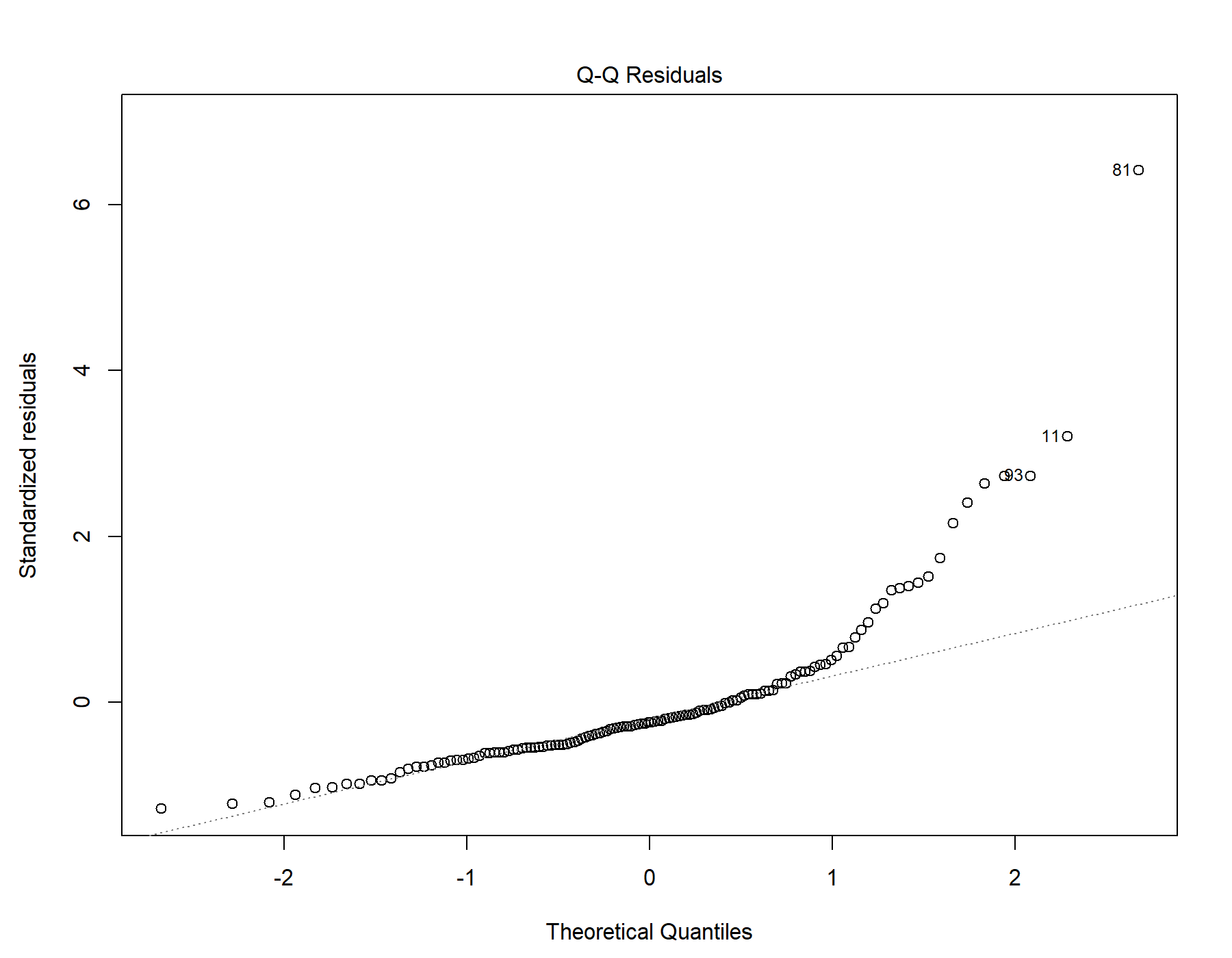

Supplemental Figure 13. Normal distribution quantile-quantile plot for T1w+T2w WML segmentation.

#### 5.5. Residuals vs leverage

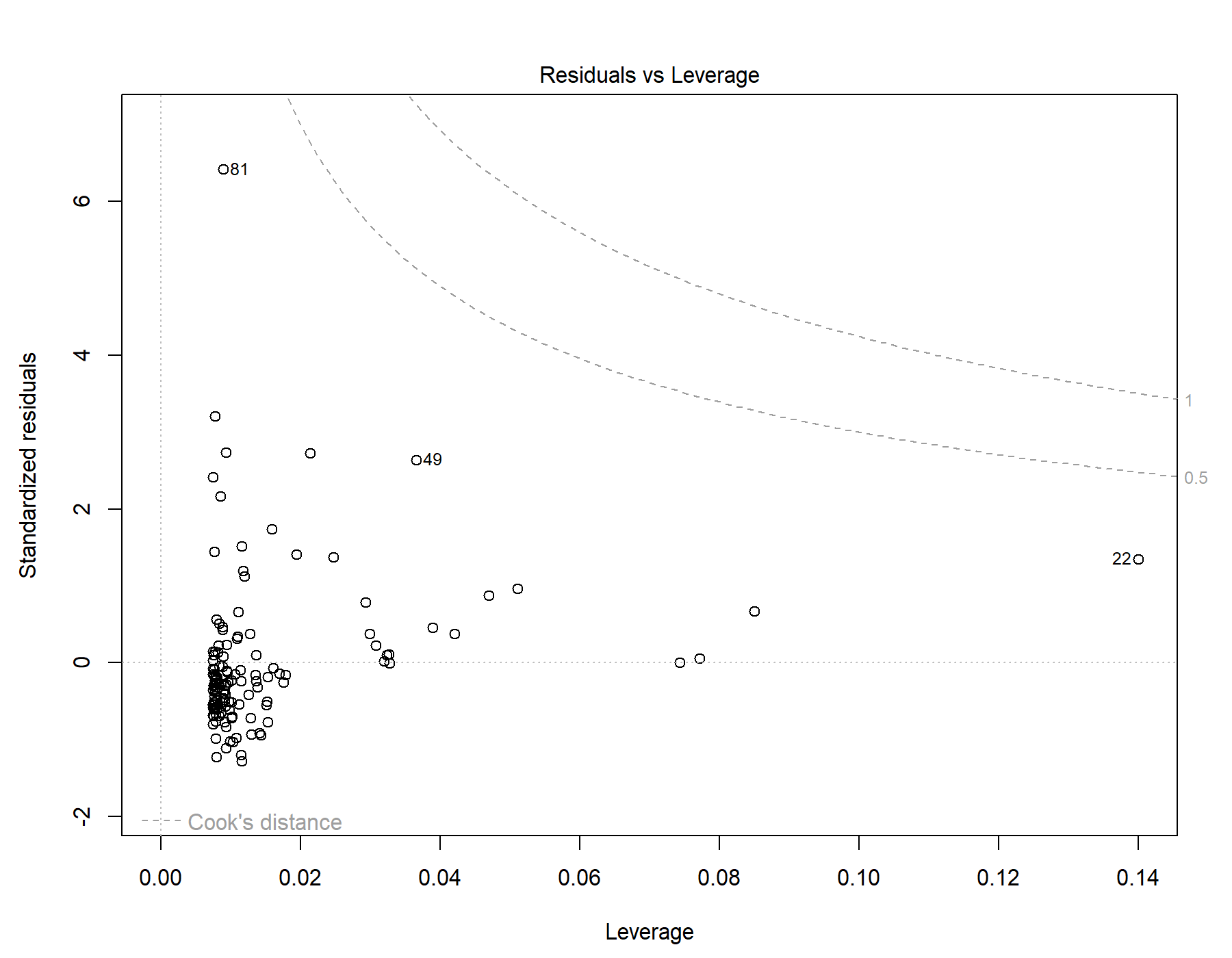

Supplemental Figure 14. Residuals vs leverage diagnostics for T1w+T2w WML segmentation.

### 6. Global | All subjects | T1w-alone WML vol | SDNN.LF._log | Exploratory unadjusted model

Model: T1w-alone WML vol ~ SDNN.LF._log

#### 6.1. Standardized coefficients

| **Term** | **Estimate** | **Std. Error** | **t** | **p** | **β** |
| --- | --- | --- | --- | --- | --- |
| (Intercept) | 4.027e-20 | 6.589e-05 | 6.112e-16 | 1.000 |  |
| SDNN.LF._log | -4.207e-04 | 1.668e-04 | -2.52 | .013 | -0.214 |

***Supplemental Table 25.*** *Standardized coefficients values for T1w-alone WML segmentation.*

#### 6.2. 95% confidence intervals

| **Term** | **Lower 95%** | **Upper 95%** |
| --- | --- | --- |
| (Intercept) | -1.303e-04 | 1.303e-04 |
| SDNN.LF._log | -7.507e-04 | -9.081e-05 |

***Supplemental Table 26.*** *95% confidence intervals values for T1w-alone WML segmentation.*

#### 6.3. Breusch-Pagan test

| **Test** | **Statistic** | **df** | **p** |
| --- | --- | --- | --- |
| Breusch-Pagan | 1.199 | 1 | .273 |

***Supplemental Table 27.*** *Breusch-Pagan test values for T1w-alone WML segmentation.*

#### 6.4. Normal Q-Q plot

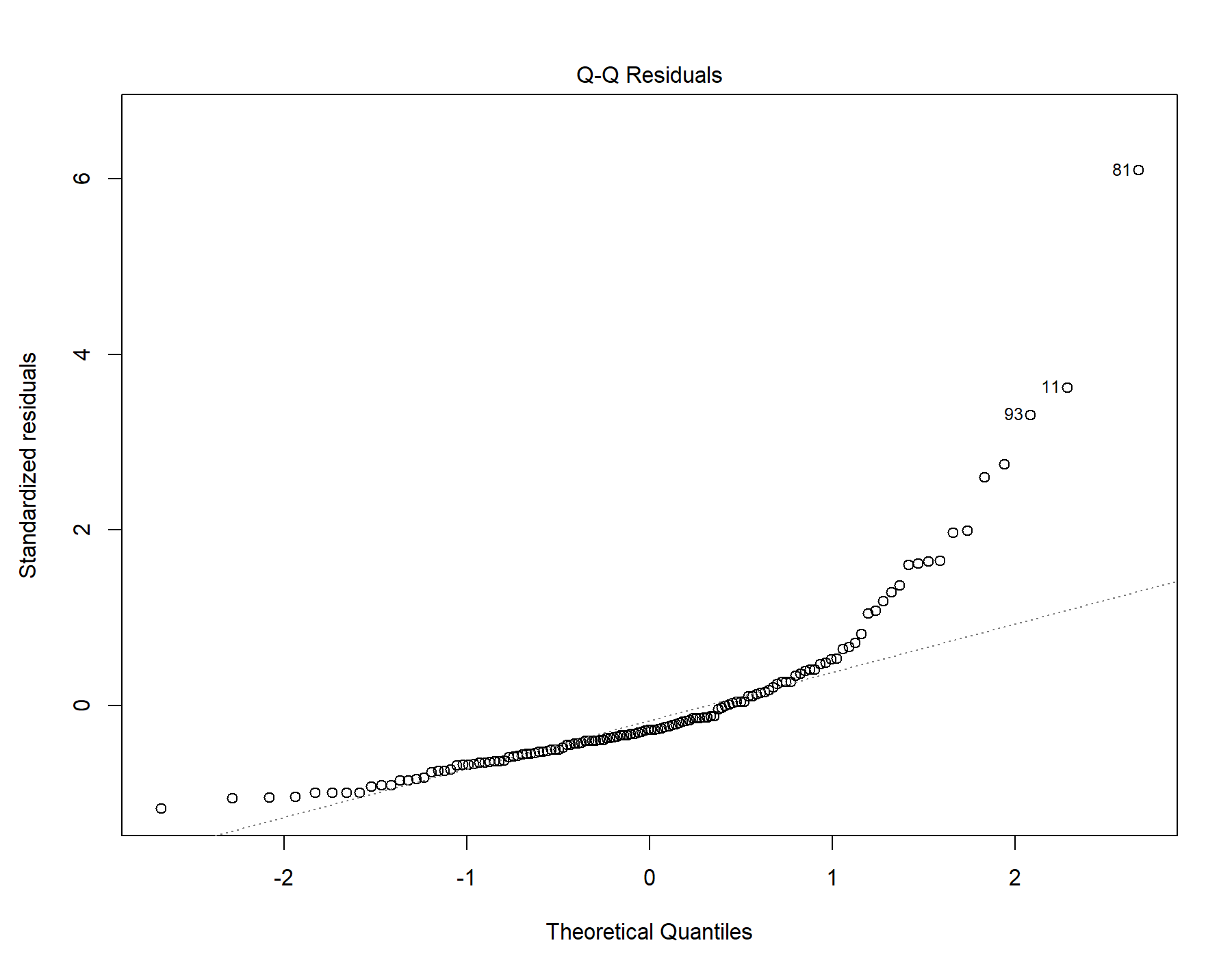

Supplemental Figure 15. Normal distribution quantile-quantile plot for T1w-alone WML segmentation.

#### 6.5. Residuals vs leverage

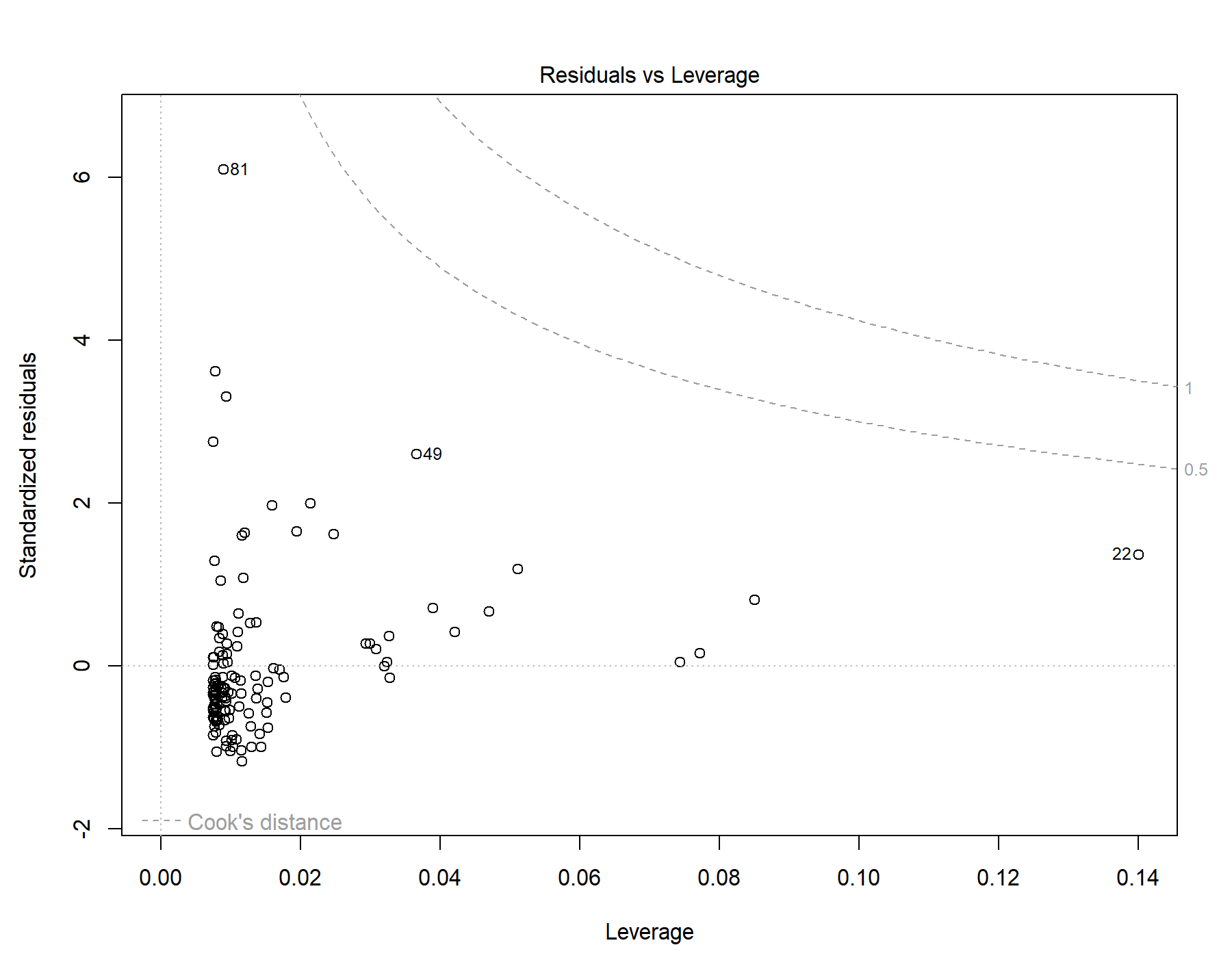

Supplemental Figure 16. Residuals vs leverage diagnostics for T1w-alone WML segmentation.

### 7. Global | All subjects | T1w+FLAIR WML vol | SDNN.HF._log | Exploratory unadjusted model

Model: T1w+FLAIR WML vol ~ SDNN.HF._log

#### 7.1. Standardized coefficients

| **Term** | **Estimate** | **Std. Error** | **t** | **p** | **β** |
| --- | --- | --- | --- | --- | --- |
| (Intercept) | -6.979e-20 | 6.830e-05 | -1.022e-15 | 1.000 |  |
| SDNN.HF._log | -7.001e-04 | 1.790e-04 | -3.91 | <.001 | -0.322 |

***Supplemental Table 28.*** *Standardized coefficients values for T1w+FLAIR WML segmentation.*

#### 7.2. 95% confidence intervals

| **Term** | **Lower 95%** | **Upper 95%** |
| --- | --- | --- |
| (Intercept) | -1.351e-04 | 1.351e-04 |
| SDNN.HF._log | -0.00105 | -3.460e-04 |

***Supplemental Table 29.*** *95% confidence intervals values for T1w+FLAIR WML segmentation.*

#### 7.3. Breusch-Pagan test

| **Test** | **Statistic** | **df** | **p** |
| --- | --- | --- | --- |
| Breusch-Pagan | 1.667 | 1 | .197 |

***Supplemental Table 30.*** *Breusch-Pagan test values for T1w+FLAIR WML segmentation.*

#### 7.4. Normal Q-Q plot

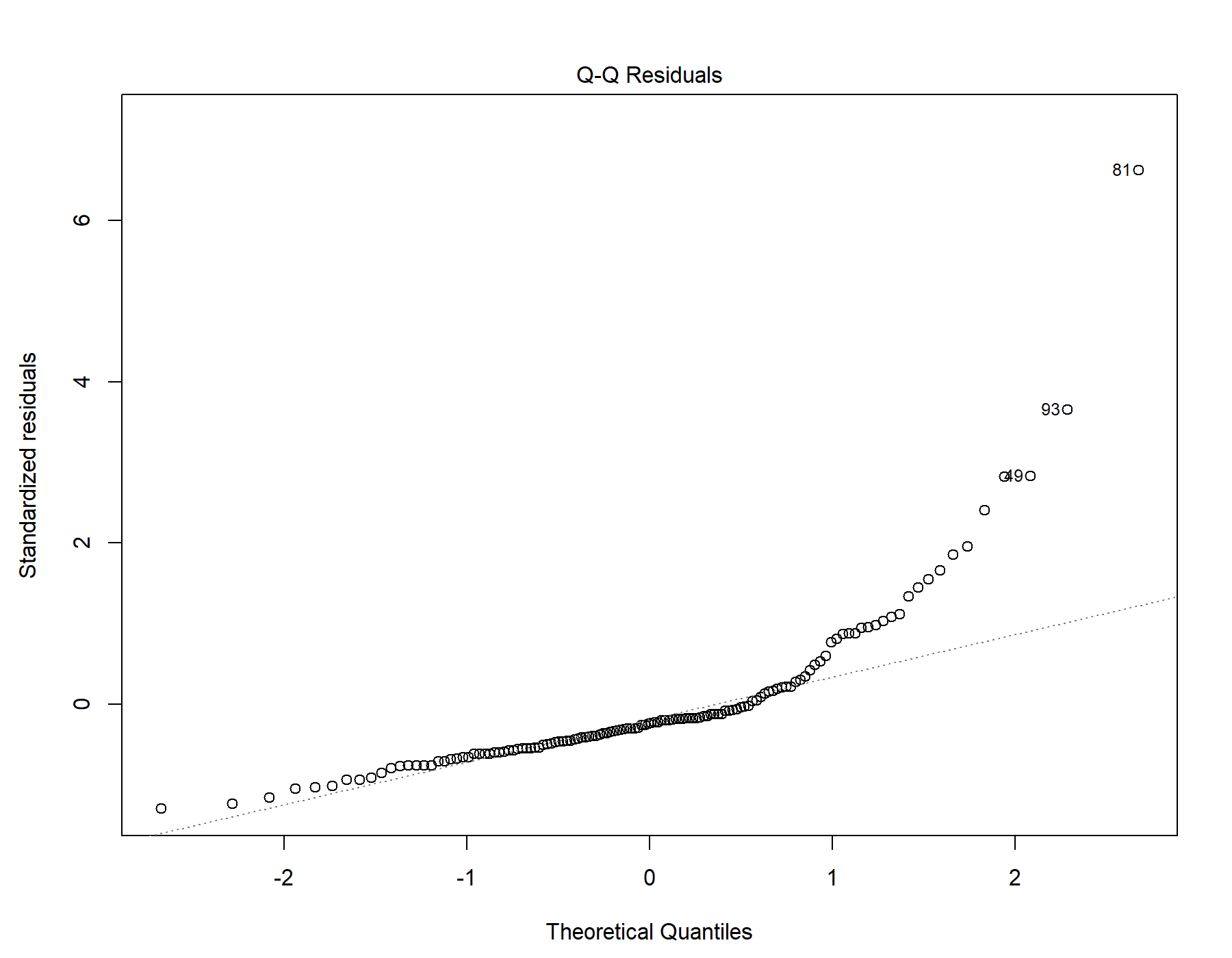

Supplemental Figure 17. Normal distribution quantile-quantile plot for T1w+FLAIR WML segmentation.

#### 7.5. Residuals vs leverage

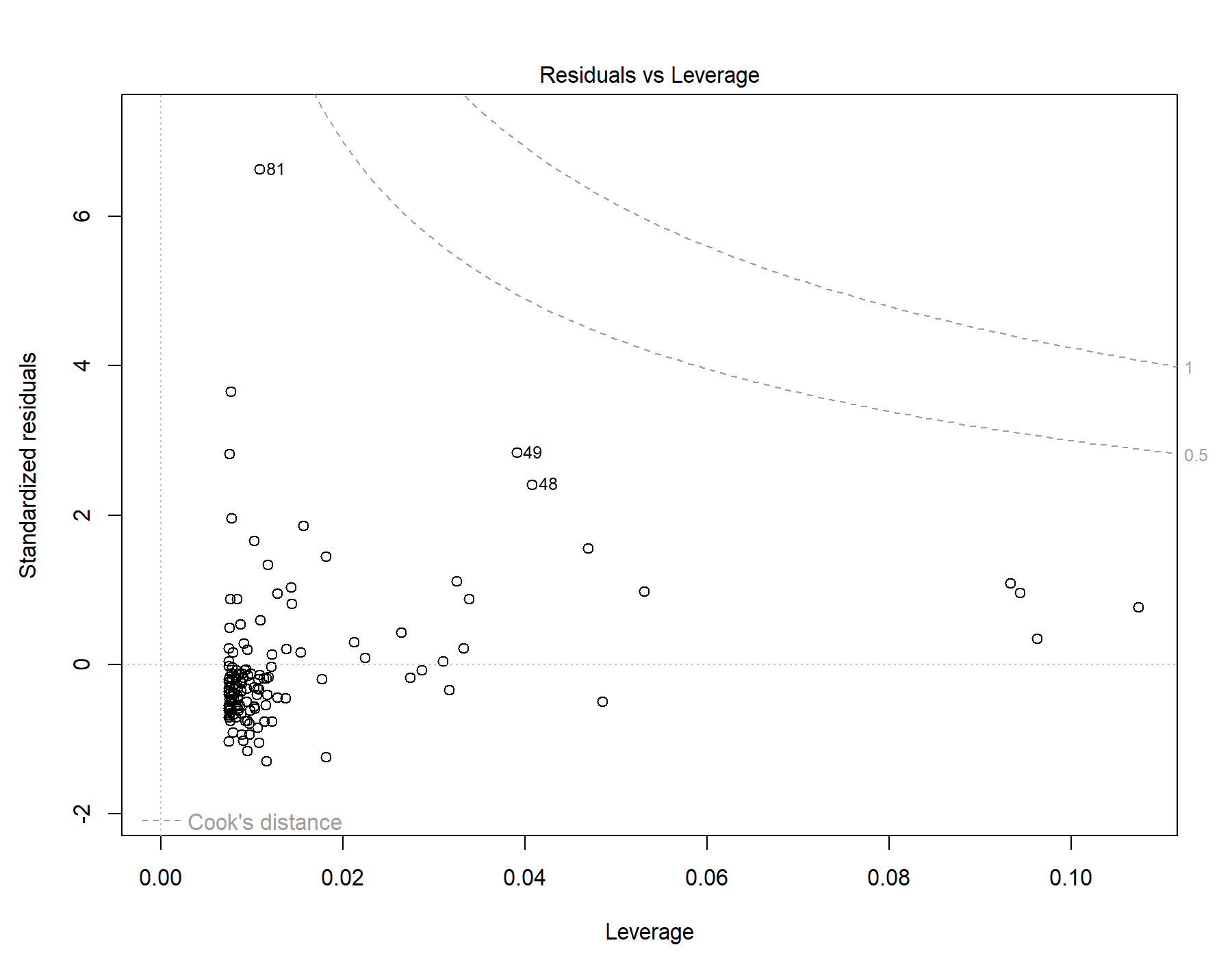

Supplemental Figure 18. Residuals vs leverage diagnostics for T1w+FLAIR WML segmentation.

### 8. Global | All subjects | T1w+T2w WML vol | SDNN.HF._log | Exploratory unadjusted model

Model: T1w+T2w WML vol ~ SDNN.HF._log

#### 8.1. Standardized coefficients

| **Term** | **Estimate** | **Std. Error** | **t** | **p** | **β** |
| --- | --- | --- | --- | --- | --- |
| (Intercept) | 6.758e-21 | 7.775e-05 | 8.692e-17 | 1.000 |  |
| SDNN.HF._log | -7.898e-04 | 2.037e-04 | -3.88 | <.001 | -0.320 |

***Supplemental Table 31.*** *Standardized coefficients values for T1w+T2w WML segmentation.*

#### 8.2. 95% confidence intervals

| **Term** | **Lower 95%** | **Upper 95%** |
| --- | --- | --- |
| (Intercept) | -1.538e-04 | 1.538e-04 |
| SDNN.HF._log | -0.00119 | -3.868e-04 |

***Supplemental Table 32.*** *95% confidence intervals values for T1w+T2w WML segmentation.*

#### 8.3. Breusch-Pagan test

| **Test** | **Statistic** | **df** | **p** |
| --- | --- | --- | --- |
| Breusch-Pagan | 1.453 | 1 | .228 |

***Supplemental Table 33.*** *Breusch-Pagan test values for T1w+T2w WML segmentation.*

#### 8.4. Normal Q-Q plot

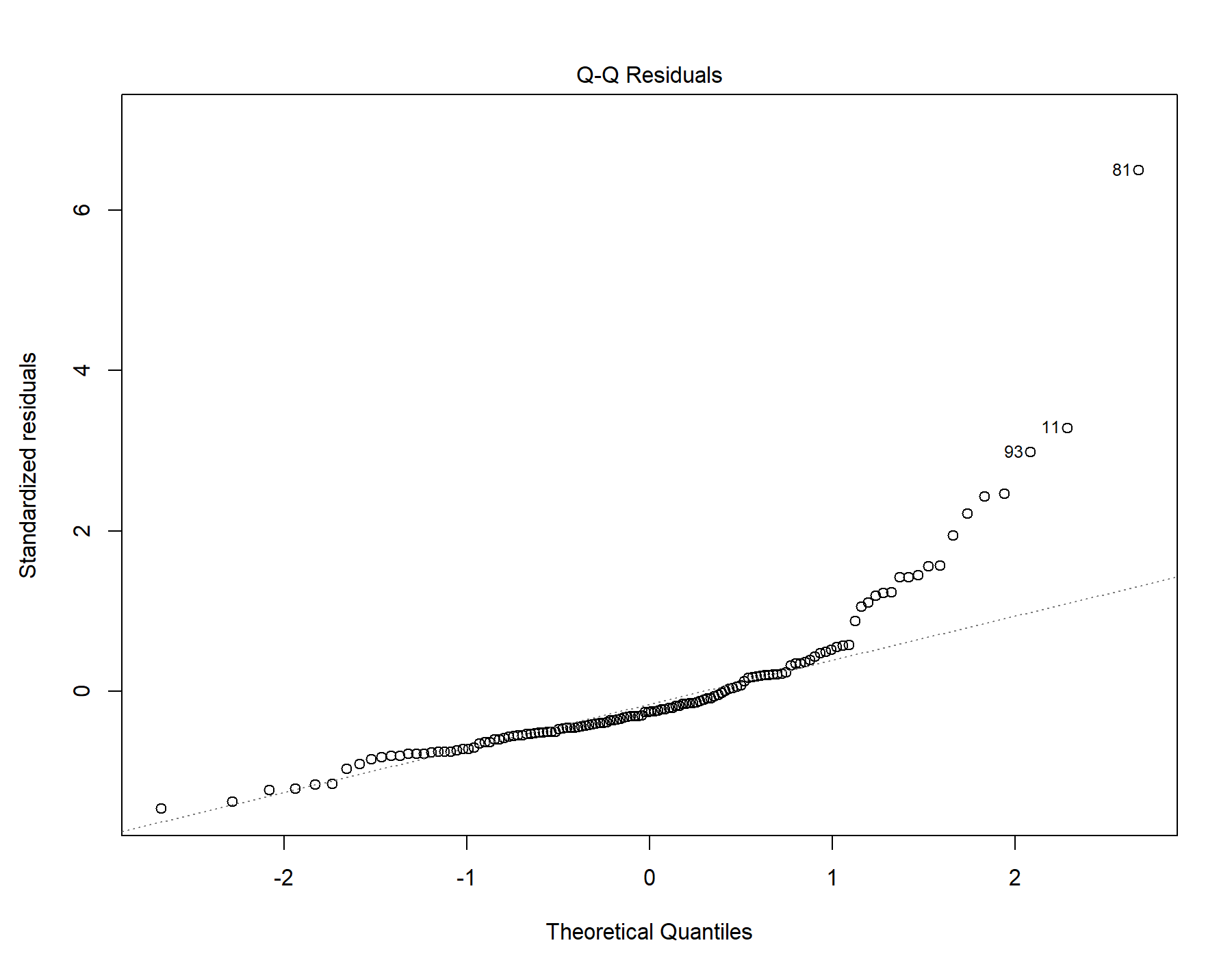

Supplemental Figure 19. Normal distribution quantile-quantile plot for T1w+T2w WML segmentation.

#### 8.5. Residuals vs leverage

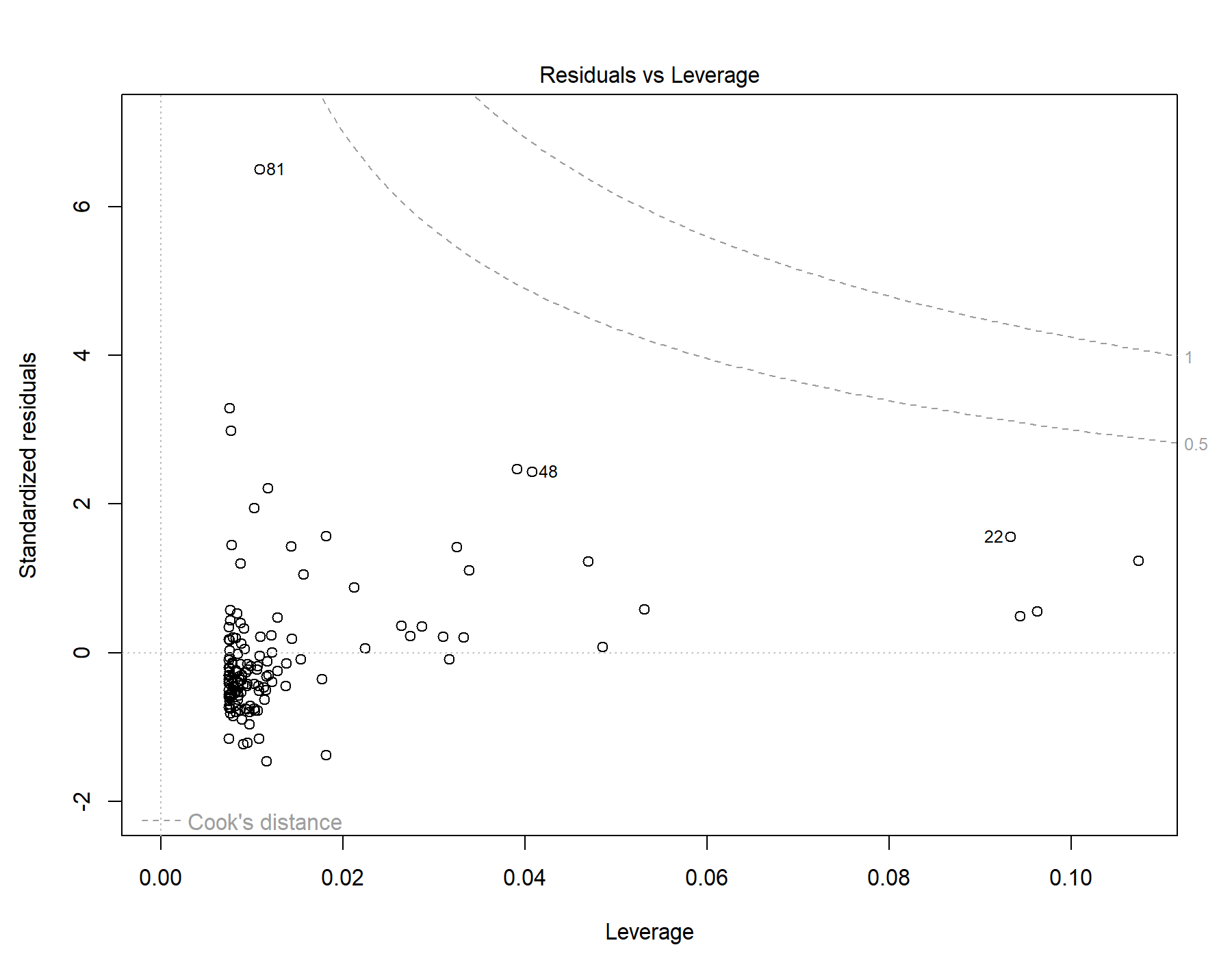

Supplemental Figure 20. Residuals vs leverage diagnostics for T1w+T2w WML segmentation.

### 9. Global | All subjects | T1w-alone WML vol | SDNN.HF._log | Exploratory unadjusted model

Model: T1w-alone WML vol ~ SDNN.HF._log

#### 9.1. Standardized coefficients

| **Term** | **Estimate** | **Std. Error** | **t** | **p** | **β** |
| --- | --- | --- | --- | --- | --- |
| (Intercept) | -6.781e-21 | 6.423e-05 | -1.056e-16 | 1.000 |  |
| SDNN.HF._log | -6.211e-04 | 1.683e-04 | -3.69 | <.001 | -0.306 |

***Supplemental Table 34.*** *Standardized coefficients values for T1w-alone WML segmentation.*

#### 9.2. 95% confidence intervals

| **Term** | **Lower 95%** | **Upper 95%** |
| --- | --- | --- |
| (Intercept) | -1.270e-04 | 1.270e-04 |
| SDNN.HF._log | -9.540e-04 | -2.882e-04 |

***Supplemental Table 35.*** *95% confidence intervals values for T1w-alone WML segmentation.*

#### 9.3. Breusch-Pagan test

| **Test** | **Statistic** | **df** | **p** |
| --- | --- | --- | --- |
| Breusch-Pagan | 1.07 | 1 | .301 |

***Supplemental Table 36.*** *Breusch-Pagan test values for T1w-alone WML segmentation.*

#### 9.4. Normal Q-Q plot

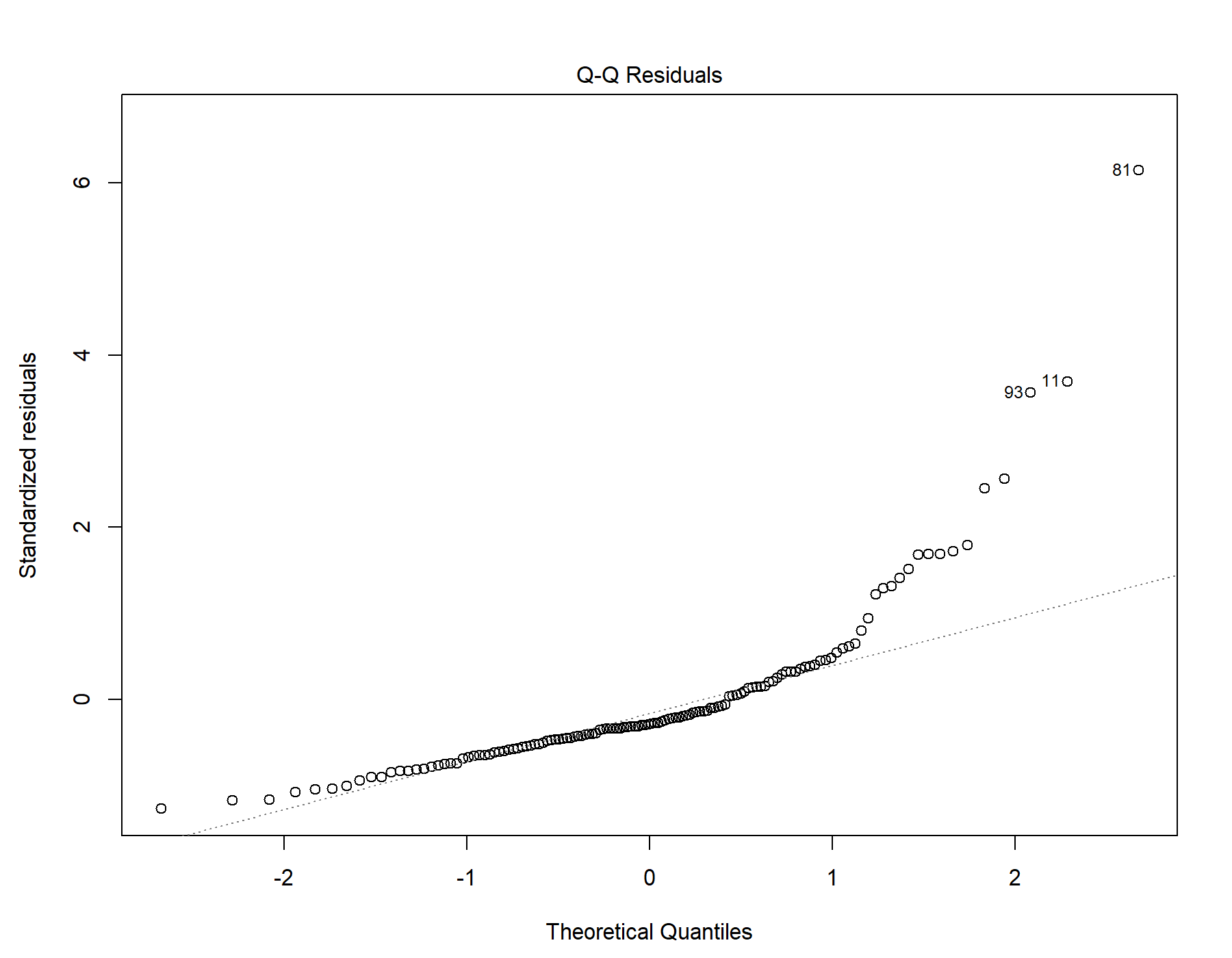

Supplemental Figure 21. Normal distribution quantile-quantile plot for T1w-alone WML segmentation.

#### 9.5. Residuals vs leverage

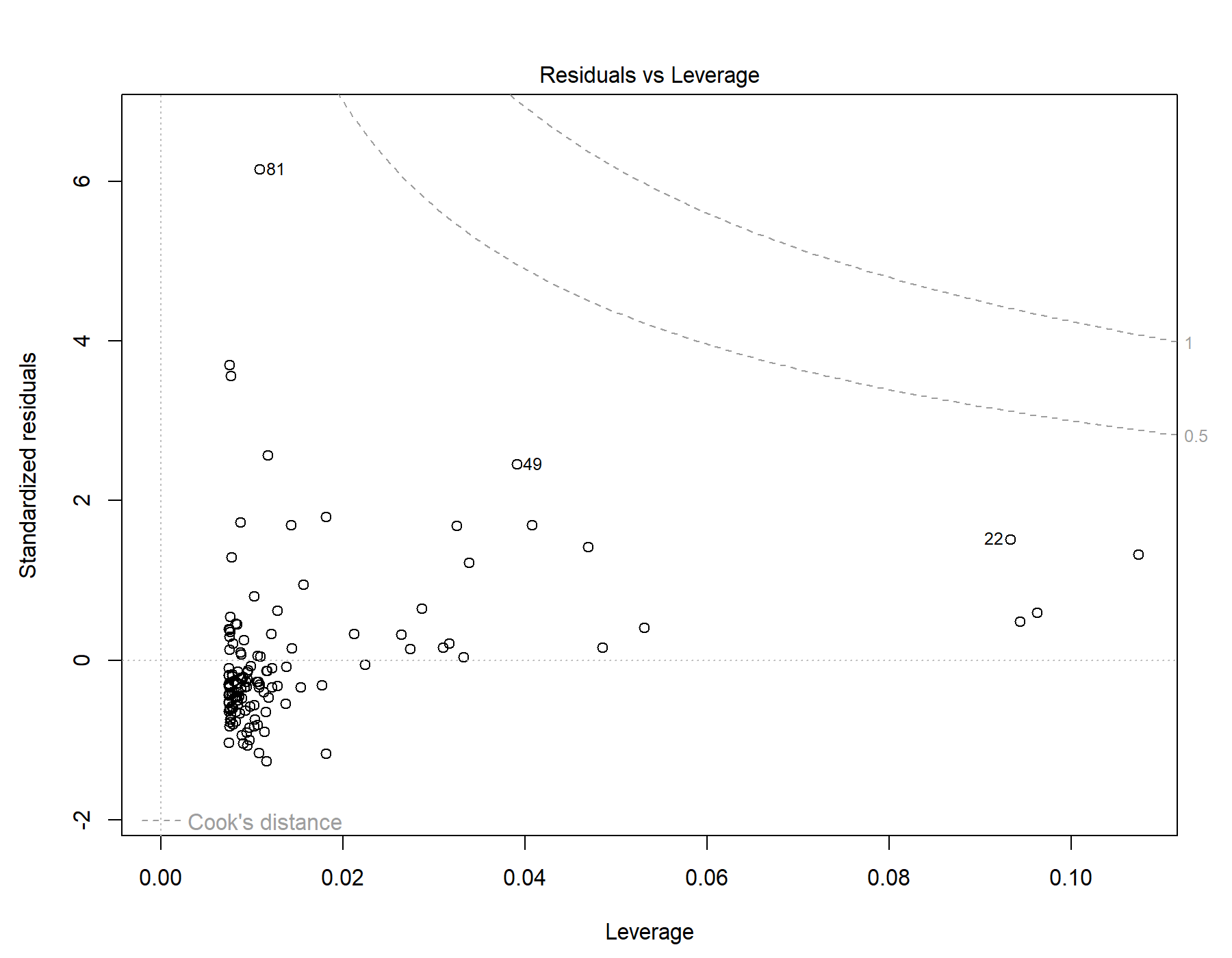

Supplemental Figure 22. Residuals vs leverage diagnostics for T1w-alone WML segmentation.

### 10. Global | All subjects | T1w+FLAIR WML vol | RMSSD | Primary interaction model

Model: T1w+FLAIR WML vol ~ RMSSD + Age + Sex + Age * RMSSD + Sex * RMSSD

#### 10.1. Standardized coefficients

| **Term** | **Estimate** | **Std. Error** | **t** | **p** | **β** |
| --- | --- | --- | --- | --- | --- |
| (Intercept) | -5.424e-04 | 2.019e-04 | -2.69 | .008 |  |
| RMSSD | 9.694e-07 | 5.996e-06 | 0.162 | .872 | 0.0485 |
| Age | 1.530e-04 | 1.605e-05 | 9.54 | <.001 | 0.650 |
| Sex | -6.078e-06 | 1.210e-04 | -0.0502 | .960 | -0.00338 |
| RMSSD:Age | -5.988e-07 | 3.242e-07 | -1.85 | .067 | -0.191 |
| RMSSD:Sex | -1.050e-07 | 3.478e-06 | -0.0302 | .976 | -0.00970 |

***Supplemental Table 37.*** *Standardized coefficients values for T1w+FLAIR WML segmentation.*

#### 10.2. 95% confidence intervals

| **Term** | **Lower 95%** | **Upper 95%** |
| --- | --- | --- |
| (Intercept) | -9.418e-04 | -1.430e-04 |
| RMSSD | -1.089e-05 | 1.283e-05 |
| Age | 1.213e-04 | 1.848e-04 |
| Sex | -2.455e-04 | 2.334e-04 |
| RMSSD:Age | -1.240e-06 | 4.276e-08 |
| RMSSD:Sex | -6.987e-06 | 6.777e-06 |

***Supplemental Table 38.*** *95% confidence intervals values for T1w+FLAIR WML segmentation.*

#### 10.3. Breusch-Pagan test

| **Test** | **Statistic** | **df** | **p** |
| --- | --- | --- | --- |
| Breusch-Pagan | 19.883 | 5 | .001 |

***Supplemental Table 39.*** *Breusch-Pagan test values for T1w+FLAIR WML segmentation.*

#### 10.4. Normal Q-Q plot

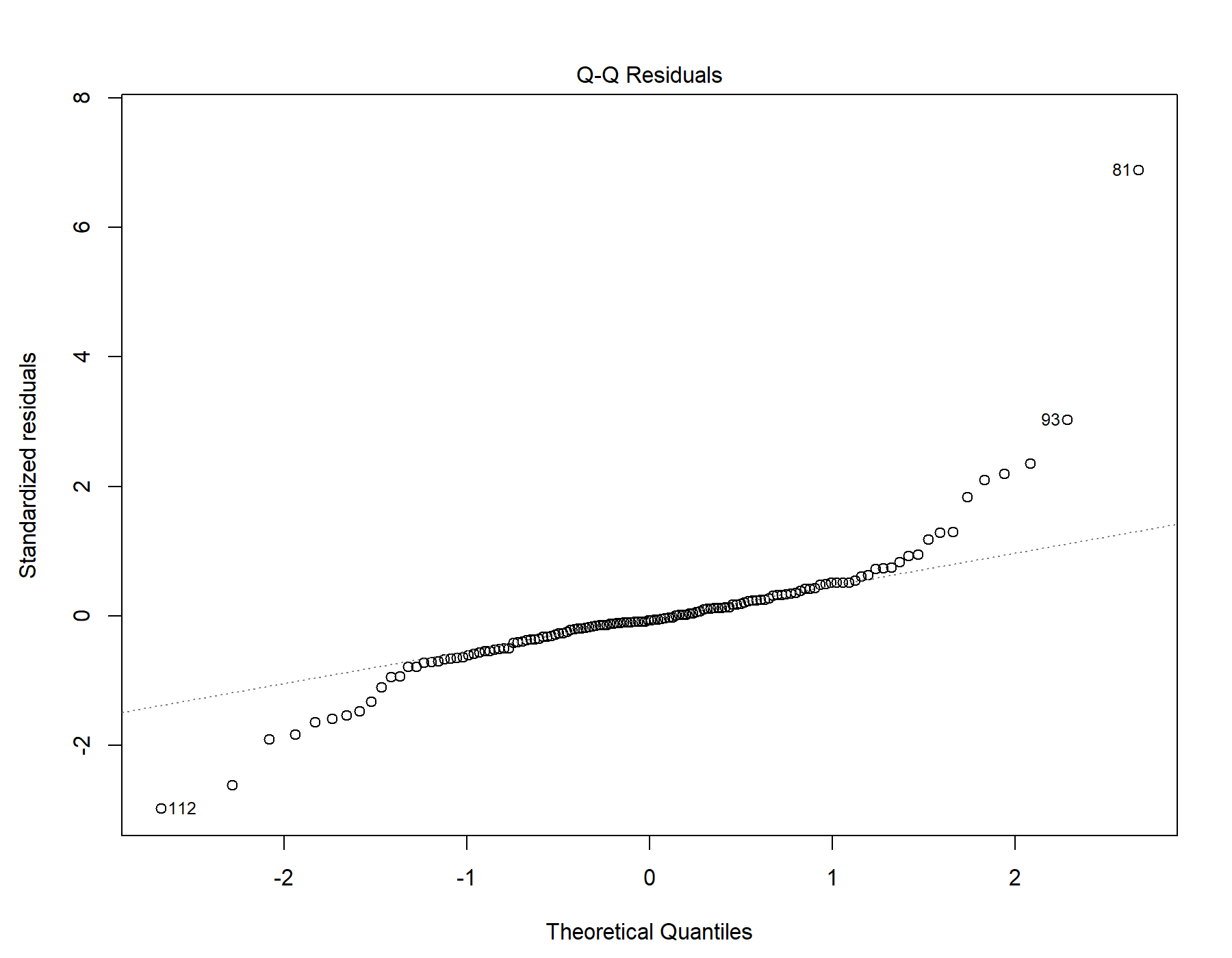

Supplemental Figure 23. Normal distribution quantile-quantile plot for T1w+FLAIR WML segmentation.

#### 10.5. Residuals vs leverage

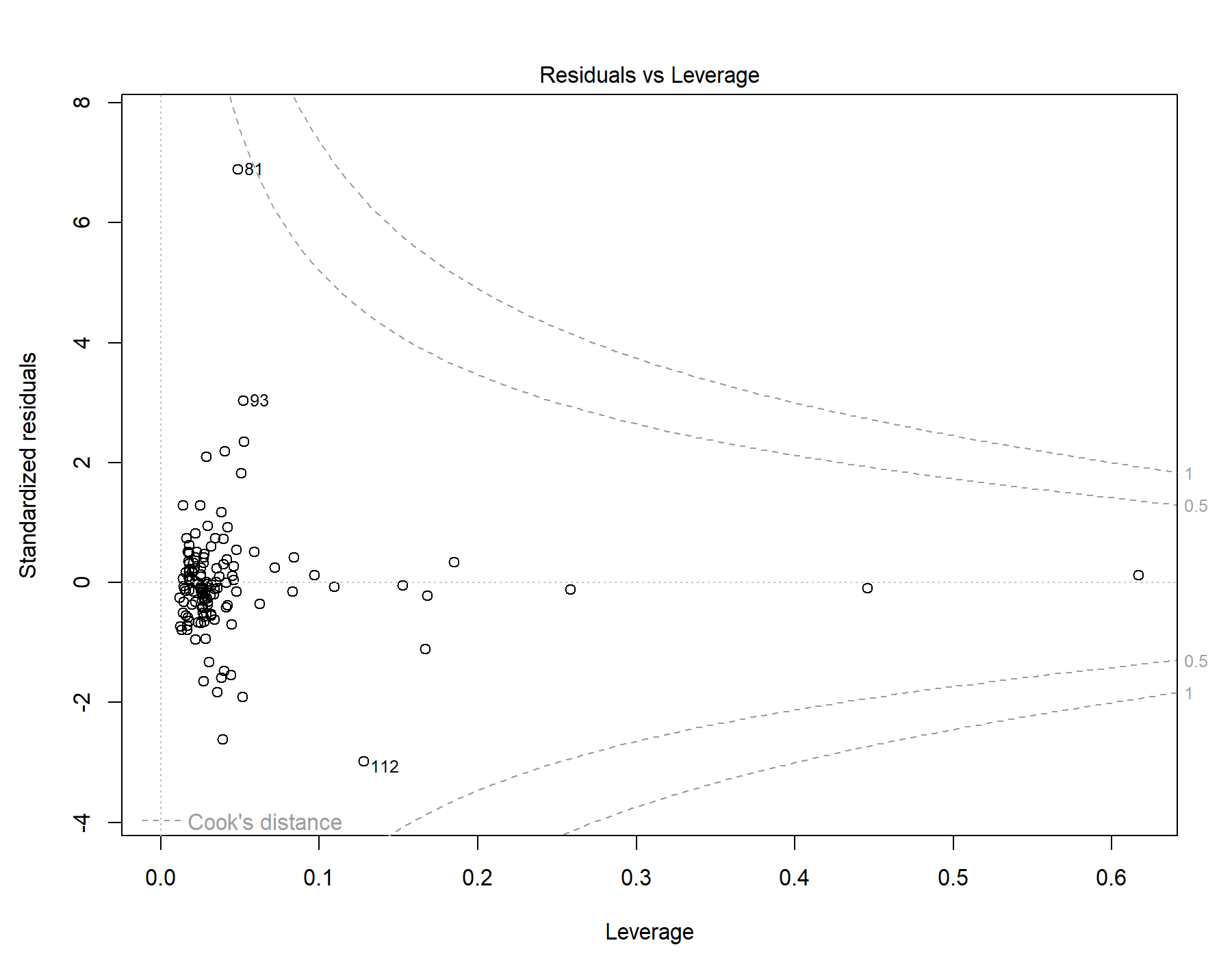

Supplemental Figure 24. Residuals vs leverage diagnostics for T1w+FLAIR WML segmentation.

### 11. Global | All subjects | T1w+T2w WML vol | RMSSD | Primary interaction model

Model: T1w+T2w WML vol ~ RMSSD + Age + Sex + Age *	RMSSD + Sex * RMSSD

#### 11.1. Standardized coefficients

| **Term** | **Estimate** | **Std. Error** | **t** | **p** | **β** |
| --- | --- | --- | --- | --- | --- |
| (Intercept) | -5.397e-04 | 2.214e-04 | -2.44 | .016 |  |
| RMSSD | 4.281e-06 | 6.575e-06 | 0.651 | .516 | 0.188 |
| Age | 1.888e-04 | 1.760e-05 | 10.7 | <.001 | 0.705 |
| Sex | -8.050e-05 | 1.327e-04 | -0.607 | .545 | -0.0393 |
| RMSSD:Age | -6.160e-07 | 3.556e-07 | -1.73 | .086 | -0.173 |
| RMSSD:Sex | -1.247e-06 | 3.814e-06 | -0.327 | .744 | -0.101 |

***Supplemental Table 40.*** *Standardized coefficients values for T1w+T2w WML segmentation.*

#### 11.2. 95% confidence intervals

| **Term** | **Lower 95%** | **Upper 95%** |
| --- | --- | --- |
| (Intercept) | -9.777e-04 | -1.016e-04 |
| RMSSD | -8.729e-06 | 1.729e-05 |
| Age | 1.540e-04 | 2.236e-04 |
| Sex | -3.431e-04 | 1.821e-04 |
| RMSSD:Age | -1.320e-06 | 8.751e-08 |
| RMSSD:Sex | -8.795e-06 | 6.300e-06 |

***Supplemental Table 41.*** *95% confidence intervals values for T1w+T2w WML segmentation.*

#### 11.3. Breusch-Pagan test

| **Test** | **Statistic** | **df** | **p** |
| --- | --- | --- | --- |
| Breusch-Pagan | 18.505 | 5 | .002 |

***Supplemental Table 42.*** *Breusch-Pagan test values for T1w+T2w WML segmentation.*

#### 11.4. Normal Q-Q plot

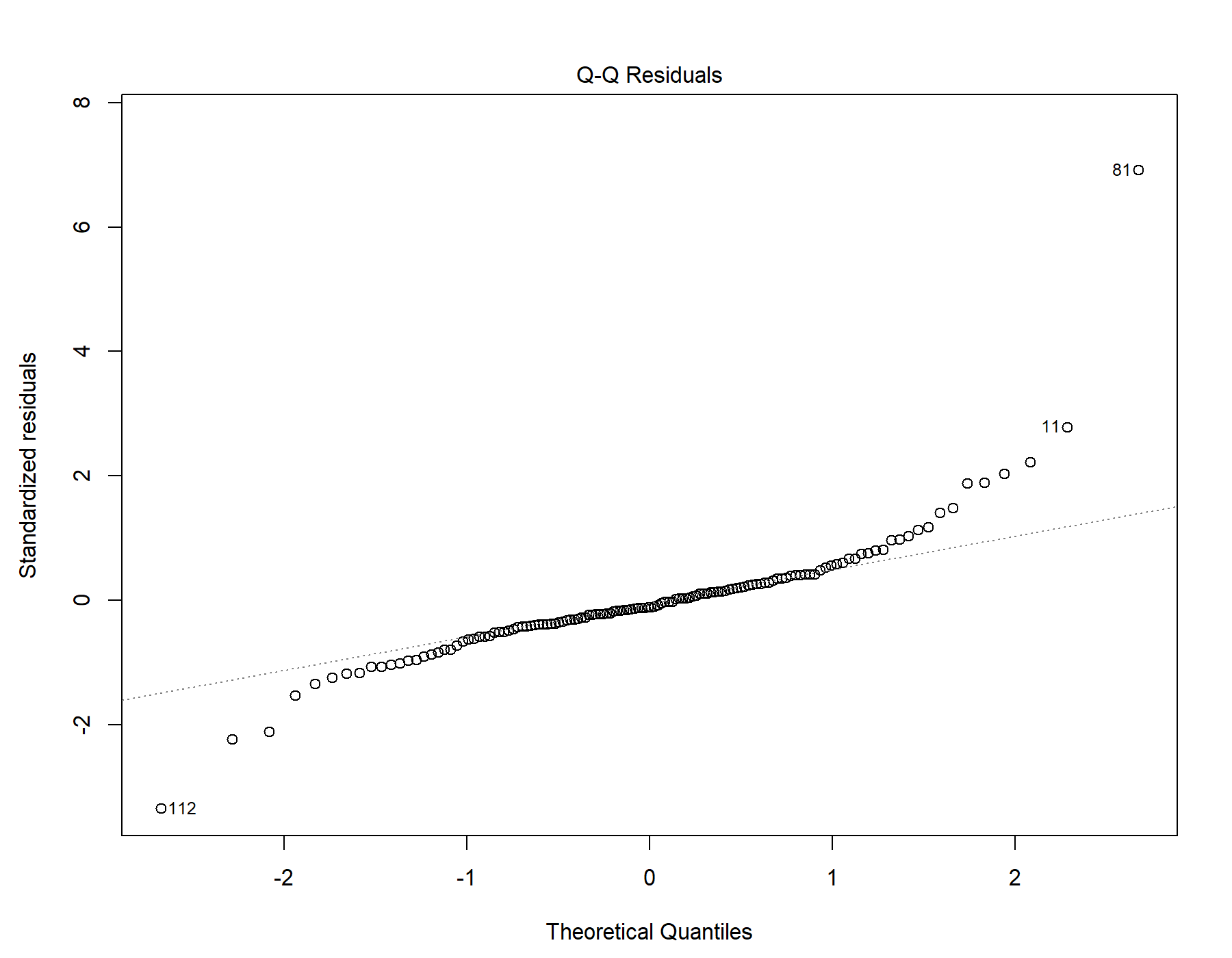

Supplemental Figure 25. Normal distribution quantile-quantile plot for T1w+T2w WML segmentation.

#### 11.5. Residuals vs leverage

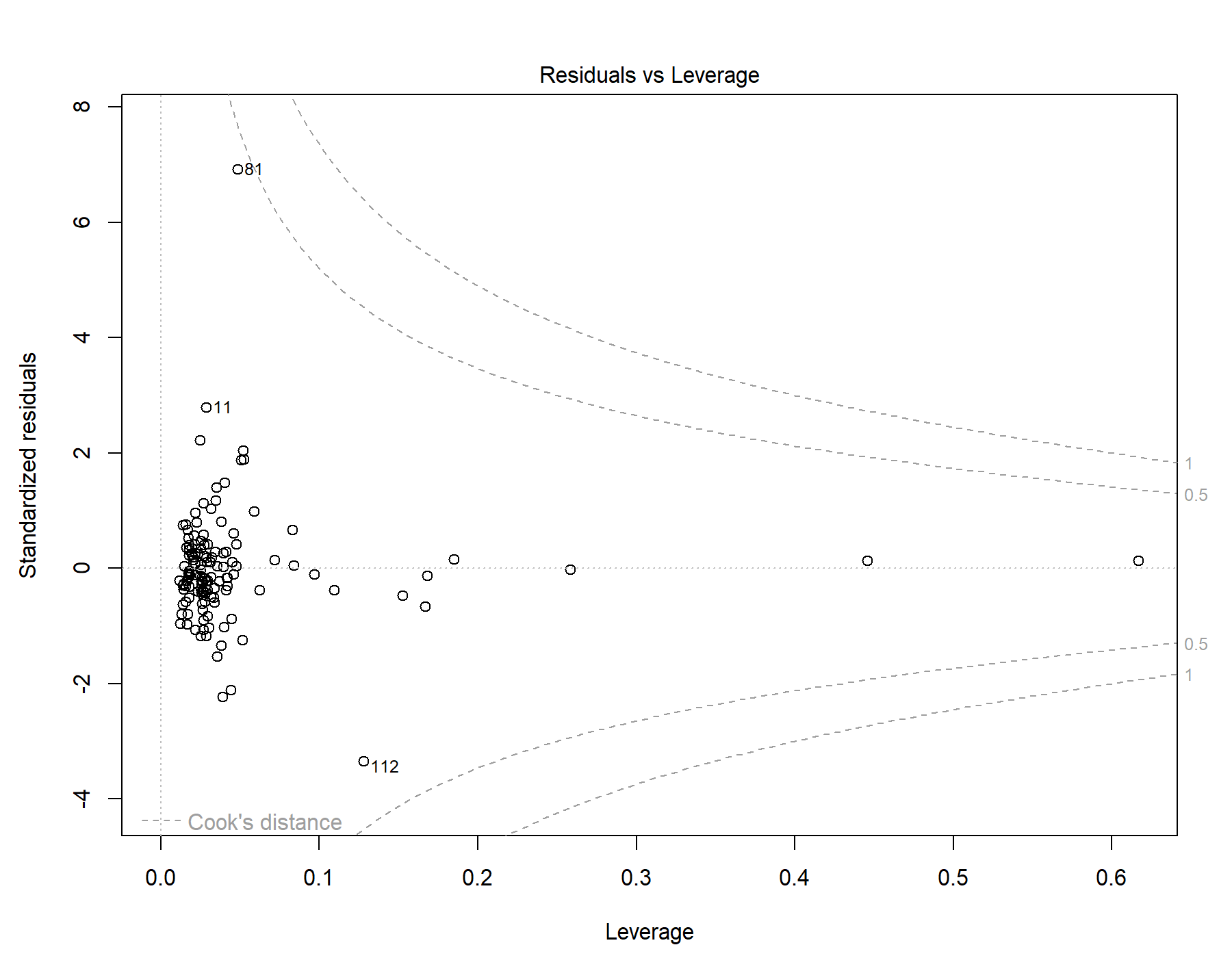

Supplemental Figure 26. Residuals vs leverage diagnostics for T1w+T2w WML segmentation.

### 12. Global | All subjects | T1w-alone WML vol | RMSSD | Primary interaction model

Model: T1w-alone WML vol ~ RMSSD + Age + Sex + Age *	RMSSD + Sex * RMSSD

#### 12.1. Standardized coefficients

| **Term** | **Estimate** | **Std. Error** | **t** | **p** | **β** |
| --- | --- | --- | --- | --- | --- |
| (Intercept) | -4.174e-04 | 1.870e-04 | -2.23 | .027 |  |
| RMSSD | 3.154e-06 | 5.555e-06 | 0.568 | .571 | 0.169 |
| Age | 1.519e-04 | 1.487e-05 | 10.2 | <.001 | 0.690 |
| Sex | -7.423e-05 | 1.121e-04 | -0.662 | .509 | -0.0441 |
| RMSSD:Age | -4.670e-07 | 3.004e-07 | -1.55 | .122 | -0.160 |
| RMSSD:Sex | -9.367e-07 | 3.222e-06 | -0.291 | .772 | -0.0926 |

***Supplemental Table 43.*** *Standardized coefficients values for T1w-alone WML segmentation.*

#### 12.2. 95% confidence intervals

| **Term** | **Lower 95%** | **Upper 95%** |
| --- | --- | --- |
| (Intercept) | -7.874e-04 | -4.729e-05 |
| RMSSD | -7.836e-06 | 1.415e-05 |
| Age | 1.225e-04 | 1.813e-04 |
| Sex | -2.961e-04 | 1.476e-04 |
| RMSSD:Age | -1.061e-06 | 1.273e-07 |
| RMSSD:Sex | -7.313e-06 | 5.440e-06 |

***Supplemental Table 44.*** *95% confidence intervals values for T1w-alone WML segmentation.*

#### 12.3. Breusch-Pagan test

| **Test** | **Statistic** | **df** | **p** |
| --- | --- | --- | --- |
| Breusch-Pagan | 20.303 | 5 | .001 |

***Supplemental Table 45.*** *Breusch-Pagan test values for T1w-alone WML segmentation.*

#### 12.4. Normal Q-Q plot

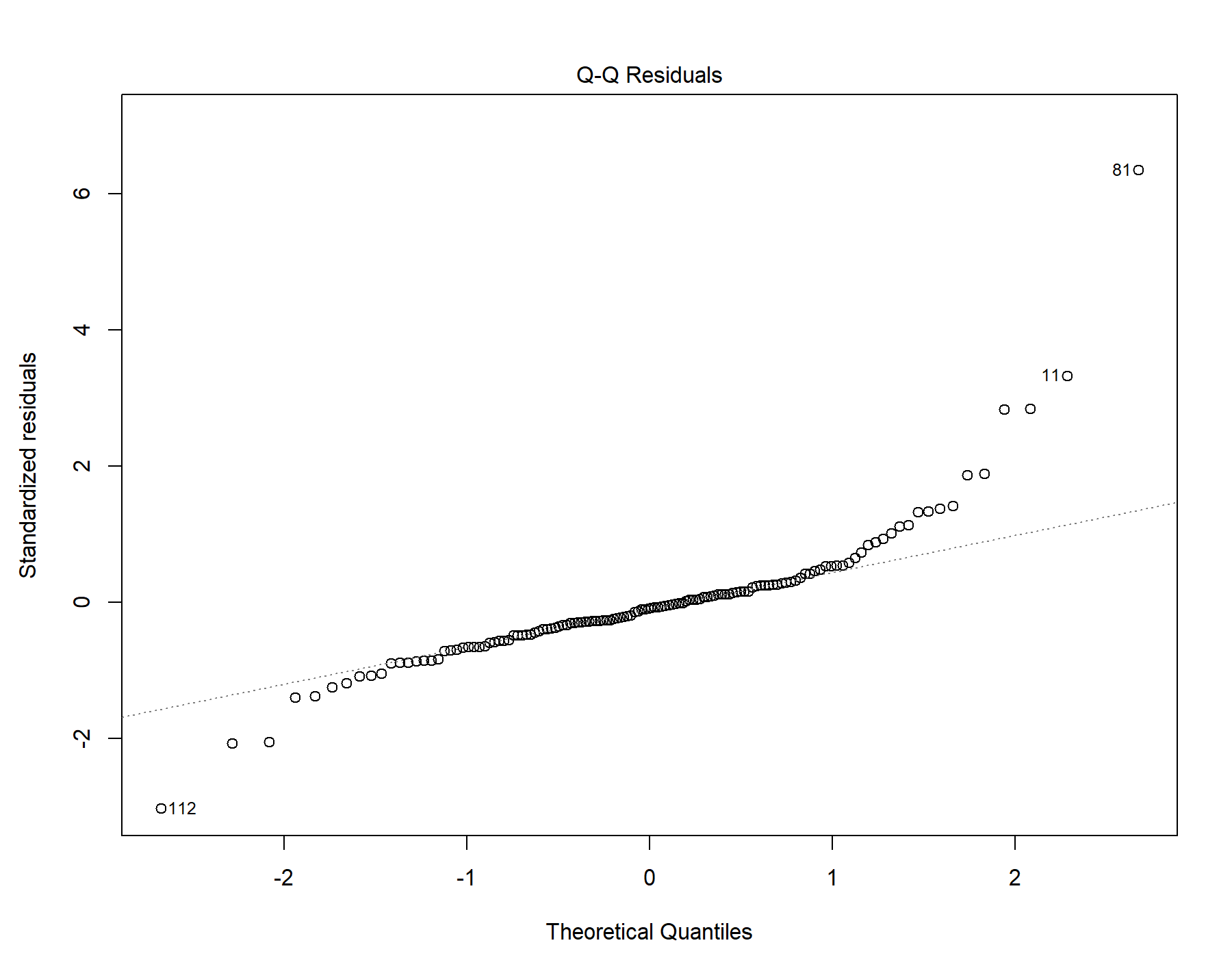

Supplemental Figure 27. Normal distribution quantile-quantile plot for T1w-alone WML segmentation.

#### 12.5. Residuals vs leverage

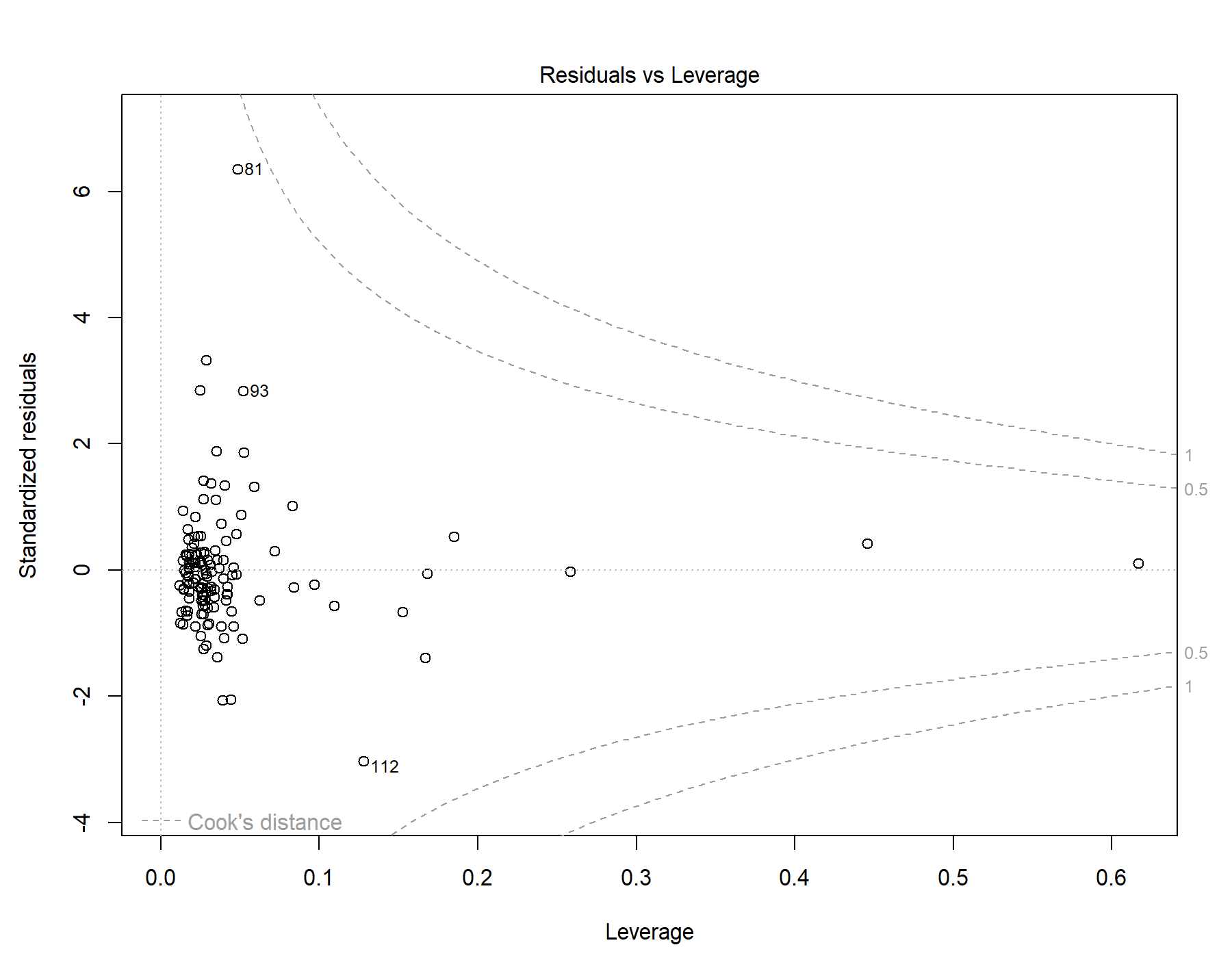

Supplemental Figure 28. Residuals vs leverage diagnostics for T1w-alone WML segmentation.

### 13. Global | All subjects | T1w+FLAIR WML vol | SDNN.LF._log | Primary interaction model

Model: T1w+FLAIR WML vol ~ SDNN.LF._log + Age + Sex + Age * SDNN.LF._log + Sex * SDNN.LF._log

#### 13.1. Standardized coefficients

| **Term** | **Estimate** | **Std. Error** | **t** | **p** | **β** |
| --- | --- | --- | --- | --- | --- |
| (Intercept) | -4.938e-04 | 1.967e-04 | -2.51 | .013 |  |
| SDNN.LF._log | -0.00104 | 6.162e-04 | -1.68 | .095 | -0.494 |
| Age | 1.572e-04 | 1.569e-05 | 10.0 | <.001 | 0.668 |
| Sex | -4.433e-05 | 1.176e-04 | -0.377 | .707 | -0.0246 |
| SDNN.LF._log:Age | -8.544e-05 | 3.299e-05 | -2.59 | .011 | -0.282 |
| SDNN.LF._log:Sex | 7.260e-04 | 3.562e-04 | 2.04 | .044 | 0.640 |

***Supplemental Table 46.*** *Standardized coefficients values for T1w+FLAIR WML segmentation.*

#### 13.2. 95% confidence intervals

| **Term** | **Lower 95%** | **Upper 95%** |
| --- | --- | --- |
| (Intercept) | -8.831e-04 | -1.045e-04 |
| SDNN.LF._log | -0.00226 | 1.816e-04 |
| Age | 1.262e-04 | 1.883e-04 |
| Sex | -2.770e-04 | 1.883e-04 |
| SDNN.LF._log:Age | -1.507e-04 | -2.017e-05 |
| SDNN.LF._log:Sex | 2.126e-05 | 0.00143 |

***Supplemental Table 47.*** *95% confidence intervals values for T1w+FLAIR WML segmentation.*

#### 13.3. Breusch-Pagan test

| **Test** | **Statistic** | **df** | **p** |
| --- | --- | --- | --- |
| Breusch-Pagan | 16.931 | 5 | .005 |

***Supplemental Table 48.*** *Breusch-Pagan test values for T1w+FLAIR WML segmentation.*

#### 13.4. Normal Q-Q plot

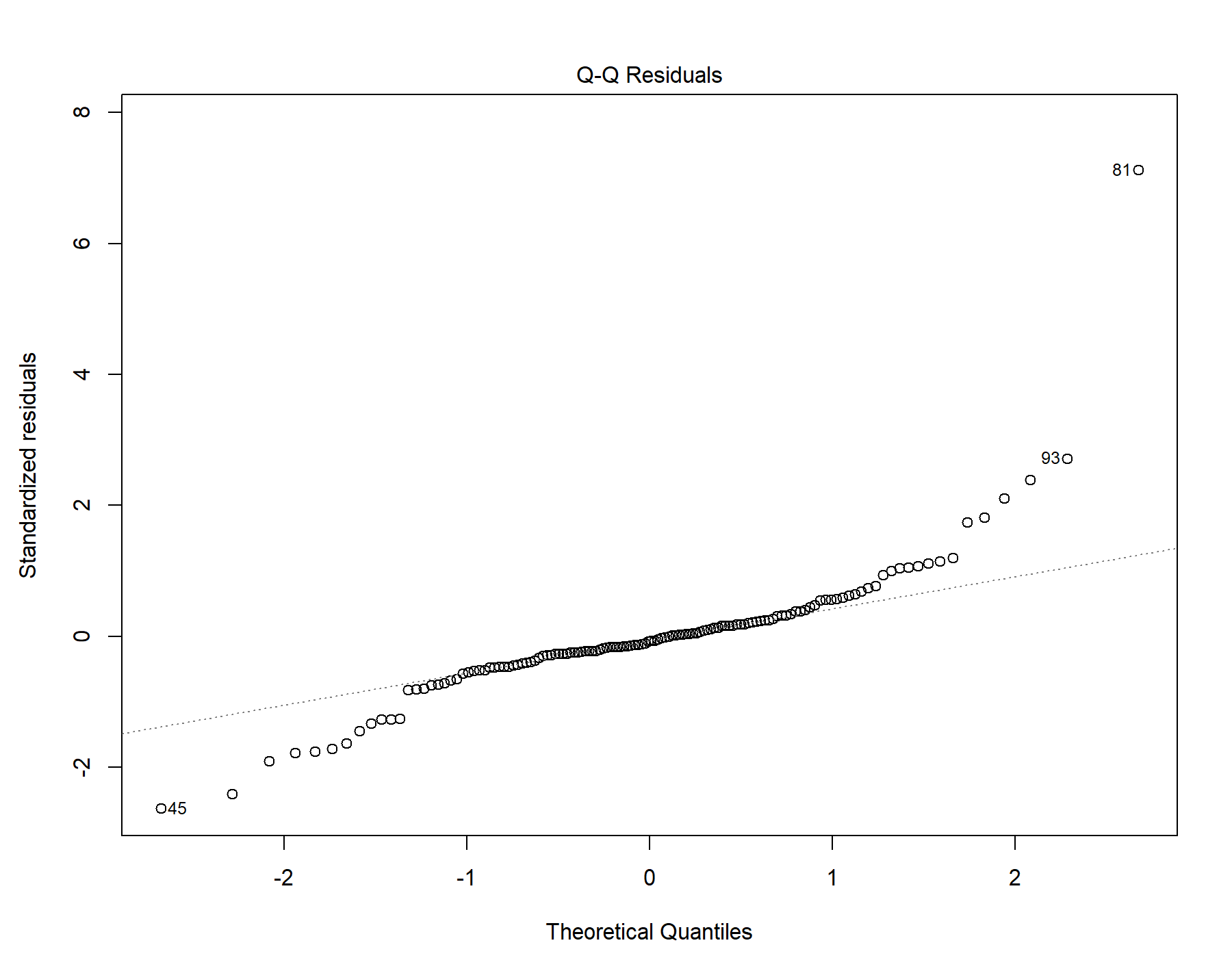

Supplemental Figure 29. Normal distribution quantile-quantile plot for T1w+FLAIR WML segmentation.

#### 13.5. Residuals vs leverage

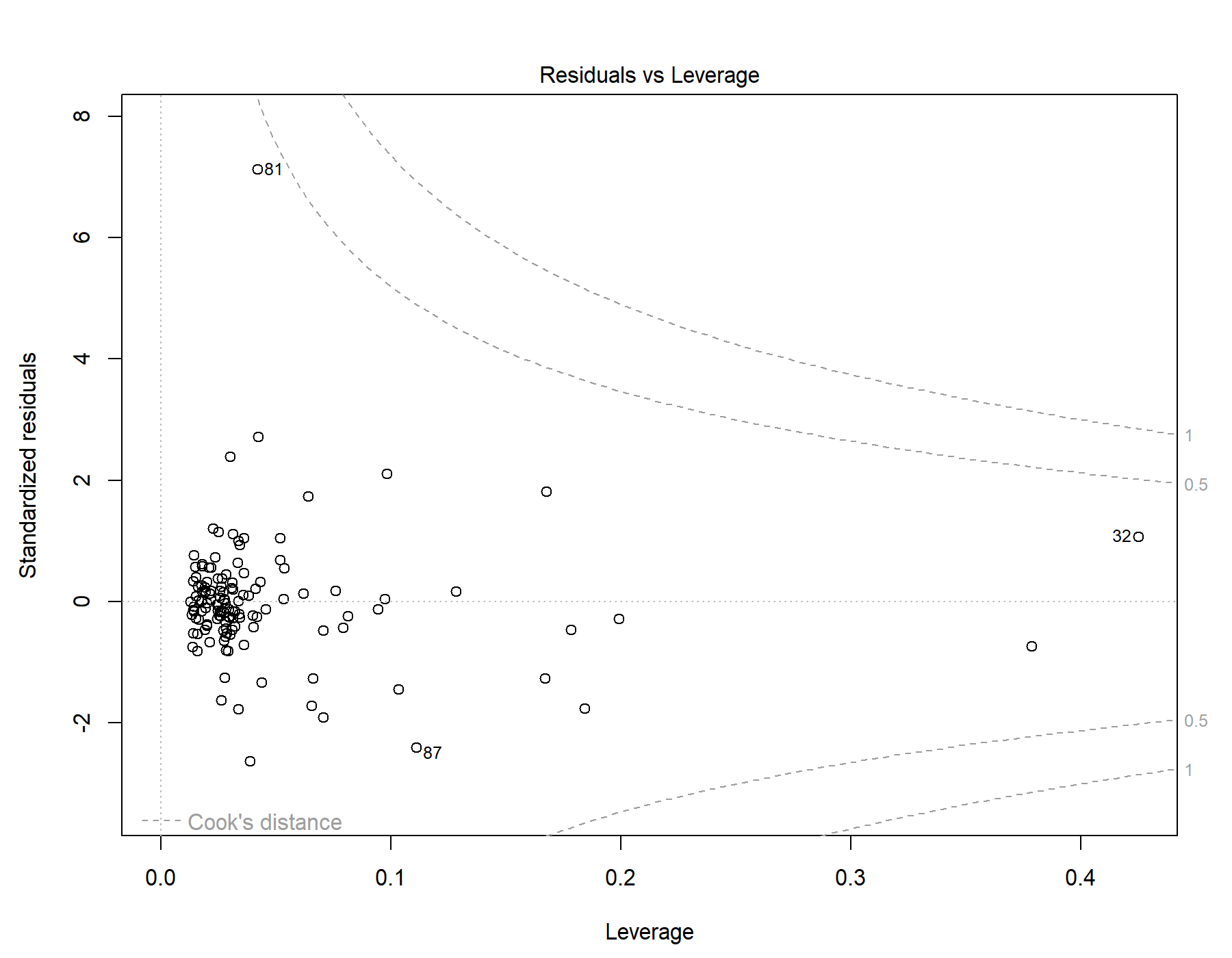

Supplemental Figure 30. Residuals vs leverage diagnostics for T1w+FLAIR WML segmentation.

### 14. Global | All subjects | T1w+T2w WML vol | SDNN.LF._log | Primary interaction model

Model: T1w+T2w WML vol ~ SDNN.LF._log + Age + Sex + Age * SDNN.LF._log + Sex * SDNN.LF._log

#### 14.1. Standardized coefficients

| **Term** | **Estimate** | **Std. Error** | **t** | **p** | **β** |
| --- | --- | --- | --- | --- | --- |
| (Intercept) | -5.236e-04 | 2.153e-04 | -2.43 | .016 |  |
| SDNN.LF._log | -8.105e-04 | 6.743e-04 | -1.20 | .232 | -0.340 |
| Age | 1.900e-04 | 1.717e-05 | 11.1 | <.001 | 0.710 |
| Sex | -9.635e-05 | 1.287e-04 | -0.749 | .455 | -0.0471 |
| SDNN.LF._log:Age | -9.542e-05 | 3.610e-05 | -2.64 | .009 | -0.277 |
| SDNN.LF._log:Sex | 6.502e-04 | 3.898e-04 | 1.67 | .098 | 0.504 |

***Supplemental Table 49.*** *Standardized coefficients values for T1w+T2w WML segmentation.*

#### 14.2. 95% confidence intervals

| **Term** | **Lower 95%** | **Upper 95%** |
| --- | --- | --- |
| (Intercept) | -9.496e-04 | -9.757e-05 |
| SDNN.LF._log | -0.00214 | 5.238e-04 |
| Age | 1.560e-04 | 2.240e-04 |
| Sex | -3.509e-04 | 1.582e-04 |
| SDNN.LF._log:Age | -1.669e-04 | -2.398e-05 |
| SDNN.LF._log:Sex | -1.210e-04 | 0.00142 |

***Supplemental Table 50.*** *95% confidence intervals values for T1w+T2w WML segmentation.*

#### 14.3. Breusch-Pagan test

| **Test** | **Statistic** | **df** | **p** |
| --- | --- | --- | --- |
| Breusch-Pagan | 13.815 | 5 | .017 |

***Supplemental Table 51.*** *Breusch-Pagan test values for T1w+T2w WML segmentation.*

#### 14.4. Normal Q-Q plot

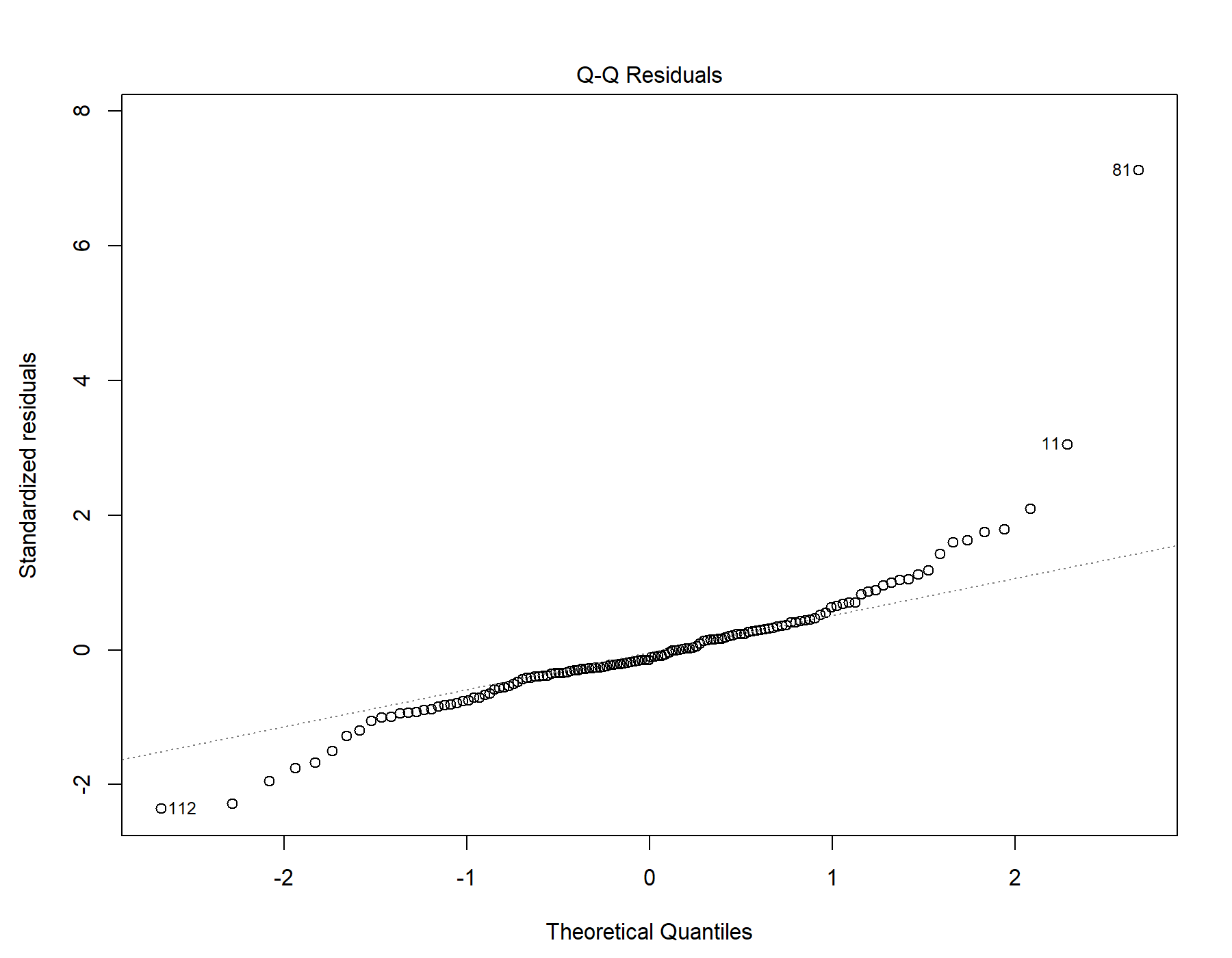

Supplemental Figure 31. Normal distribution quantile-quantile plot for T1w+T2w WML segmentation.

#### 14.5. Residuals vs leverage

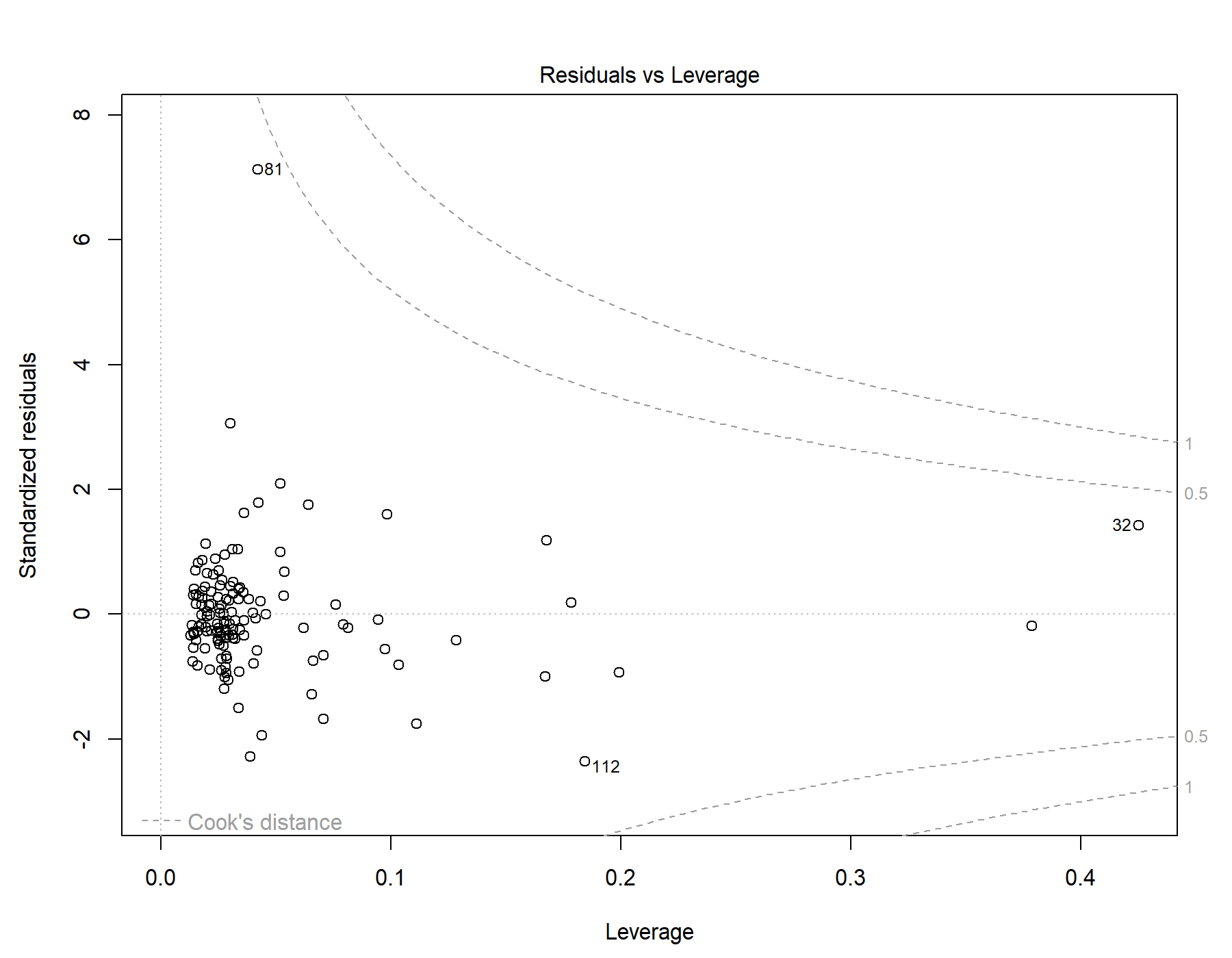

Supplemental Figure 32. Residuals vs leverage diagnostics for T1w+T2w WML segmentation.

### 15. Global | All subjects | T1w-alone WML vol | SDNN.LF._log | Primary interaction model

Model: T1w-alone WML vol ~ SDNN.LF._log + Age + Sex + Age * SDNN.LF._log + Sex * SDNN.LF._log

#### 15.1. Standardized coefficients

| **Term** | **Estimate** | **Std. Error** | **t** | **p** | **β** |
| --- | --- | --- | --- | --- | --- |
| (Intercept) | -4.012e-04 | 1.815e-04 | -2.21 | .029 |  |
| SDNN.LF._log | -6.430e-04 | 5.684e-04 | -1.13 | .260 | -0.328 |
| Age | 1.524e-04 | 1.448e-05 | 10.5 | <.001 | 0.693 |
| Sex | -8.889e-05 | 1.085e-04 | -0.820 | .414 | -0.0529 |
| SDNN.LF._log:Age | -7.904e-05 | 3.043e-05 | -2.60 | .011 | -0.279 |
| SDNN.LF._log:Sex | 5.180e-04 | 3.285e-04 | 1.58 | .117 | 0.489 |

***Supplemental Table 52.*** *Standardized coefficients values for T1w-alone WML segmentation.*

#### 15.2. 95% confidence intervals

| **Term** | **Lower 95%** | **Upper 95%** |
| --- | --- | --- |
| (Intercept) | -7.603e-04 | -4.212e-05 |
| SDNN.LF._log | -0.00177 | 4.817e-04 |
| Age | 1.238e-04 | 1.811e-04 |
| Sex | -3.035e-04 | 1.257e-04 |
| SDNN.LF._log:Age | -1.393e-04 | -1.882e-05 |
| SDNN.LF._log:Sex | -1.321e-04 | 0.00117 |

***Supplemental Table 53.*** *95% confidence intervals values for T1w-alone WML segmentation.*

#### 15.3. Breusch-Pagan test

| **Test** | **Statistic** | **df** | **p** |
| --- | --- | --- | --- |
| Breusch-Pagan | 15.515 | 5 | .008 |

***Supplemental Table 54.*** *Breusch-Pagan test values for T1w-alone WML segmentation.*

#### 15.4. Normal Q-Q plot

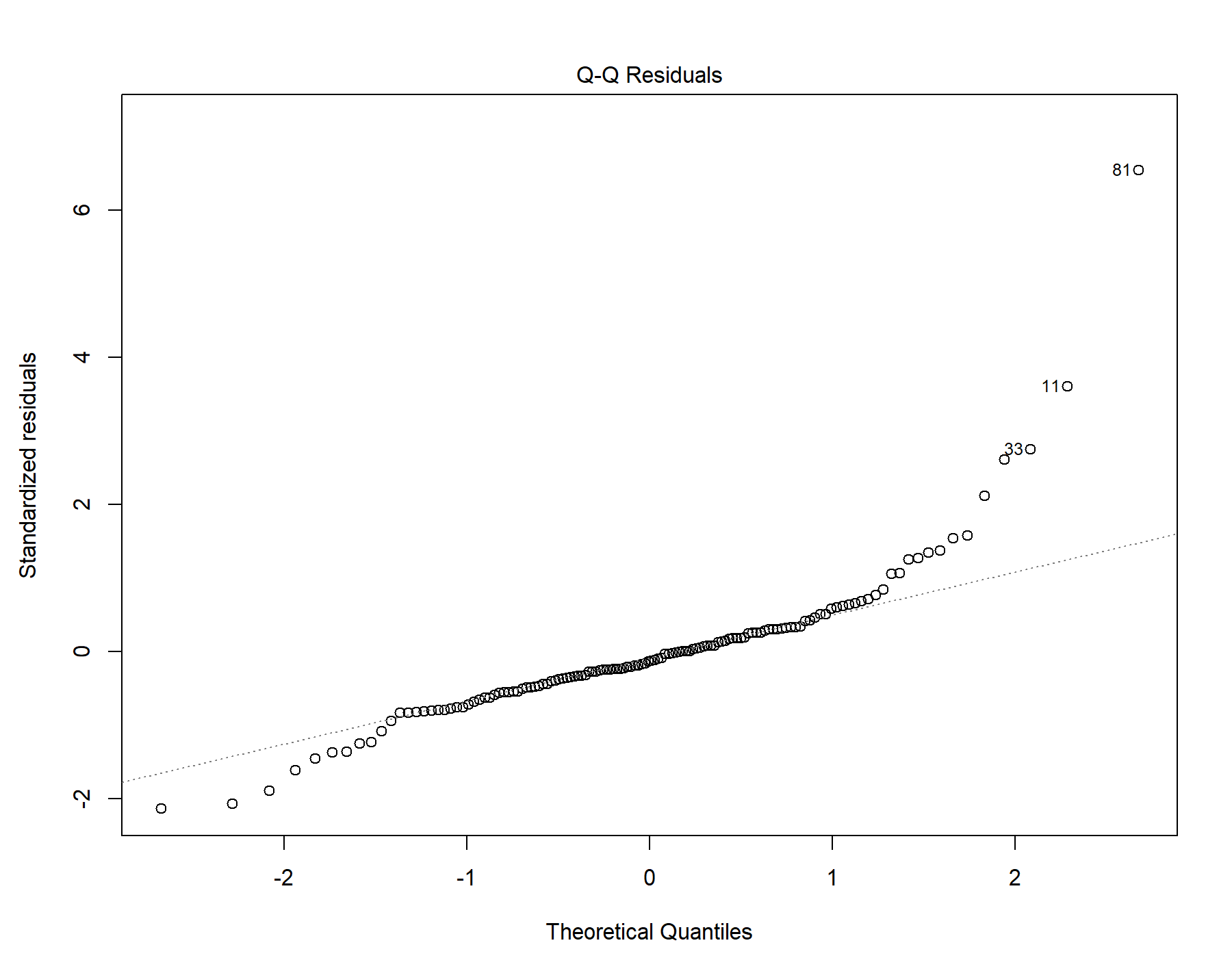

Supplemental Figure 33. Normal distribution quantile-quantile plot for T1w-alone WML segmentation.

#### 15.5. Residuals vs leverage

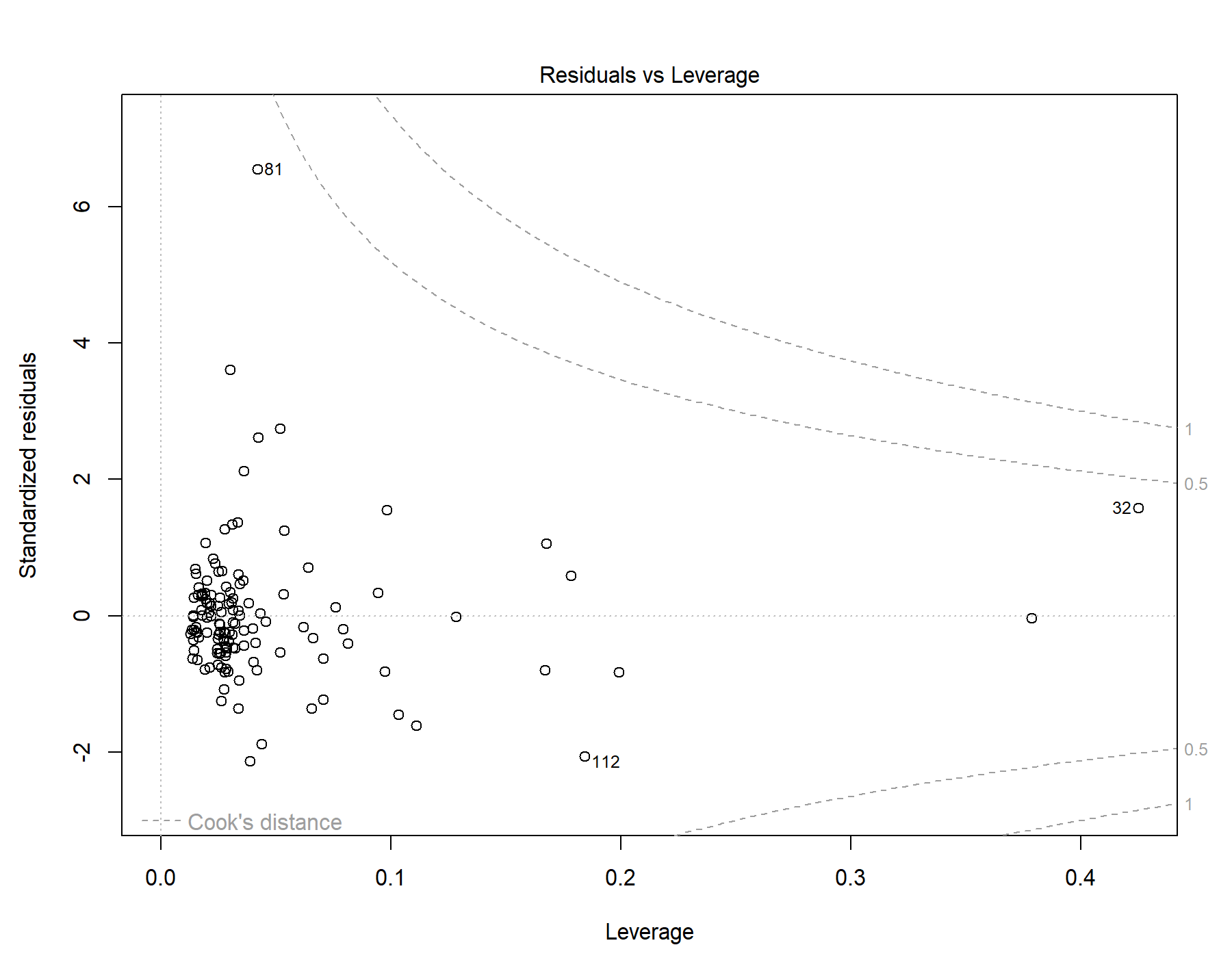

Supplemental Figure 34. Residuals vs leverage diagnostics for T1w-alone WML segmentation.

### 16. Global | All subjects | T1w+FLAIR WML vol | SDNN.HF._log | Primary interaction model

Model: T1w+FLAIR WML vol ~ SDNN.HF._log + Age + Sex + Age * SDNN.HF._log + Sex * SDNN.HF._log

#### 16.1. Standardized coefficients

| **Term** | **Estimate** | **Std. Error** | **t** | **p** | **β** |
| --- | --- | --- | --- | --- | --- |
| (Intercept) | -5.278e-04 | 1.997e-04 | -2.64 | .009 |  |
| SDNN.HF._log | -2.584e-04 | 6.458e-04 | -0.400 | .690 | -0.119 |
| Age | 1.481e-04 | 1.683e-05 | 8.80 | <.001 | 0.629 |
| Sex | -1.181e-05 | 1.204e-04 | -0.0980 | .922 | -0.00656 |
| SDNN.HF._log:Age | -6.663e-05 | 3.725e-05 | -1.79 | .076 | -0.209 |
| SDNN.HF._log:Sex | 2.066e-04 | 3.877e-04 | 0.533 | .595 | 0.175 |

***Supplemental Table 55.*** *Standardized coefficients values for T1w+FLAIR WML segmentation.*

#### 16.2. 95% confidence intervals

| **Term** | **Lower 95%** | **Upper 95%** |
| --- | --- | --- |
| (Intercept) | -9.230e-04 | -1.327e-04 |
| SDNN.HF._log | -0.00154 | 0.00102 |
| Age | 1.148e-04 | 1.814e-04 |
| Sex | -2.501e-04 | 2.264e-04 |
| SDNN.HF._log:Age | -1.403e-04 | 7.079e-06 |
| SDNN.HF._log:Sex | -5.606e-04 | 9.738e-04 |

***Supplemental Table 56.*** *95% confidence intervals values for T1w+FLAIR WML segmentation.*

#### 16.3. Breusch-Pagan test

| **Test** | **Statistic** | **df** | **p** |
| --- | --- | --- | --- |
| Breusch-Pagan | 17.013 | 5 | .004 |

***Supplemental Table 57.*** *Breusch-Pagan test values for T1w+FLAIR WML segmentation.*

#### 16.4. Normal Q-Q plot

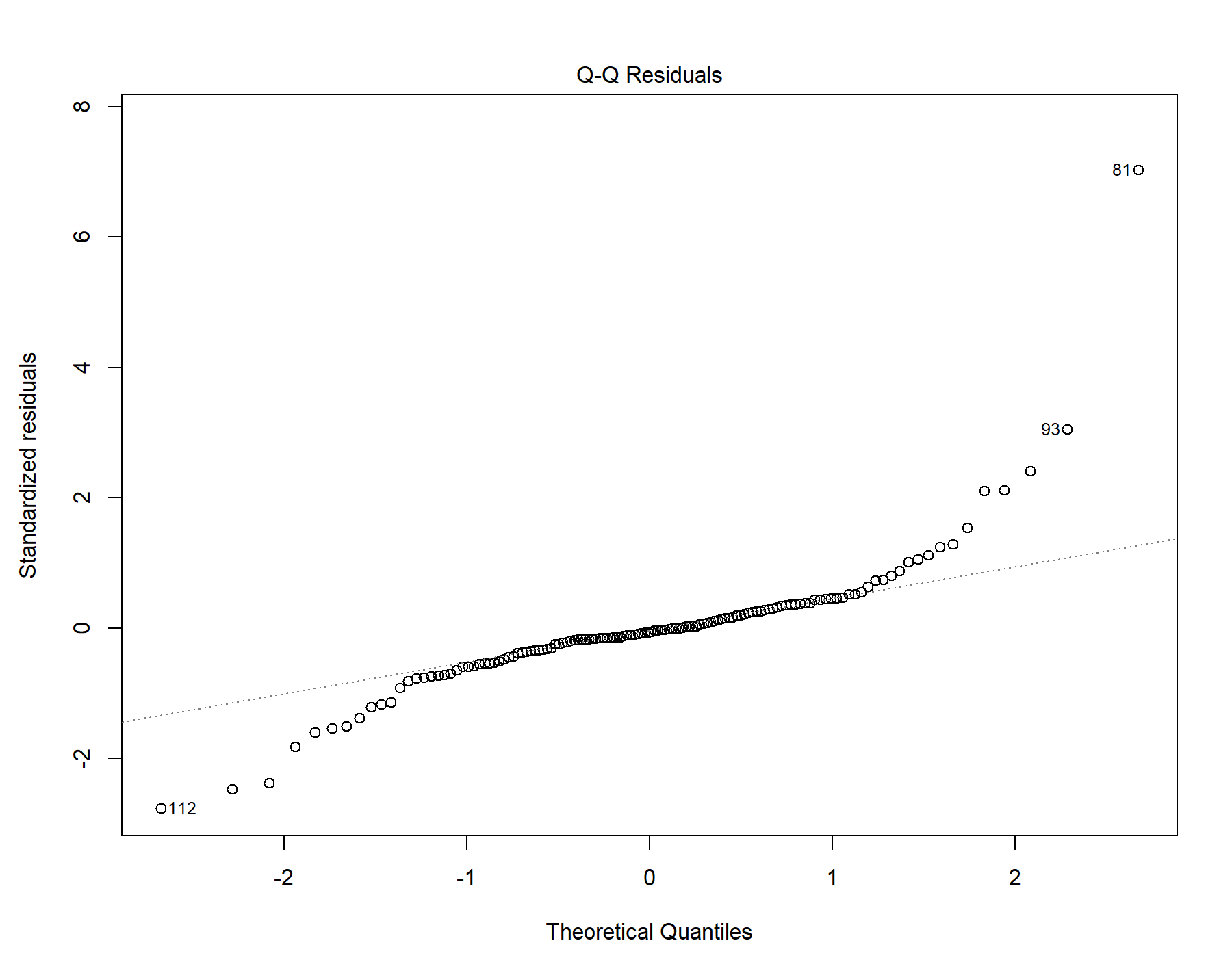

Supplemental Figure 35. Normal distribution quantile-quantile plot for T1w+FLAIR WML segmentation.

#### 16.5. Residuals vs leverage

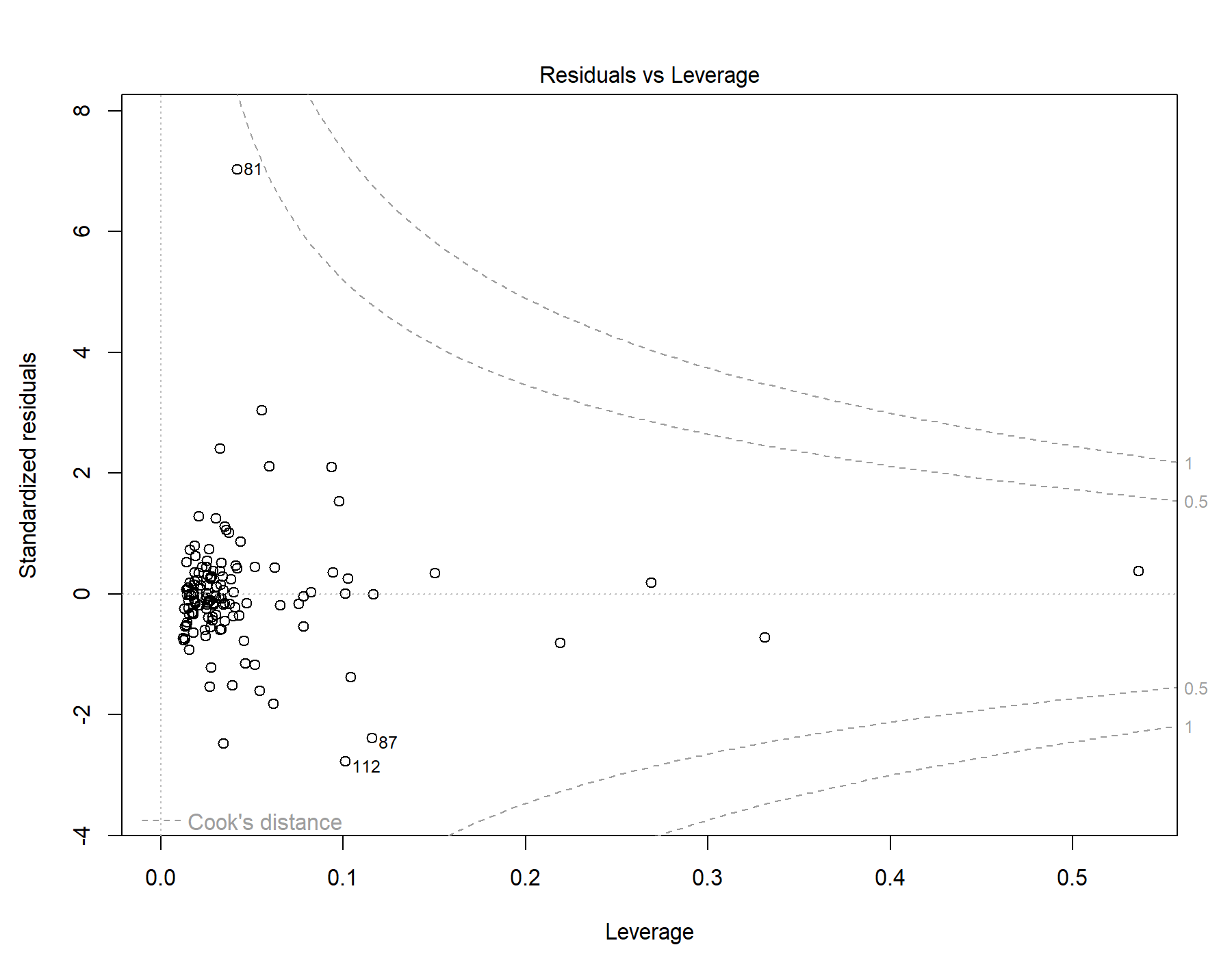

Supplemental Figure 36. Residuals vs leverage diagnostics for T1w+FLAIR WML segmentation.

### 17. Global | All subjects | T1w+T2w WML vol | SDNN.HF._log | Primary interaction model

Model: T1w+T2w WML vol ~ SDNN.HF._log + Age + Sex + Age * SDNN.HF._log + Sex * SDNN.HF._log

#### 17.1. Standardized coefficients

| **Term** | **Estimate** | **Std. Error** | **t** | **p** | **β** |
| --- | --- | --- | --- | --- | --- |
| (Intercept) | -5.494e-04 | 2.173e-04 | -2.53 | .013 |  |
| SDNN.HF._log | 2.392e-04 | 7.026e-04 | 0.340 | .734 | 0.0968 |
| Age | 1.796e-04 | 1.830e-05 | 9.81 | <.001 | 0.671 |
| Sex | -6.448e-05 | 1.310e-04 | -0.492 | .623 | -0.0315 |
| SDNN.HF._log:Age | -7.026e-05 | 4.052e-05 | -1.73 | .085 | -0.194 |
| SDNN.HF._log:Sex | -5.419e-05 | 4.218e-04 | -0.128 | .898 | -0.0405 |

***Supplemental Table 58.*** *Standardized coefficients values for T1w+T2w WML segmentation.*

#### 17.2. 95% confidence intervals

| **Term** | **Lower 95%** | **Upper 95%** |
| --- | --- | --- |
| (Intercept) | -9.793e-04 | -1.195e-04 |
| SDNN.HF._log | -0.00115 | 0.00163 |
| Age | 1.434e-04 | 2.158e-04 |
| Sex | -3.237e-04 | 1.947e-04 |
| SDNN.HF._log:Age | -1.504e-04 | 9.921e-06 |
| SDNN.HF._log:Sex | -8.888e-04 | 7.804e-04 |

***Supplemental Table 59.*** *95% confidence intervals values for T1w+T2w WML segmentation.*

#### 17.3. Breusch-Pagan test

| **Test** | **Statistic** | **df** | **p** |
| --- | --- | --- | --- |
| Breusch-Pagan | 14.528 | 5 | .013 |

***Supplemental Table 60.*** *Breusch-Pagan test values for T1w+T2w WML segmentation.*

#### 17.4. Normal Q-Q plot

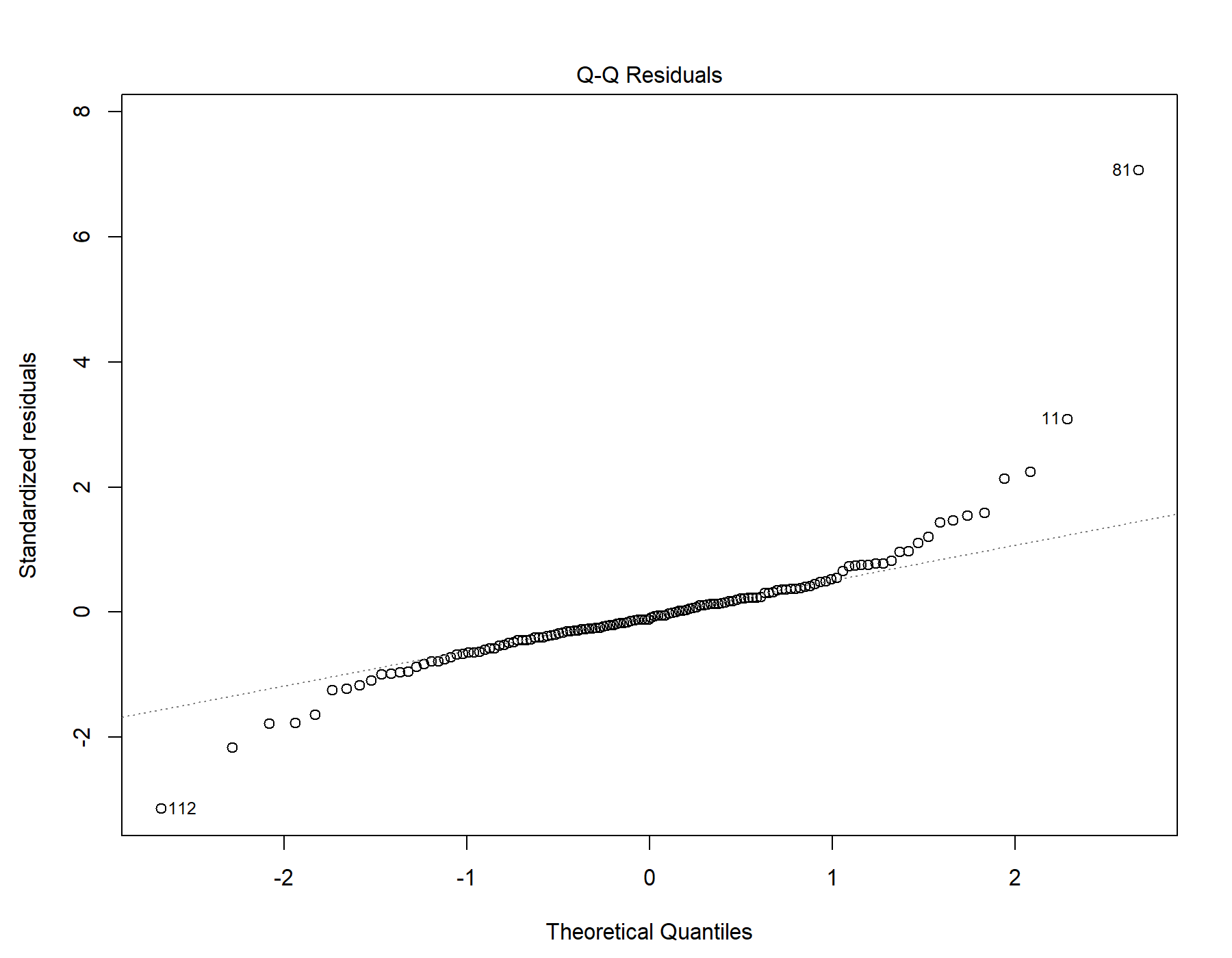

Supplemental Figure 37. Normal distribution quantile-quantile plot for T1w+T2w WML segmentation.

#### 17.5. Residuals vs leverage

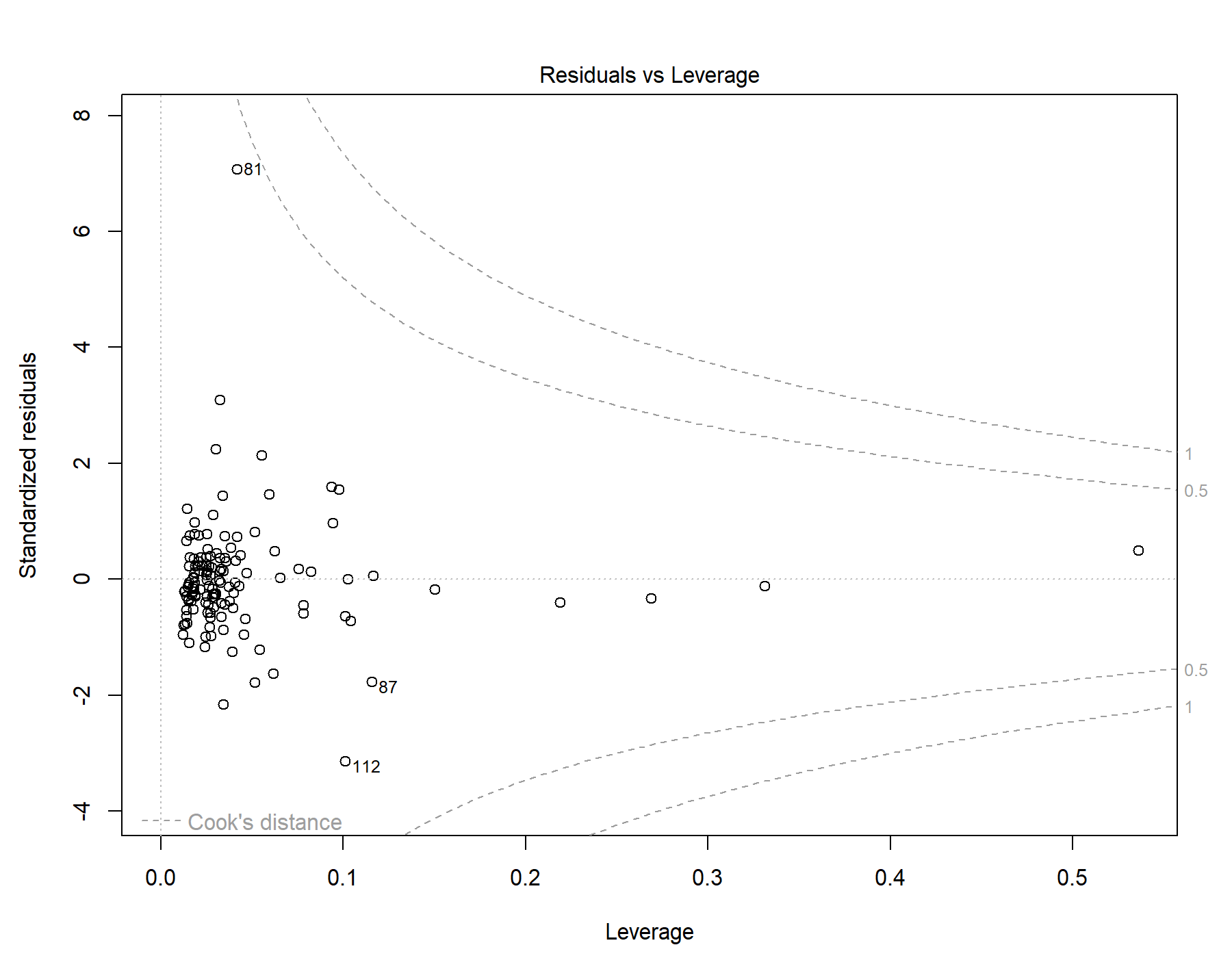

Supplemental Figure 38. Residuals vs leverage diagnostics for T1w+T2w WML segmentation.

### 18. Global | All subjects | T1w-alone WML vol | SDNN.HF._log | Primary interaction model

Model: T1w-alone WML vol ~ SDNN.HF._log + Age + Sex + Age * SDNN.HF._log + Sex * SDNN.HF._log

#### 18.1. Standardized coefficients

| **Term** | **Estimate** | **Std. Error** | **t** | **p** | **β** |
| --- | --- | --- | --- | --- | --- |
| (Intercept) | -4.239e-04 | 1.840e-04 | -2.30 | .023 |  |
| SDNN.HF._log | 1.063e-04 | 5.952e-04 | 0.179 | .859 | 0.0523 |
| Age | 1.456e-04 | 1.551e-05 | 9.39 | <.001 | 0.662 |
| Sex | -6.410e-05 | 1.110e-04 | -0.578 | .565 | -0.0381 |
| SDNN.HF._log:Age | -5.316e-05 | 3.433e-05 | -1.55 | .124 | -0.179 |
| SDNN.HF._log:Sex | 4.215e-06 | 3.573e-04 | 0.0118 | .991 | 0.00383 |

***Supplemental Table 61.*** *Standardized coefficients values for T1w-alone WML segmentation.*

#### 18.2. 95% confidence intervals

| **Term** | **Lower 95%** | **Upper 95%** |
| --- | --- | --- |
| (Intercept) | -7.880e-04 | -5.972e-05 |
| SDNN.HF._log | -0.00107 | 0.00128 |
| Age | 1.149e-04 | 1.763e-04 |
| Sex | -2.837e-04 | 1.555e-04 |
| SDNN.HF._log:Age | -1.211e-04 | 1.476e-05 |
| SDNN.HF._log:Sex | -7.028e-04 | 7.112e-04 |

***Supplemental Table 62.*** *95% confidence intervals values for T1w-alone WML segmentation.*

#### 18.3. Breusch-Pagan test

| **Test** | **Statistic** | **df** | **p** |
| --- | --- | --- | --- |
| Breusch-Pagan | 16.824 | 5 | .005 |

***Supplemental Table 63.*** *Breusch-Pagan test values for T1w-alone WML segmentation.*

#### 18.4. Normal Q-Q plot

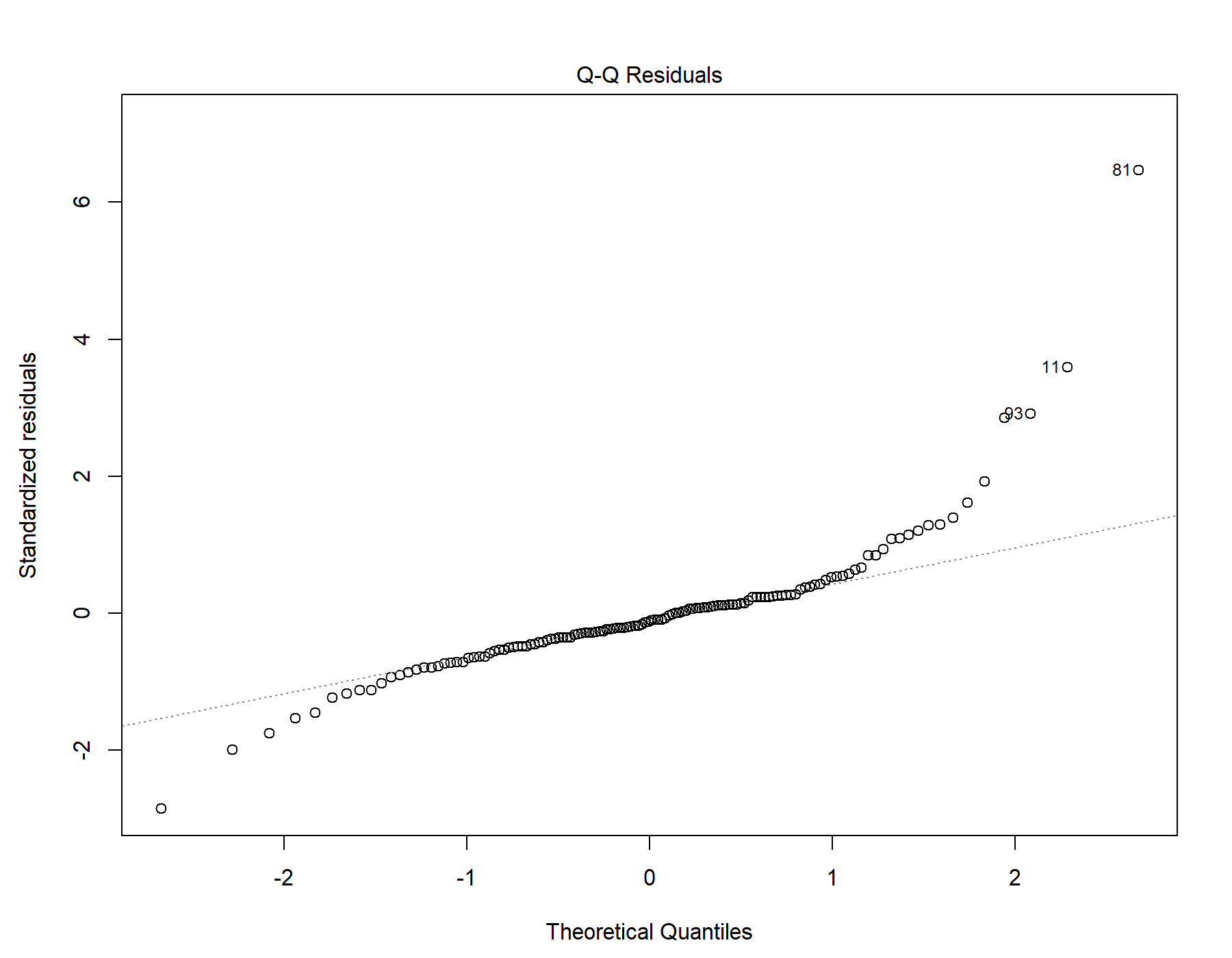

Supplemental Figure 39. Normal distribution quantile-quantile plot for T1w-alone WML segmentation.

#### 18.5. Residuals vs leverage

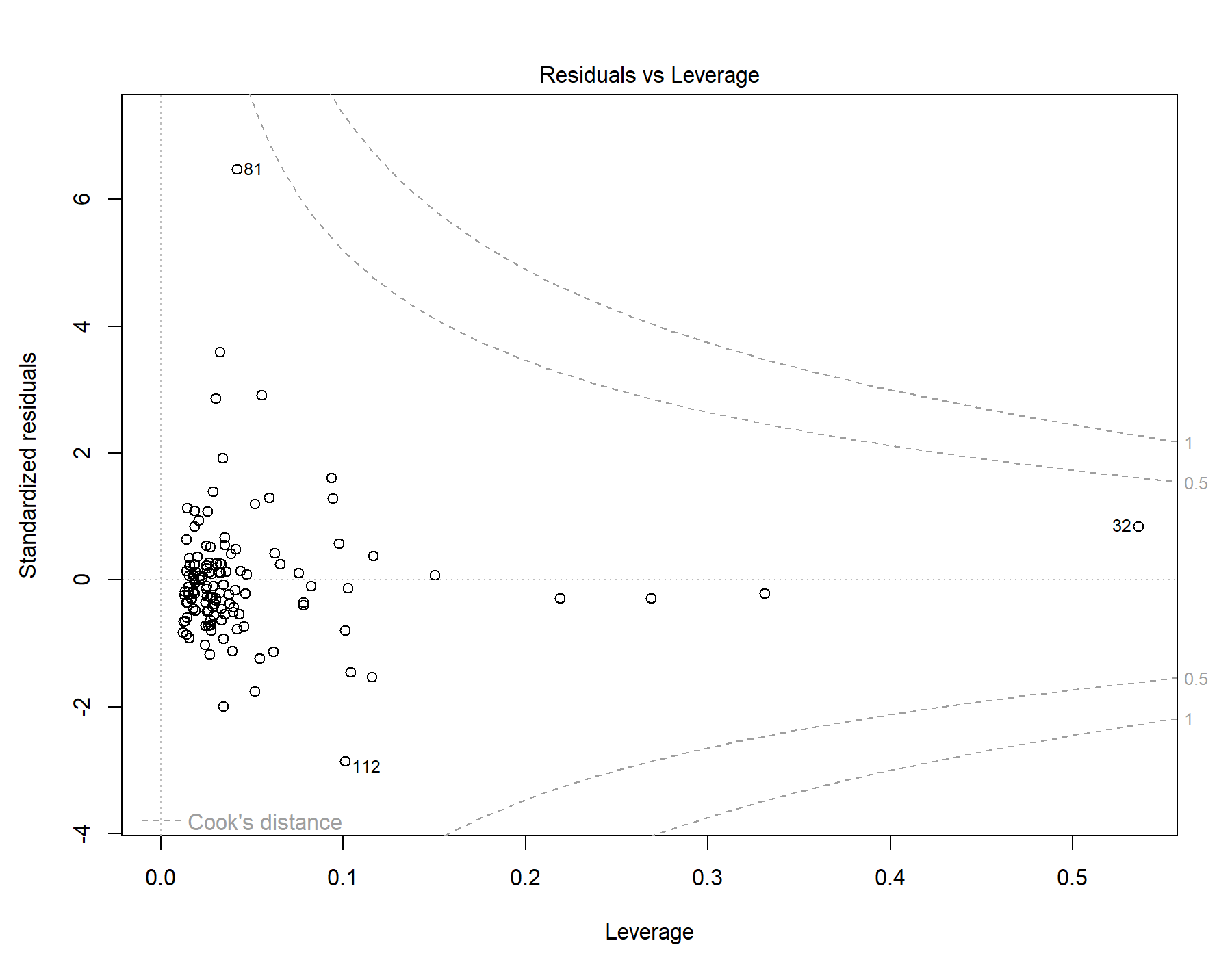

Supplemental Figure 40. Residuals vs leverage diagnostics for T1w-alone WML segmentation.

### 19. Global | Female only | T1w+FLAIR WML vol | RMSSD | Exploratory restricted interaction model

Model: T1w+FLAIR WML vol ~ RMSSD + Age + Age *	RMSSD

#### 19.1. Standardized coefficients

| **Term** | **Estimate** | **Std. Error** | **t** | **p** | **β** |
| --- | --- | --- | --- | --- | --- |
| (Intercept) | -4.499e-04 | 1.235e-04 | -3.64 | <.001 |  |
| RMSSD | -1.044e-05 | 3.294e-06 | -3.17 | .003 | -0.872 |
| Age | 3.055e-04 | 7.789e-05 | 3.92 | <.001 | 1.47 |
| RMSSD:Age | 9.277e-06 | 2.843e-06 | 3.26 | .002 | 1.58 |

***Supplemental Table 64.*** *Standardized coefficients values for T1w+FLAIR WML segmentation.*

#### 19.2. 95% confidence intervals

| **Term** | **Lower 95%** | **Upper 95%** |
| --- | --- | --- |
| (Intercept) | -7.003e-04 | -1.996e-04 |
| RMSSD | -1.712e-05 | -3.769e-06 |
| Age | 1.477e-04 | 4.633e-04 |
| RMSSD:Age | 3.516e-06 | 1.504e-05 |

***Supplemental Table 65.*** *95% confidence intervals values for T1w+FLAIR WML segmentation.*

#### 19.3. Breusch-Pagan test

| **Test** | **Statistic** | **df** | **p** |
| --- | --- | --- | --- |
| Breusch-Pagan | 2.013 | 3 | .570 |

***Supplemental Table 66.*** *Breusch-Pagan test values for T1w+FLAIR WML segmentation.*

#### 19.4. Normal Q-Q plot

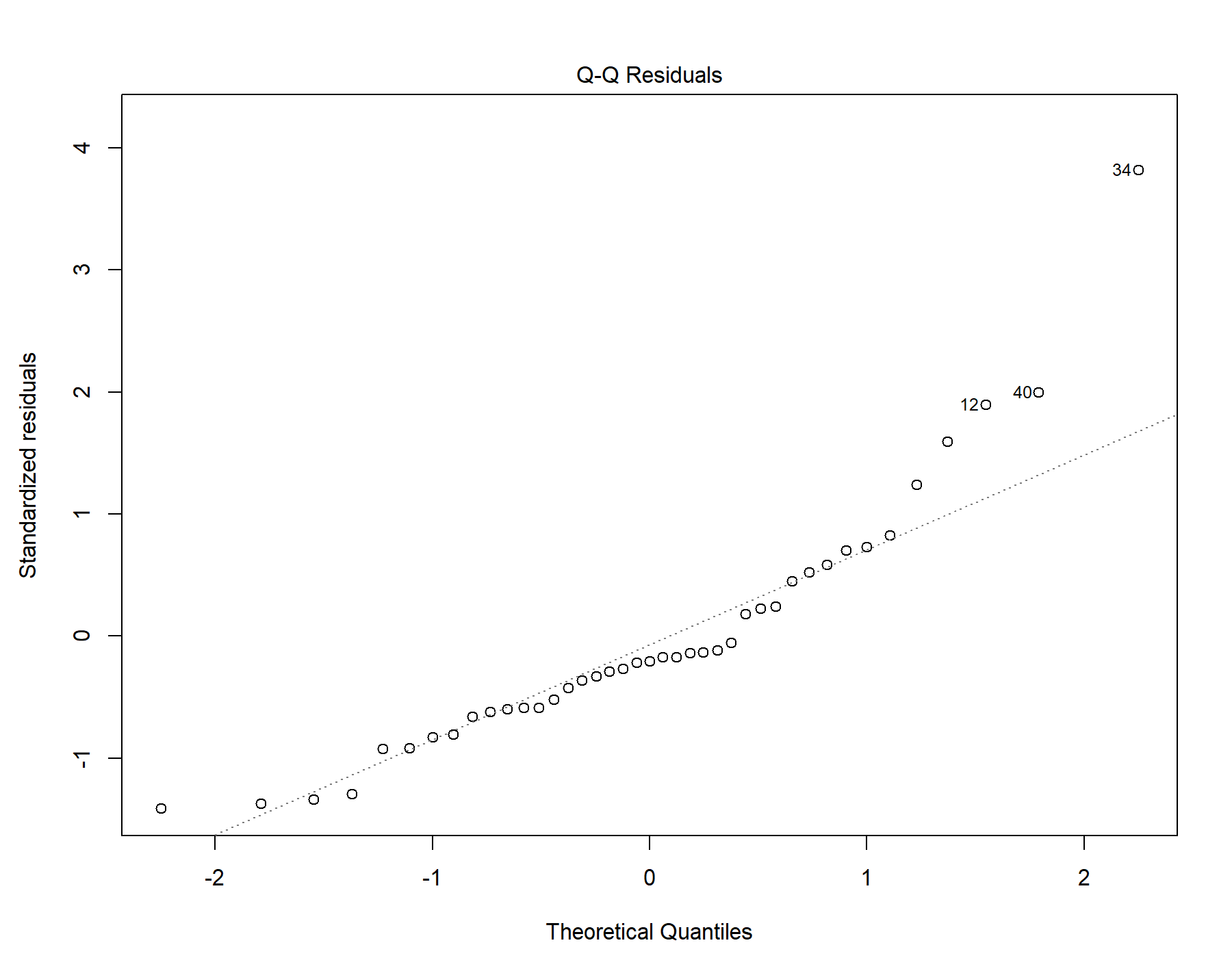

Supplemental Figure 41. Normal distribution quantile-quantile plot for T1w+FLAIR WML segmentation.

#### 19.5. Residuals vs leverage

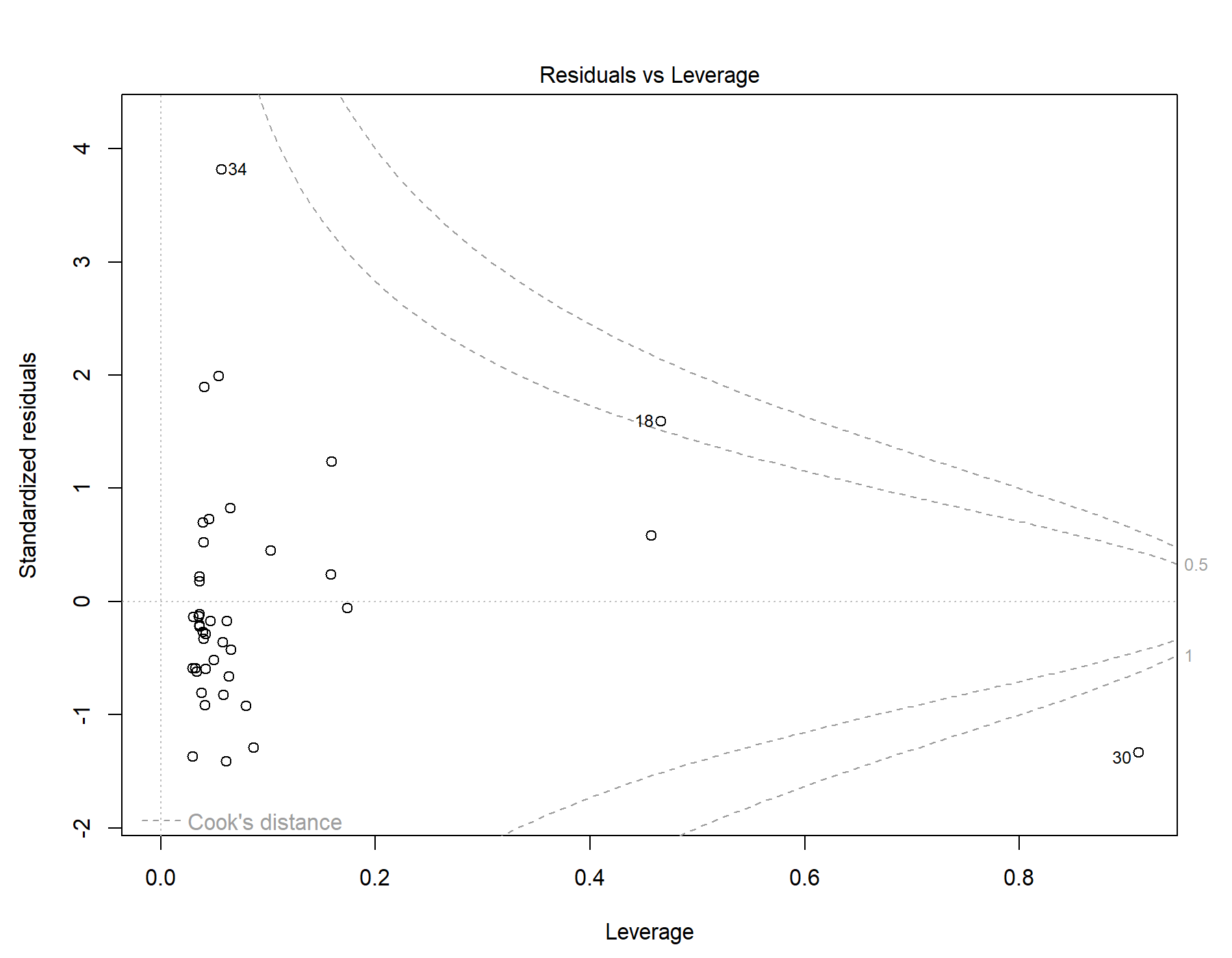

Supplemental Figure 42. Residuals vs leverage diagnostics for T1w+FLAIR WML segmentation.

### 20. Global | Female only | T1w+T2w WML vol | RMSSD | Exploratory restricted interaction model

Model: T1w+T2w WML vol ~ RMSSD + Age + Age *	RMSSD

#### 20.1. Standardized coefficients

| **Term** | **Estimate** | **Std. Error** | **t** | **p** | **β** |
| --- | --- | --- | --- | --- | --- |
| (Intercept) | -5.665e-04 | 1.534e-04 | -3.69 | <.001 |  |
| RMSSD | -1.073e-05 | 4.091e-06 | -2.62 | .013 | -0.730 |
| Age | 3.833e-04 | 9.673e-05 | 3.96 | <.001 | 1.50 |
| RMSSD:Age | 1.151e-05 | 3.531e-06 | 3.26 | .002 | 1.60 |

***Supplemental Table 67.*** *Standardized coefficients values for T1w+T2w WML segmentation.*

#### 20.2. 95% confidence intervals

| **Term** | **Lower 95%** | **Upper 95%** |
| --- | --- | --- |
| (Intercept) | -8.774e-04 | -2.556e-04 |
| RMSSD | -1.902e-05 | -2.445e-06 |
| Age | 1.873e-04 | 5.793e-04 |
| RMSSD:Age | 4.352e-06 | 1.866e-05 |

***Supplemental Table 68.*** *95% confidence intervals values for T1w+T2w WML segmentation.*

#### 20.3. Breusch-Pagan test

| **Test** | **Statistic** | **df** | **p** |
| --- | --- | --- | --- |
| Breusch-Pagan | 2.47 | 3 | .481 |

***Supplemental Table 69.*** *Breusch-Pagan test values for T1w+T2w WML segmentation.*

#### 20.4. Normal Q-Q plot

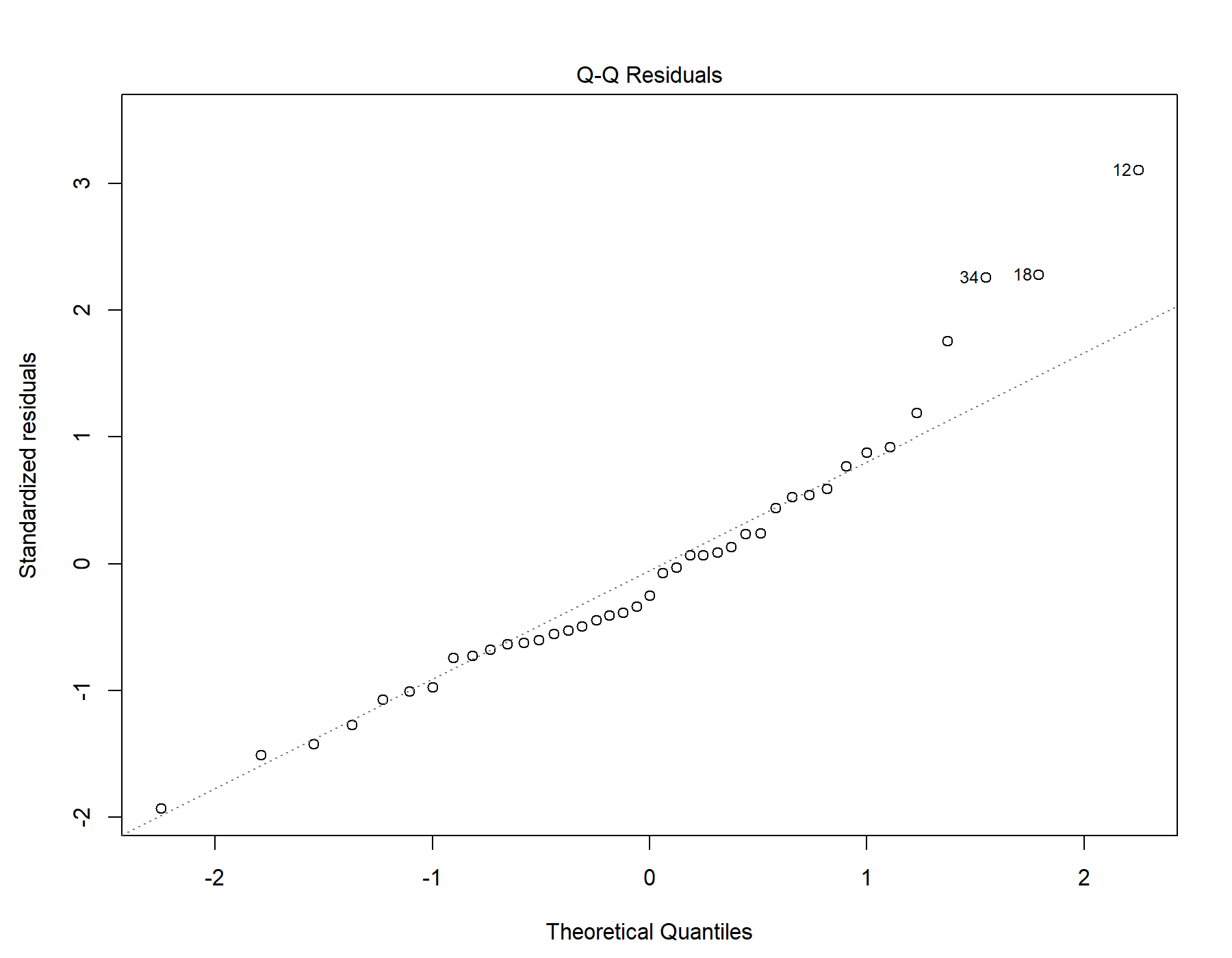

Supplemental Figure 43. Normal distribution quantile-quantile plot for T1w+T2w WML segmentation.

#### 20.5. Residuals vs leverage

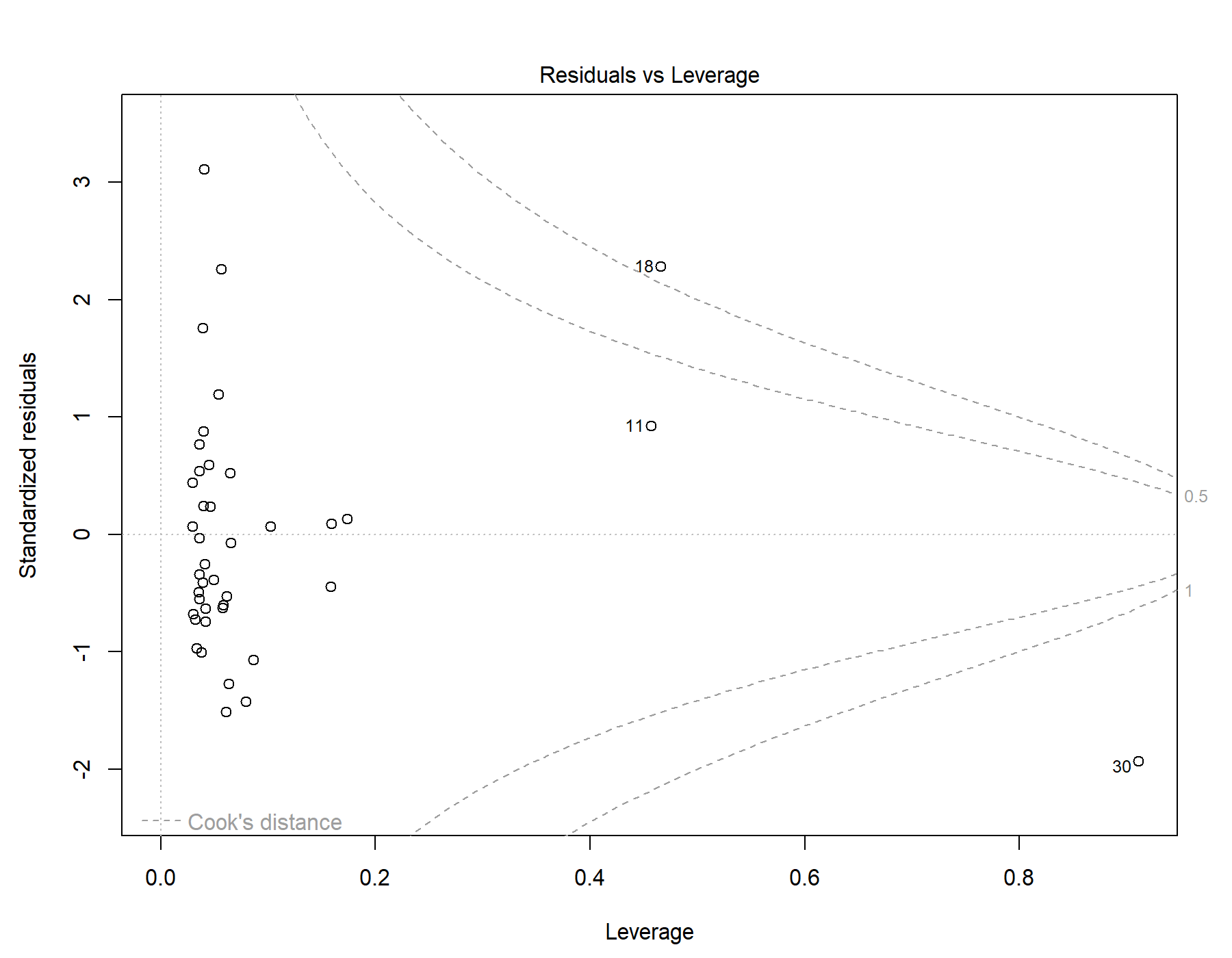

Supplemental Figure 44. Residuals vs leverage diagnostics for T1w+T2w WML segmentation.

### 21. Global | Female only | T1w-alone WML vol | RMSSD | Exploratory restricted interaction model

Model: T1w-alone WML vol ~ RMSSD + Age + Age *	RMSSD

#### 21.1. Standardized coefficients

| **Term** | **Estimate** | **Std. Error** | **t** | **p** | **β** |
| --- | --- | --- | --- | --- | --- |
| (Intercept) | -4.390e-04 | 1.543e-04 | -2.85 | .007 |  |
| RMSSD | -7.822e-06 | 4.113e-06 | -1.90 | .065 | -0.564 |
| Age | 2.927e-04 | 9.724e-05 | 3.01 | .005 | 1.21 |
| RMSSD:Age | 8.361e-06 | 3.550e-06 | 2.36 | .024 | 1.23 |

***Supplemental Table 70.*** *Standardized coefficients values for T1w-alone WML segmentation.*

#### 21.2. 95% confidence intervals

| **Term** | **Lower 95%** | **Upper 95%** |
| --- | --- | --- |
| (Intercept) | -7.515e-04 | -1.264e-04 |
| RMSSD | -1.615e-05 | 5.114e-07 |
| Age | 9.570e-05 | 4.898e-04 |
| RMSSD:Age | 1.169e-06 | 1.555e-05 |

***Supplemental Table 71.*** *95% confidence intervals values for T1w-alone WML segmentation.*

#### 21.3. Breusch-Pagan test

| **Test** | **Statistic** | **df** | **p** |
| --- | --- | --- | --- |
| Breusch-Pagan | 2.796 | 3 | .424 |

***Supplemental Table 72.*** *Breusch-Pagan test values for T1w-alone WML segmentation.*

#### 21.4. Normal Q-Q plot

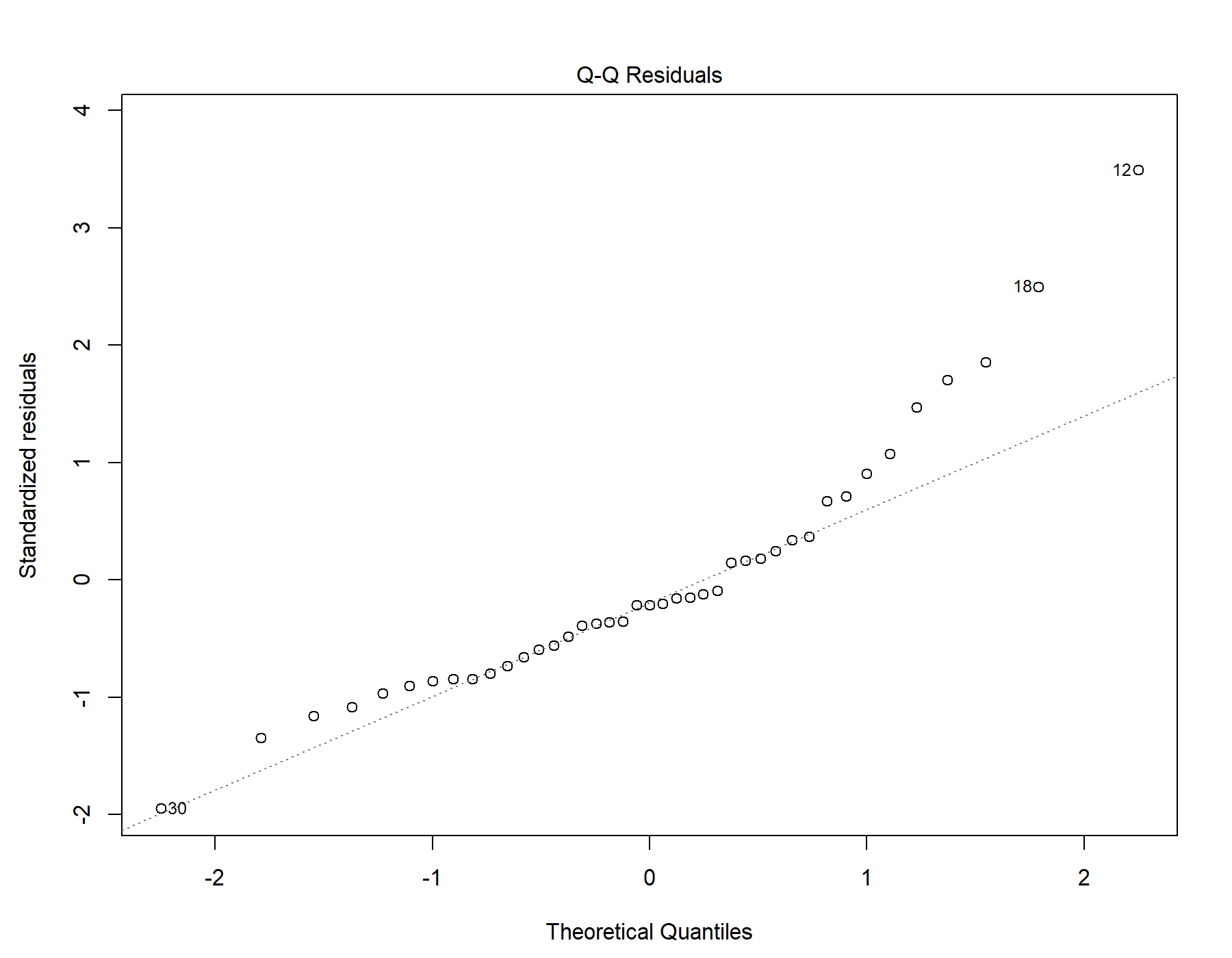

Supplemental Figure 45. Normal distribution quantile-quantile plot for T1w-alone WML segmentation.

#### 21.5. Residuals vs leverage

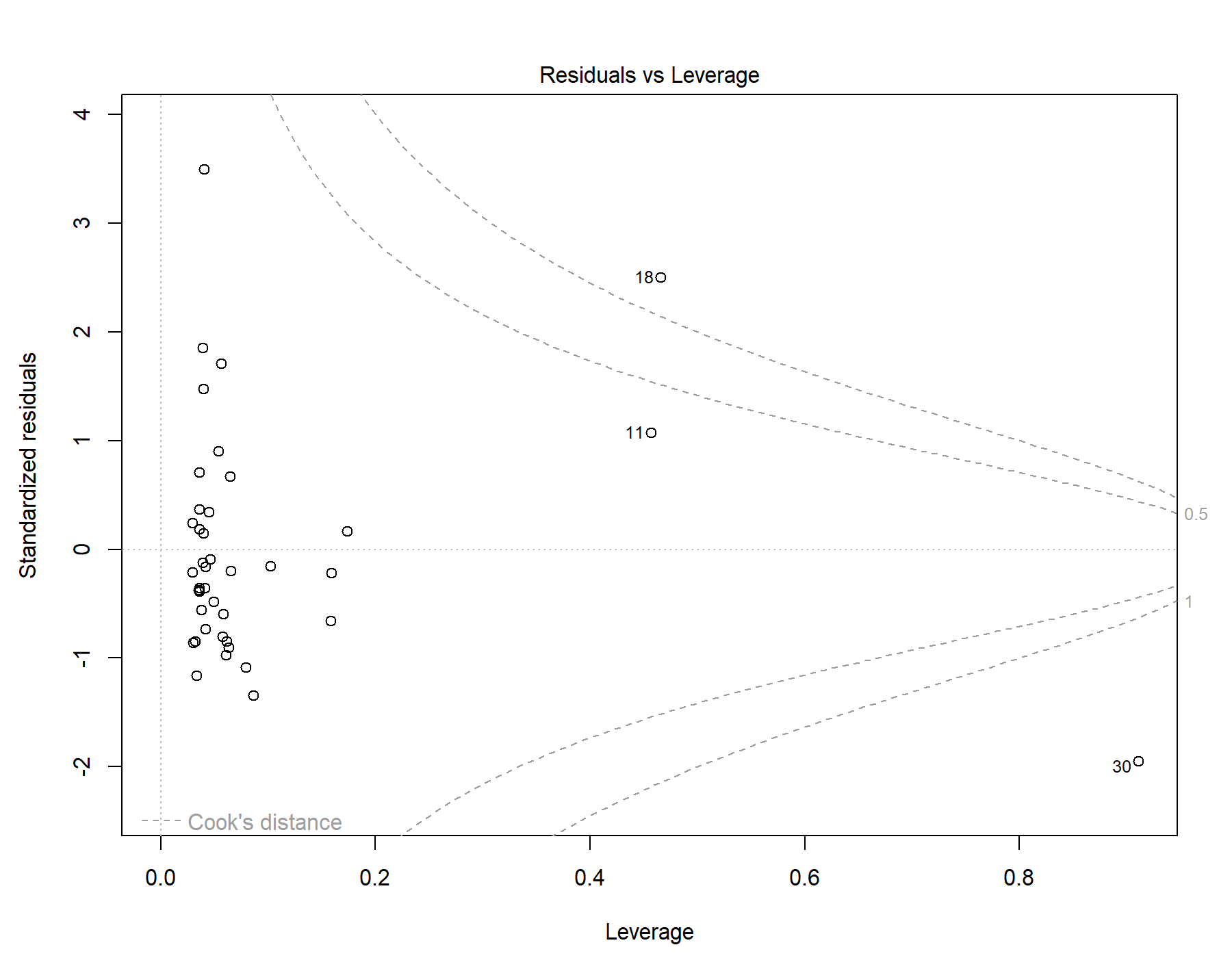

Supplemental Figure 46. Residuals vs leverage diagnostics for T1w-alone WML segmentation.

### 22. Global | Female only | T1w+FLAIR WML vol | SDNN.LF._log | Exploratory restricted interaction model

Model: T1w+FLAIR WML vol ~ SDNN.LF._log + Age +	Age * SDNN.LF._log

#### 22.1. Standardized coefficients

| **Term** | **Estimate** | **Std. Error** | **t** | **p** | **β** |
| --- | --- | --- | --- | --- | --- |
| (Intercept) | -1.642e-04 | 7.275e-05 | -2.26 | .030 |  |
| SDNN.LF._log | -2.247e-04 | 2.114e-04 | -1.06 | .295 | -0.179 |
| Age | 8.266e-05 | 2.724e-05 | 3.03 | .004 | 0.396 |
| SDNN.LF._log:Age | -1.342e-04 | 5.796e-05 | -2.32 | .026 | -0.392 |

***Supplemental Table 73.*** *Standardized coefficients values for T1w+FLAIR WML segmentation.*

#### 22.2. 95% confidence intervals

| **Term** | **Lower 95%** | **Upper 95%** |
| --- | --- | --- |
| (Intercept) | -3.116e-04 | -1.679e-05 |
| SDNN.LF._log | -6.531e-04 | 2.037e-04 |
| Age | 2.747e-05 | 1.378e-04 |
| SDNN.LF._log:Age | -2.516e-04 | -1.676e-05 |

***Supplemental Table 74.*** *95% confidence intervals values for T1w+FLAIR WML segmentation.*

#### 22.3. Breusch-Pagan test

| **Test** | **Statistic** | **df** | **p** |
| --- | --- | --- | --- |
| Breusch-Pagan | 4.489 | 3 | .213 |

***Supplemental Table 75.*** *Breusch-Pagan test values for T1w+FLAIR WML segmentation.*

#### 22.4. Normal Q-Q plot

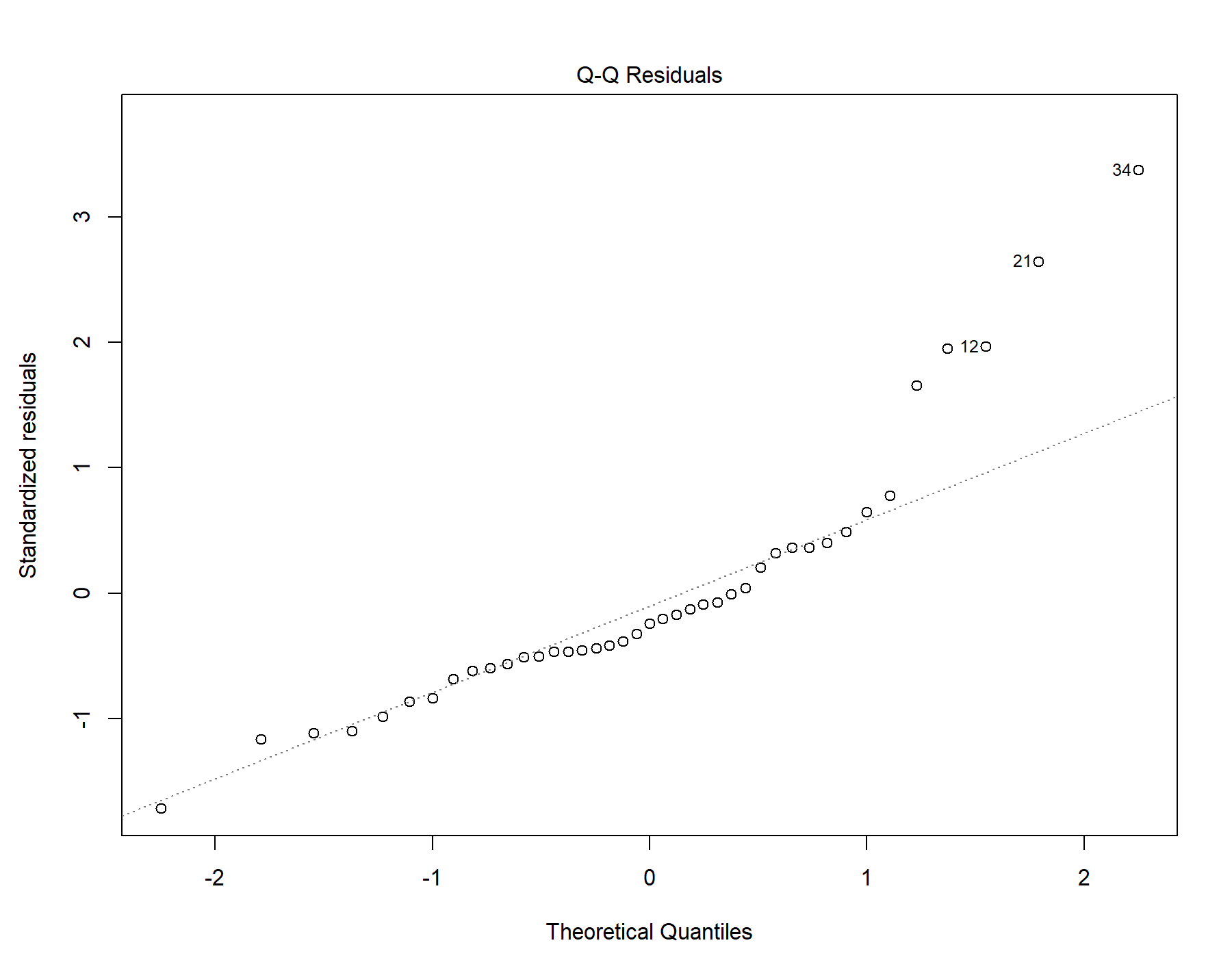

Supplemental Figure 47. Normal distribution quantile-quantile plot for T1w+FLAIR WML segmentation.

#### 22.5. Residuals vs leverage

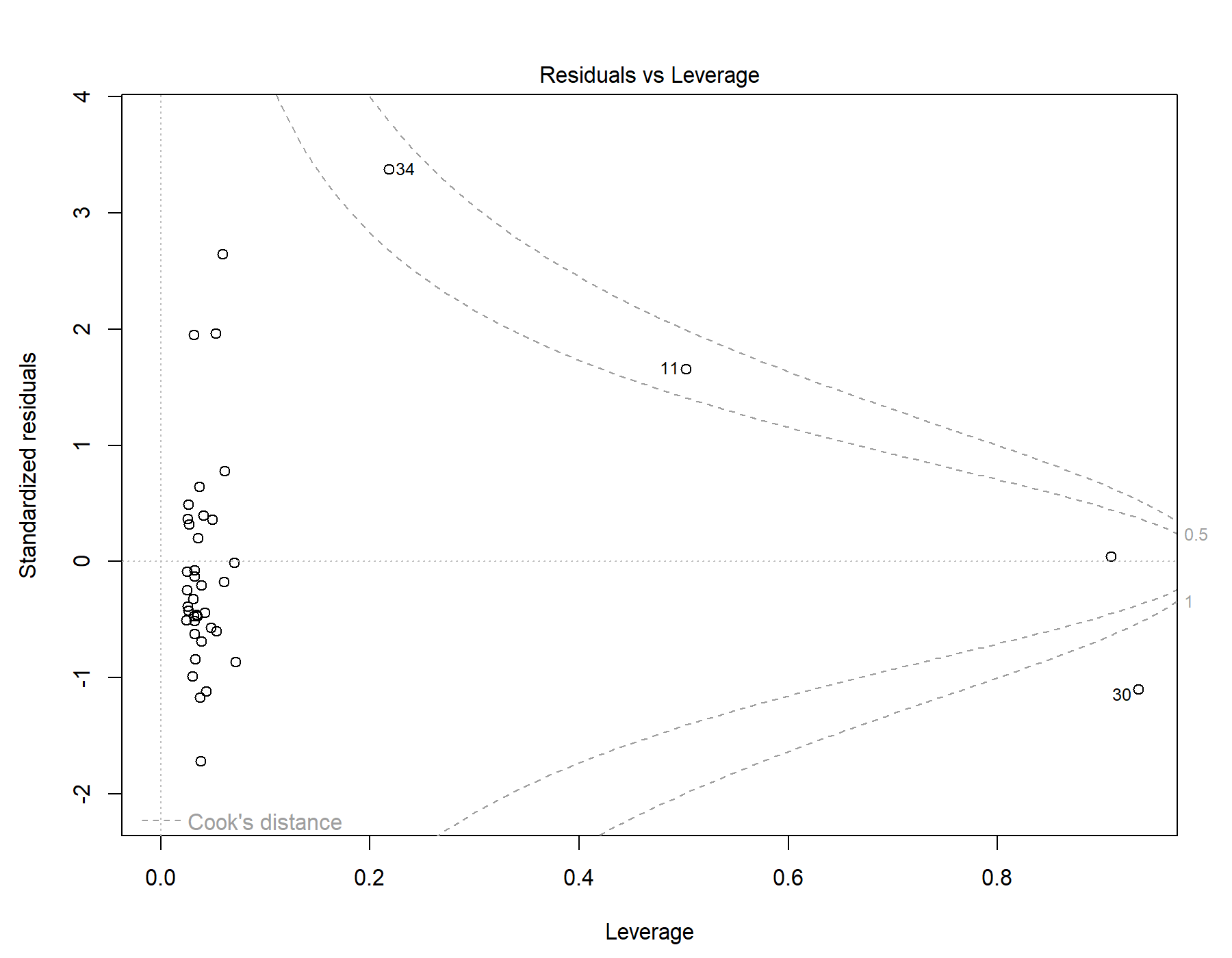

Supplemental Figure 48. Residuals vs leverage diagnostics for T1w+FLAIR WML segmentation.

### 23. Global | Female only | T1w+T2w WML vol | SDNN.LF._log | Exploratory restricted interaction model

Model: T1w+T2w WML vol ~ SDNN.LF._log + Age +	Age * SDNN.LF._log

#### 23.1. Standardized coefficients

| **Term** | **Estimate** | **Std. Error** | **t** | **p** | **β** |
| --- | --- | --- | --- | --- | --- |
| (Intercept) | -2.196e-04 | 8.971e-05 | -2.45 | .019 |  |
| SDNN.LF._log | 1.825e-04 | 2.607e-04 | 0.700 | .488 | 0.118 |
| Age | 1.092e-04 | 3.359e-05 | 3.25 | .002 | 0.426 |
| SDNN.LF._log:Age | -2.543e-04 | 7.148e-05 | -3.56 | .001 | -0.605 |

***Supplemental Table 76.*** *Standardized coefficients values for T1w+T2w WML segmentation.*

#### 23.2. 95% confidence intervals

| **Term** | **Lower 95%** | **Upper 95%** |
| --- | --- | --- |
| (Intercept) | -4.014e-04 | -3.787e-05 |
| SDNN.LF._log | -3.458e-04 | 7.108e-04 |
| Age | 4.110e-05 | 1.772e-04 |
| SDNN.LF._log:Age | -3.992e-04 | -1.095e-04 |

***Supplemental Table 77.*** *95% confidence intervals values for T1w+T2w WML segmentation.*

#### 23.3. Breusch-Pagan test

| **Test** | **Statistic** | **df** | **p** |
| --- | --- | --- | --- |
| Breusch-Pagan | 3.722 | 3 | .293 |

***Supplemental Table 78.*** *Breusch-Pagan test values for T1w+T2w WML segmentation.*

#### 23.4. Normal Q-Q plot

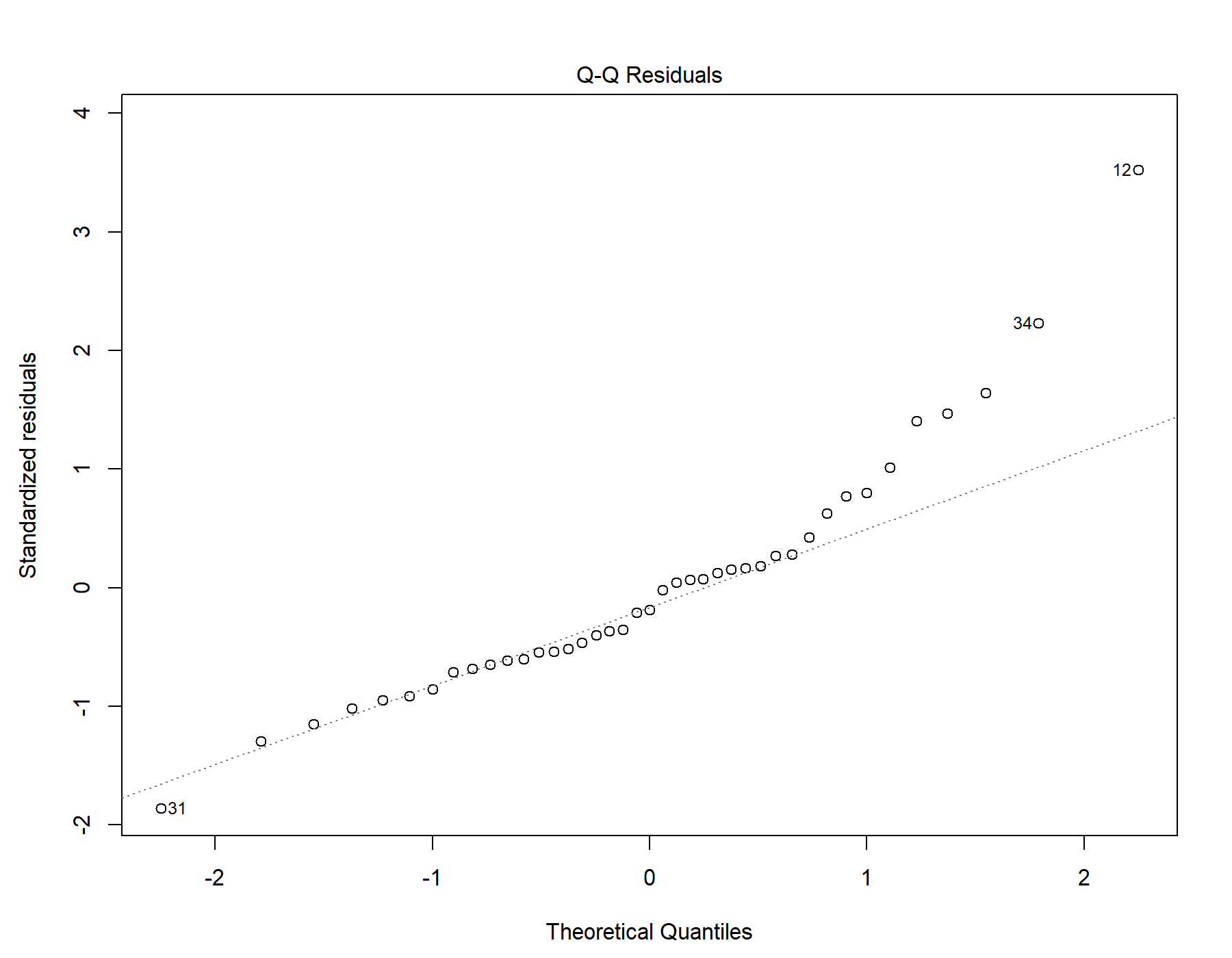

Supplemental Figure 49. Normal distribution quantile-quantile plot for T1w+T2w WML segmentation.

#### 23.5. Residuals vs leverage

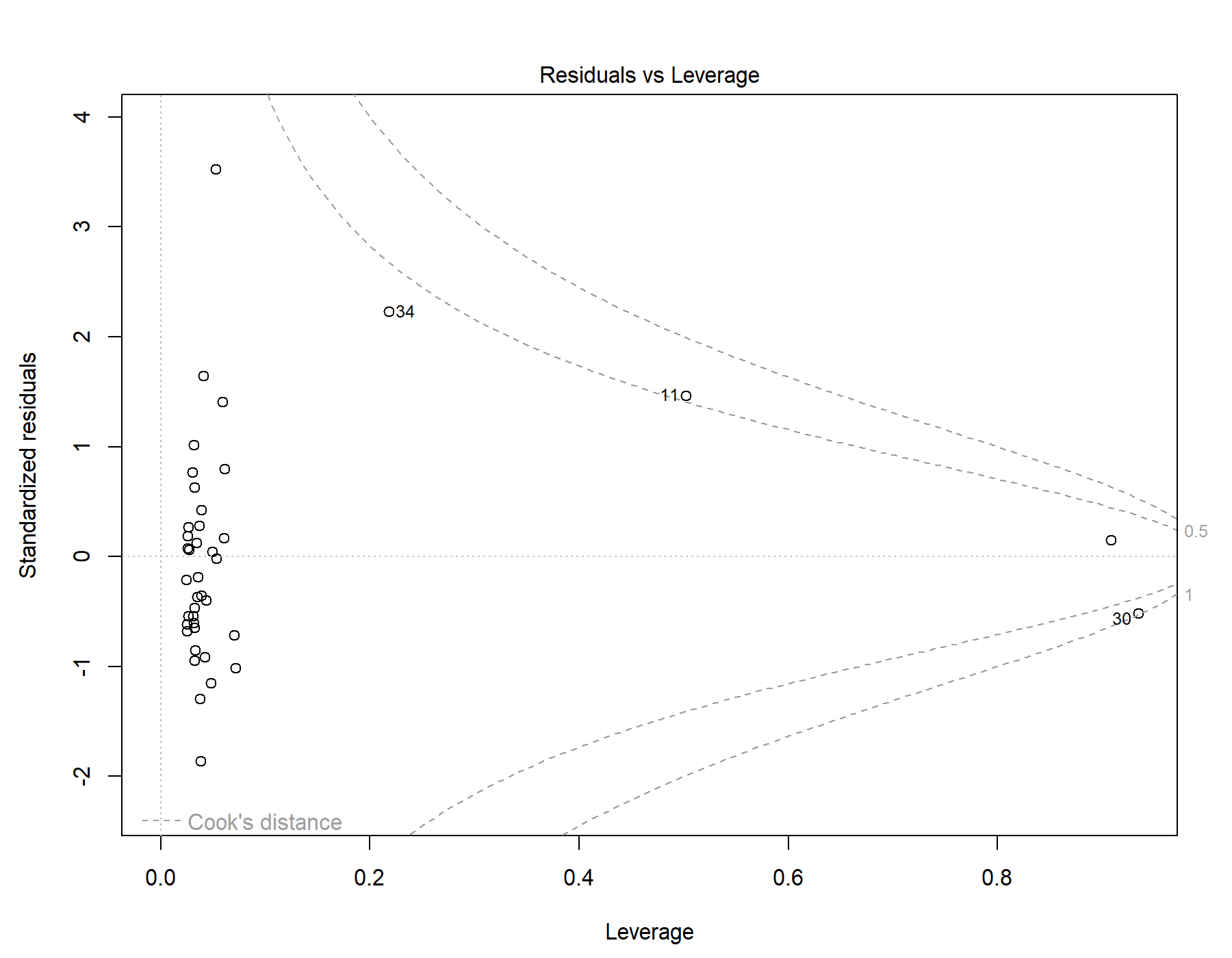

Supplemental Figure 50. Residuals vs leverage diagnostics for T1w+T2w WML segmentation.

### 24. Global | Female only | T1w-alone WML vol | SDNN.LF._log | Exploratory restricted interaction model

Model: T1w-alone WML vol ~ SDNN.LF._log + Age + Age *	SDNN.LF._log

#### 24.1. Standardized coefficients

| **Term** | **Estimate** | **Std. Error** | **t** | **p** | **β** |
| --- | --- | --- | --- | --- | --- |
| (Intercept) | -1.976e-04 | 8.779e-05 | -2.25 | .030 |  |
| SDNN.LF._log | 2.142e-04 | 2.551e-04 | 0.840 | .406 | 0.147 |
| Age | 9.816e-05 | 3.287e-05 | 2.99 | .005 | 0.406 |
| SDNN.LF._log:Age | -2.322e-04 | 6.995e-05 | -3.32 | .002 | -0.586 |

***Supplemental Table 79.*** *Standardized coefficients values for T1w-alone WML segmentation.*

#### 24.2. 95% confidence intervals

| **Term** | **Lower 95%** | **Upper 95%** |
| --- | --- | --- |
| (Intercept) | -3.755e-04 | -1.976e-05 |
| SDNN.LF._log | -3.027e-04 | 7.312e-04 |
| Age | 3.156e-05 | 1.648e-04 |
| SDNN.LF._log:Age | -3.740e-04 | -9.052e-05 |

***Supplemental Table 80.*** *95% confidence intervals values for T1w-alone WML segmentation.*

#### 24.3. Breusch-Pagan test

| **Test** | **Statistic** | **df** | **p** |
| --- | --- | --- | --- |
| Breusch-Pagan | 2.111 | 3 | .550 |

***Supplemental Table 81.*** *Breusch-Pagan test values for T1w-alone WML segmentation.*

#### 24.4. Normal Q-Q plot

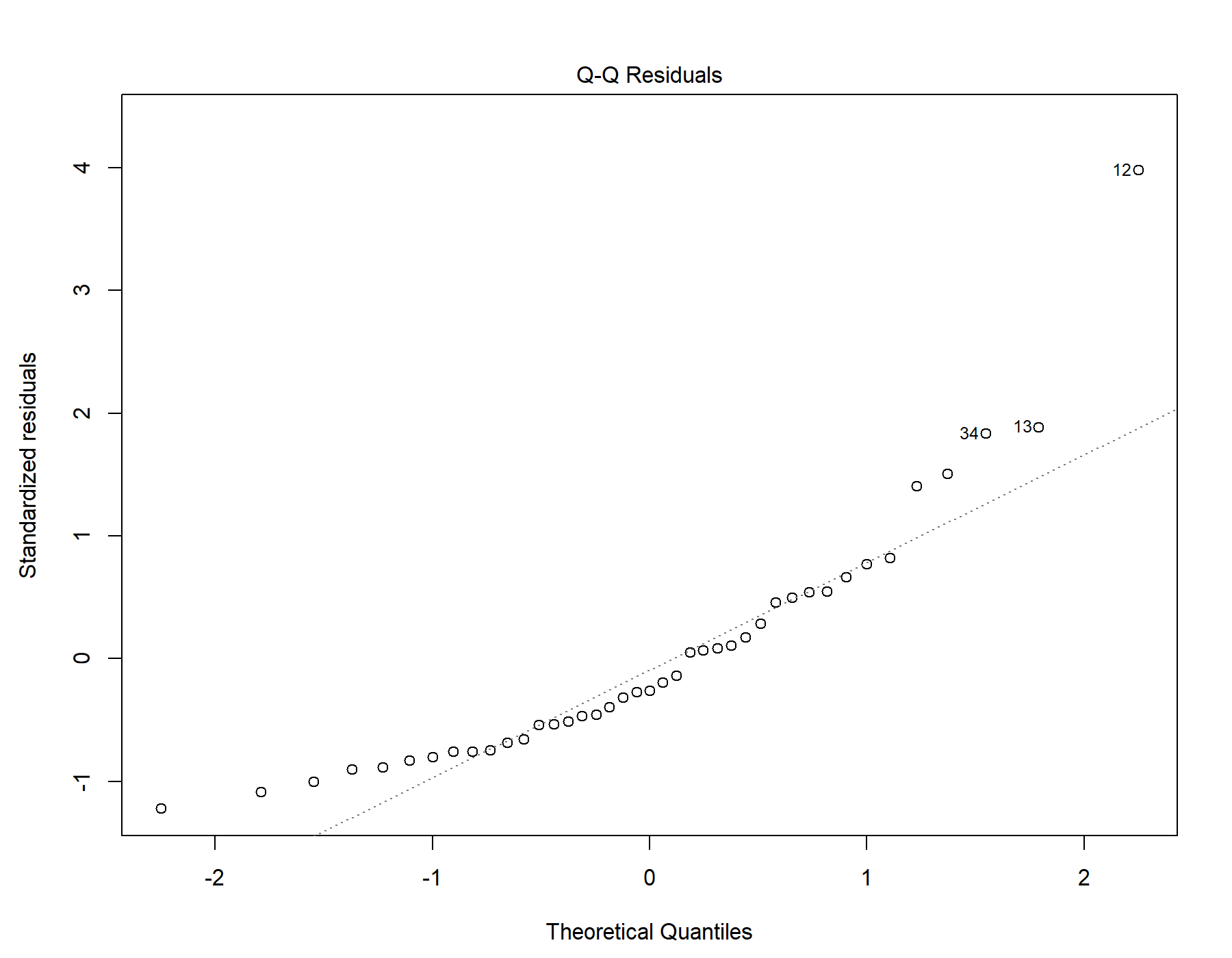

Supplemental Figure 51. Normal distribution quantile-quantile plot for T1w-alone WML segmentation.

#### 24.5. Residuals vs leverage

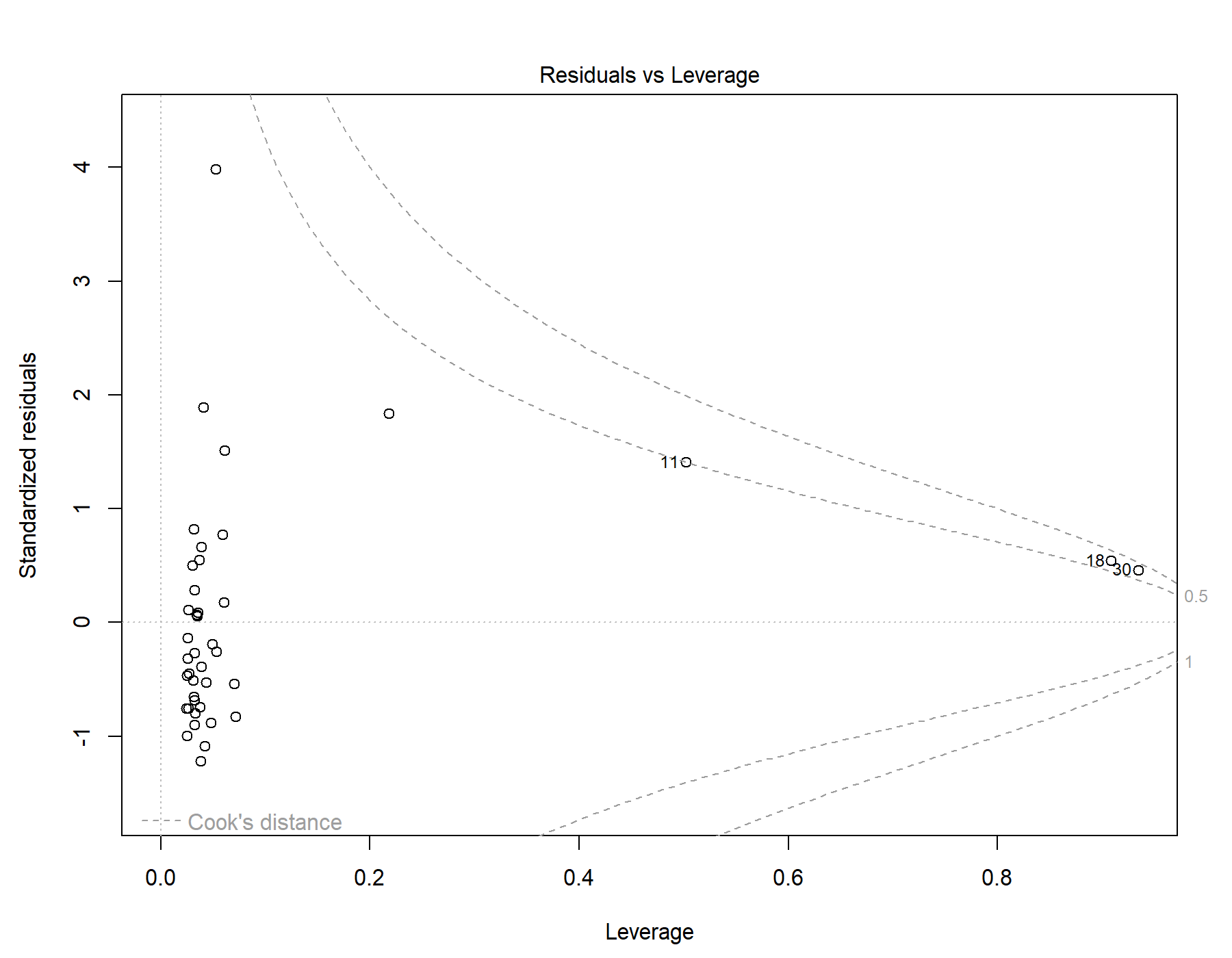

Supplemental Figure 52. Residuals vs leverage diagnostics for T1w-alone WML segmentation.

### 25. Global | Female only | T1w+FLAIR WML vol | SDNN.HF._log | Exploratory restricted interaction model

Model: T1w+FLAIR WML vol ~ SDNN.HF._log + Age +	Age * SDNN.HF._log

#### 25.1. Standardized coefficients

| **Term** | **Estimate** | **Std. Error** | **t** | **p** | **β** |
| --- | --- | --- | --- | --- | --- |
| (Intercept) | 7.066e-05 | 1.127e-04 | 0.627 | .534 |  |
| SDNN.HF._log | 3.942e-04 | 3.347e-04 | 1.18 | .246 | 0.309 |
| Age | -8.238e-05 | 6.751e-05 | -1.22 | .230 | -0.395 |
| SDNN.HF._log:Age | -6.118e-04 | 2.544e-04 | -2.40 | .021 | -1.03 |

***Supplemental Table 82.*** *Standardized coefficients values for T1w+FLAIR WML segmentation.*

#### 25.2. 95% confidence intervals

| **Term** | **Lower 95%** | **Upper 95%** |
| --- | --- | --- |
| (Intercept) | -1.576e-04 | 2.989e-04 |
| SDNN.HF._log | -2.840e-04 | 0.00107 |
| Age | -2.192e-04 | 5.441e-05 |
| SDNN.HF._log:Age | -0.00113 | -9.621e-05 |

***Supplemental Table 83.*** *95% confidence intervals values for T1w+FLAIR WML segmentation.*

#### 25.3. Breusch-Pagan test

| **Test** | **Statistic** | **df** | **p** |
| --- | --- | --- | --- |
| Breusch-Pagan | 5.46 | 3 | .141 |

***Supplemental Table 84.*** *Breusch-Pagan test values for T1w+FLAIR WML segmentation.*

#### 25.4. Normal Q-Q plot

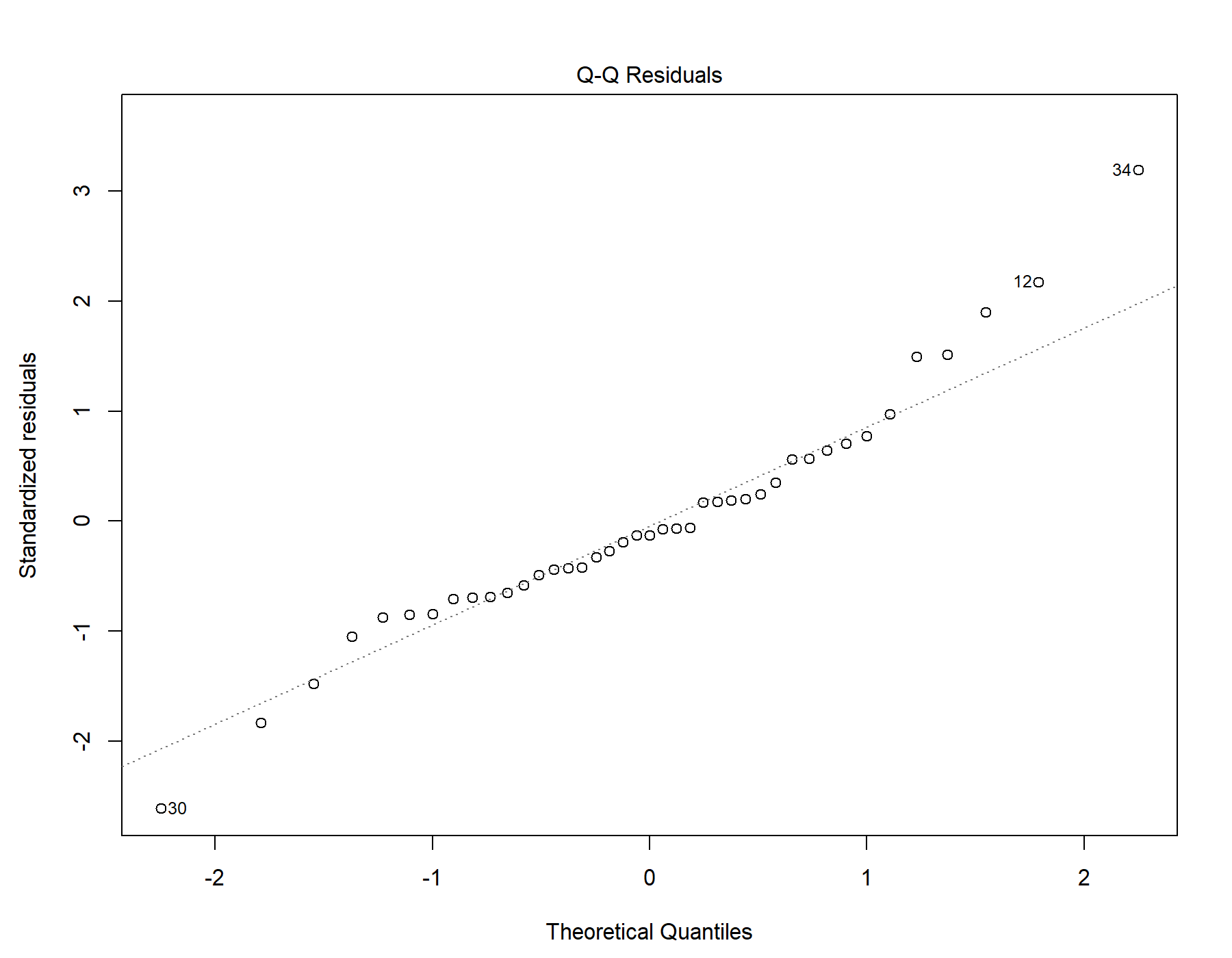

Supplemental Figure 53. Normal distribution quantile-quantile plot for T1w+FLAIR WML segmentation.

#### 25.5. Residuals vs leverage

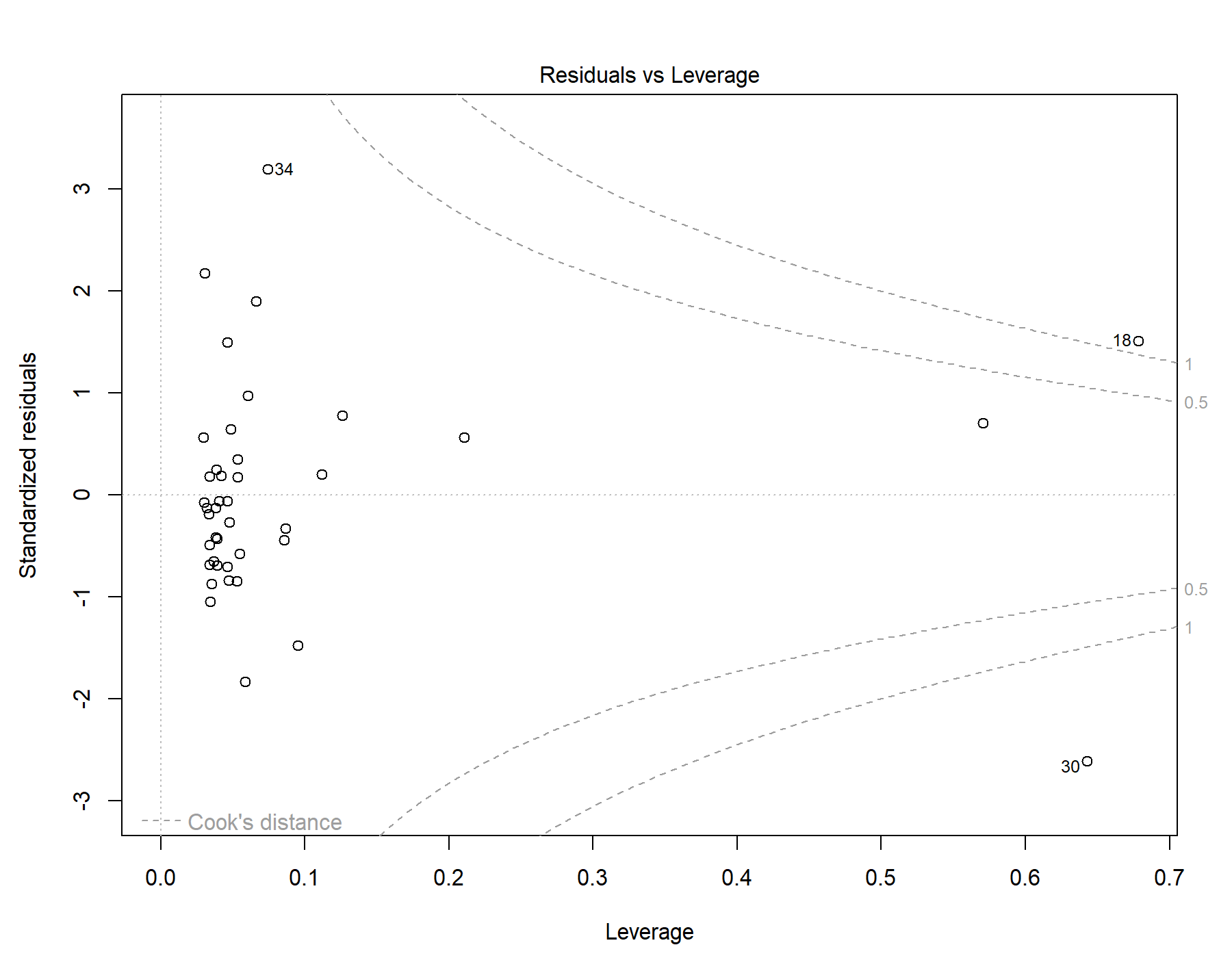

Supplemental Figure 54. Residuals vs leverage diagnostics for T1w+FLAIR WML segmentation.

### 26. Global | Female only | T1w+T2w WML vol | SDNN.HF._log | Exploratory restricted interaction model

Model: T1w+T2w WML vol ~ SDNN.HF._log + Age +	Age * SDNN.HF._log

#### 26.1. Standardized coefficients

| **Term** | **Estimate** | **Std. Error** | **t** | **p** | **β** |
| --- | --- | --- | --- | --- | --- |
| (Intercept) | 8.834e-05 | 1.423e-04 | 0.621 | .538 |  |
| SDNN.HF._log | 8.192e-04 | 4.227e-04 | 1.94 | .060 | 0.523 |
| Age | -1.057e-04 | 8.525e-05 | -1.24 | .223 | -0.412 |
| SDNN.HF._log:Age | -8.011e-04 | 3.213e-04 | -2.49 | .017 | -1.10 |

***Supplemental Table 85.*** *Standardized coefficients values for T1w+T2w WML segmentation.*

#### 26.2. 95% confidence intervals

| **Term** | **Lower 95%** | **Upper 95%** |
| --- | --- | --- |
| (Intercept) | -1.999e-04 | 3.766e-04 |
| SDNN.HF._log | -3.718e-05 | 0.00168 |
| Age | -2.784e-04 | 6.704e-05 |
| SDNN.HF._log:Age | -0.00145 | -1.500e-04 |

***Supplemental Table 86.*** *95% confidence intervals values for T1w+T2w WML segmentation.*

#### 26.3. Breusch-Pagan test

| **Test** | **Statistic** | **df** | **p** |
| --- | --- | --- | --- |
| Breusch-Pagan | 2.981 | 3 | .395 |

***Supplemental Table 87.*** *Breusch-Pagan test values for T1w+T2w WML segmentation.*

#### 26.4. Normal Q-Q plot

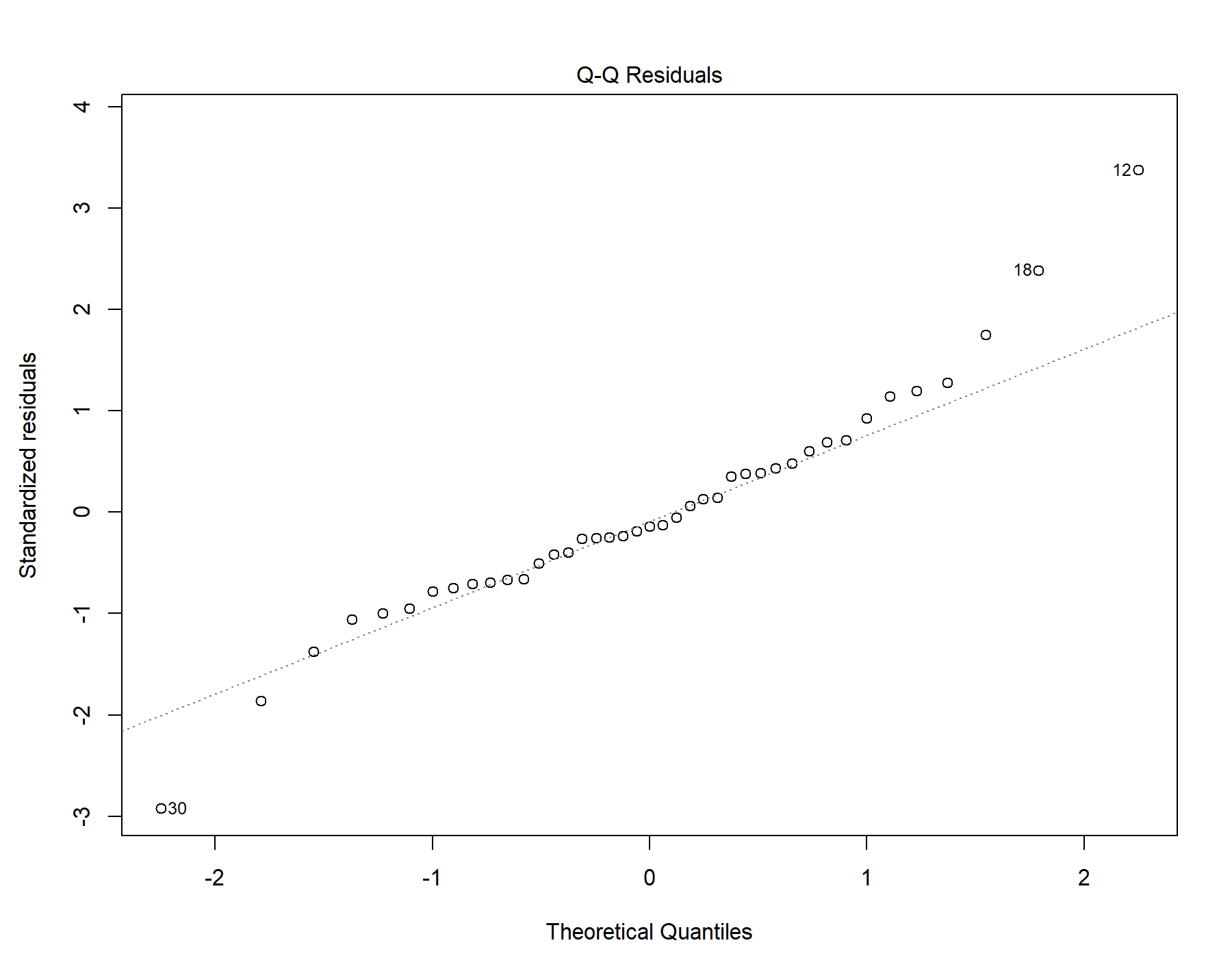

Supplemental Figure 55. Normal distribution quantile-quantile plot for T1w+T2w WML segmentation.

#### 26.5. Residuals vs leverage

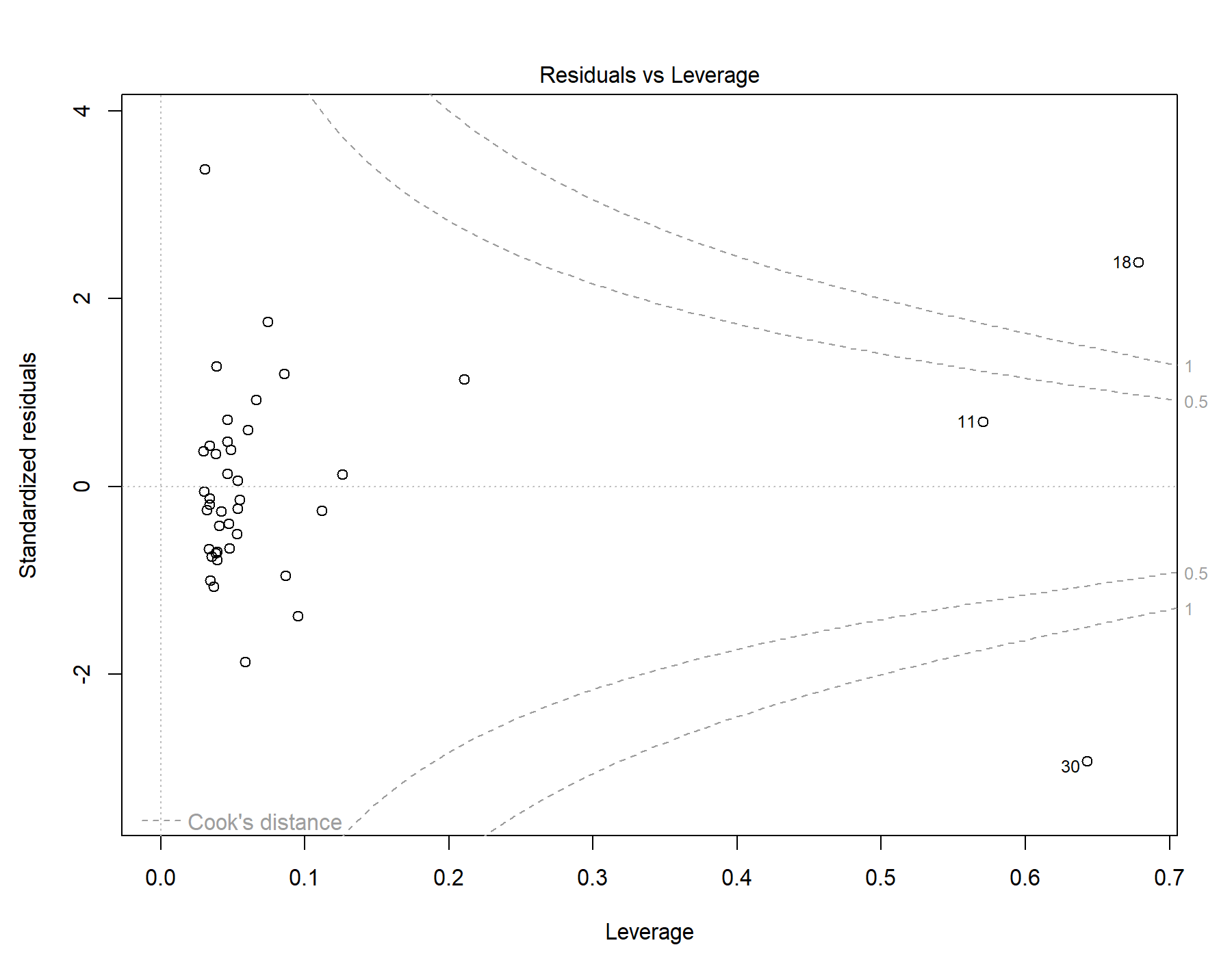

Supplemental Figure 56. Residuals vs leverage diagnostics for T1w+T2w WML segmentation.

### 27. Global | Female only | T1w-alone WML vol | SDNN.HF._log | Exploratory restricted interaction model

Model: T1w-alone WML vol ~ SDNN.HF._log + Age + Age *	SDNN.HF._log

#### 27.1. Standardized coefficients

| **Term** | **Estimate** | **Std. Error** | **t** | **p** | **β** |
| --- | --- | --- | --- | --- | --- |
| (Intercept) | 1.130e-04 | 1.333e-04 | 0.848 | .402 |  |
| SDNN.HF._log | 8.243e-04 | 3.959e-04 | 2.08 | .044 | 0.558 |
| Age | -1.197e-04 | 7.985e-05 | -1.50 | .142 | -0.495 |
| SDNN.HF._log:Age | -8.174e-04 | 3.010e-04 | -2.72 | .010 | -1.19 |

***Supplemental Table 88.*** *Standardized coefficients values for T1w-alone WML segmentation.*

#### 27.2. 95% confidence intervals

| **Term** | **Lower 95%** | **Upper 95%** |
| --- | --- | --- |
| (Intercept) | -1.570e-04 | 3.830e-04 |
| SDNN.HF._log | 2.212e-05 | 0.00163 |
| Age | -2.815e-04 | 4.209e-05 |
| SDNN.HF._log:Age | -0.00143 | -2.076e-04 |

***Supplemental Table 89.*** *95% confidence intervals values for T1w-alone WML segmentation.*

#### 27.3. Breusch-Pagan test

| **Test** | **Statistic** | **df** | **p** |
| --- | --- | --- | --- |
| Breusch-Pagan | 0.908 | 3 | .824 |

***Supplemental Table 90.*** *Breusch-Pagan test values for T1w-alone WML segmentation.*

#### 27.4. Normal Q-Q plot

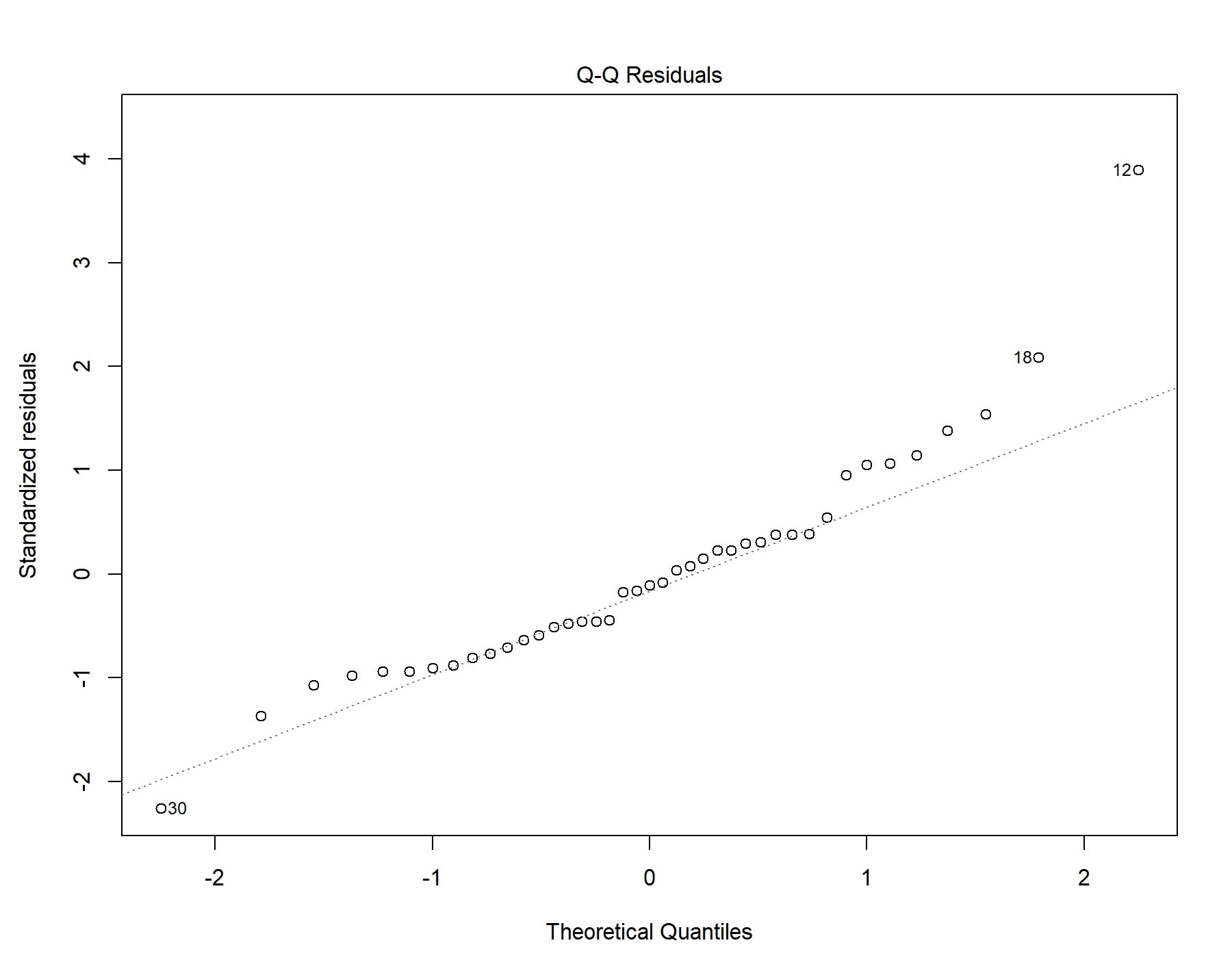

Supplemental Figure 57. Normal distribution quantile-quantile plot for T1w-alone WML segmentation.

#### 27.5. Residuals vs leverage

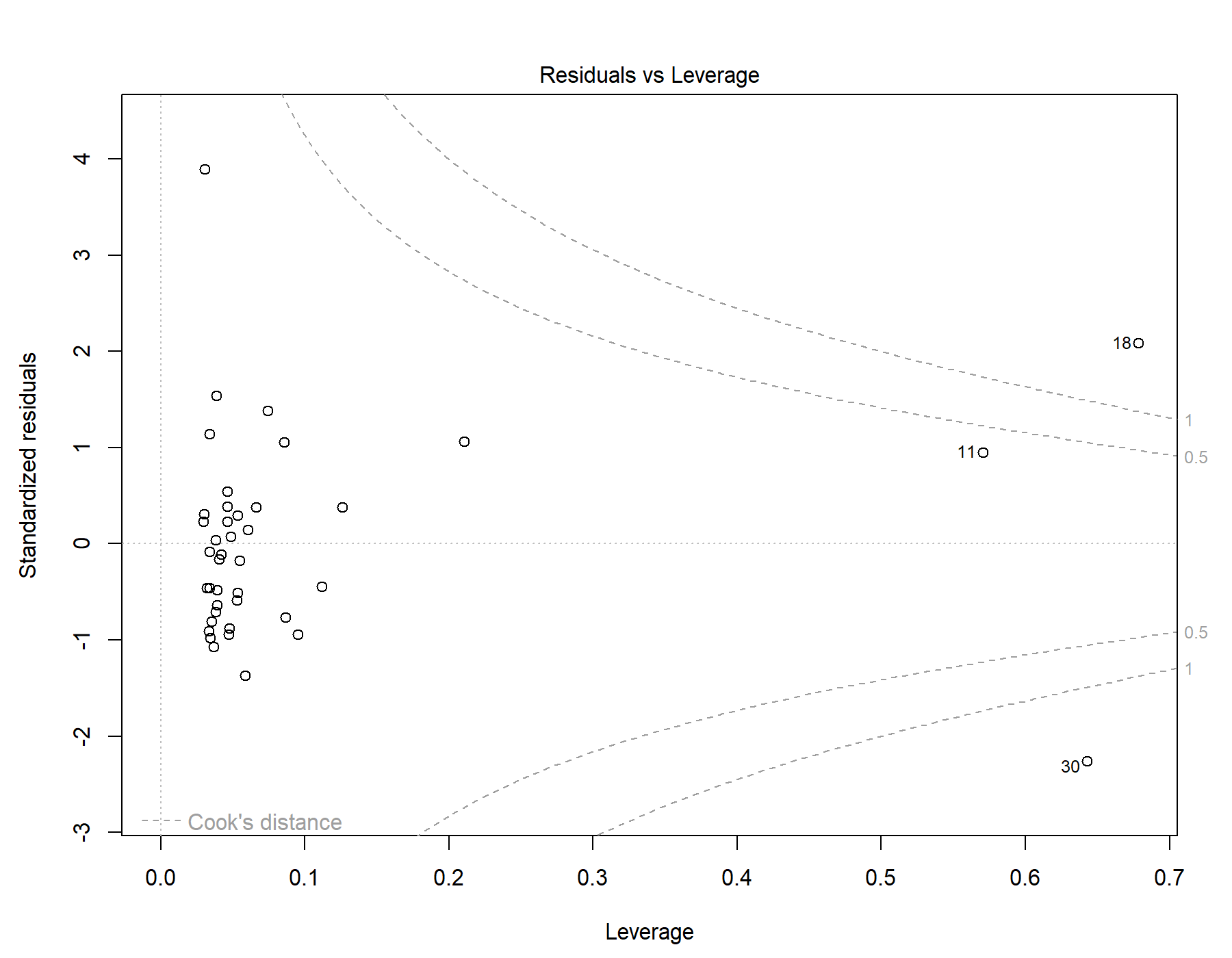

Supplemental Figure 58. Residuals vs leverage diagnostics for T1w-alone WML segmentation.

### 28. Global | Male only | T1w+FLAIR WML vol | RMSSD | Exploratory restricted interaction model

Model: T1w+FLAIR WML vol ~ RMSSD + Age + Age *	RMSSD

#### 28.1. Standardized coefficients

| **Term** | **Estimate** | **Std. Error** | **t** | **p** | **β** |
| --- | --- | --- | --- | --- | --- |
| (Intercept) | -7.110e-04 | 1.046e-04 | -6.80 | <.001 |  |
| RMSSD | 1.980e-06 | 2.601e-06 | 0.761 | .448 | 0.0938 |
| Age | 1.619e-04 | 1.859e-05 | 8.71 | <.001 | 0.661 |
| RMSSD:Age | -7.734e-07 | 3.645e-07 | -2.12 | .037 | -0.260 |

***Supplemental Table 91.*** *Standardized coefficients values for T1w+FLAIR WML segmentation.*

#### 28.2. 95% confidence intervals

| **Term** | **Lower 95%** | **Upper 95%** |
| --- | --- | --- |
| (Intercept) | -9.188e-04 | -5.032e-04 |
| RMSSD | -3.188e-06 | 7.149e-06 |
| Age | 1.250e-04 | 1.989e-04 |
| RMSSD:Age | -1.498e-06 | -4.909e-08 |

***Supplemental Table 92.*** *95% confidence intervals values for T1w+FLAIR WML segmentation.*

#### 28.3. Breusch-Pagan test

| **Test** | **Statistic** | **df** | **p** |
| --- | --- | --- | --- |
| Breusch-Pagan | 12.99 | 3 | .005 |

***Supplemental Table 93.*** *Breusch-Pagan test values for T1w+FLAIR WML segmentation.*

#### 28.4. Normal Q-Q plot

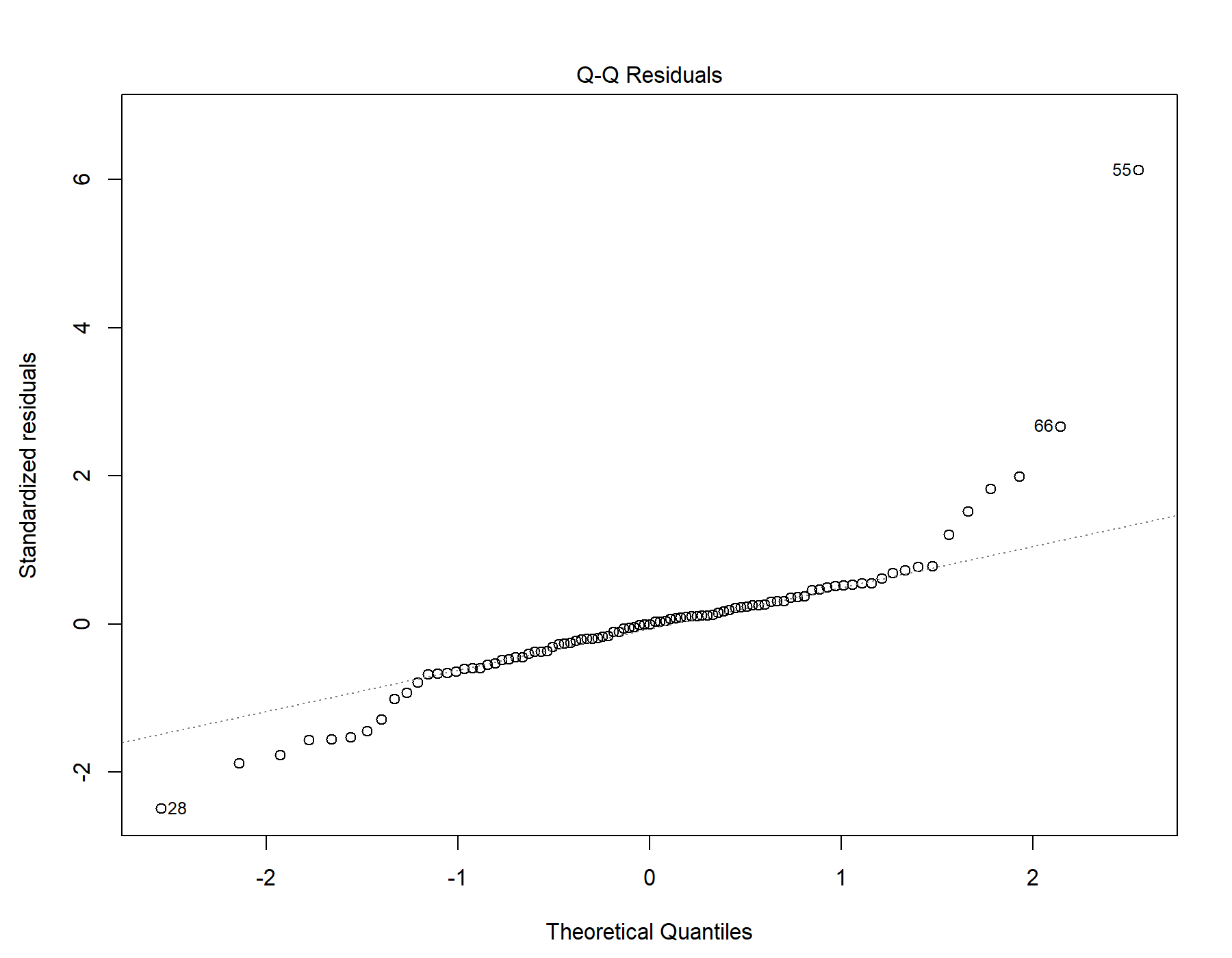

Supplemental Figure 59. Normal distribution quantile-quantile plot for T1w+FLAIR WML segmentation.

#### 28.5. Residuals vs leverage

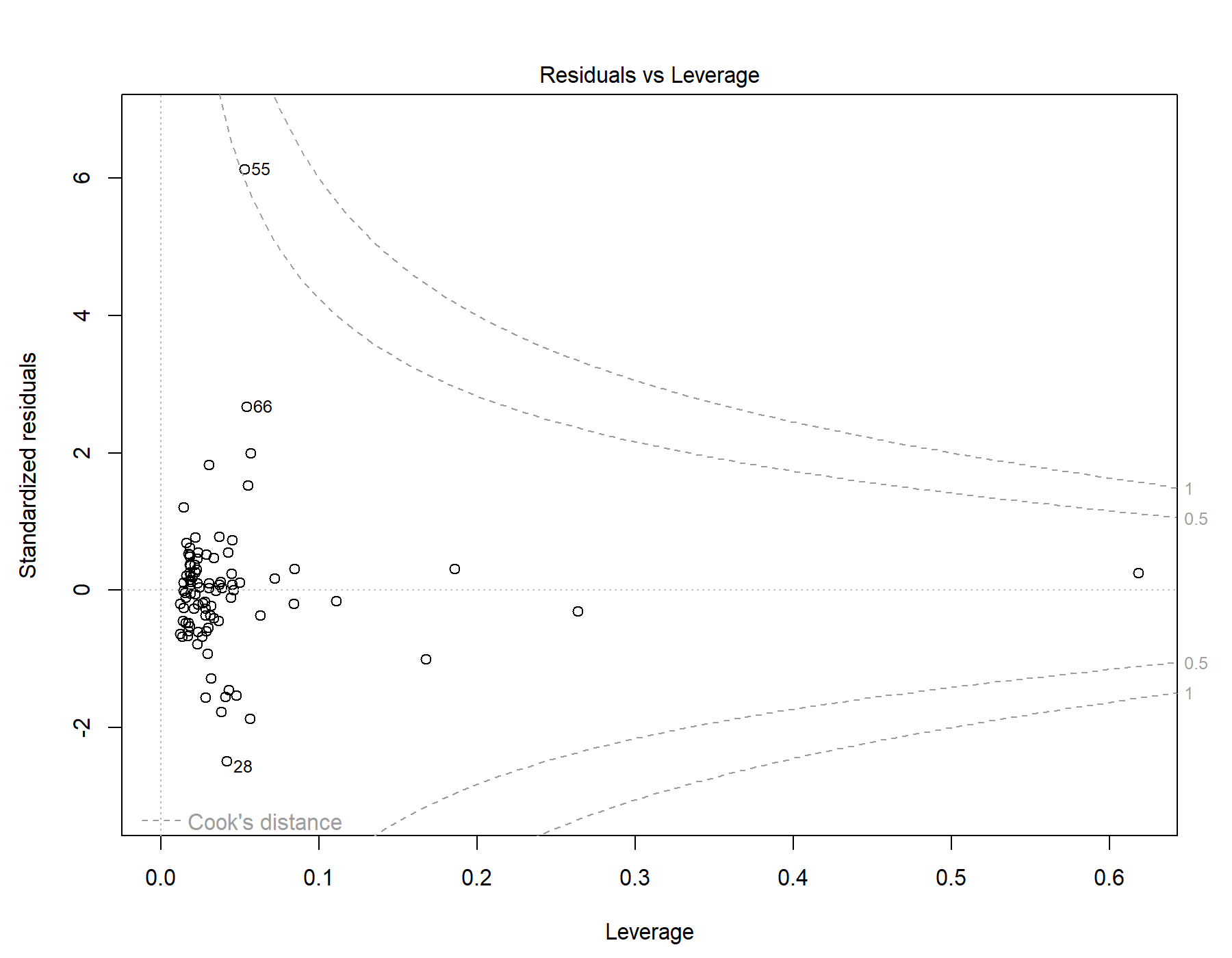

Supplemental Figure 60. Residuals vs leverage diagnostics for T1w+FLAIR WML segmentation.

### 29. Global | Male only | T1w+T2w WML vol | RMSSD | Exploratory restricted interaction model

Model: T1w+T2w WML vol ~ RMSSD + Age + Age *	RMSSD

#### 29.1. Standardized coefficients

| **Term** | **Estimate** | **Std. Error** | **t** | **p** | **β** |
| --- | --- | --- | --- | --- | --- |
| (Intercept) | -8.689e-04 | 1.122e-04 | -7.74 | <.001 |  |
| RMSSD | 3.207e-06 | 2.790e-06 | 1.15 | .253 | 0.134 |
| Age | 1.992e-04 | 1.994e-05 | 9.99 | <.001 | 0.718 |
| RMSSD:Age | -8.200e-07 | 3.910e-07 | -2.10 | .039 | -0.243 |

***Supplemental Table 94.*** *Standardized coefficients values for T1w+T2w WML segmentation.*

#### 29.2. 95% confidence intervals

| **Term** | **Lower 95%** | **Upper 95%** |
| --- | --- | --- |
| (Intercept) | -0.00109 | -6.460e-04 |
| RMSSD | -2.337e-06 | 8.751e-06 |
| Age | 1.596e-04 | 2.388e-04 |
| RMSSD:Age | -1.597e-06 | -4.304e-08 |

***Supplemental Table 95.*** *95% confidence intervals values for T1w+T2w WML segmentation.*

#### 29.3. Breusch-Pagan test

| **Test** | **Statistic** | **df** | **p** |
| --- | --- | --- | --- |
| Breusch-Pagan | 11.431 | 3 | .010 |

***Supplemental Table 96.*** *Breusch-Pagan test values for T1w+T2w WML segmentation.*

#### 29.4. Normal Q-Q plot

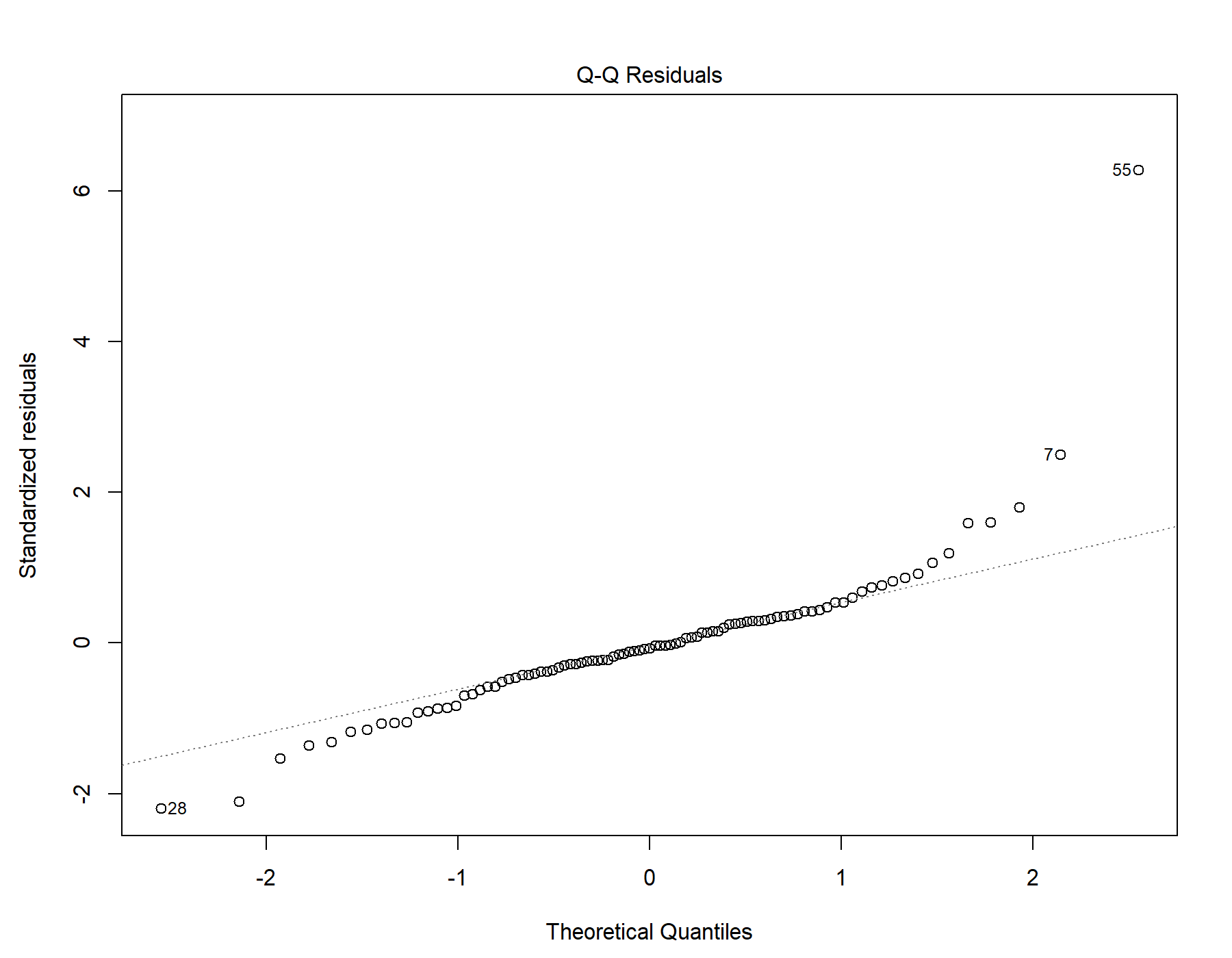

Supplemental Figure 61. Normal distribution quantile-quantile plot for T1w+T2w WML segmentation.

#### 29.5. Residuals vs leverage

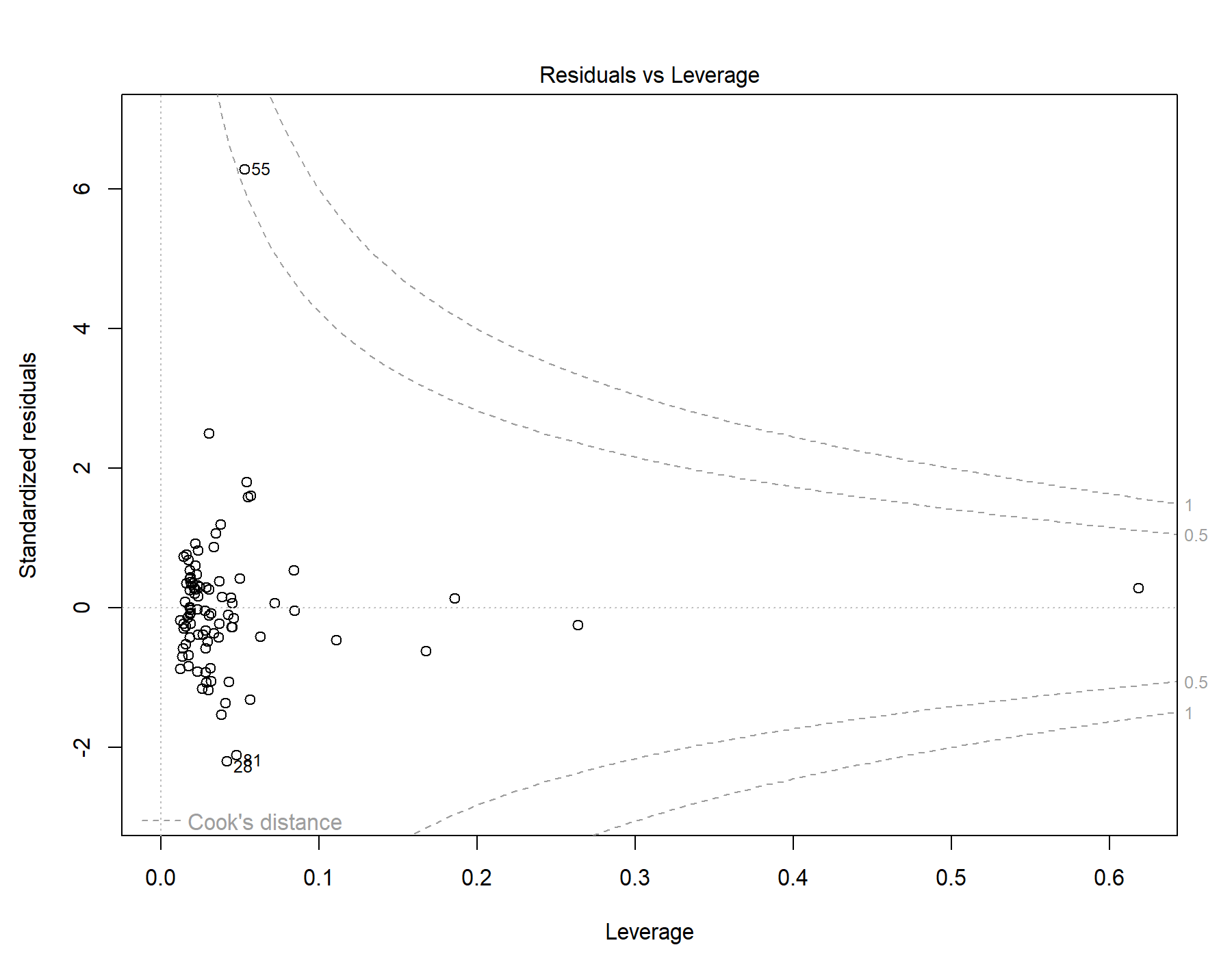

Supplemental Figure 62. Residuals vs leverage diagnostics for T1w+T2w WML segmentation.

### 30. Global | Male only | T1w-alone WML vol | RMSSD | Exploratory restricted interaction model

Model: T1w-alone WML vol ~ RMSSD + Age + Age *	RMSSD

#### 30.1. Standardized coefficients

| **Term** | **Estimate** | **Std. Error** | **t** | **p** | **β** |
| --- | --- | --- | --- | --- | --- |
| (Intercept) | -6.940e-04 | 9.305e-05 | -7.46 | <.001 |  |
| RMSSD | 2.325e-06 | 2.314e-06 | 1.00 | .318 | 0.120 |
| Age | 1.595e-04 | 1.654e-05 | 9.65 | <.001 | 0.708 |
| RMSSD:Age | -6.168e-07 | 3.243e-07 | -1.90 | .060 | -0.225 |

***Supplemental Table 97.*** *Standardized coefficients values for T1w-alone WML segmentation.*

#### 30.2. 95% confidence intervals

| **Term** | **Lower 95%** | **Upper 95%** |
| --- | --- | --- |
| (Intercept) | -8.789e-04 | -5.091e-04 |
| RMSSD | -2.273e-06 | 6.923e-06 |
| Age | 1.266e-04 | 1.924e-04 |
| RMSSD:Age | -1.261e-06 | 2.760e-08 |

***Supplemental Table 98.*** *95% confidence intervals values for T1w-alone WML segmentation.*

#### 30.3. Breusch-Pagan test

| **Test** | **Statistic** | **df** | **p** |
| --- | --- | --- | --- |
| Breusch-Pagan | 13.402 | 3 | .004 |

***Supplemental Table 99.*** *Breusch-Pagan test values for T1w-alone WML segmentation.*

#### 30.4. Normal Q-Q plot

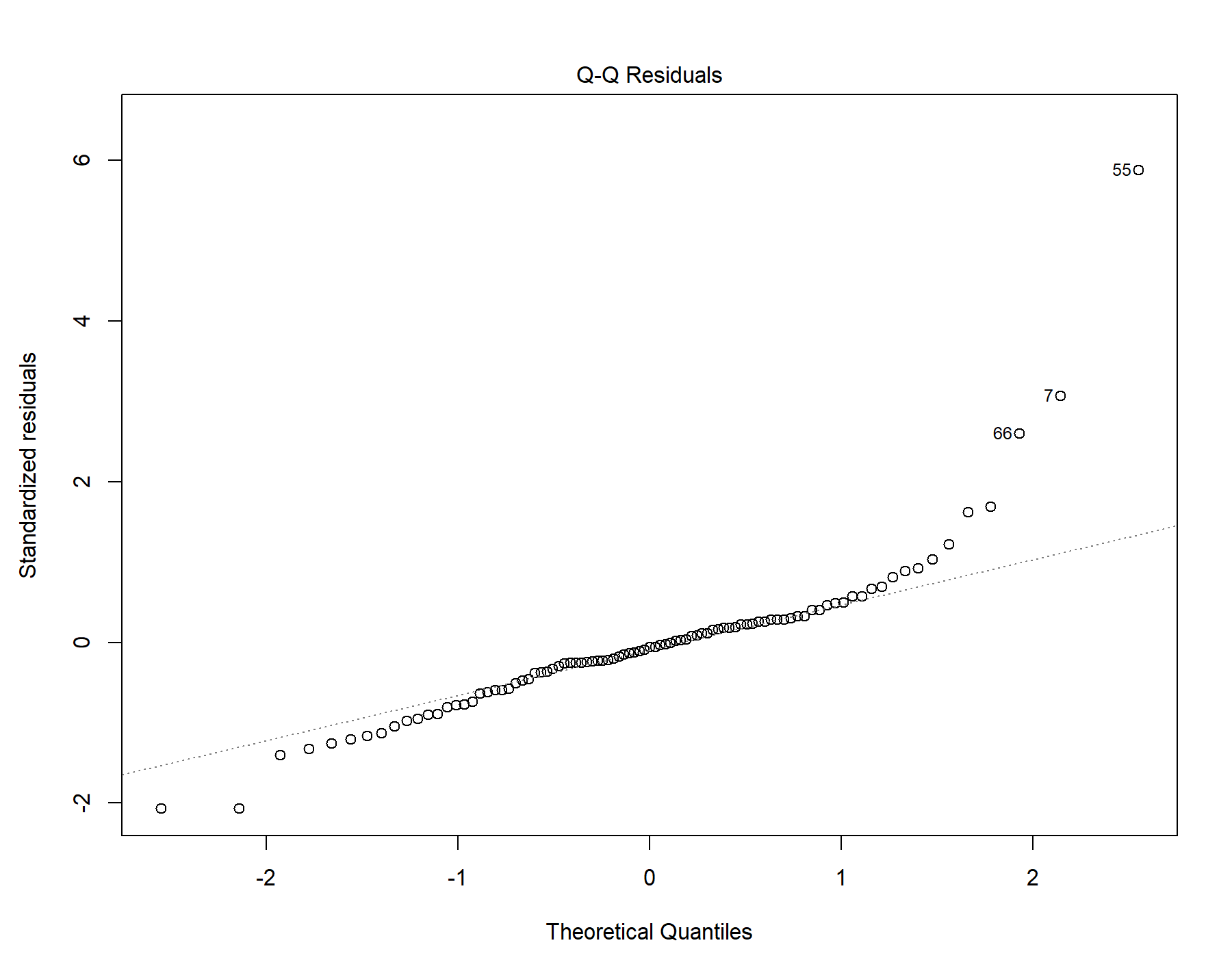

Supplemental Figure 63. Normal distribution quantile-quantile plot for T1w-alone WML segmentation.

#### 30.5. Residuals vs leverage

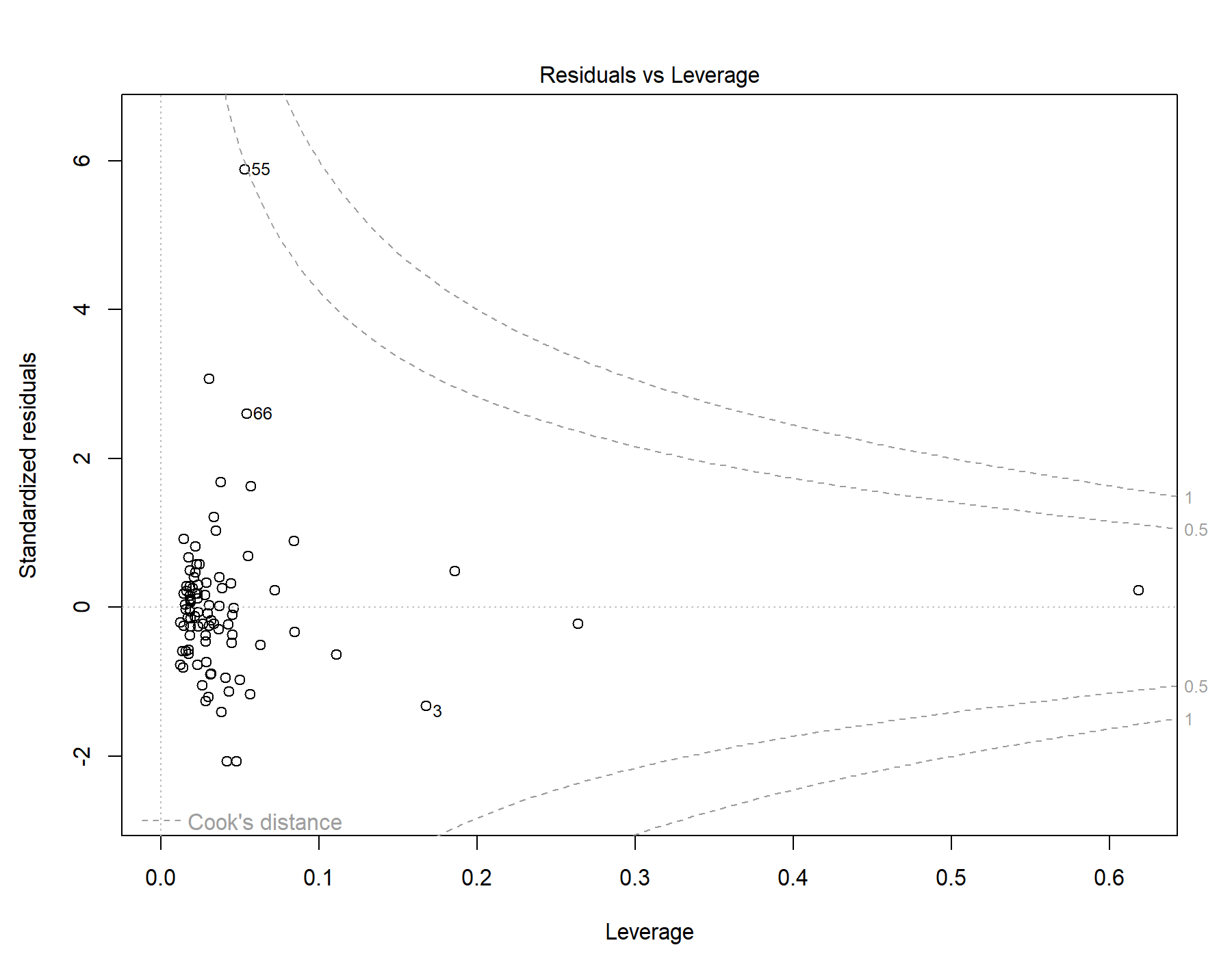

Supplemental Figure 64. Residuals vs leverage diagnostics for T1w-alone WML segmentation.

### 31. Global | Male only | T1w+FLAIR WML vol | SDNN.LF._log | Exploratory restricted interaction model

Model: T1w+FLAIR WML vol ~ SDNN.LF._log + Age +	Age * SDNN.LF._log

#### 31.1. Standardized coefficients

| **Term** | **Estimate** | **Std. Error** | **t** | **p** | **β** |
| --- | --- | --- | --- | --- | --- |
| (Intercept) | -7.265e-04 | 1.058e-04 | -6.87 | <.001 |  |
| SDNN.LF._log | 3.650e-04 | 2.959e-04 | 1.23 | .221 | 0.165 |
| Age | 1.669e-04 | 1.876e-05 | 8.90 | <.001 | 0.681 |
| SDNN.LF._log:Age | -7.448e-05 | 3.947e-05 | -1.89 | .062 | -0.252 |

***Supplemental Table 100.*** *Standardized coefficients values for T1w+FLAIR WML segmentation.*

#### 31.2. 95% confidence intervals

| **Term** | **Lower 95%** | **Upper 95%** |
| --- | --- | --- |
| (Intercept) | -9.367e-04 | -5.163e-04 |
| SDNN.LF._log | -2.230e-04 | 9.529e-04 |
| Age | 1.296e-04 | 2.042e-04 |
| SDNN.LF._log:Age | -1.529e-04 | 3.955e-06 |

***Supplemental Table 101.*** *95% confidence intervals values for T1w+FLAIR WML segmentation.*

#### 31.3. Breusch-Pagan test

| **Test** | **Statistic** | **df** | **p** |
| --- | --- | --- | --- |
| Breusch-Pagan | 11.764 | 3 | .008 |

***Supplemental Table 102.*** *Breusch-Pagan test values for T1w+FLAIR WML segmentation.*

#### 31.4. Normal Q-Q plot

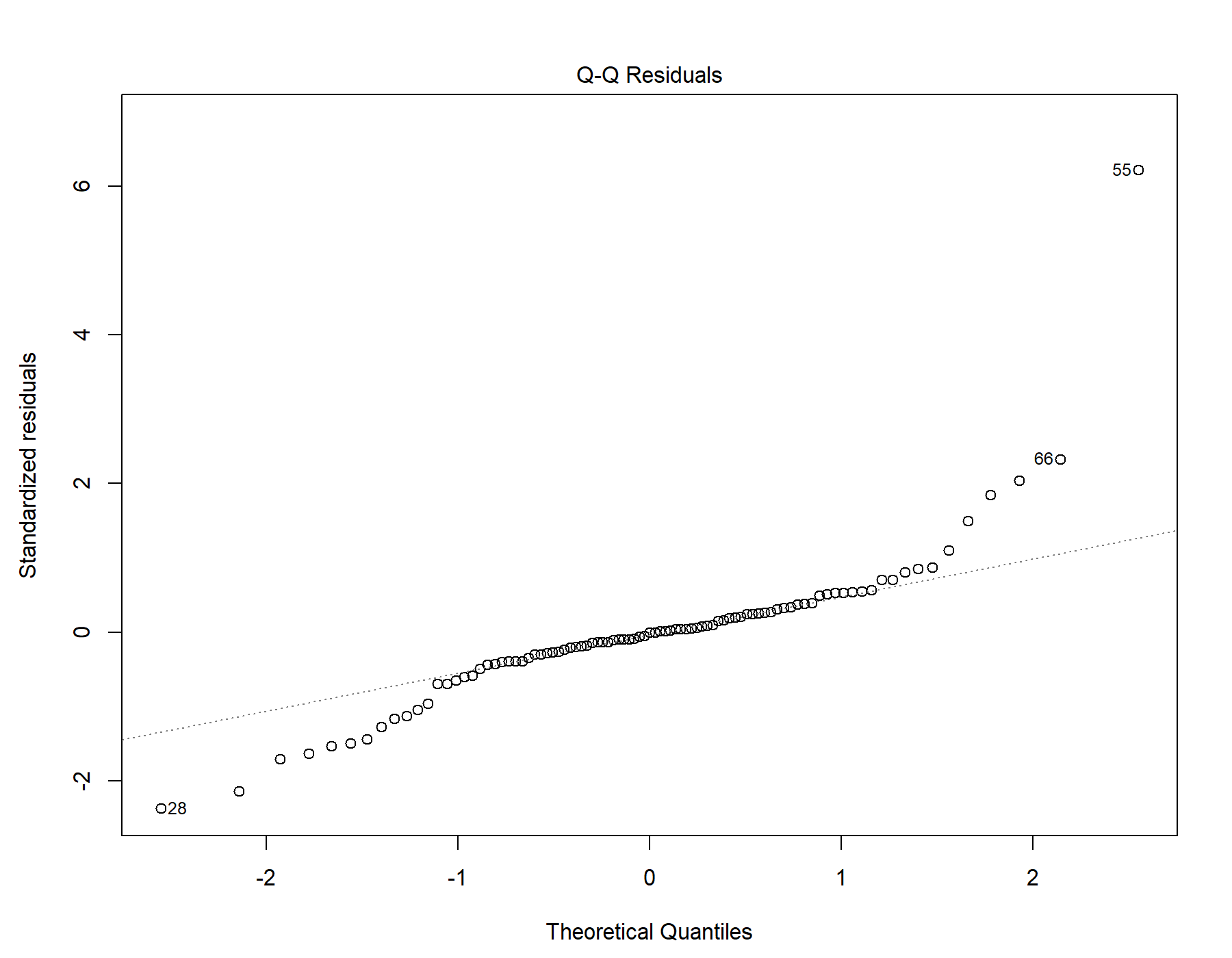

Supplemental Figure 65. Normal distribution quantile-quantile plot for T1w+FLAIR WML segmentation.

#### 31.5. Residuals vs leverage

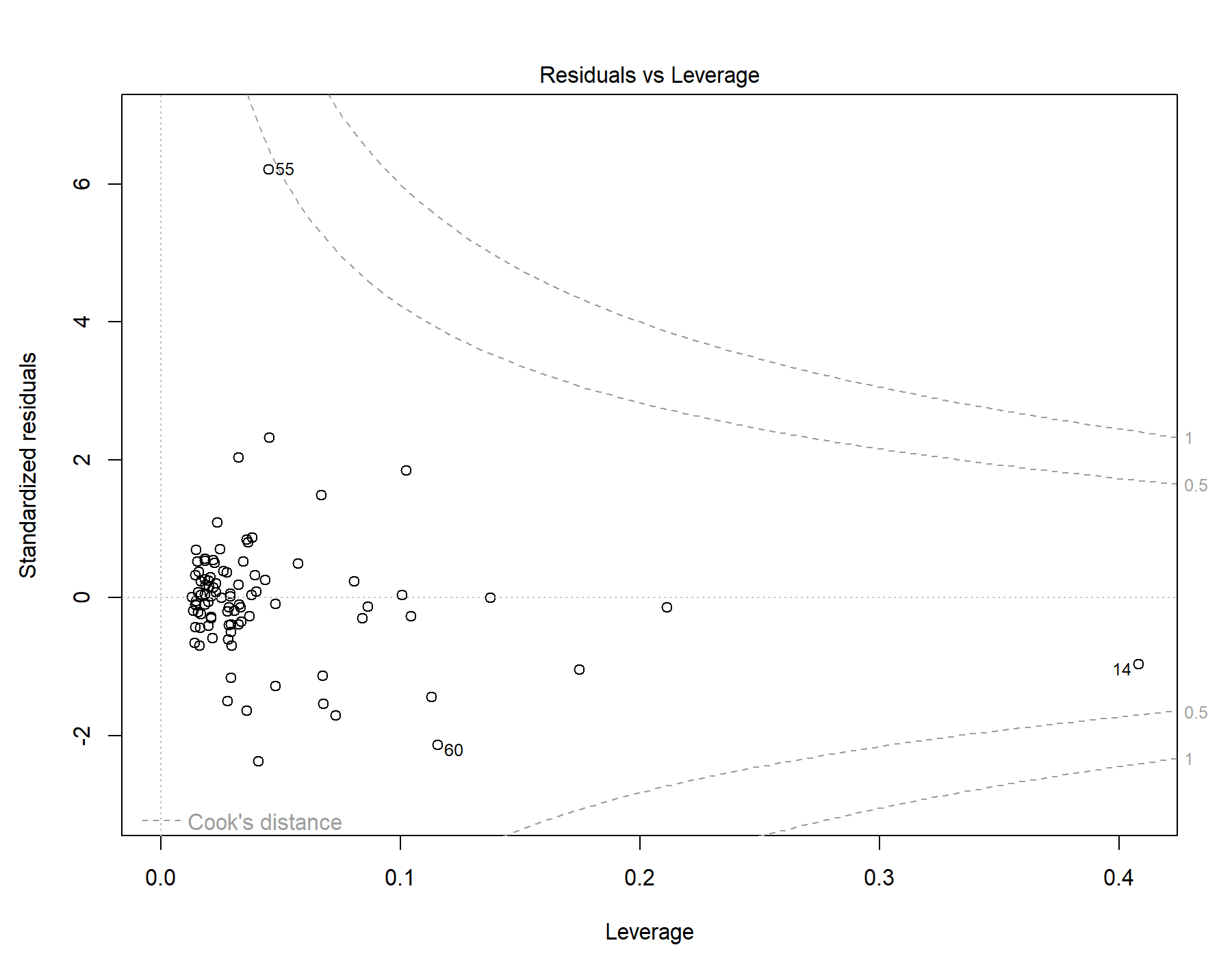

Supplemental Figure 66. Residuals vs leverage diagnostics for T1w+FLAIR WML segmentation.

### 32. Global | Male only | T1w+T2w WML vol | SDNN.LF._log | Exploratory restricted interaction model

Model: T1w+T2w WML vol ~ SDNN.LF._log + Age +	Age * SDNN.LF._log

#### 32.1. Standardized coefficients

| **Term** | **Estimate** | **Std. Error** | **t** | **p** | **β** |
| --- | --- | --- | --- | --- | --- |
| (Intercept) | -8.720e-04 | 1.130e-04 | -7.72 | <.001 |  |
| SDNN.LF._log | 3.661e-04 | 3.160e-04 | 1.16 | .250 | 0.146 |
| Age | 2.019e-04 | 2.003e-05 | 10.1 | <.001 | 0.728 |
| SDNN.LF._log:Age | -7.171e-05 | 4.216e-05 | -1.70 | .092 | -0.214 |

***Supplemental Table 103.*** *Standardized coefficients values for T1w+T2w WML segmentation.*

#### 32.2. 95% confidence intervals

| **Term** | **Lower 95%** | **Upper 95%** |
| --- | --- | --- |
| (Intercept) | -0.00110 | -6.475e-04 |
| SDNN.LF._log | -2.618e-04 | 9.941e-04 |
| Age | 1.621e-04 | 2.417e-04 |
| SDNN.LF._log:Age | -1.555e-04 | 1.206e-05 |

***Supplemental Table 104.*** *95% confidence intervals values for T1w+T2w WML segmentation.*

#### 32.3. Breusch-Pagan test

| **Test** | **Statistic** | **df** | **p** |
| --- | --- | --- | --- |
| Breusch-Pagan | 9.345 | 3 | .025 |

***Supplemental Table 105.*** *Breusch-Pagan test values for T1w+T2w WML segmentation.*

#### 32.4. Normal Q-Q plot

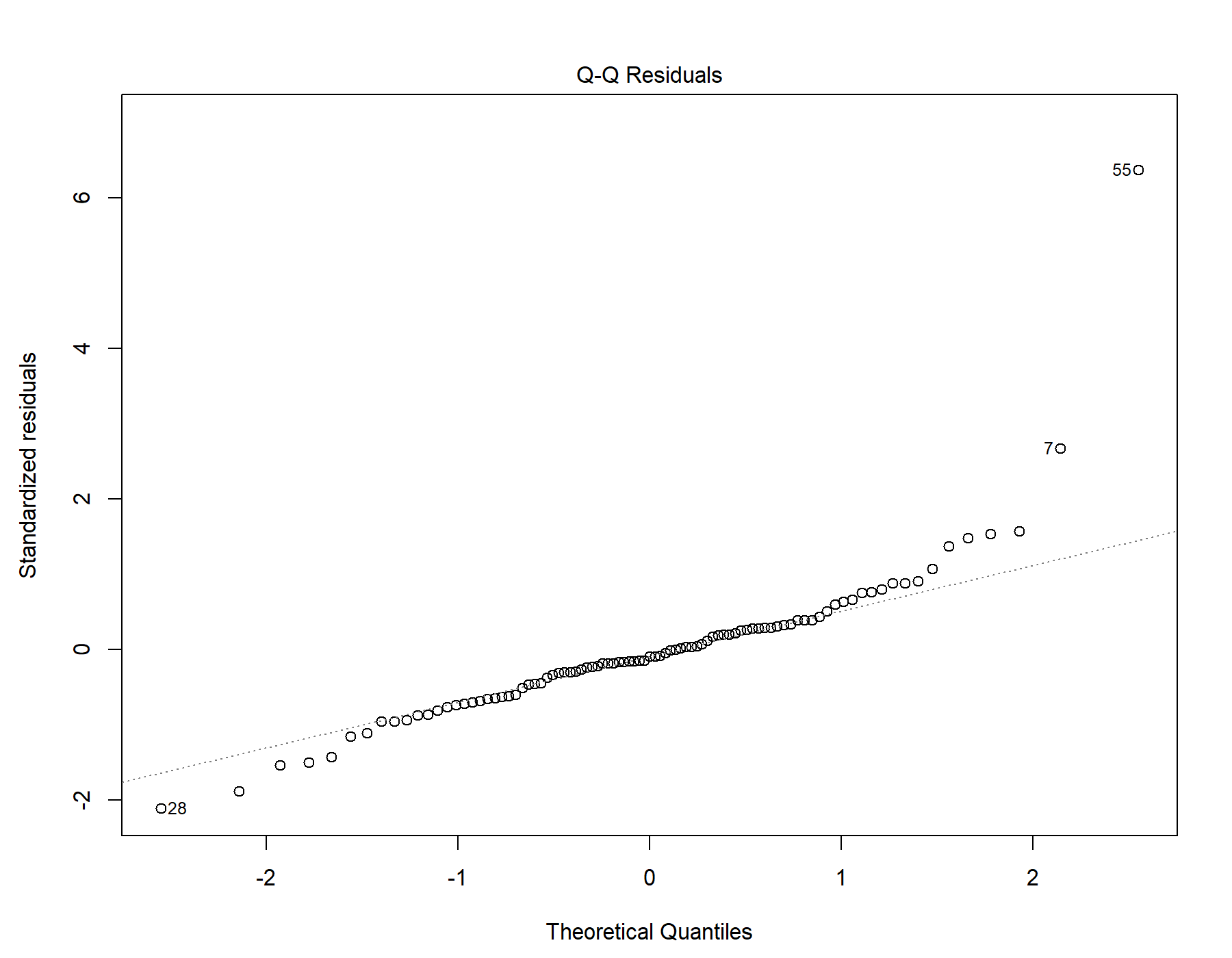

Supplemental Figure 67. Normal distribution quantile-quantile plot for T1w+T2w WML segmentation.

#### 32.5. Residuals vs leverage

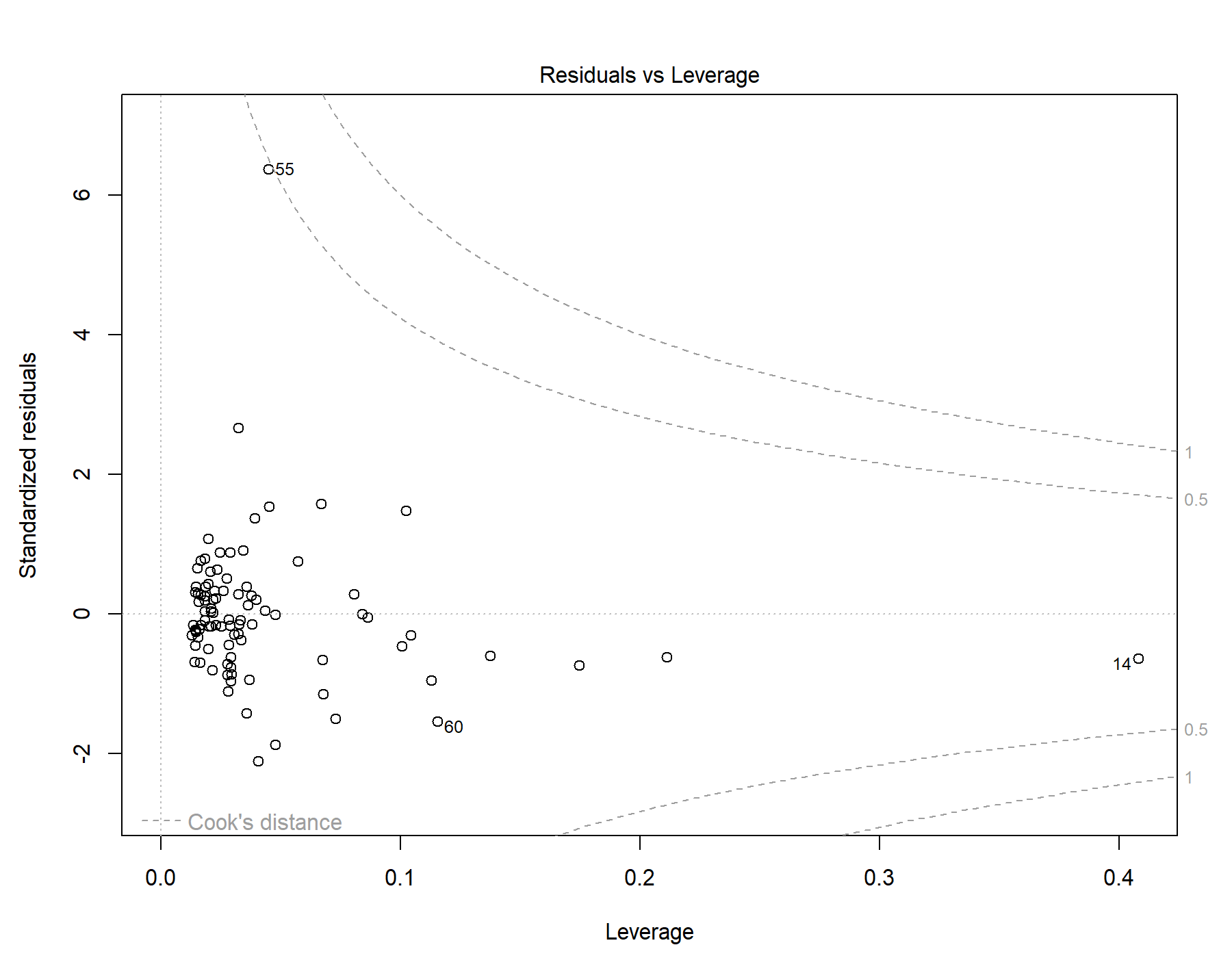

Supplemental Figure 68. Residuals vs leverage diagnostics for T1w+T2w WML segmentation.

### 33. Global | Male only | T1w-alone WML vol | SDNN.LF._log | Exploratory restricted interaction model

Model: T1w-alone WML vol ~ SDNN.LF._log + Age + Age *	SDNN.LF._log

#### 33.1. Standardized coefficients

| **Term** | **Estimate** | **Std. Error** | **t** | **p** | **β** |
| --- | --- | --- | --- | --- | --- |
| (Intercept) | -6.964e-04 | 9.331e-05 | -7.46 | <.001 |  |
| SDNN.LF._log | 2.792e-04 | 2.610e-04 | 1.07 | .288 | 0.137 |
| Age | 1.612e-04 | 1.655e-05 | 9.74 | <.001 | 0.715 |
| SDNN.LF._log:Age | -5.797e-05 | 3.482e-05 | -1.66 | .099 | -0.213 |

***Supplemental Table 106.*** *Standardized coefficients values for T1w-alone WML segmentation.*

#### 33.2. 95% confidence intervals

| **Term** | **Lower 95%** | **Upper 95%** |
| --- | --- | --- |
| (Intercept) | -8.818e-04 | -5.110e-04 |
| SDNN.LF._log | -2.395e-04 | 7.978e-04 |
| Age | 1.283e-04 | 1.941e-04 |
| SDNN.LF._log:Age | -1.272e-04 | 1.122e-05 |

***Supplemental Table 107.*** *95% confidence intervals values for T1w-alone WML segmentation.*

#### 33.3. Breusch-Pagan test

| **Test** | **Statistic** | **df** | **p** |
| --- | --- | --- | --- |
| Breusch-Pagan | 11.183 | 3 | .011 |

***Supplemental Table 108.*** *Breusch-Pagan test values for T1w-alone WML segmentation.*

#### 33.4. Normal Q-Q plot

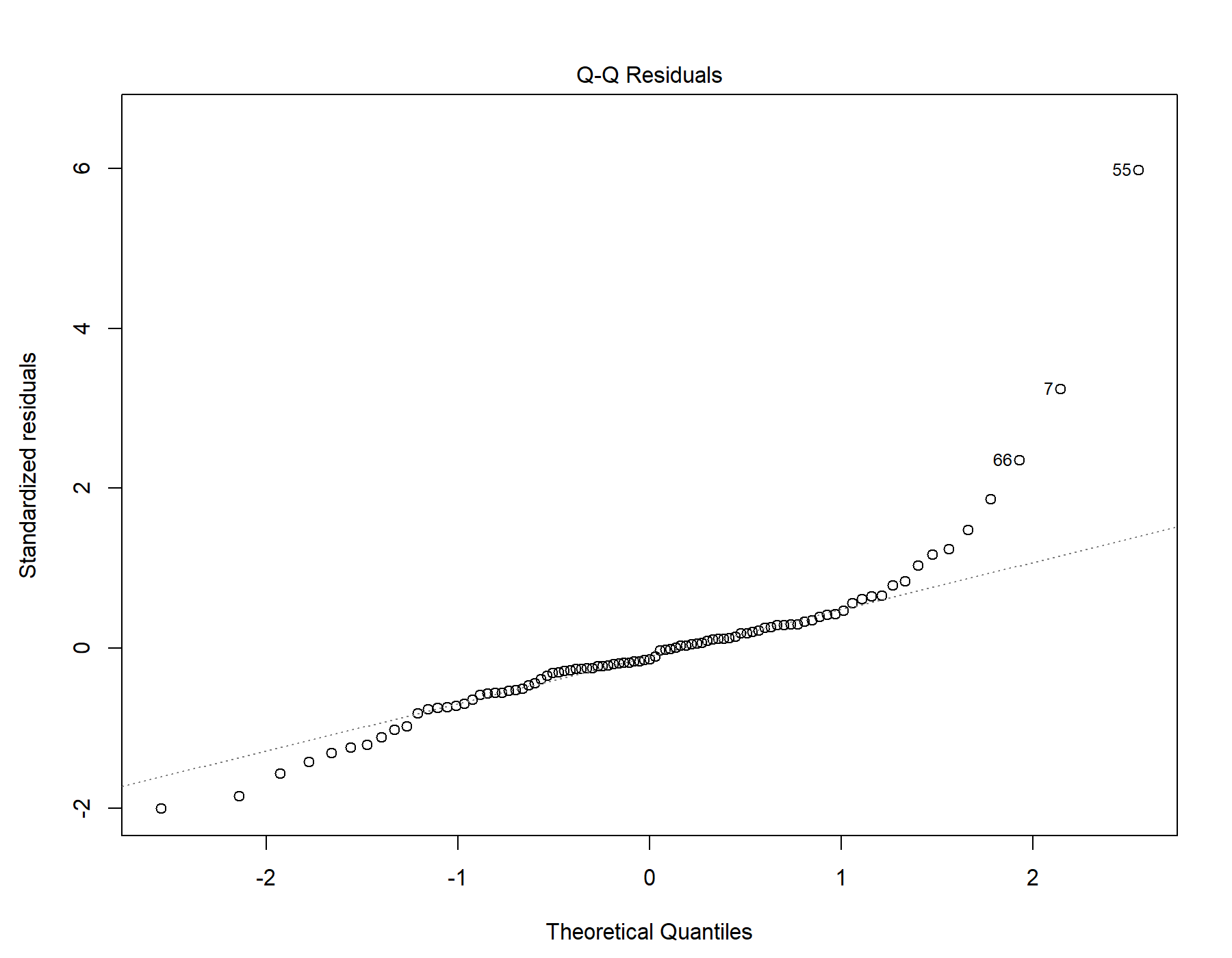

Supplemental Figure 69. Normal distribution quantile-quantile plot for T1w-alone WML segmentation.

#### 33.5. Residuals vs leverage

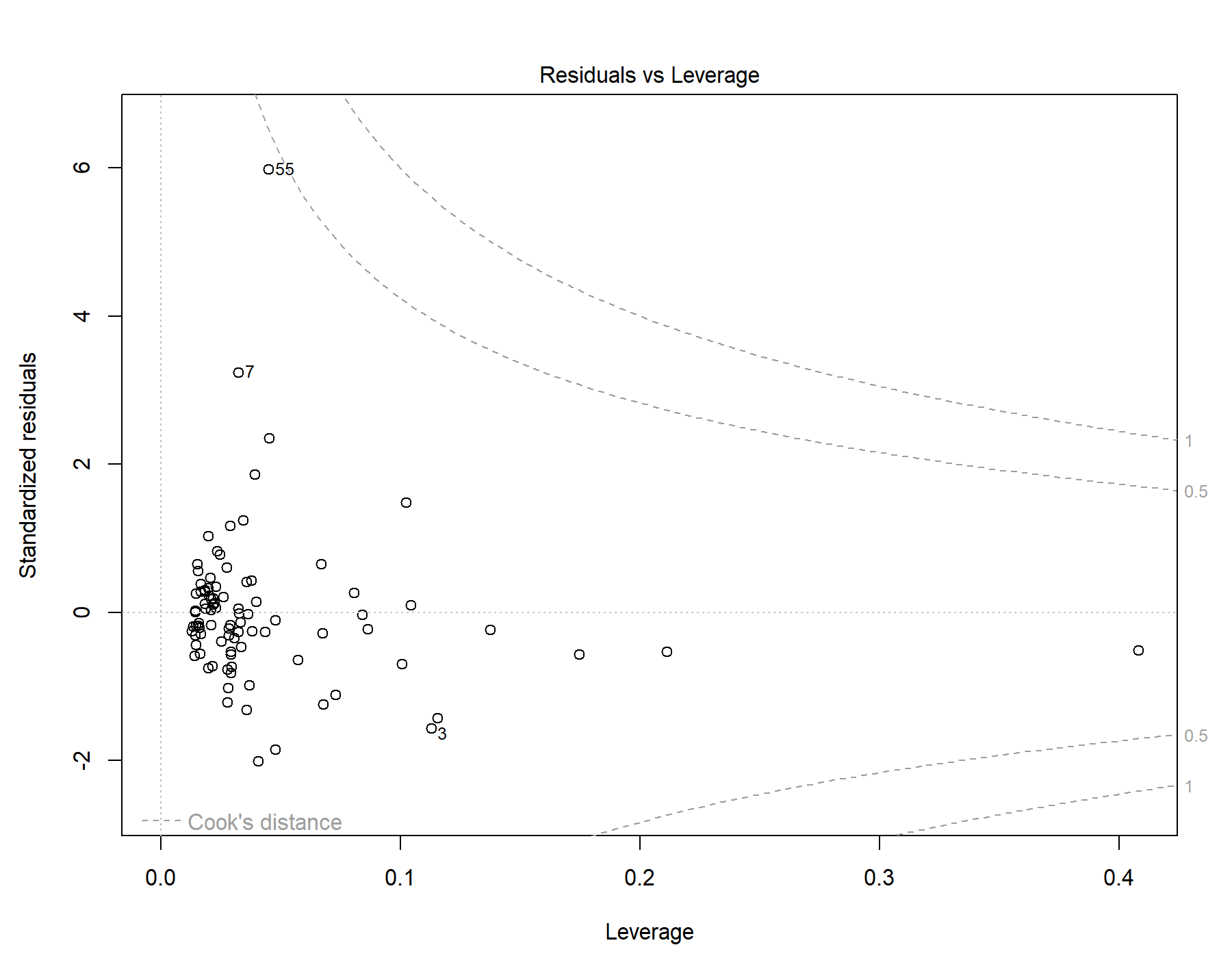

Supplemental Figure 70. Residuals vs leverage diagnostics for T1w-alone WML segmentation.

### 34. Global | Male only | T1w+FLAIR WML vol | SDNN.HF._log | Exploratory restricted interaction model

Model: T1w+FLAIR WML vol ~ SDNN.HF._log + Age +	Age * SDNN.HF._log

#### 34.1. Standardized coefficients

| **Term** | **Estimate** | **Std. Error** | **t** | **p** | **β** |
| --- | --- | --- | --- | --- | --- |
| (Intercept) | -7.102e-04 | 1.093e-04 | -6.50 | <.001 |  |
| SDNN.HF._log | 2.434e-04 | 3.194e-04 | 0.762 | .448 | 0.106 |
| Age | 1.593e-04 | 1.966e-05 | 8.10 | <.001 | 0.650 |
| SDNN.HF._log:Age | -7.423e-05 | 4.200e-05 | -1.77 | .081 | -0.244 |

***Supplemental Table 109.*** *Standardized coefficients values for T1w+FLAIR WML segmentation.*

#### 34.2. 95% confidence intervals

| **Term** | **Lower 95%** | **Upper 95%** |
| --- | --- | --- |
| (Intercept) | -9.273e-04 | -4.932e-04 |
| SDNN.HF._log | -3.912e-04 | 8.780e-04 |
| Age | 1.202e-04 | 1.983e-04 |
| SDNN.HF._log:Age | -1.577e-04 | 9.212e-06 |

***Supplemental Table 110.*** *95% confidence intervals values for T1w+FLAIR WML segmentation.*

#### 34.3. Breusch-Pagan test

| **Test** | **Statistic** | **df** | **p** |
| --- | --- | --- | --- |
| Breusch-Pagan | 11.111 | 3 | .011 |

***Supplemental Table 111.*** *Breusch-Pagan test values for T1w+FLAIR WML segmentation.*

#### 34.4. Normal Q-Q plot

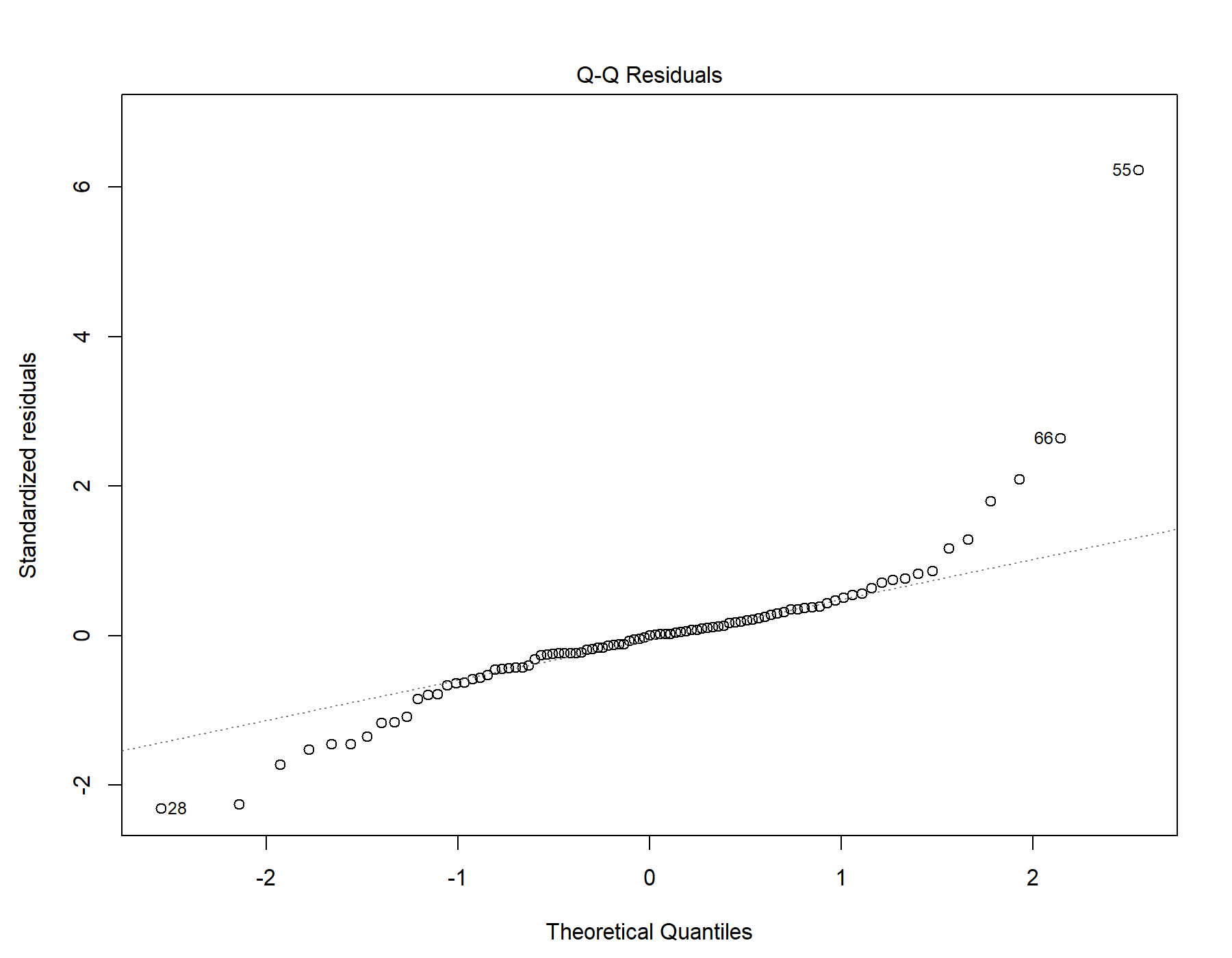

Supplemental Figure 71. Normal distribution quantile-quantile plot for T1w+FLAIR WML segmentation.

#### 34.5. Residuals vs leverage

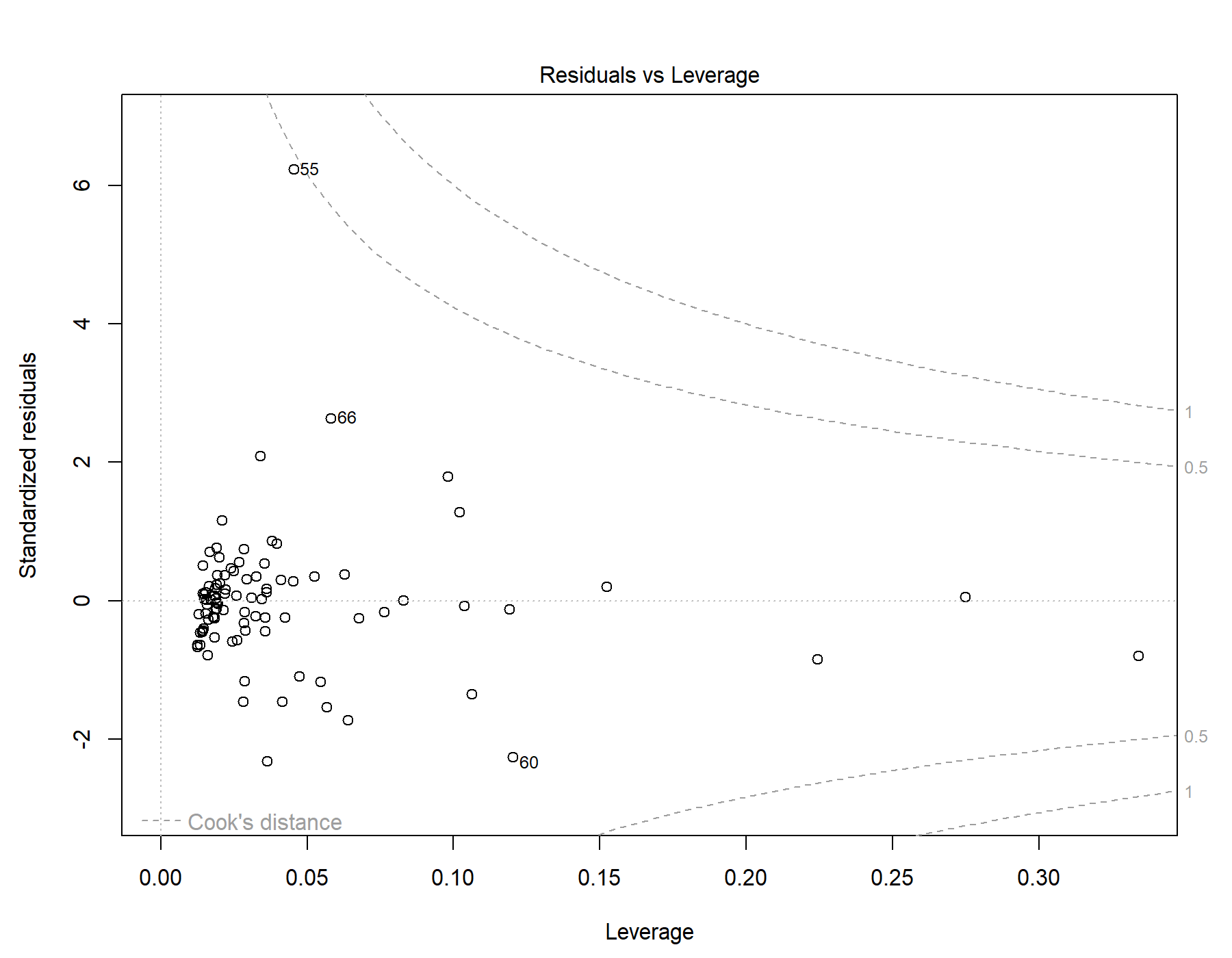

Supplemental Figure 72. Residuals vs leverage diagnostics for T1w+FLAIR WML segmentation.

### 35. Global | Male only | T1w+T2w WML vol | SDNN.HF._log | Exploratory restricted interaction model

Model: T1w+T2w WML vol ~ SDNN.HF._log + Age +	Age * SDNN.HF._log

#### 35.1. Standardized coefficients

| **Term** | **Estimate** | **Std. Error** | **t** | **p** | **β** |
| --- | --- | --- | --- | --- | --- |
| (Intercept) | -8.491e-04 | 1.160e-04 | -7.32 | <.001 |  |
| SDNN.HF._log | 2.229e-04 | 3.391e-04 | 0.657 | .513 | 0.0857 |
| Age | 1.920e-04 | 2.088e-05 | 9.20 | <.001 | 0.692 |
| SDNN.HF._log:Age | -7.773e-05 | 4.459e-05 | -1.74 | .085 | -0.225 |

***Supplemental Table 112.*** *Standardized coefficients values for T1w+T2w WML segmentation.*

#### 35.2. 95% confidence intervals

| **Term** | **Lower 95%** | **Upper 95%** |
| --- | --- | --- |
| (Intercept) | -0.00108 | -6.186e-04 |
| SDNN.HF._log | -4.510e-04 | 8.967e-04 |
| Age | 1.505e-04 | 2.335e-04 |
| SDNN.HF._log:Age | -1.663e-04 | 1.088e-05 |

***Supplemental Table 113.*** *95% confidence intervals values for T1w+T2w WML segmentation.*

#### 35.3. Breusch-Pagan test

| **Test** | **Statistic** | **df** | **p** |
| --- | --- | --- | --- |
| Breusch-Pagan | 8.873 | 3 | .031 |

***Supplemental Table 114.*** *Breusch-Pagan test values for T1w+T2w WML segmentation.*

#### 35.4. Normal Q-Q plot

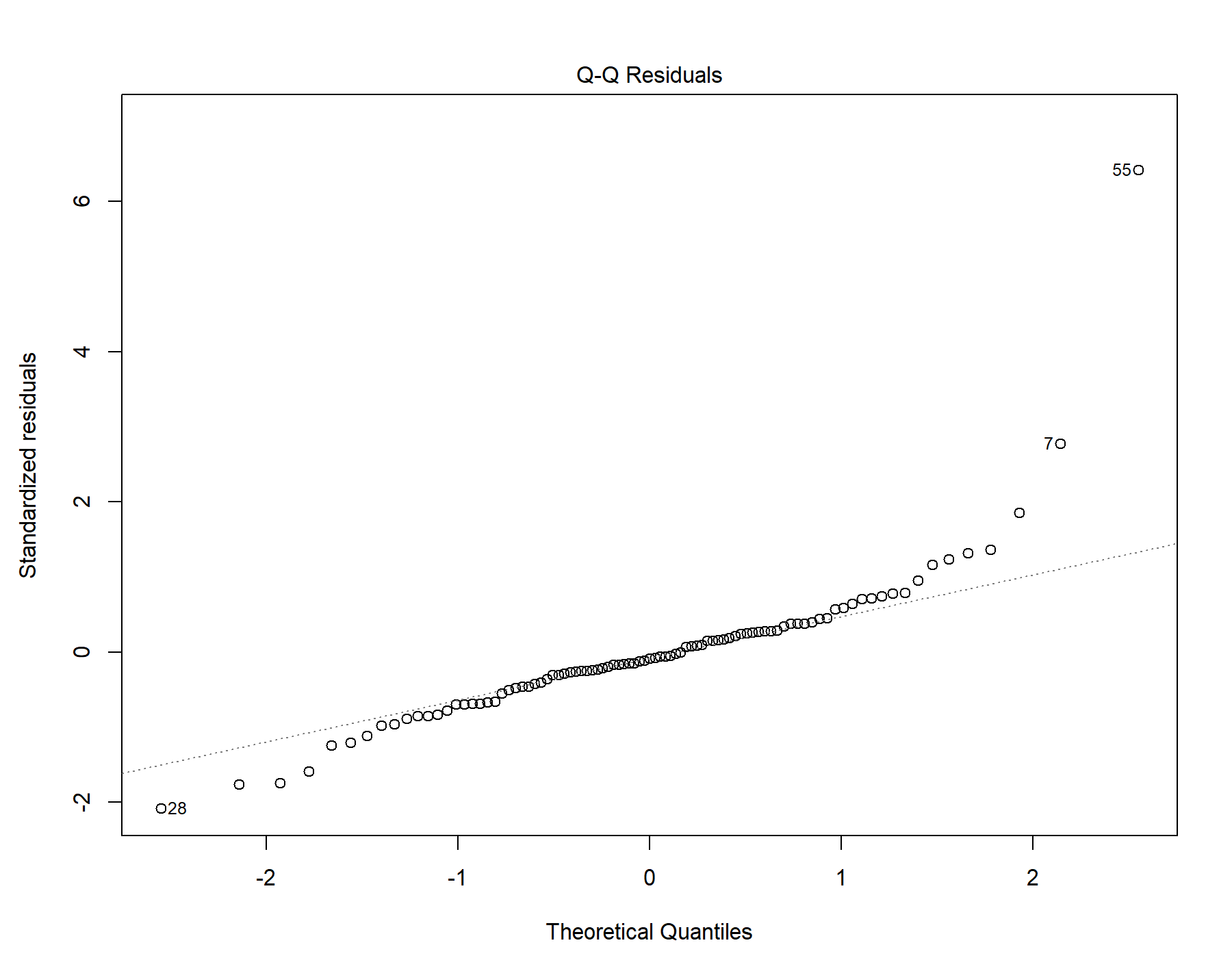

Supplemental Figure 73. Normal distribution quantile-quantile plot for T1w+T2w WML segmentation.

#### 35.5. Residuals vs leverage

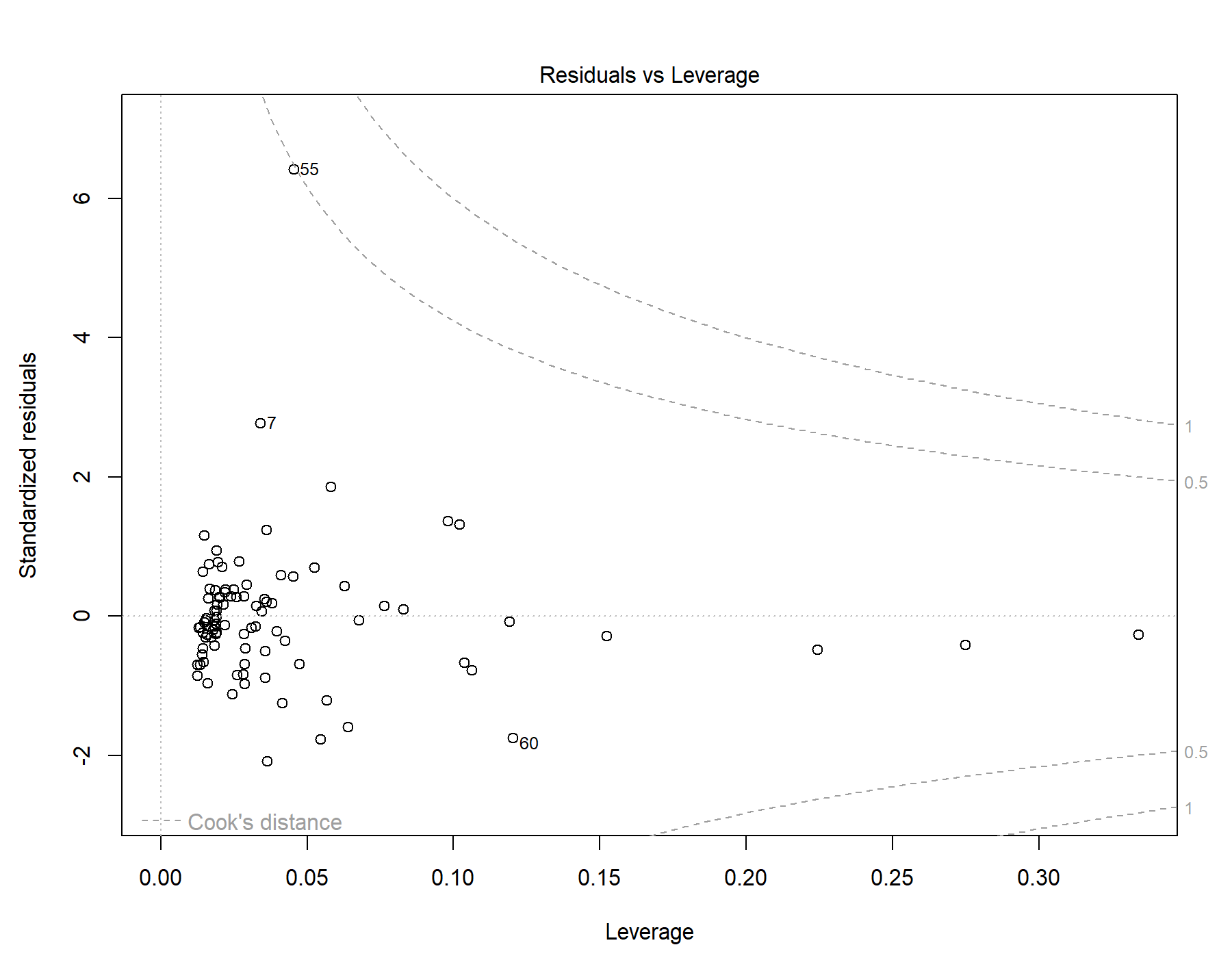

Supplemental Figure 74. Residuals vs leverage diagnostics for T1w+T2w WML segmentation.

### 36. Global | Male only | T1w-alone WML vol | SDNN.HF._log | Exploratory restricted interaction model

Model: T1w-alone WML vol ~ SDNN.HF._log + Age + Age *	SDNN.HF._log

#### 36.1. Standardized coefficients

| **Term** | **Estimate** | **Std. Error** | **t** | **p** | **β** |
| --- | --- | --- | --- | --- | --- |
| (Intercept) | -6.820e-04 | 9.641e-05 | -7.07 | <.001 |  |
| SDNN.HF._log | 1.739e-04 | 2.818e-04 | 0.617 | .539 | 0.0824 |
| Age | 1.549e-04 | 1.735e-05 | 8.93 | <.001 | 0.688 |
| SDNN.HF._log:Age | -5.717e-05 | 3.706e-05 | -1.54 | .126 | -0.204 |

***Supplemental Table 115.*** *Standardized coefficients values for T1w-alone WML segmentation.*

#### 36.2. 95% confidence intervals

| **Term** | **Lower 95%** | **Upper 95%** |
| --- | --- | --- |
| (Intercept) | -8.736e-04 | -4.904e-04 |
| SDNN.HF._log | -3.861e-04 | 7.339e-04 |
| Age | 1.205e-04 | 1.894e-04 |
| SDNN.HF._log:Age | -1.308e-04 | 1.646e-05 |

***Supplemental Table 116.*** *95% confidence intervals values for T1w-alone WML segmentation.*

#### 36.3. Breusch-Pagan test

| **Test** | **Statistic** | **df** | **p** |
| --- | --- | --- | --- |
| Breusch-Pagan | 10.966 | 3 | .012 |

***Supplemental Table 117.*** *Breusch-Pagan test values for T1w-alone WML segmentation.*

#### 36.4. Normal Q-Q plot

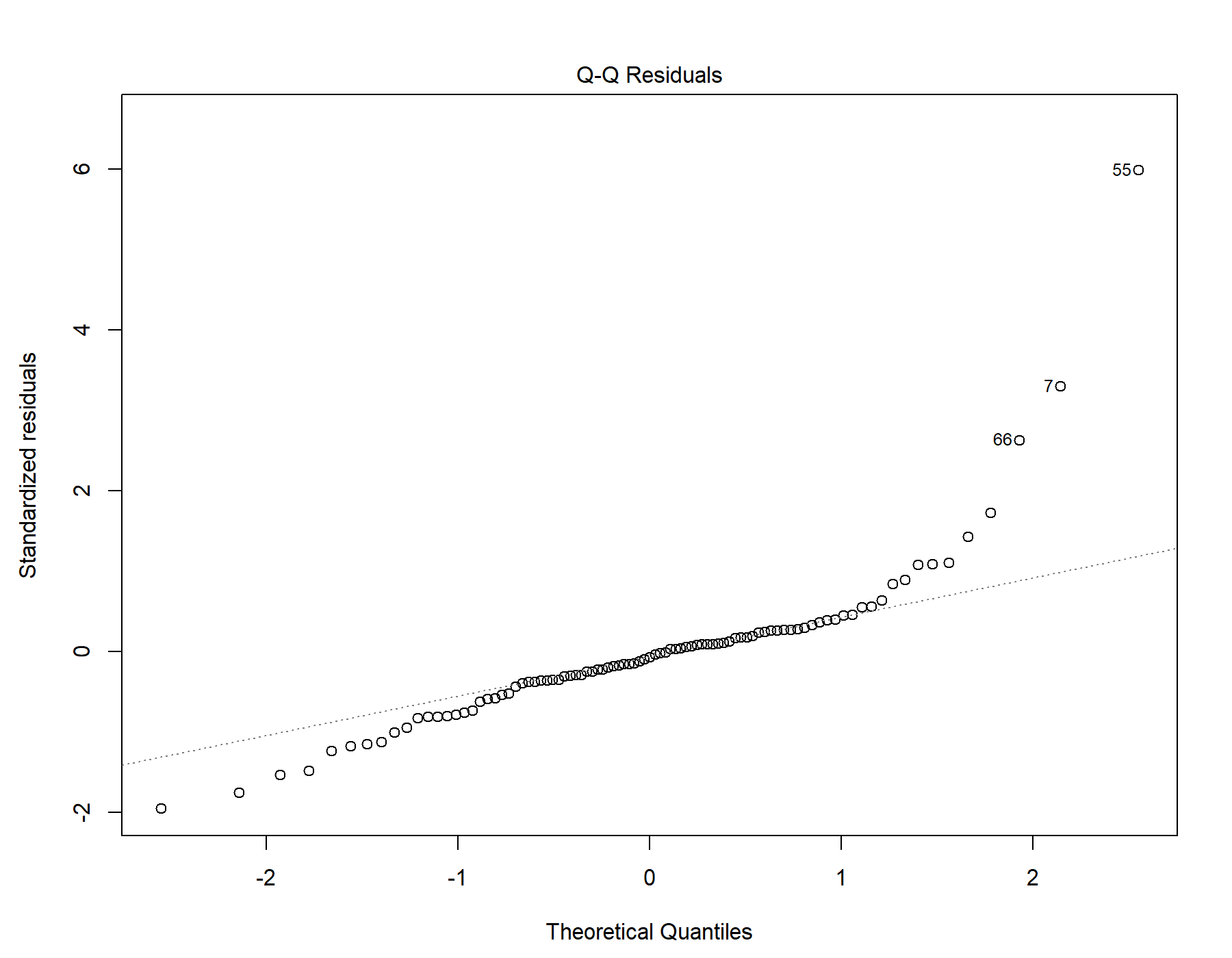

Supplemental Figure 75. Normal distribution quantile-quantile plot for T1w-alone WML segmentation.

#### 36.5. Residuals vs leverage

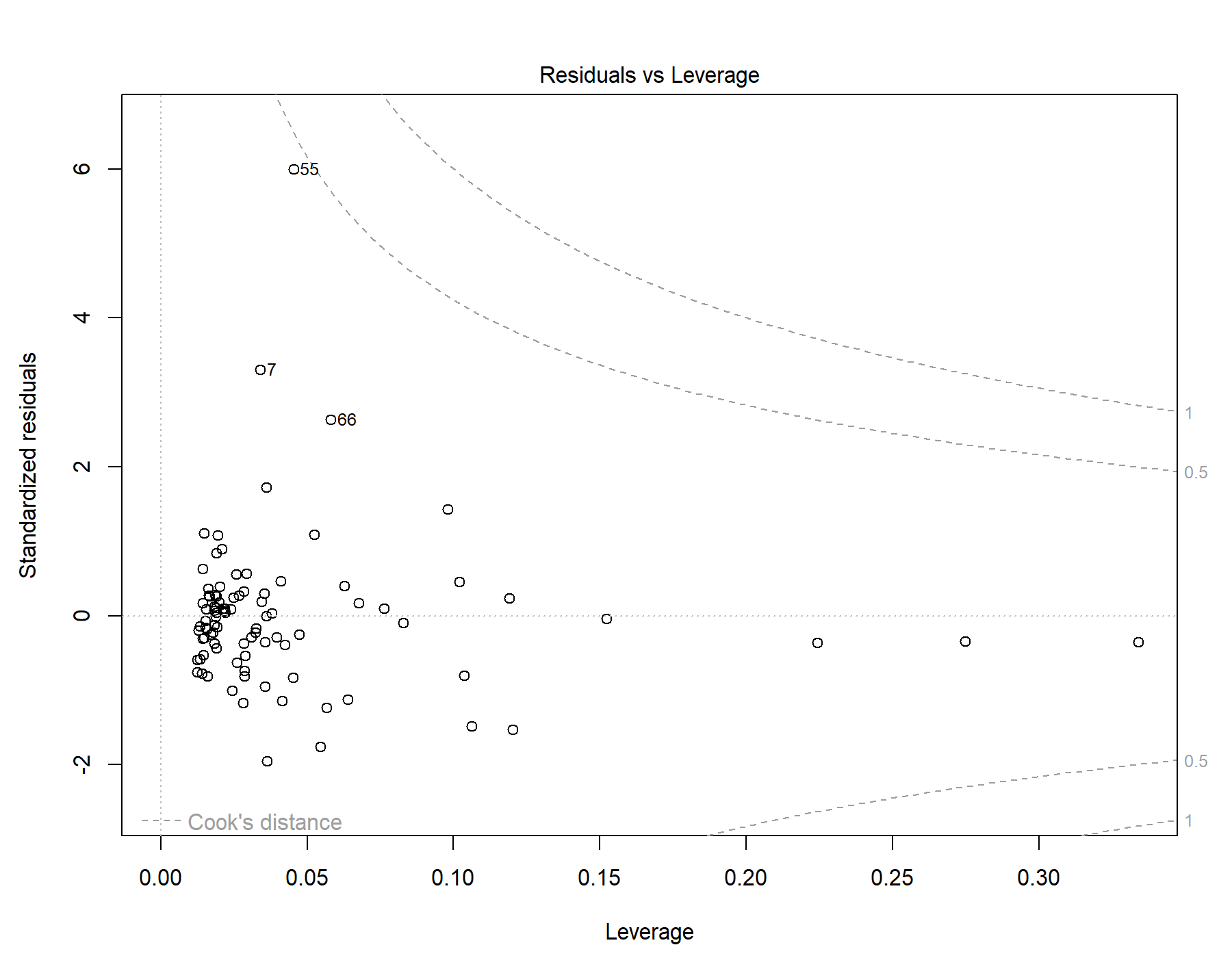

Supplemental Figure 76. Residuals vs leverage diagnostics for T1w-alone WML segmentation.

### 37. Global | Young only | T1w+FLAIR WML vol | RMSSD | Exploratory restricted interaction model

Model: T1w+FLAIR WML vol ~ RMSSD + Age + Sex +	Age * RMSSD + Sex * RMSSD

#### 37.1. Standardized coefficients

| **Term** | **Estimate** | **Std. Error** | **t** | **p** | **β** |
| --- | --- | --- | --- | --- | --- |
| (Intercept) | -7.639e-05 | 1.166e-04 | -0.655 | .514 |  |
| RMSSD | -1.710e-06 | 3.266e-06 | -0.524 | .602 | -0.214 |
| Age | 6.499e-05 | 4.464e-05 | 1.46 | .149 | 0.155 |
| Sex | -2.701e-05 | 6.223e-05 | -0.434 | .665 | -0.0450 |
| RMSSD:Age | -1.348e-06 | 1.597e-06 | -0.844 | .401 | -0.244 |
| RMSSD:Sex | 1.865e-06 | 1.773e-06 | 1.05 | .296 | 0.404 |

***Supplemental Table 118.*** *Standardized coefficients values for T1w+FLAIR WML segmentation.*

#### 37.2. 95% confidence intervals

| **Term** | **Lower 95%** | **Upper 95%** |
| --- | --- | --- |
| (Intercept) | -3.078e-04 | 1.551e-04 |
| RMSSD | -8.190e-06 | 4.771e-06 |
| Age | -2.361e-05 | 1.536e-04 |
| Sex | -1.505e-04 | 9.648e-05 |
| RMSSD:Age | -4.518e-06 | 1.822e-06 |
| RMSSD:Sex | -1.654e-06 | 5.383e-06 |

***Supplemental Table 119.*** *95% confidence intervals values for T1w+FLAIR WML segmentation.*

#### 37.3. Breusch-Pagan test

| **Test** | **Statistic** | **df** | **p** |
| --- | --- | --- | --- |
| Breusch-Pagan | 5.962 | 5 | .310 |

***Supplemental Table 120.*** *Breusch-Pagan test values for T1w+FLAIR WML segmentation.*

#### 37.4. Normal Q-Q plot

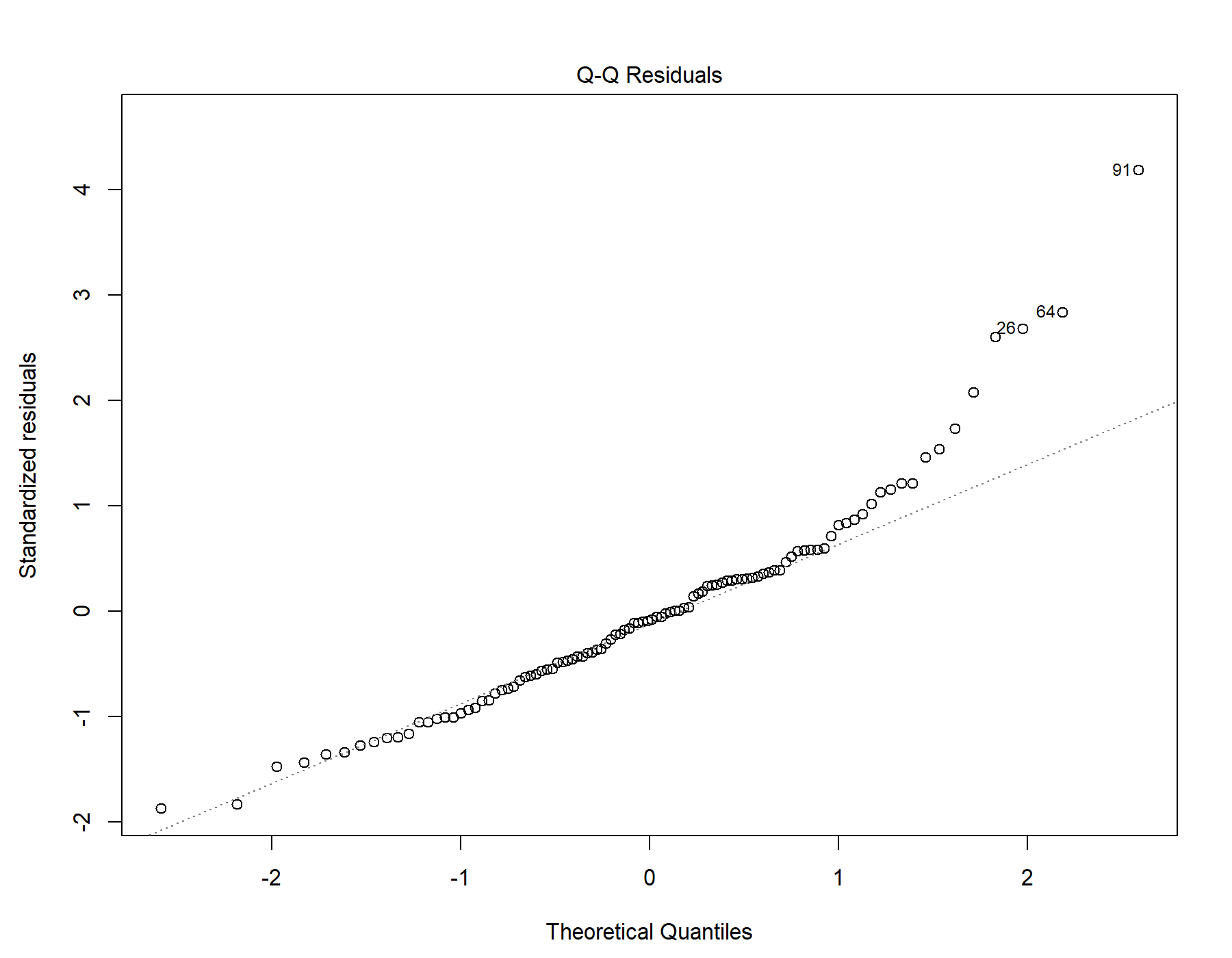

Supplemental Figure 77. Normal distribution quantile-quantile plot for T1w+FLAIR WML segmentation.

#### 37.5. Residuals vs leverage

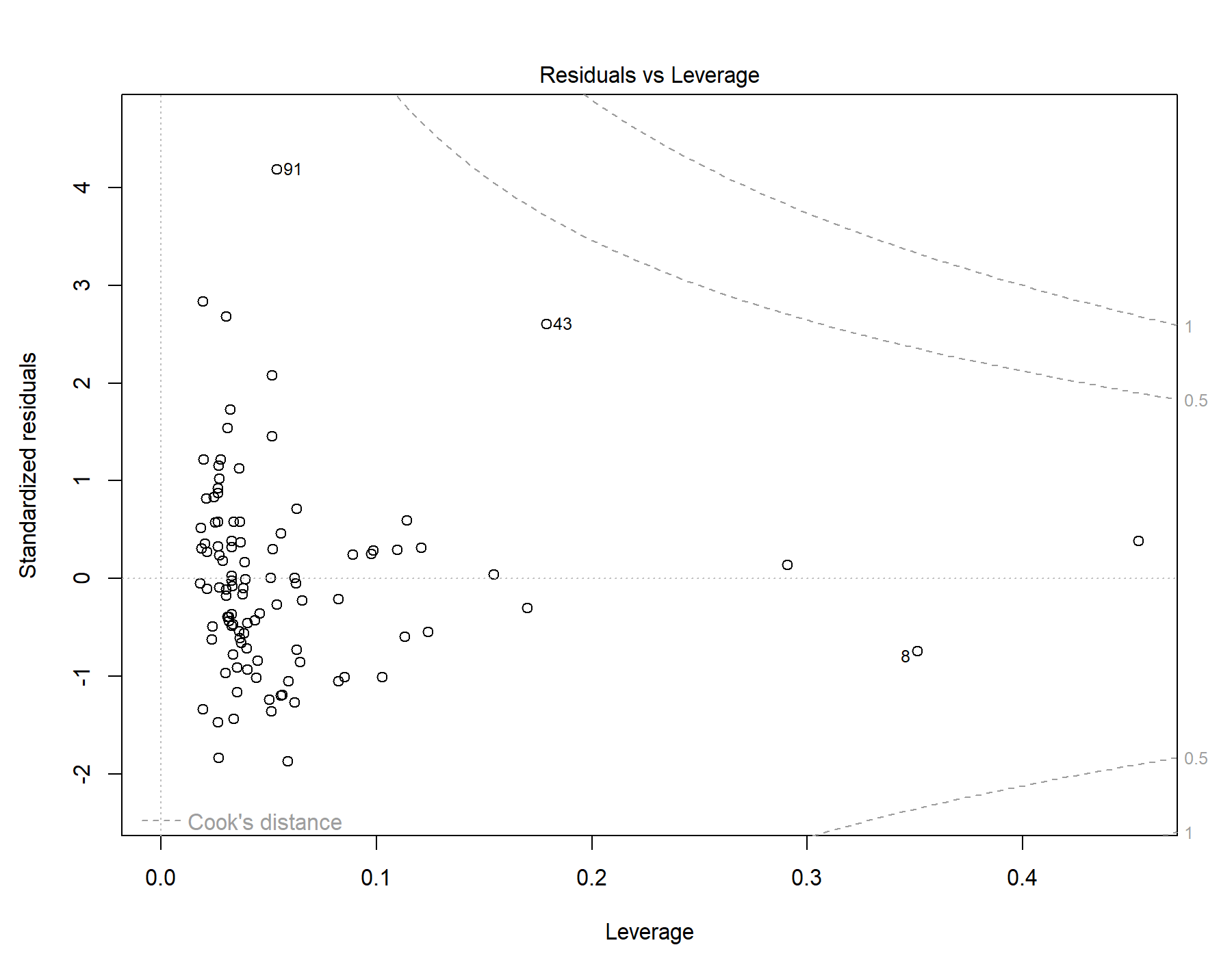

Supplemental Figure 78. Residuals vs leverage diagnostics for T1w+FLAIR WML segmentation.

### 38. Global | Young only | T1w+T2w WML vol | RMSSD | Exploratory restricted interaction model

Model: T1w+T2w WML vol ~ RMSSD + Age + Sex + Age *	RMSSD + Sex * RMSSD

#### 38.1. Standardized coefficients

| **Term** | **Estimate** | **Std. Error** | **t** | **p** | **β** |
| --- | --- | --- | --- | --- | --- |
| (Intercept) | 2.844e-05 | 1.370e-04 | 0.208 | .836 |  |
| RMSSD | 1.577e-06 | 3.837e-06 | 0.411 | .682 | 0.168 |
| Age | 7.579e-05 | 5.245e-05 | 1.44 | .152 | 0.153 |
| Sex | -1.028e-04 | 7.311e-05 | -1.41 | .163 | -0.145 |
| RMSSD:Age | -1.932e-06 | 1.877e-06 | -1.03 | .306 | -0.297 |
| RMSSD:Sex | 1.119e-06 | 2.083e-06 | 0.537 | .593 | 0.206 |

***Supplemental Table 121.*** *Standardized coefficients values for T1w+T2w WML segmentation.*

#### 38.2. 95% confidence intervals

| **Term** | **Lower 95%** | **Upper 95%** |
| --- | --- | --- |
| (Intercept) | -2.435e-04 | 3.004e-04 |
| RMSSD | -6.037e-06 | 9.192e-06 |
| Age | -2.830e-05 | 1.799e-04 |
| Sex | -2.479e-04 | 4.230e-05 |
| RMSSD:Age | -5.656e-06 | 1.793e-06 |
| RMSSD:Sex | -3.015e-06 | 5.252e-06 |

***Supplemental Table 122.*** *95% confidence intervals values for T1w+T2w WML segmentation.*

#### 38.3. Breusch-Pagan test

| **Test** | **Statistic** | **df** | **p** |
| --- | --- | --- | --- |
| Breusch-Pagan | 3.222 | 5 | .666 |

***Supplemental Table 123.*** *Breusch-Pagan test values for T1w+T2w WML segmentation.*

#### 38.4. Normal Q-Q plot

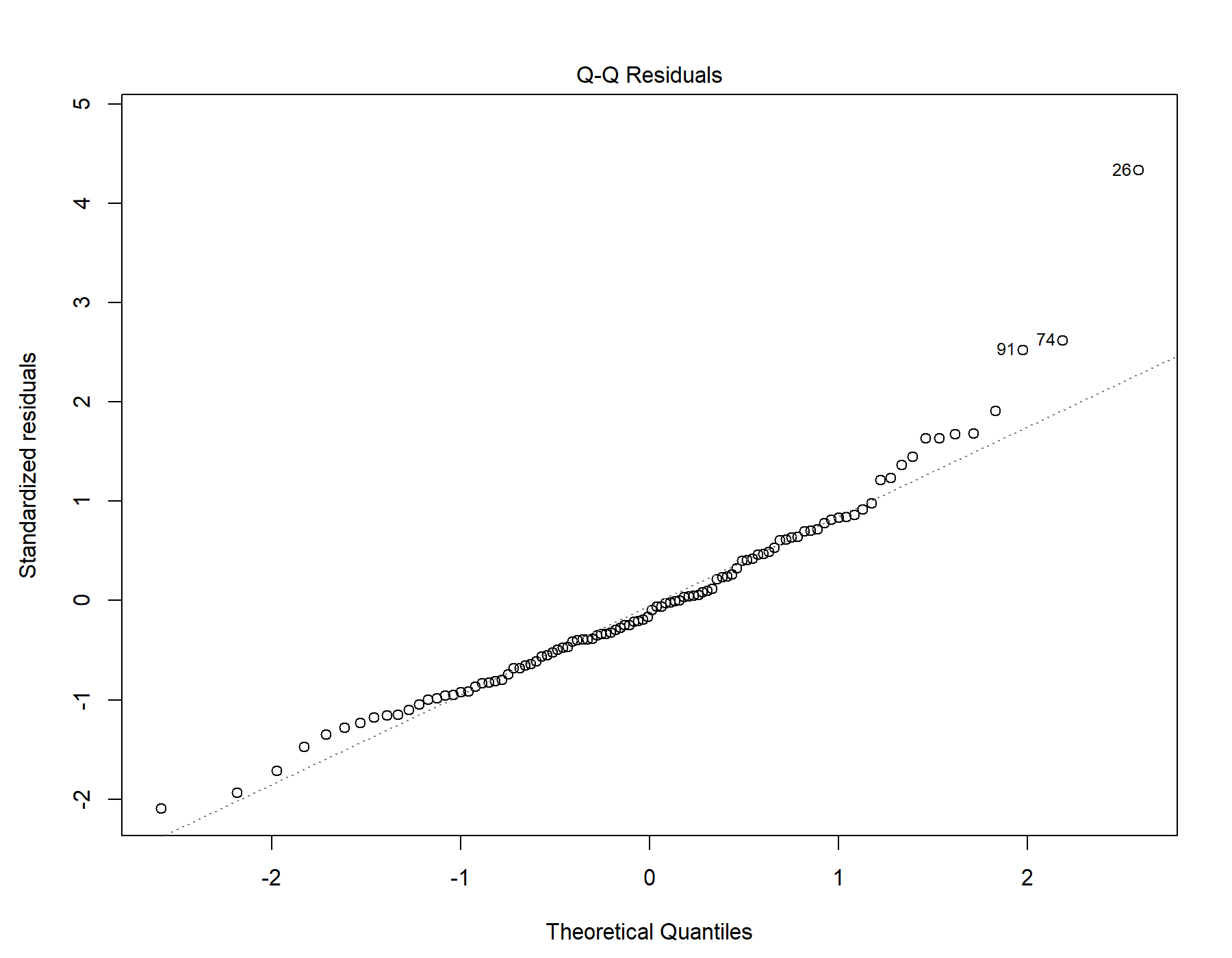

Supplemental Figure 79. Normal distribution quantile-quantile plot for T1w+T2w WML segmentation.

#### 38.5. Residuals vs leverage

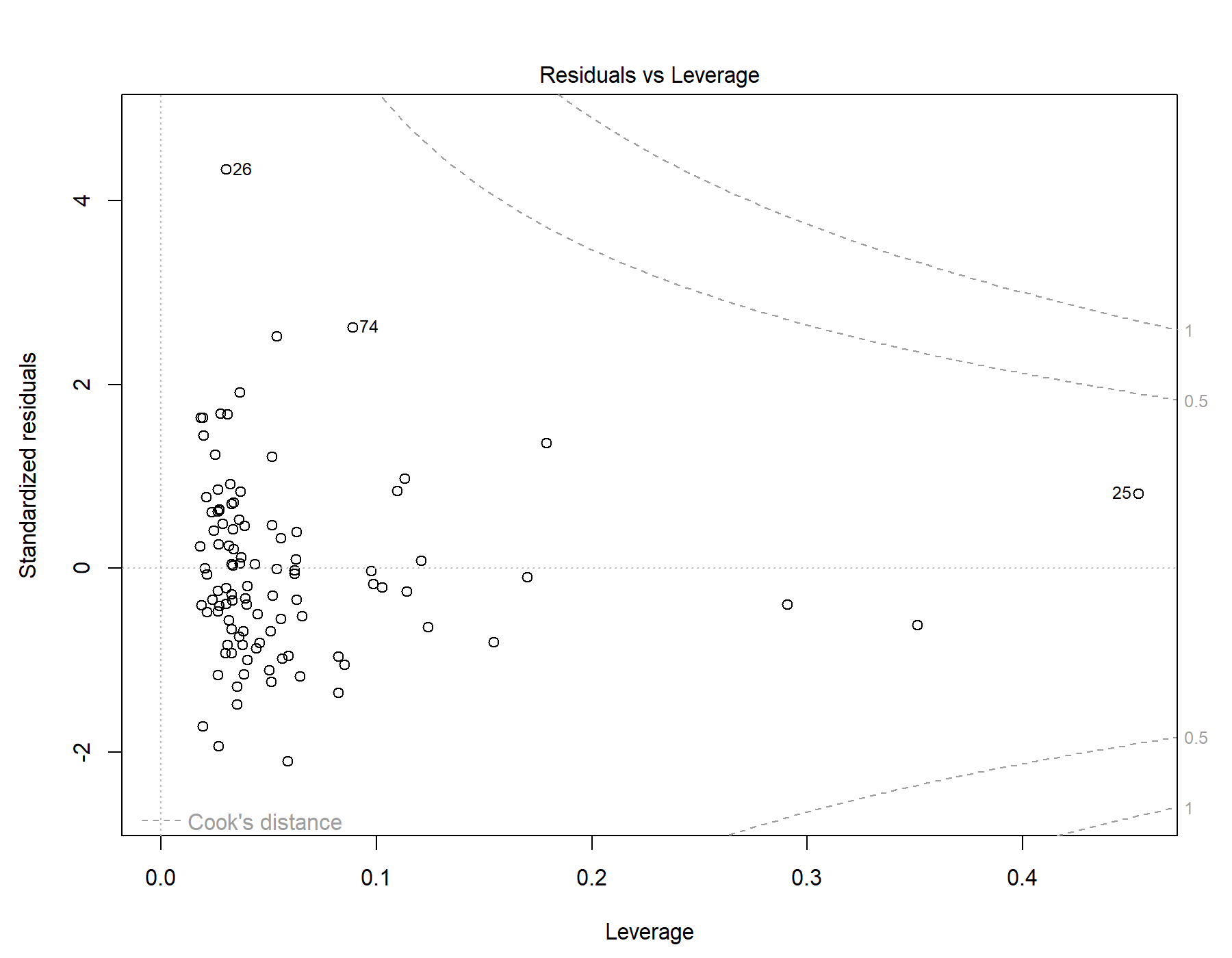

Supplemental Figure 80. Residuals vs leverage diagnostics for T1w+T2w WML segmentation.

### 39. Global | Young only | T1w-alone WML vol | RMSSD | Exploratory restricted interaction model

Model: T1w-alone WML vol ~ RMSSD + Age + Sex + Age *	RMSSD + Sex * RMSSD

#### 39.1. Standardized coefficients

| **Term** | **Estimate** | **Std. Error** | **t** | **p** | **β** |
| --- | --- | --- | --- | --- | --- |
| (Intercept) | 5.808e-05 | 1.208e-04 | 0.481 | .632 |  |
| RMSSD | 1.014e-06 | 3.382e-06 | 0.300 | .765 | 0.124 |
| Age | 3.990e-05 | 4.623e-05 | 0.863 | .390 | 0.0925 |
| Sex | -8.251e-05 | 6.444e-05 | -1.28 | .203 | -0.134 |
| RMSSD:Age | -1.421e-06 | 1.654e-06 | -0.859 | .392 | -0.251 |
| RMSSD:Sex | 8.584e-07 | 1.836e-06 | 0.468 | .641 | 0.181 |

***Supplemental Table 124.*** *Standardized coefficients values for T1w-alone WML segmentation.*

#### 39.2. 95% confidence intervals

| **Term** | **Lower 95%** | **Upper 95%** |
| --- | --- | --- |
| (Intercept) | -1.816e-04 | 2.978e-04 |
| RMSSD | -5.697e-06 | 7.725e-06 |
| Age | -5.184e-05 | 1.316e-04 |
| Sex | -2.104e-04 | 4.536e-05 |
| RMSSD:Age | -4.703e-06 | 1.862e-06 |
| RMSSD:Sex | -2.785e-06 | 4.502e-06 |

***Supplemental Table 125.*** *95% confidence intervals values for T1w-alone WML segmentation.*

#### 39.3. Breusch-Pagan test

| **Test** | **Statistic** | **df** | **p** |
| --- | --- | --- | --- |
| Breusch-Pagan | 3.616 | 5 | .606 |

***Supplemental Table 126.*** *Breusch-Pagan test values for T1w-alone WML segmentation.*

#### 39.4. Normal Q-Q plot

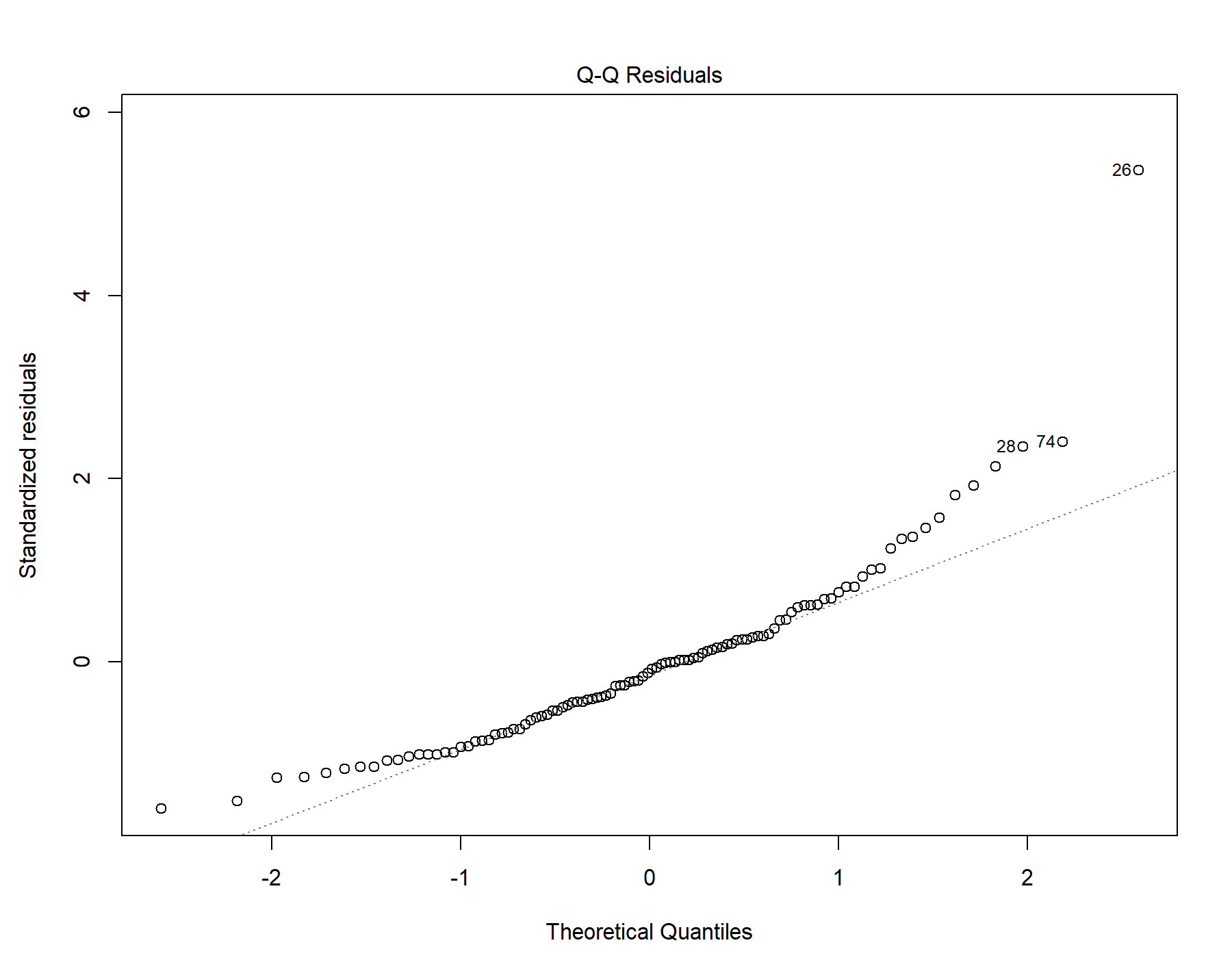

Supplemental Figure 81. Normal distribution quantile-quantile plot for T1w-alone WML segmentation.

#### 39.5. Residuals vs leverage

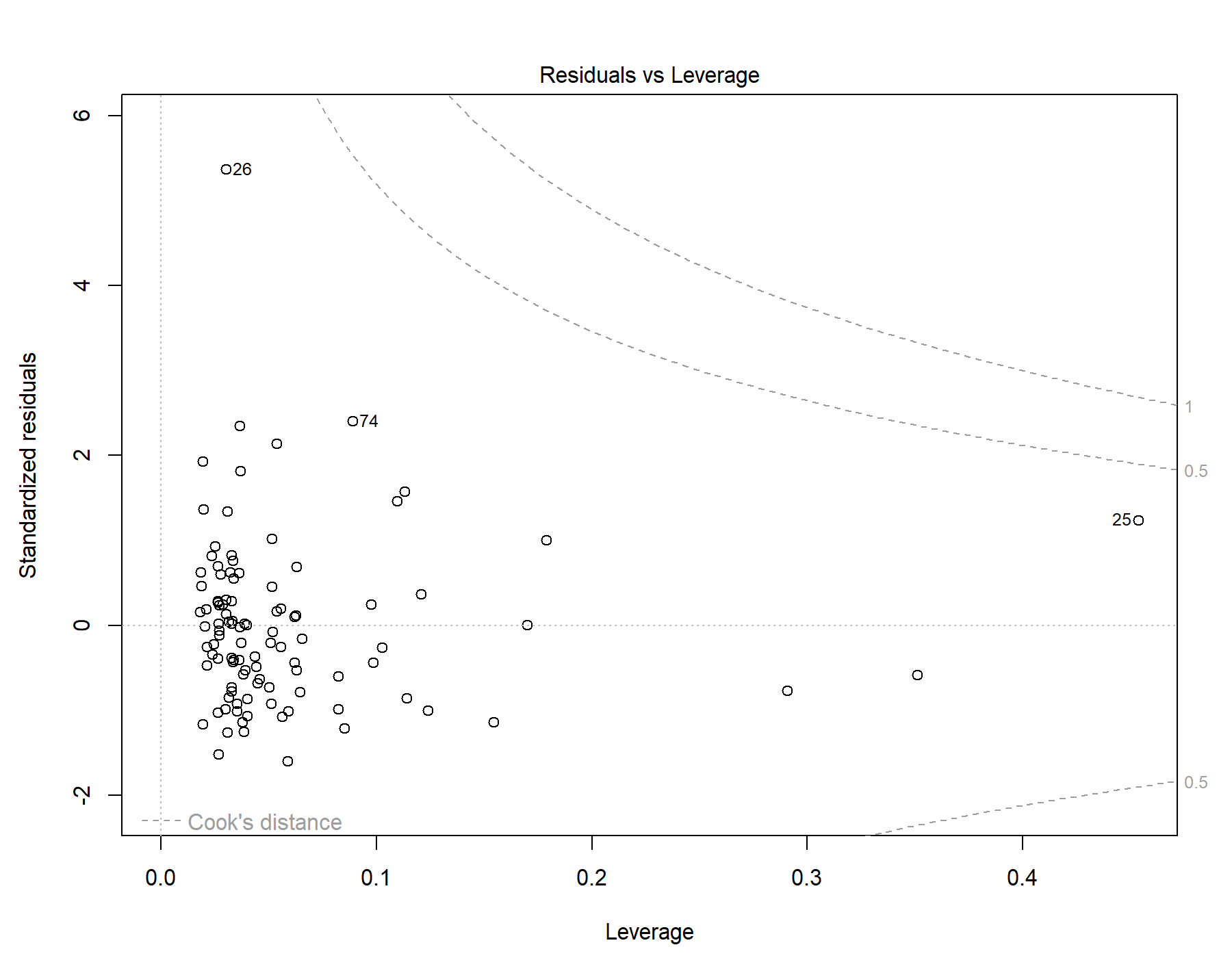

Supplemental Figure 82. Residuals vs leverage diagnostics for T1w-alone WML segmentation.

### 40. Global | Young only | T1w+FLAIR WML vol | SDNN.LF._log | Exploratory restricted interaction model

Model: T1w+FLAIR WML vol ~ SDNN.LF._log + Age + Sex + Age * SDNN.LF._log + Sex * SDNN.LF._log

#### 40.1. Standardized coefficients

| **Term** | **Estimate** | **Std. Error** | **t** | **p** | **β** |
| --- | --- | --- | --- | --- | --- |
| (Intercept) | -6.334e-05 | 1.119e-04 | -0.566 | .573 |  |
| SDNN.LF._log | -9.267e-04 | 3.514e-04 | -2.64 | .010 | -1.02 |
| Age | 8.060e-05 | 4.212e-05 | 1.91 | .059 | 0.192 |
| Sex | -4.379e-05 | 5.757e-05 | -0.761 | .449 | -0.0729 |
| SDNN.LF._log:Age | 1.220e-05 | 1.565e-04 | 0.0780 | .938 | 0.0210 |
| SDNN.LF._log:Sex | 5.346e-04 | 1.913e-04 | 2.79 | .006 | 1.03 |

***Supplemental Table 127.*** *Standardized coefficients values for T1w+FLAIR WML segmentation.*

#### 40.2. 95% confidence intervals

| **Term** | **Lower 95%** | **Upper 95%** |
| --- | --- | --- |
| (Intercept) | -2.855e-04 | 1.588e-04 |
| SDNN.LF._log | -0.00162 | -2.294e-04 |
| Age | -2.987e-06 | 1.642e-04 |
| Sex | -1.580e-04 | 7.046e-05 |
| SDNN.LF._log:Age | -2.983e-04 | 3.227e-04 |
| SDNN.LF._log:Sex | 1.550e-04 | 9.142e-04 |

***Supplemental Table 128.*** *95% confidence intervals values for T1w+FLAIR WML segmentation.*

#### 40.3. Breusch-Pagan test

| **Test** | **Statistic** | **df** | **p** |
| --- | --- | --- | --- |
| Breusch-Pagan | 17.424 | 5 | .004 |

***Supplemental Table 129.*** *Breusch-Pagan test values for T1w+FLAIR WML segmentation.*

#### 40.4. Normal Q-Q plot

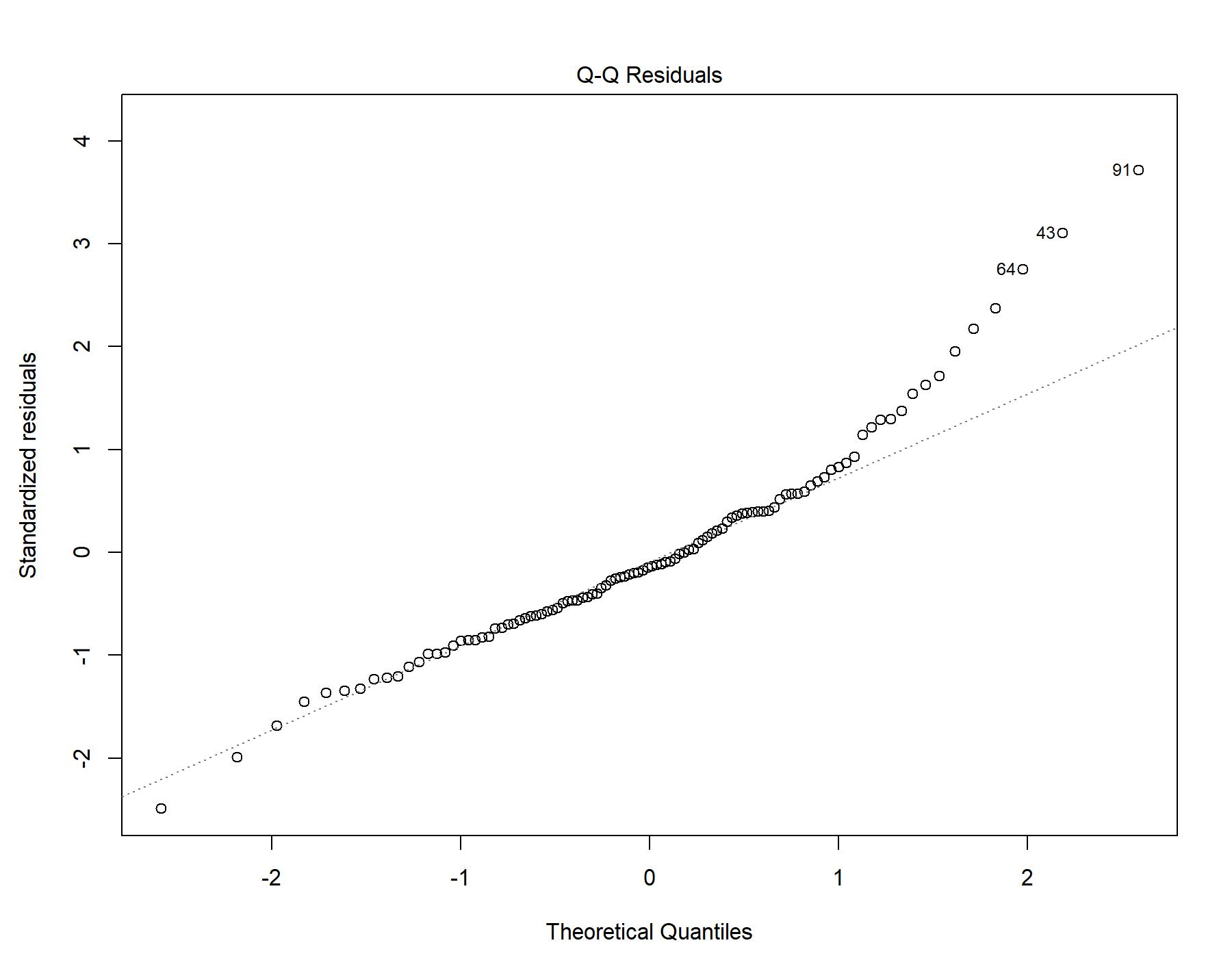

Supplemental Figure 83. Normal distribution quantile-quantile plot for T1w+FLAIR WML segmentation.

#### 40.5. Residuals vs leverage

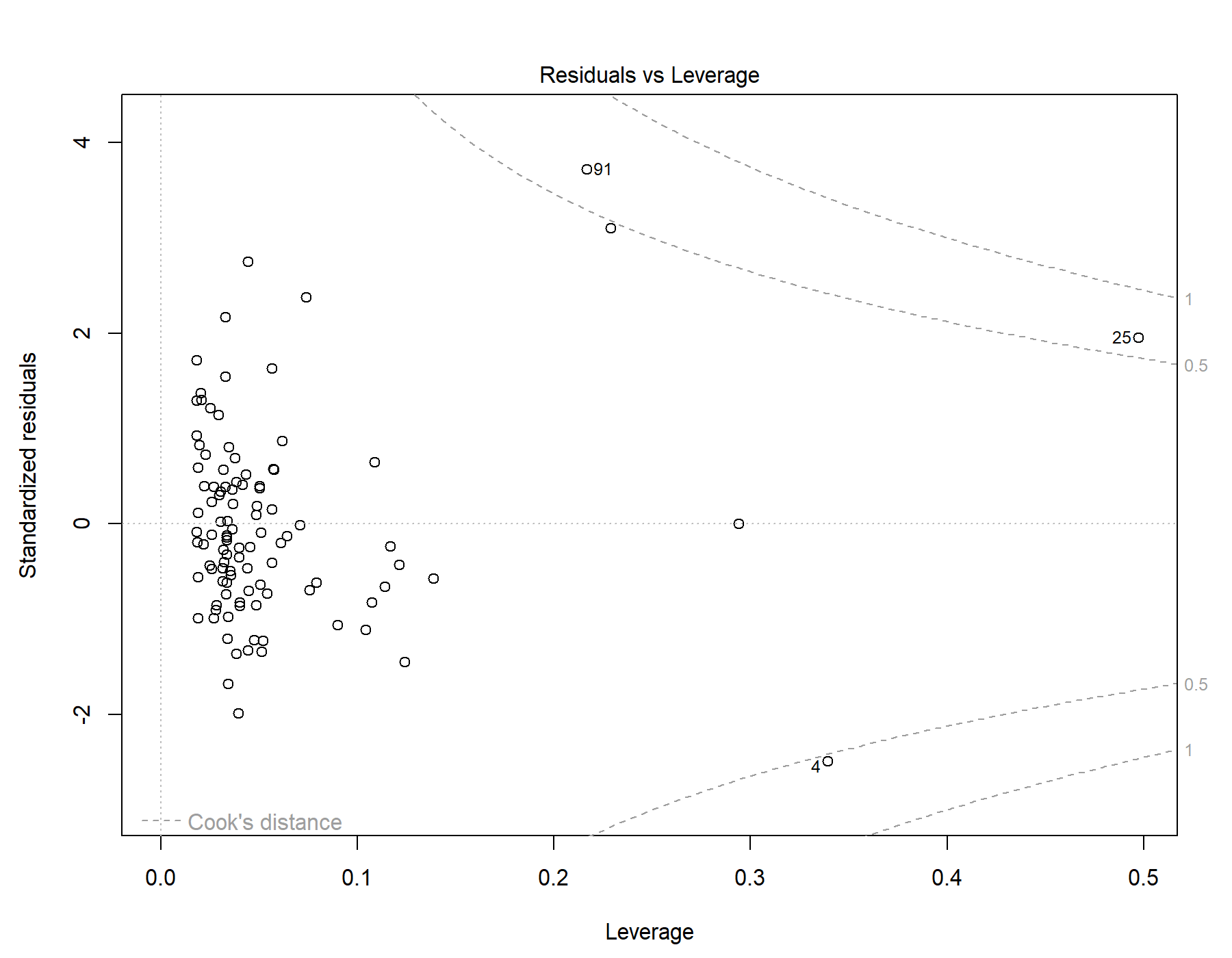

Supplemental Figure 84. Residuals vs leverage diagnostics for T1w+FLAIR WML segmentation.

### 41. Global | Young only | T1w+T2w WML vol | SDNN.LF._log | Exploratory restricted interaction model

Model: T1w+T2w WML vol ~ SDNN.LF._log + Age + Sex + Age * SDNN.LF._log + Sex * SDNN.LF._log

#### 41.1. Standardized coefficients

| **Term** | **Estimate** | **Std. Error** | **t** | **p** | **β** |
| --- | --- | --- | --- | --- | --- |
| (Intercept) | 1.656e-05 | 1.357e-04 | 0.122 | .903 |  |
| SDNN.LF._log | -5.203e-04 | 4.259e-04 | -1.22 | .225 | -0.487 |
| Age | 9.396e-05 | 5.106e-05 | 1.84 | .069 | 0.190 |
| Sex | -1.026e-04 | 6.978e-05 | -1.47 | .145 | -0.145 |
| SDNN.LF._log:Age | 1.109e-04 | 1.897e-04 | 0.585 | .560 | 0.162 |
| SDNN.LF._log:Sex | 2.595e-04 | 2.319e-04 | 1.12 | .266 | 0.424 |

***Supplemental Table 130.*** *Standardized coefficients values for T1w+T2w WML segmentation.*

#### 41.2. 95% confidence intervals

| **Term** | **Lower 95%** | **Upper 95%** |
| --- | --- | --- |
| (Intercept) | -2.527e-04 | 2.858e-04 |
| SDNN.LF._log | -0.00137 | 3.249e-04 |
| Age | -7.355e-06 | 1.953e-04 |
| Sex | -2.411e-04 | 3.585e-05 |
| SDNN.LF._log:Age | -2.655e-04 | 4.873e-04 |
| SDNN.LF._log:Sex | -2.007e-04 | 7.196e-04 |

***Supplemental Table 131.*** *95% confidence intervals values for T1w+T2w WML segmentation.*

#### 41.3. Breusch-Pagan test

| **Test** | **Statistic** | **df** | **p** |
| --- | --- | --- | --- |
| Breusch-Pagan | 6.211 | 5 | .286 |

***Supplemental Table 132.*** *Breusch-Pagan test values for T1w+T2w WML segmentation.*

#### 41.4. Normal Q-Q plot

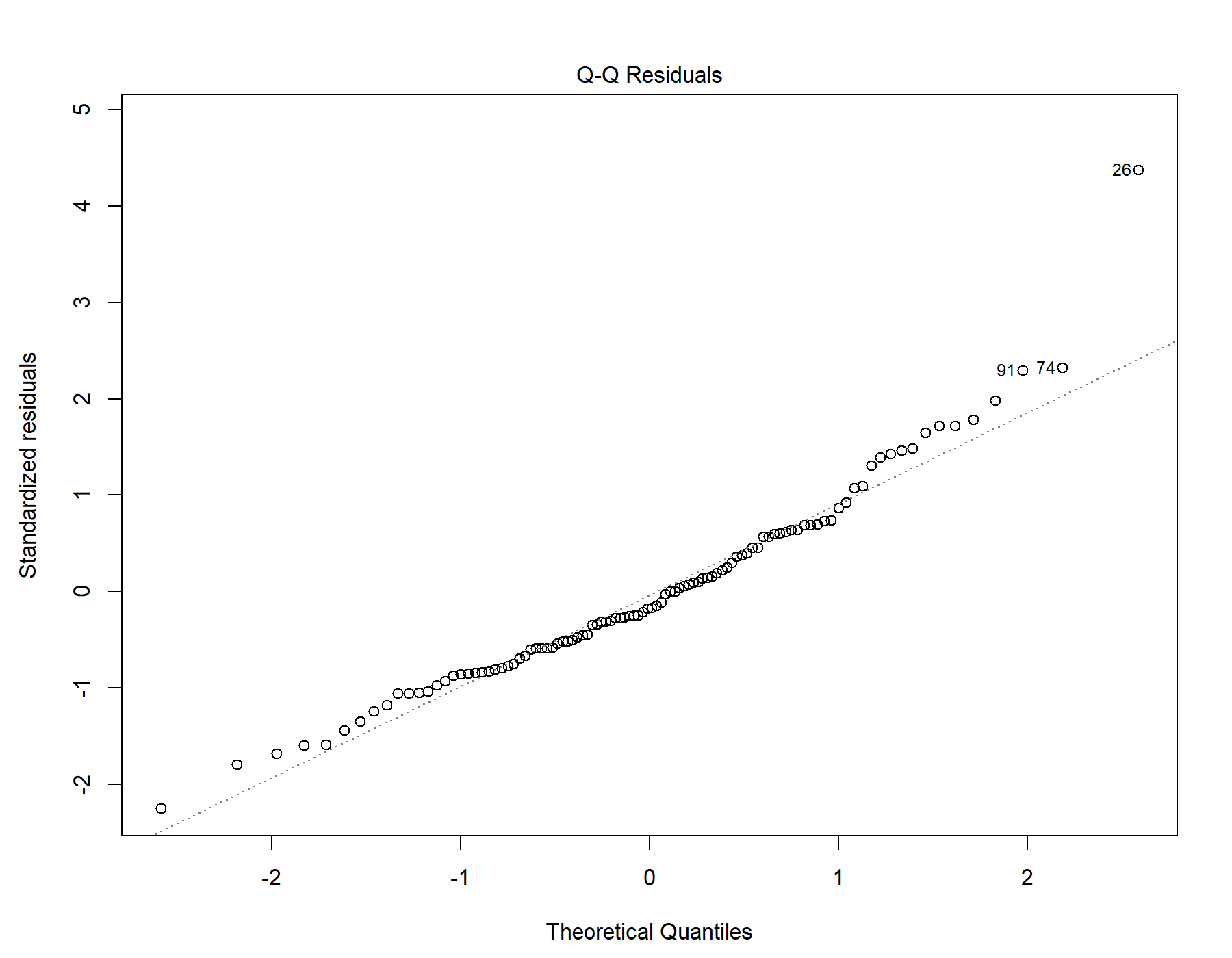

Supplemental Figure 85. Normal distribution quantile-quantile plot for T1w+T2w WML segmentation.

#### 41.5. Residuals vs leverage

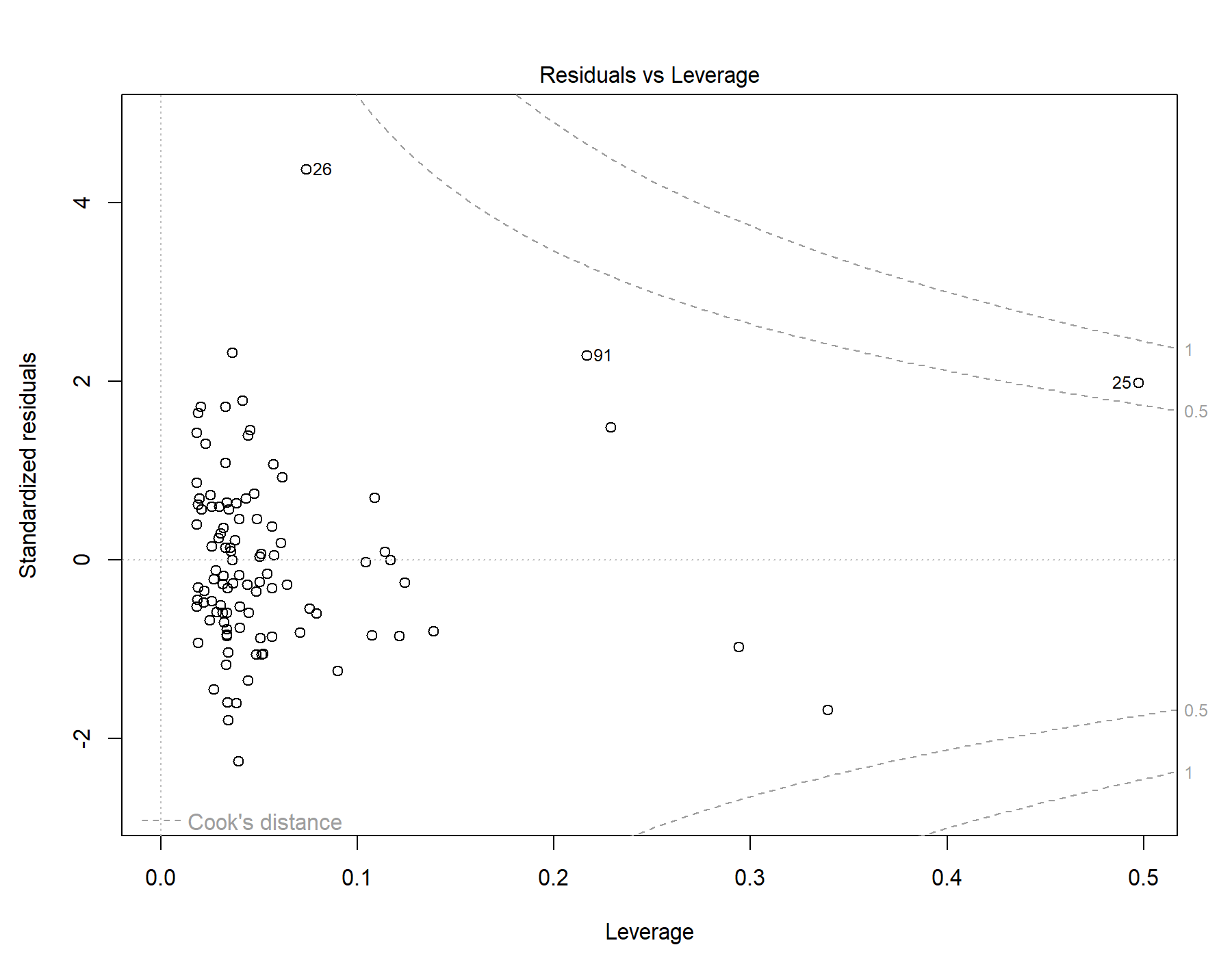

Supplemental Figure 86. Residuals vs leverage diagnostics for T1w+T2w WML segmentation.

### 42. Global | Young only | T1w-alone WML vol | SDNN.LF._log | Exploratory restricted interaction model

Model: T1w-alone WML vol ~ SDNN.LF._log + Age + Sex + Age * SDNN.LF._log + Sex * SDNN.LF._log

#### 42.1. Standardized coefficients

| **Term** | **Estimate** | **Std. Error** | **t** | **p** | **β** |
| --- | --- | --- | --- | --- | --- |
| (Intercept) | 5.135e-05 | 1.197e-04 | 0.429 | .669 |  |
| SDNN.LF._log | -3.463e-04 | 3.759e-04 | -0.921 | .359 | -0.373 |
| Age | 5.327e-05 | 4.506e-05 | 1.18 | .240 | 0.124 |
| Sex | -8.399e-05 | 6.159e-05 | -1.36 | .176 | -0.136 |
| SDNN.LF._log:Age | 6.673e-05 | 1.674e-04 | 0.399 | .691 | 0.112 |
| SDNN.LF._log:Sex | 1.803e-04 | 2.046e-04 | 0.881 | .380 | 0.339 |

***Supplemental Table 133.*** *Standardized coefficients values for T1w-alone WML segmentation.*

#### 42.2. 95% confidence intervals

| **Term** | **Lower 95%** | **Upper 95%** |
| --- | --- | --- |
| (Intercept) | -1.863e-04 | 2.890e-04 |
| SDNN.LF._log | -0.00109 | 3.996e-04 |
| Age | -3.614e-05 | 1.427e-04 |
| Sex | -2.062e-04 | 3.822e-05 |
| SDNN.LF._log:Age | -2.655e-04 | 3.989e-04 |
| SDNN.LF._log:Sex | -2.257e-04 | 5.864e-04 |

***Supplemental Table 134.*** *95% confidence intervals values for T1w-alone WML segmentation.*

#### 42.3. Breusch-Pagan test

| **Test** | **Statistic** | **df** | **p** |
| --- | --- | --- | --- |
| Breusch-Pagan | 6.536 | 5 | .257 |

***Supplemental Table 135.*** *Breusch-Pagan test values for T1w-alone WML segmentation.*

#### 42.4. Normal Q-Q plot

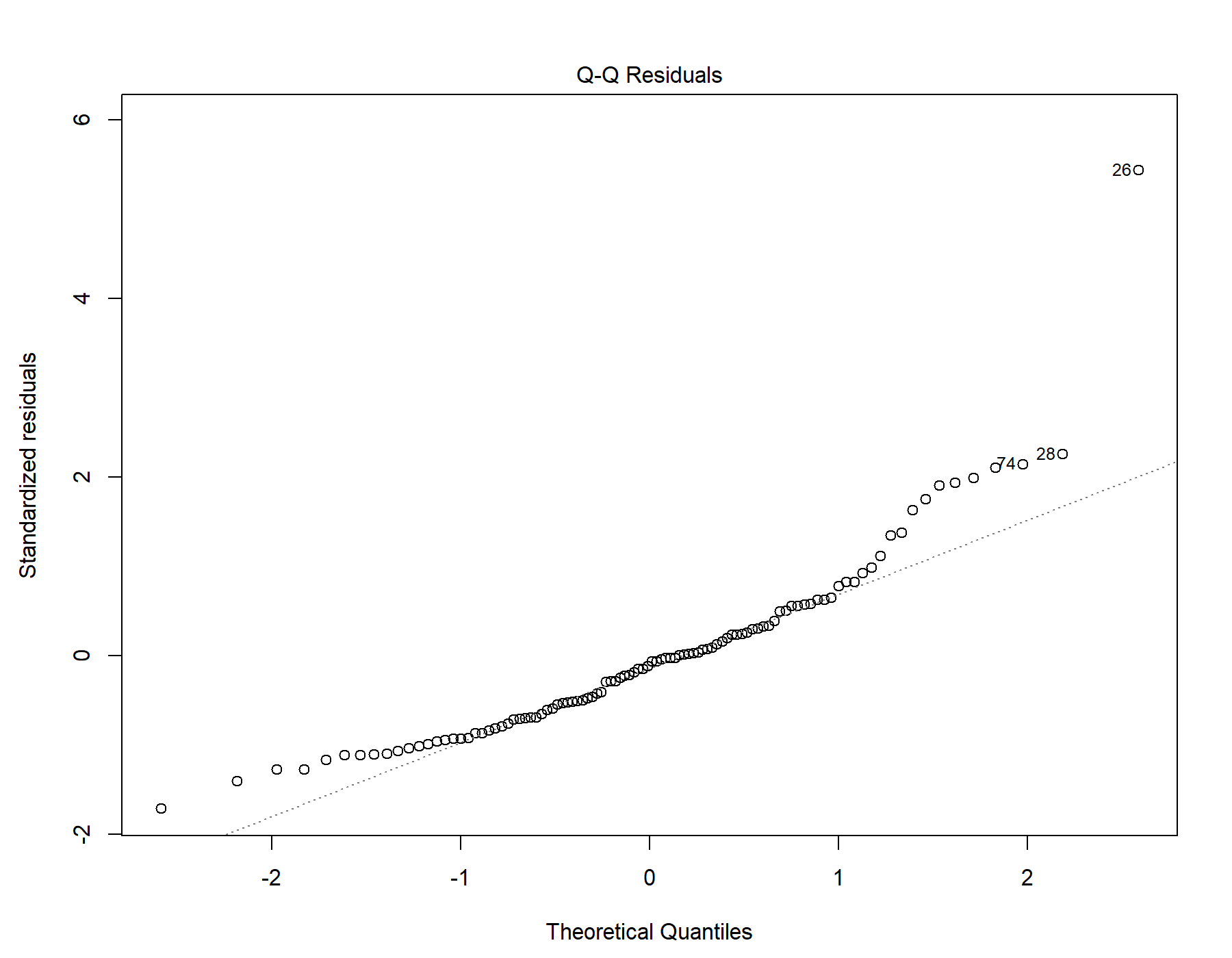

Supplemental Figure 87. Normal distribution quantile-quantile plot for T1w-alone WML segmentation.

#### 42.5. Residuals vs leverage

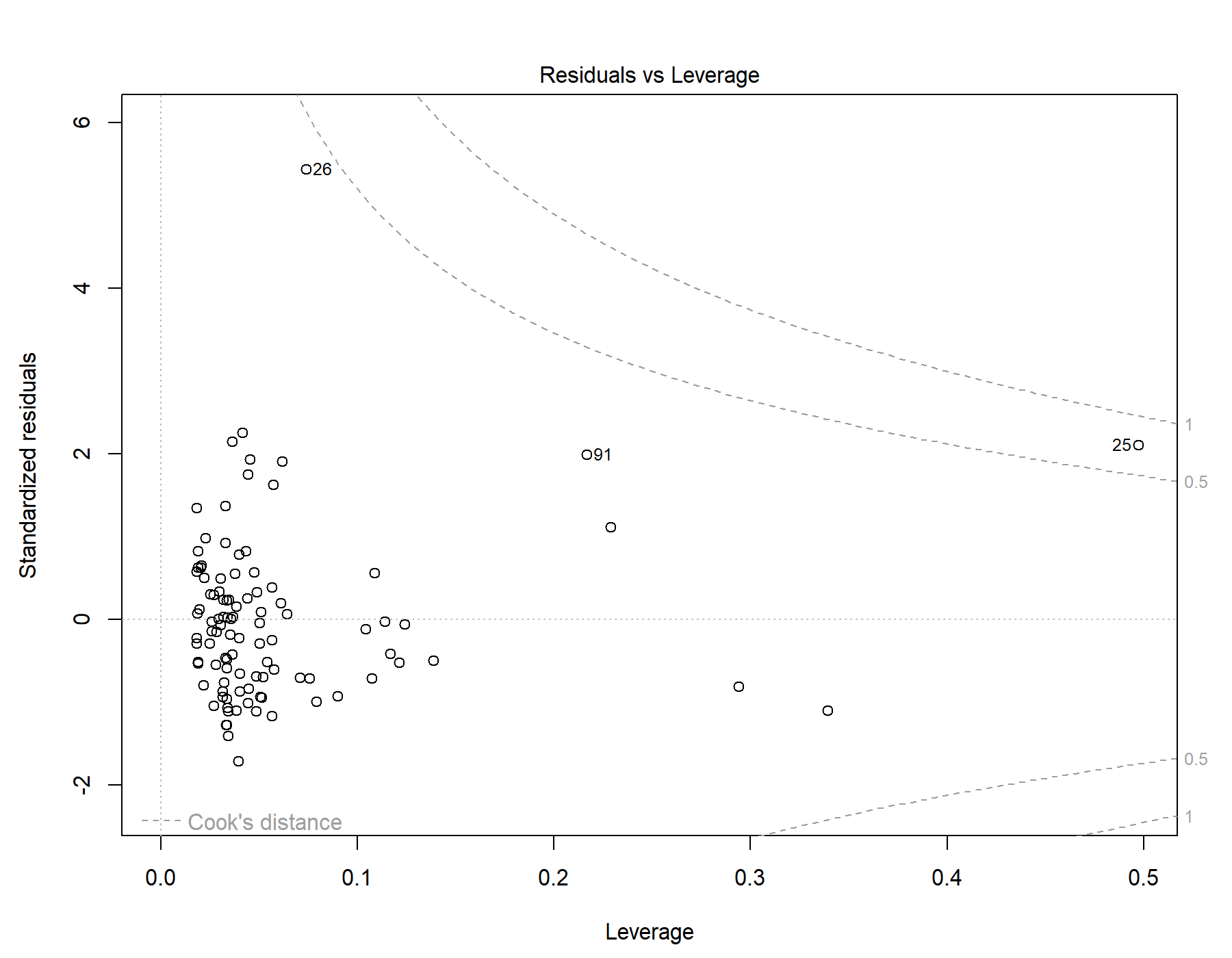

Supplemental Figure 88. Residuals vs leverage diagnostics for T1w-alone WML segmentation.

### 43. Global | Young only | T1w+FLAIR WML vol | SDNN.HF._log | Exploratory restricted interaction model

Model: T1w+FLAIR WML vol ~ SDNN.HF._log + Age + Sex + Age * SDNN.HF._log + Sex * SDNN.HF._log

#### 43.1. Standardized coefficients

| **Term** | **Estimate** | **Std. Error** | **t** | **p** | **β** |
| --- | --- | --- | --- | --- | --- |
| (Intercept) | -6.267e-05 | 1.113e-04 | -0.563 | .575 |  |
| SDNN.HF._log | -8.585e-05 | 3.447e-04 | -0.249 | .804 | -0.0917 |
| Age | 3.732e-05 | 4.332e-05 | 0.861 | .391 | 0.0888 |
| Sex | -1.903e-05 | 5.835e-05 | -0.326 | .745 | -0.0317 |
| SDNN.HF._log:Age | -4.789e-04 | 1.757e-04 | -2.73 | .008 | -0.756 |
| SDNN.HF._log:Sex | 3.963e-04 | 1.919e-04 | 2.06 | .042 | 0.715 |

***Supplemental Table 136.*** *Standardized coefficients values for T1w+FLAIR WML segmentation.*

#### 43.2. 95% confidence intervals

| **Term** | **Lower 95%** | **Upper 95%** |
| --- | --- | --- |
| (Intercept) | -2.836e-04 | 1.583e-04 |
| SDNN.HF._log | -7.699e-04 | 5.982e-04 |
| Age | -4.866e-05 | 1.233e-04 |
| Sex | -1.348e-04 | 9.676e-05 |
| SDNN.HF._log:Age | -8.276e-04 | -1.303e-04 |
| SDNN.HF._log:Sex | 1.541e-05 | 7.772e-04 |

***Supplemental Table 137.*** *95% confidence intervals values for T1w+FLAIR WML segmentation.*

#### 43.3. Breusch-Pagan test

| **Test** | **Statistic** | **df** | **p** |
| --- | --- | --- | --- |
| Breusch-Pagan | 9.404 | 5 | .094 |

***Supplemental Table 138.*** *Breusch-Pagan test values for T1w+FLAIR WML segmentation.*

#### 43.4. Normal Q-Q plot

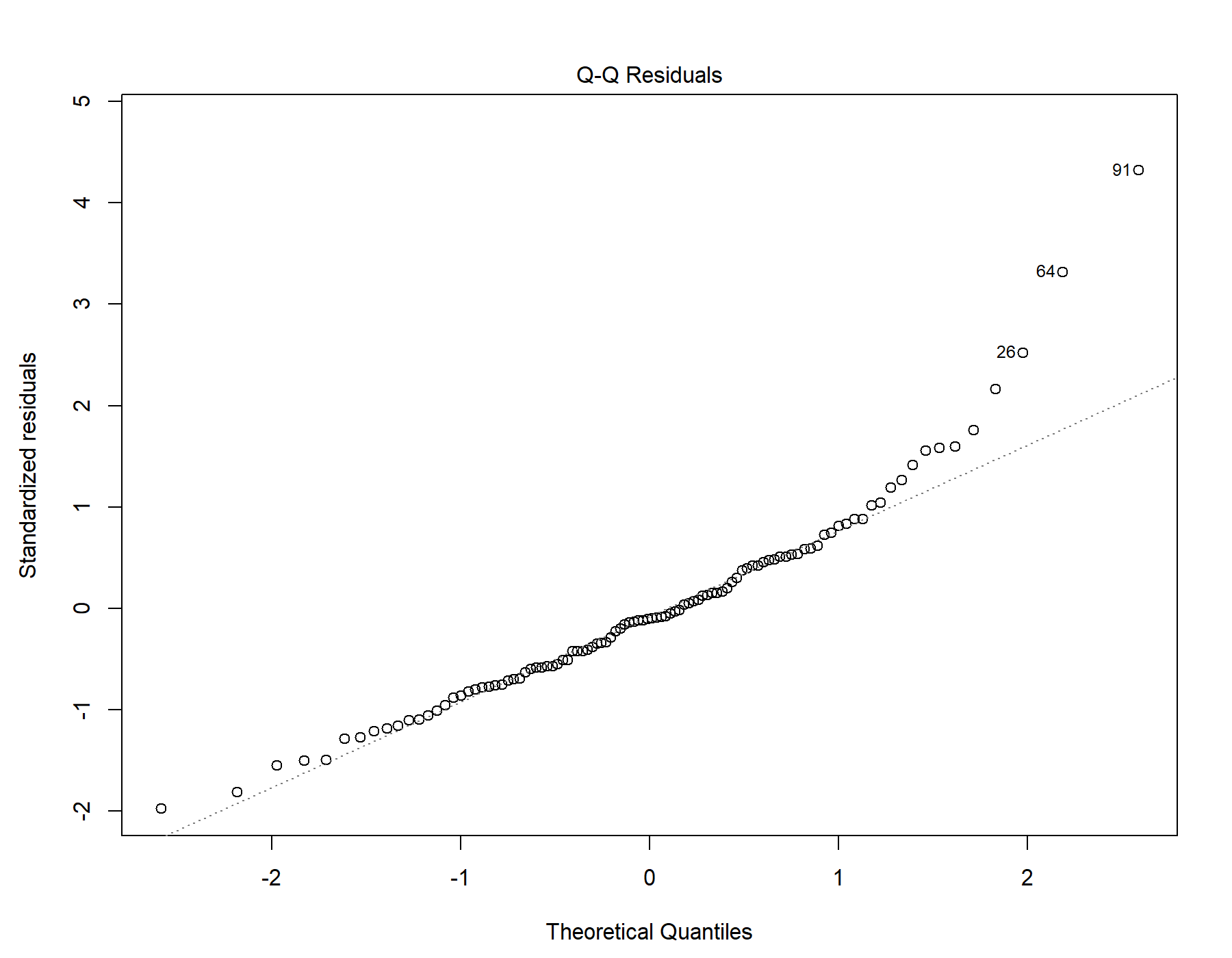

Supplemental Figure 89. Normal distribution quantile-quantile plot for T1w+FLAIR WML segmentation.

#### 43.5. Residuals vs leverage

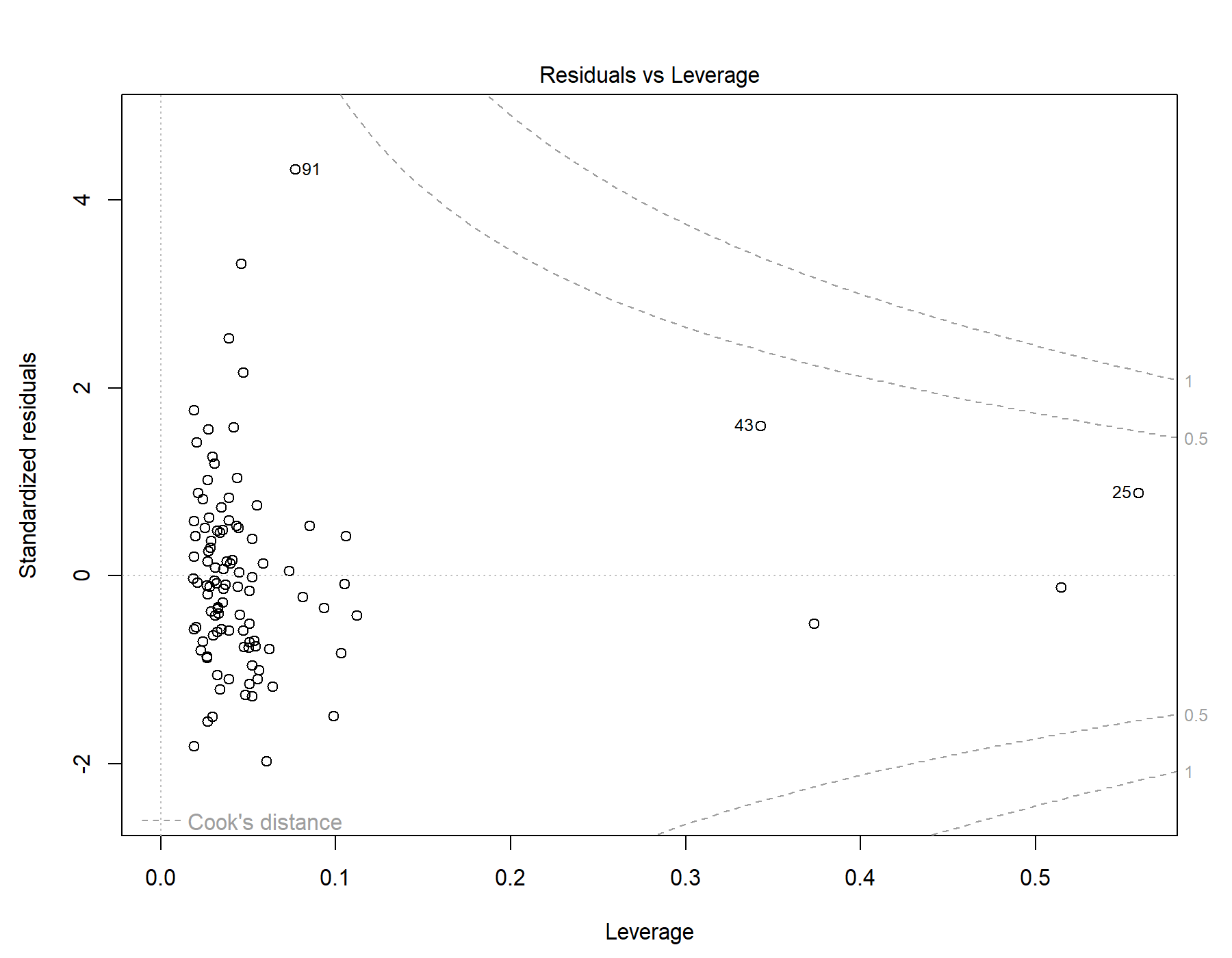

Supplemental Figure 90. Residuals vs leverage diagnostics for T1w+FLAIR WML segmentation.

### 44. Global | Young only | T1w+T2w WML vol | SDNN.HF._log | Exploratory restricted interaction model

Model: T1w+T2w WML vol ~ SDNN.HF._log + Age + Sex + Age * SDNN.HF._log + Sex * SDNN.HF._log

#### 44.1. Standardized coefficients

| **Term** | **Estimate** | **Std. Error** | **t** | **p** | **β** |
| --- | --- | --- | --- | --- | --- |
| (Intercept) | 4.379e-05 | 1.354e-04 | 0.323 | .747 |  |
| SDNN.HF._log | 3.511e-04 | 4.193e-04 | 0.837 | .404 | 0.318 |
| Age | 4.902e-05 | 5.270e-05 | 0.930 | .355 | 0.0990 |
| Sex | -8.935e-05 | 7.097e-05 | -1.26 | .211 | -0.126 |
| SDNN.HF._log:Age | -3.835e-04 | 2.137e-04 | -1.79 | .076 | -0.514 |
| SDNN.HF._log:Sex | 8.900e-05 | 2.335e-04 | 0.381 | .704 | 0.136 |

***Supplemental Table 139.*** *Standardized coefficients values for T1w+T2w WML segmentation.*

#### 44.2. 95% confidence intervals

| **Term** | **Lower 95%** | **Upper 95%** |
| --- | --- | --- |
| (Intercept) | -2.249e-04 | 3.125e-04 |
| SDNN.HF._log | -4.809e-04 | 0.00118 |
| Age | -5.556e-05 | 1.536e-04 |
| Sex | -2.302e-04 | 5.150e-05 |
| SDNN.HF._log:Age | -8.076e-04 | 4.057e-05 |
| SDNN.HF._log:Sex | -3.743e-04 | 5.523e-04 |

***Supplemental Table 140.*** *95% confidence intervals values for T1w+T2w WML segmentation.*

#### 44.3. Breusch-Pagan test

| **Test** | **Statistic** | **df** | **p** |
| --- | --- | --- | --- |
| Breusch-Pagan | 4.958 | 5 | .421 |

***Supplemental Table 141.*** *Breusch-Pagan test values for T1w+T2w WML segmentation.*

#### 44.4. Normal Q-Q plot

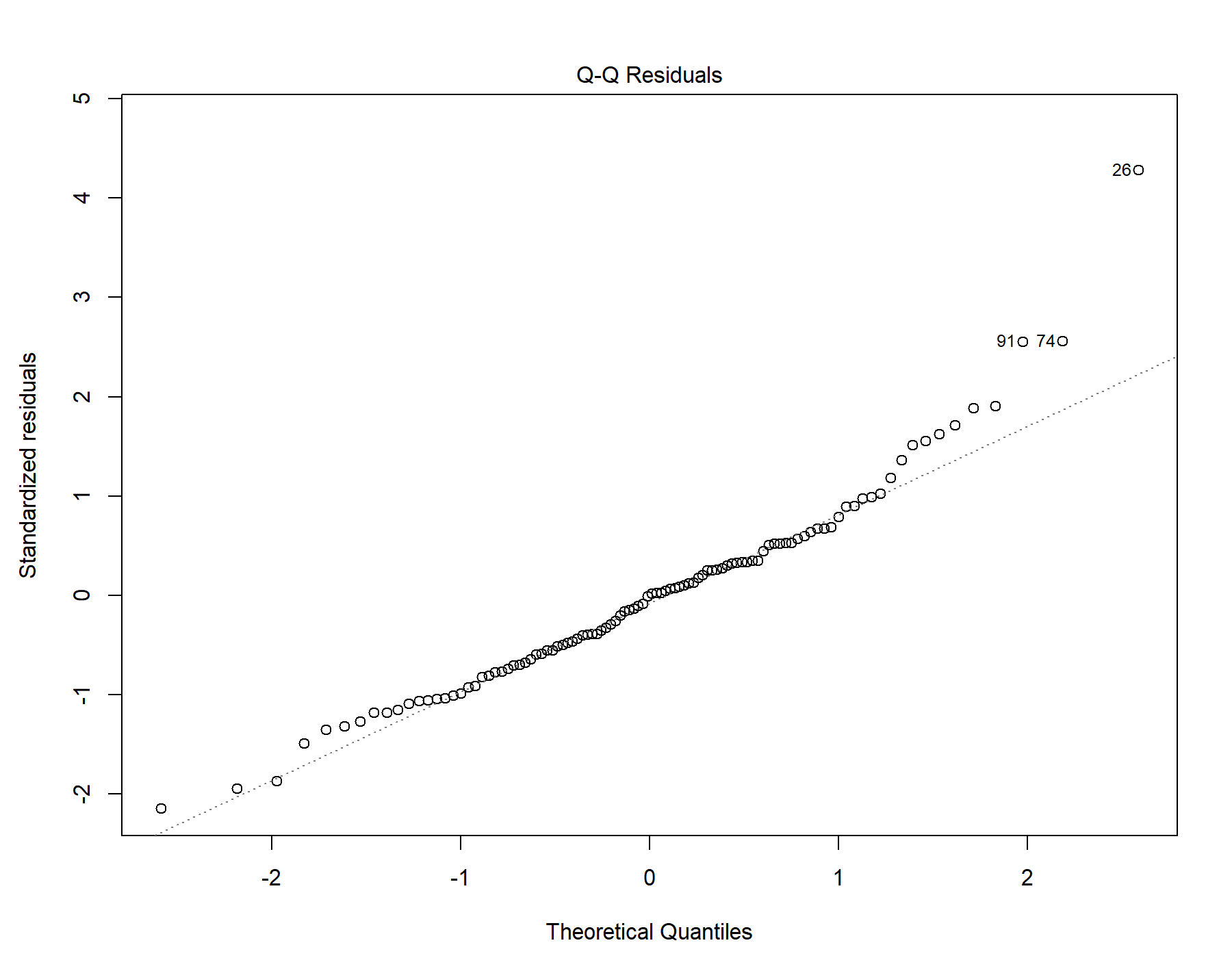

Supplemental Figure 91. Normal distribution quantile-quantile plot for T1w+T2w WML segmentation.

#### 44.5. Residuals vs leverage

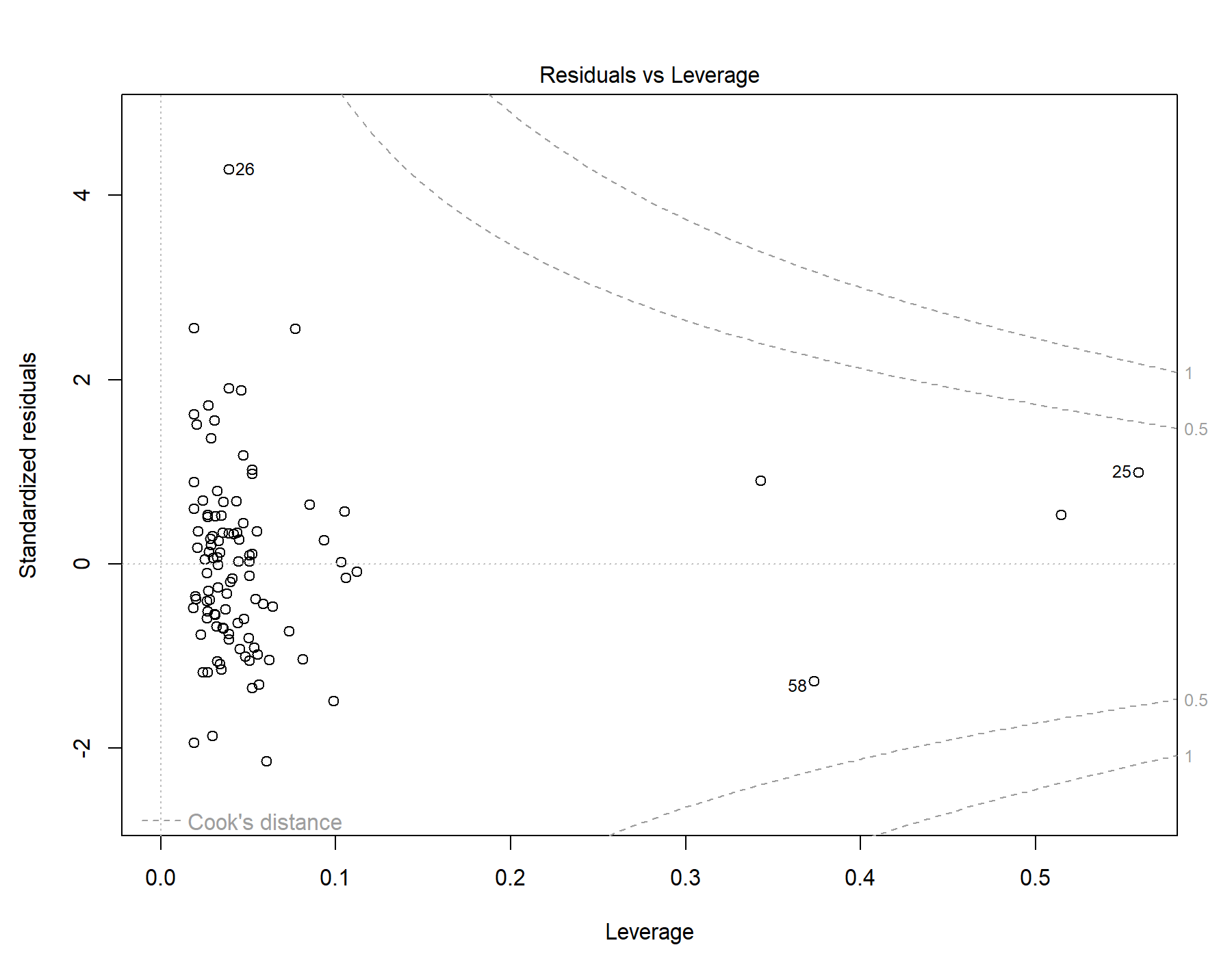

Supplemental Figure 92. Residuals vs leverage diagnostics for T1w+T2w WML segmentation.

### 45. Global | Young only | T1w-alone WML vol | SDNN.HF._log | Exploratory restricted interaction model

Model: T1w-alone WML vol ~ SDNN.HF._log + Age + Sex + Age * SDNN.HF._log + Sex * SDNN.HF._log

#### 45.1. Standardized coefficients

| **Term** | **Estimate** | **Std. Error** | **t** | **p** | **β** |
| --- | --- | --- | --- | --- | --- |
| (Intercept) | 6.658e-05 | 1.201e-04 | 0.554 | .581 |  |
| SDNN.HF._log | 1.416e-04 | 3.720e-04 | 0.381 | .704 | 0.147 |
| Age | 2.732e-05 | 4.675e-05 | 0.584 | .560 | 0.0634 |
| Sex | -7.570e-05 | 6.297e-05 | -1.20 | .232 | -0.123 |
| SDNN.HF._log:Age | -2.025e-04 | 1.896e-04 | -1.07 | .288 | -0.312 |
| SDNN.HF._log:Sex | 7.107e-05 | 2.071e-04 | 0.343 | .732 | 0.125 |

***Supplemental Table 142.*** *Standardized coefficients values for T1w-alone WML segmentation.*

#### 45.2. 95% confidence intervals

| **Term** | **Lower 95%** | **Upper 95%** |
| --- | --- | --- |
| (Intercept) | -1.718e-04 | 3.050e-04 |
| SDNN.HF._log | -5.965e-04 | 8.798e-04 |
| Age | -6.546e-05 | 1.201e-04 |
| Sex | -2.007e-04 | 4.925e-05 |
| SDNN.HF._log:Age | -5.788e-04 | 1.737e-04 |
| SDNN.HF._log:Sex | -3.400e-04 | 4.821e-04 |

***Supplemental Table 143.*** *95% confidence intervals values for T1w-alone WML segmentation.*

#### 45.3. Breusch-Pagan test

| **Test** | **Statistic** | **df** | **p** |
| --- | --- | --- | --- |
| Breusch-Pagan | 4.412 | 5 | .492 |

***Supplemental Table 144.*** *Breusch-Pagan test values for T1w-alone WML segmentation.*

#### 45.4. Normal Q-Q plot

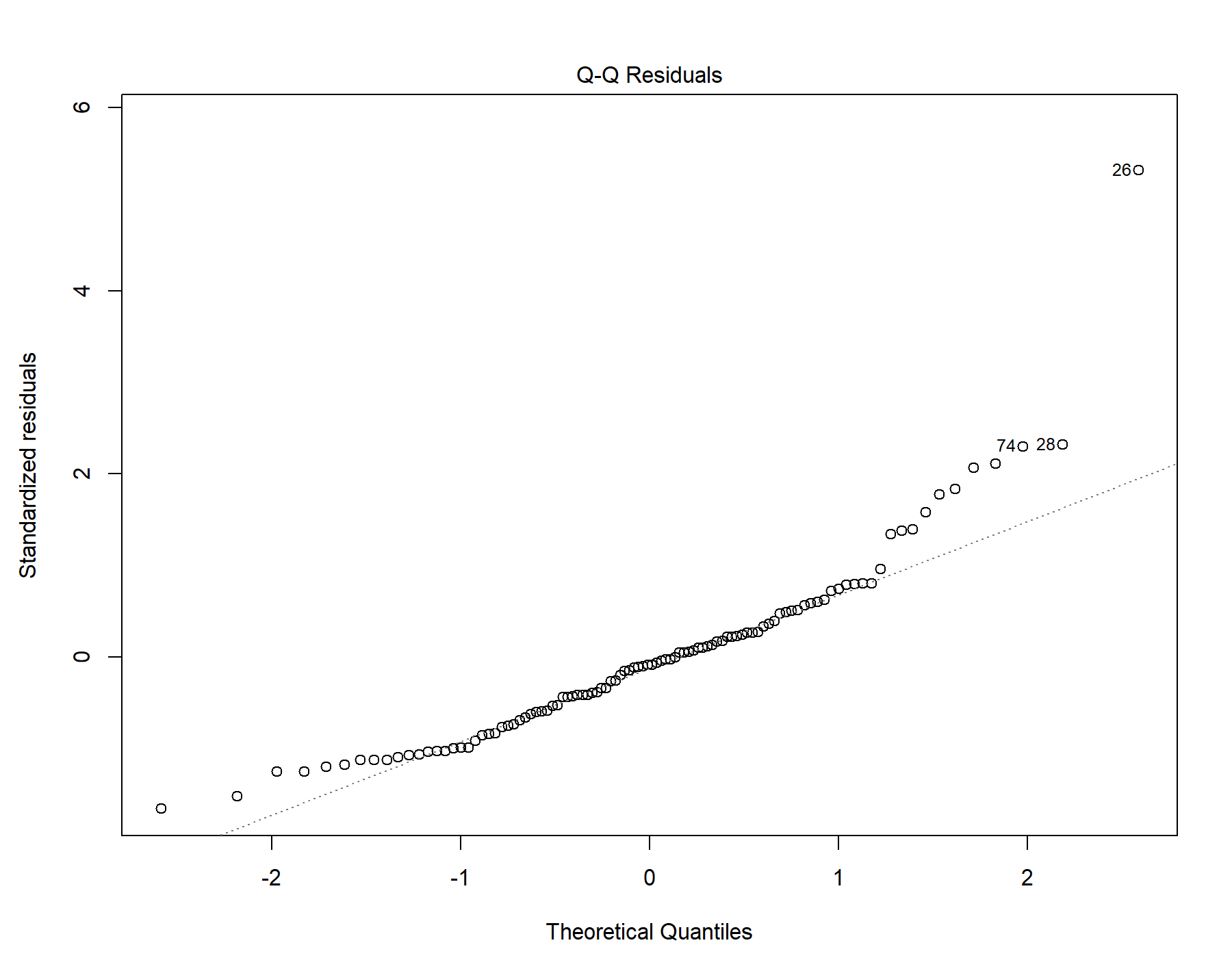

Supplemental Figure 93. Normal distribution quantile-quantile plot for T1w-alone WML segmentation.

#### 45.5. Residuals vs leverage

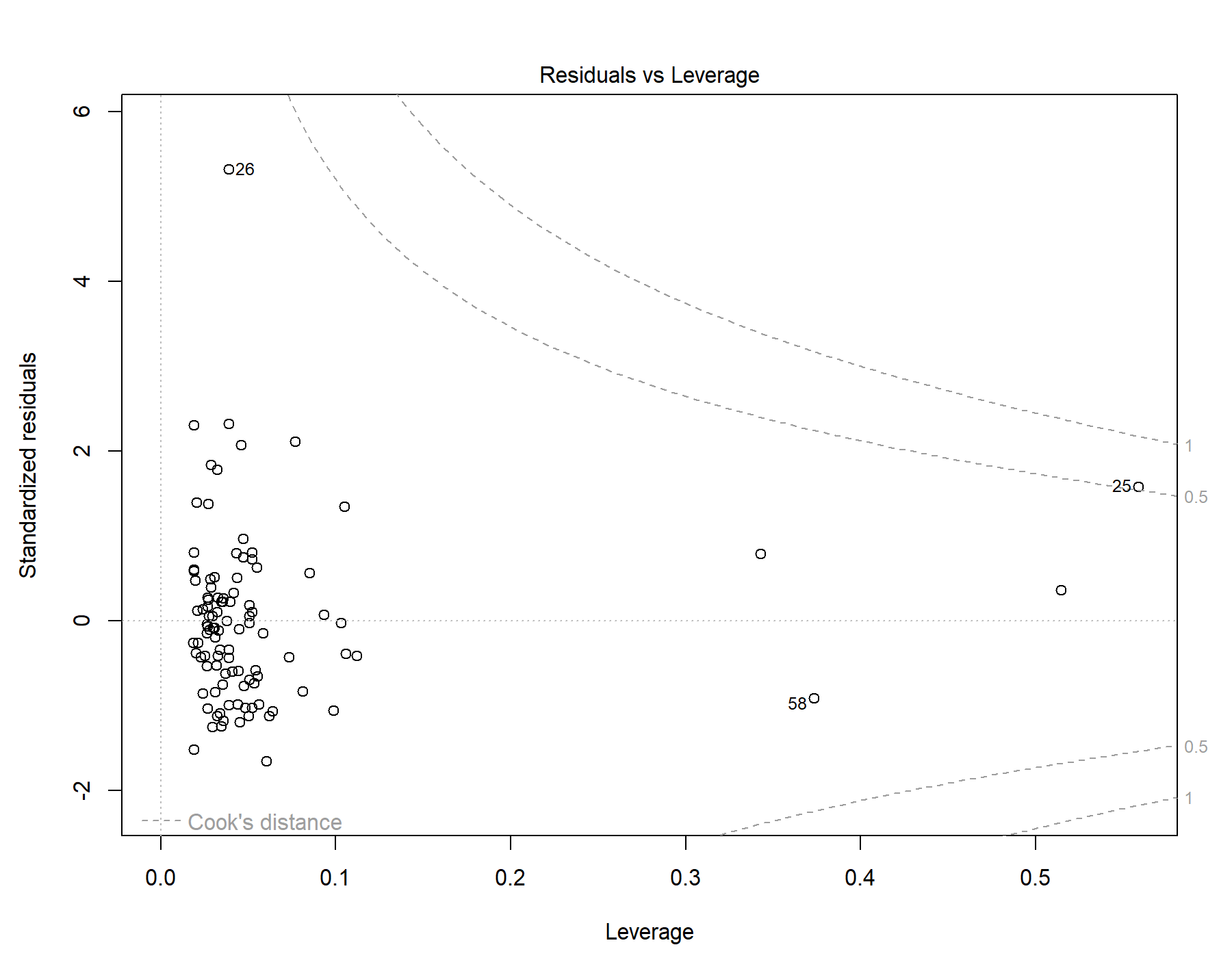

Supplemental Figure 94. Residuals vs leverage diagnostics for T1w-alone WML segmentation.

### 46. Global | Old only | T1w+FLAIR WML vol | RMSSD | Exploratory restricted interaction model

Model: T1w+FLAIR WML vol ~ RMSSD + Age + Sex +	Age * RMSSD + Sex * RMSSD

#### 46.1. Standardized coefficients

| **Term** | **Estimate** | **Std. Error** | **t** | **p** | **β** |
| --- | --- | --- | --- | --- | --- |
| (Intercept) | -3.010e-04 | 0.00390 | -0.0772 | .939 |  |
| RMSSD | 4.542e-04 | 2.705e-04 | 1.68 | .106 | 20.1 |
| Age | 6.429e-04 | 2.738e-04 | 2.35 | .027 | 0.489 |
| Sex | -0.00302 | 0.00208 | -1.45 | .160 | -0.630 |
| RMSSD:Age | 1.041e-06 | 9.953e-06 | 0.105 | .918 | 0.468 |
| RMSSD:Sex | -2.356e-04 | 1.239e-04 | -1.90 | .069 | -20.8 |

***Supplemental Table 145.*** *Standardized coefficients values for T1w+FLAIR WML segmentation.*

#### 46.2. 95% confidence intervals

| **Term** | **Lower 95%** | **Upper 95%** |
| --- | --- | --- |
| (Intercept) | -0.00835 | 0.00775 |
| RMSSD | -1.040e-04 | 0.00101 |
| Age | 7.793e-05 | 0.00121 |
| Sex | -0.00731 | 0.00127 |
| RMSSD:Age | -1.950e-05 | 2.158e-05 |
| RMSSD:Sex | -4.912e-04 | 1.999e-05 |

***Supplemental Table 146.*** *95% confidence intervals values for T1w+FLAIR WML segmentation.*

#### 46.3. Breusch-Pagan test

| **Test** | **Statistic** | **df** | **p** |
| --- | --- | --- | --- |
| Breusch-Pagan | 5.049 | 5 | .410 |

***Supplemental Table 147.*** *Breusch-Pagan test values for T1w+FLAIR WML segmentation.*

#### 46.4. Normal Q-Q plot

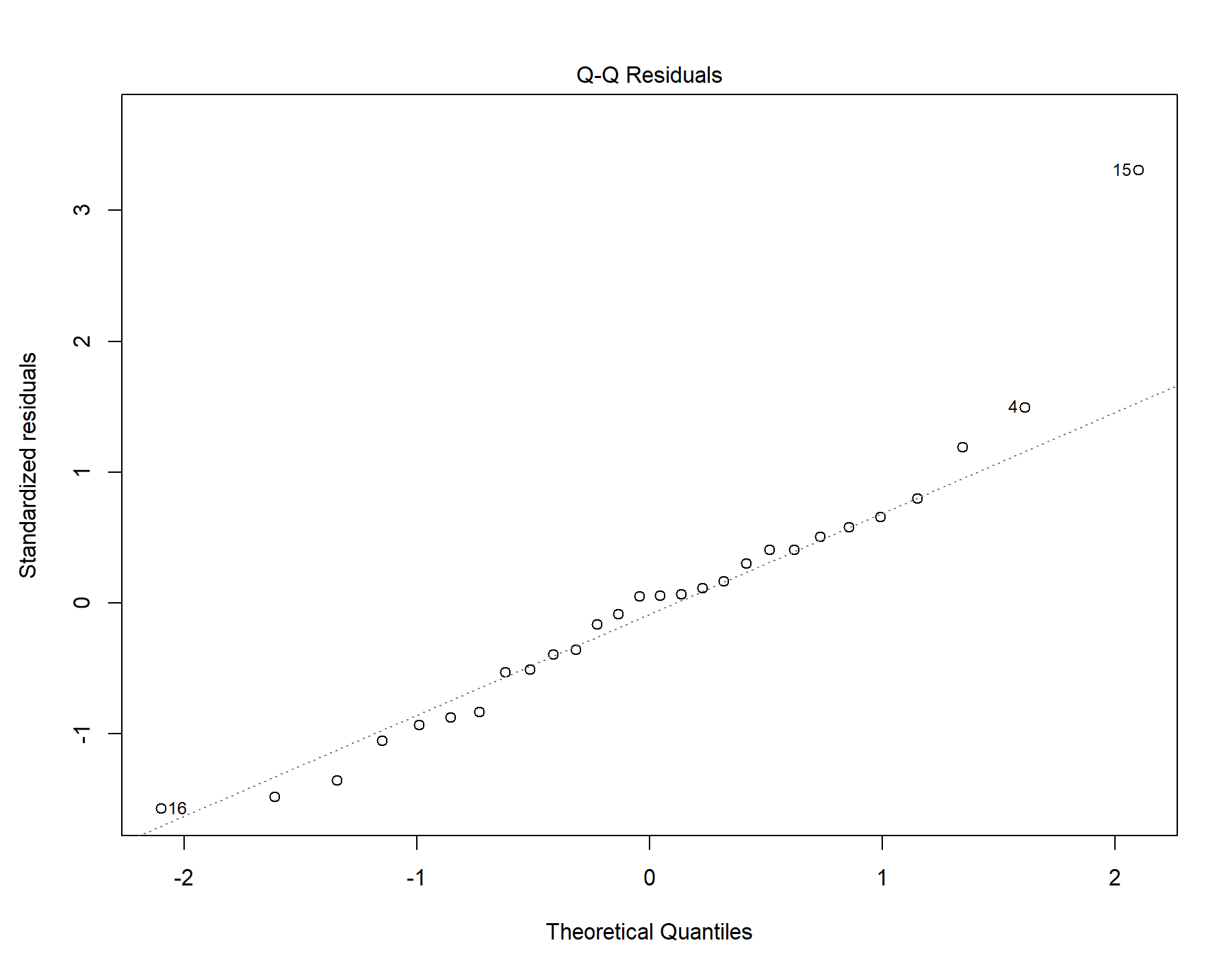

Supplemental Figure 95. Normal distribution quantile-quantile plot for T1w+FLAIR WML segmentation.

#### 46.5. Residuals vs leverage

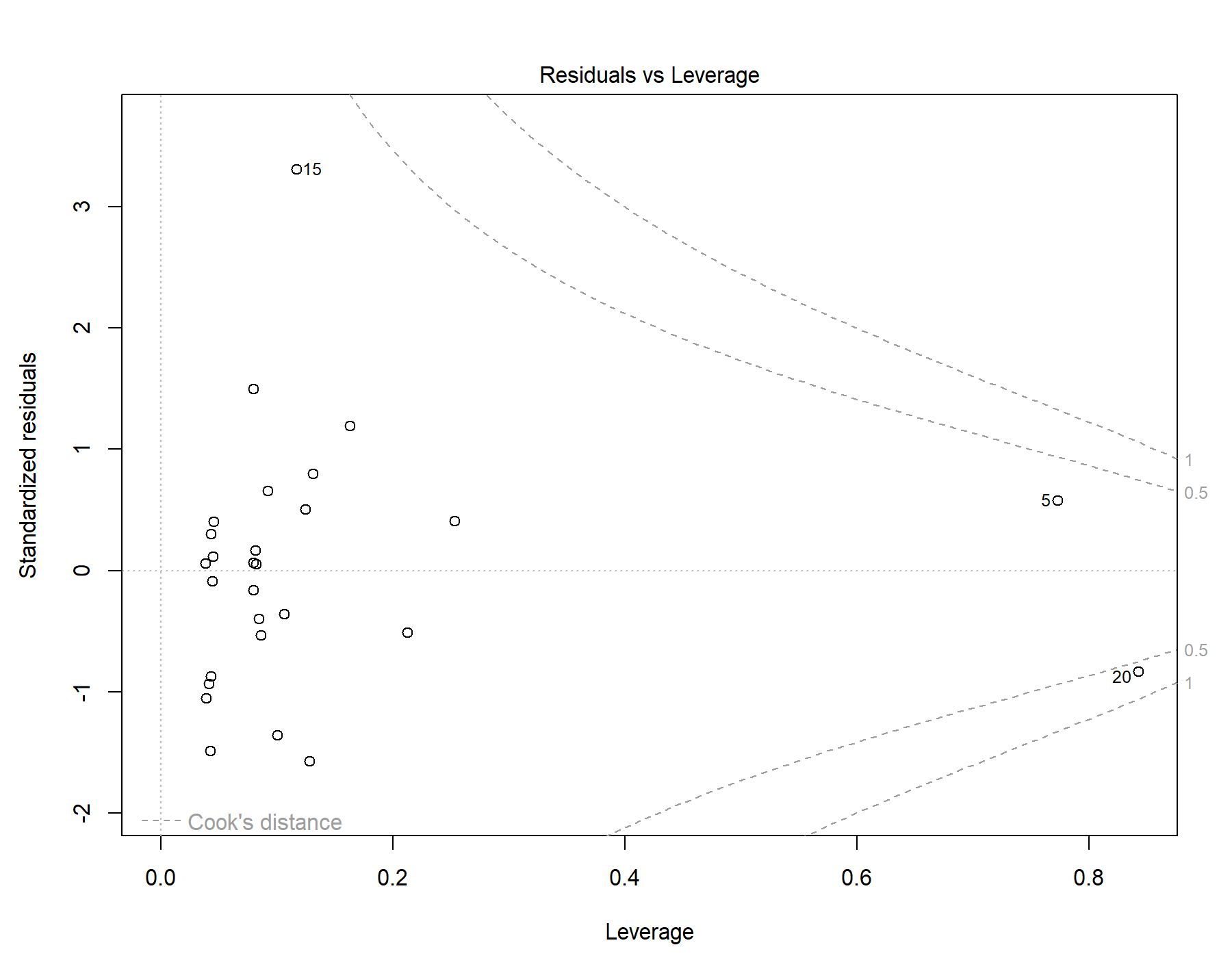

Supplemental Figure 96. Residuals vs leverage diagnostics for T1w+FLAIR WML segmentation. The 2 older female subjects are not visualized due to possessing leverage values approaching 1 in this restricted model, preventing the normal calculation of their standardized residuals.

### 47. Global | Old only | T1w+T2w WML vol | RMSSD | Exploratory restricted interaction model

Model: T1w+T2w WML vol ~ RMSSD + Age + Sex + Age *	RMSSD + Sex * RMSSD

#### 47.1. Standardized coefficients

| **Term** | **Estimate** | **Std. Error** | **t** | **p** | **β** |
| --- | --- | --- | --- | --- | --- |
| (Intercept) | 0.00119 | 0.00415 | 0.287 | .777 |  |
| RMSSD | 5.991e-04 | 2.879e-04 | 2.08 | .048 | 25. |
| Age | 6.538e-04 | 2.914e-04 | 2.24 | .034 | 0.467 |
| Sex | -0.00381 | 0.00221 | -1.72 | .098 | -0.747 |
| RMSSD:Age | -2.389e-06 | 1.060e-05 | -0.225 | .824 | -1.01 |
| RMSSD:Sex | -2.903e-04 | 1.319e-04 | -2.20 | .038 | -24.1 |

***Supplemental Table 148.*** *Standardized coefficients values for T1w+T2w WML segmentation.*

#### 47.2. 95% confidence intervals

| **Term** | **Lower 95%** | **Upper 95%** |
| --- | --- | --- |
| (Intercept) | -0.00738 | 0.00976 |
| RMSSD | 4.841e-06 | 0.00119 |
| Age | 5.232e-05 | 0.00126 |
| Sex | -0.00837 | 7.599e-04 |
| RMSSD:Age | -2.426e-05 | 1.948e-05 |
| RMSSD:Sex | -5.624e-04 | -1.815e-05 |

***Supplemental Table 149.*** *95% confidence intervals values for T1w+T2w WML segmentation.*

#### 47.3. Breusch-Pagan test

| **Test** | **Statistic** | **df** | **p** |
| --- | --- | --- | --- |
| Breusch-Pagan | 3.893 | 5 | .565 |

***Supplemental Table 150.*** *Breusch-Pagan test values for T1w+T2w WML segmentation.*

#### 47.4. Normal Q-Q plot

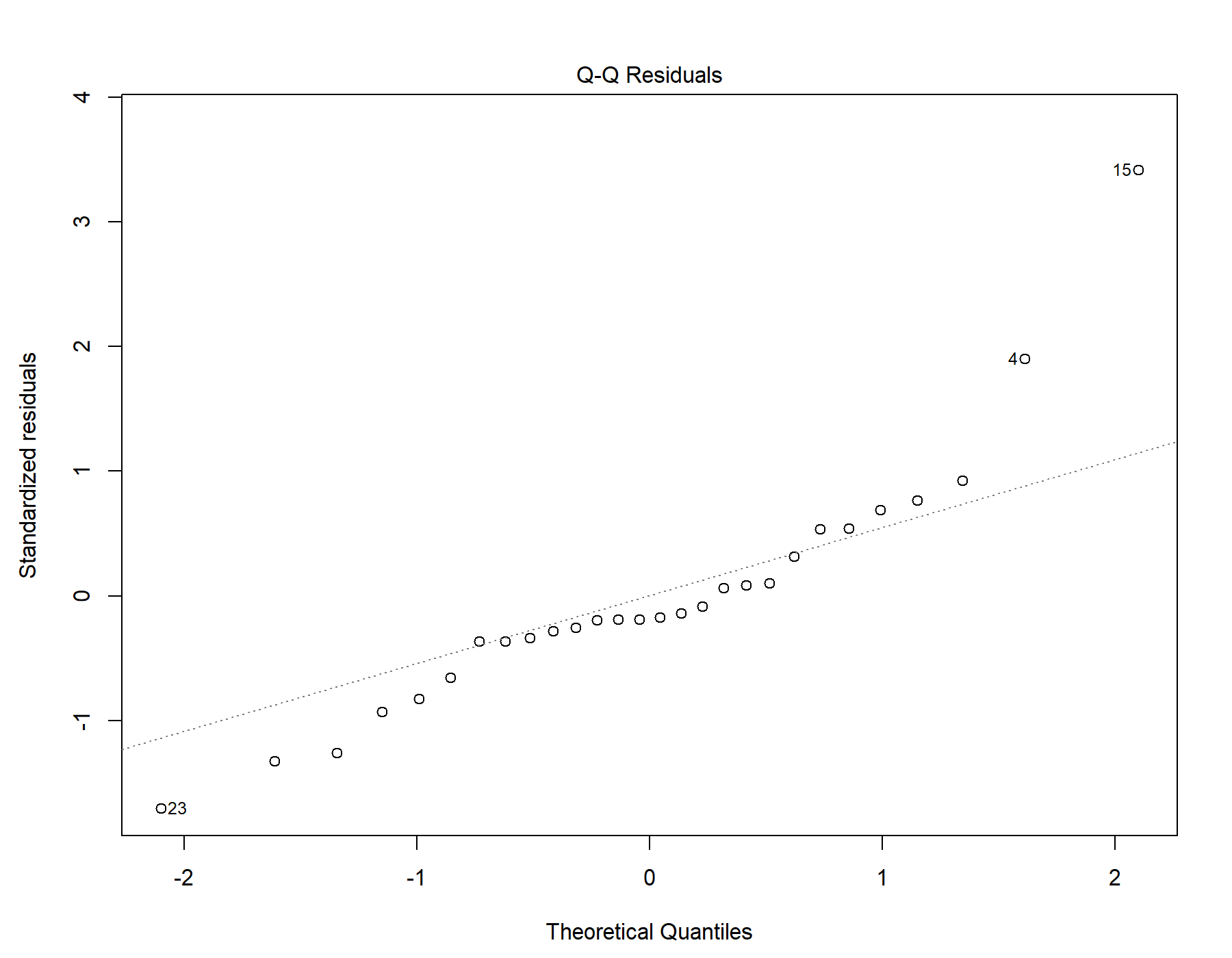

Supplemental Figure 97. Normal distribution quantile-quantile plot for T1w+T2w WML segmentation.

#### 47.5. Residuals vs leverage

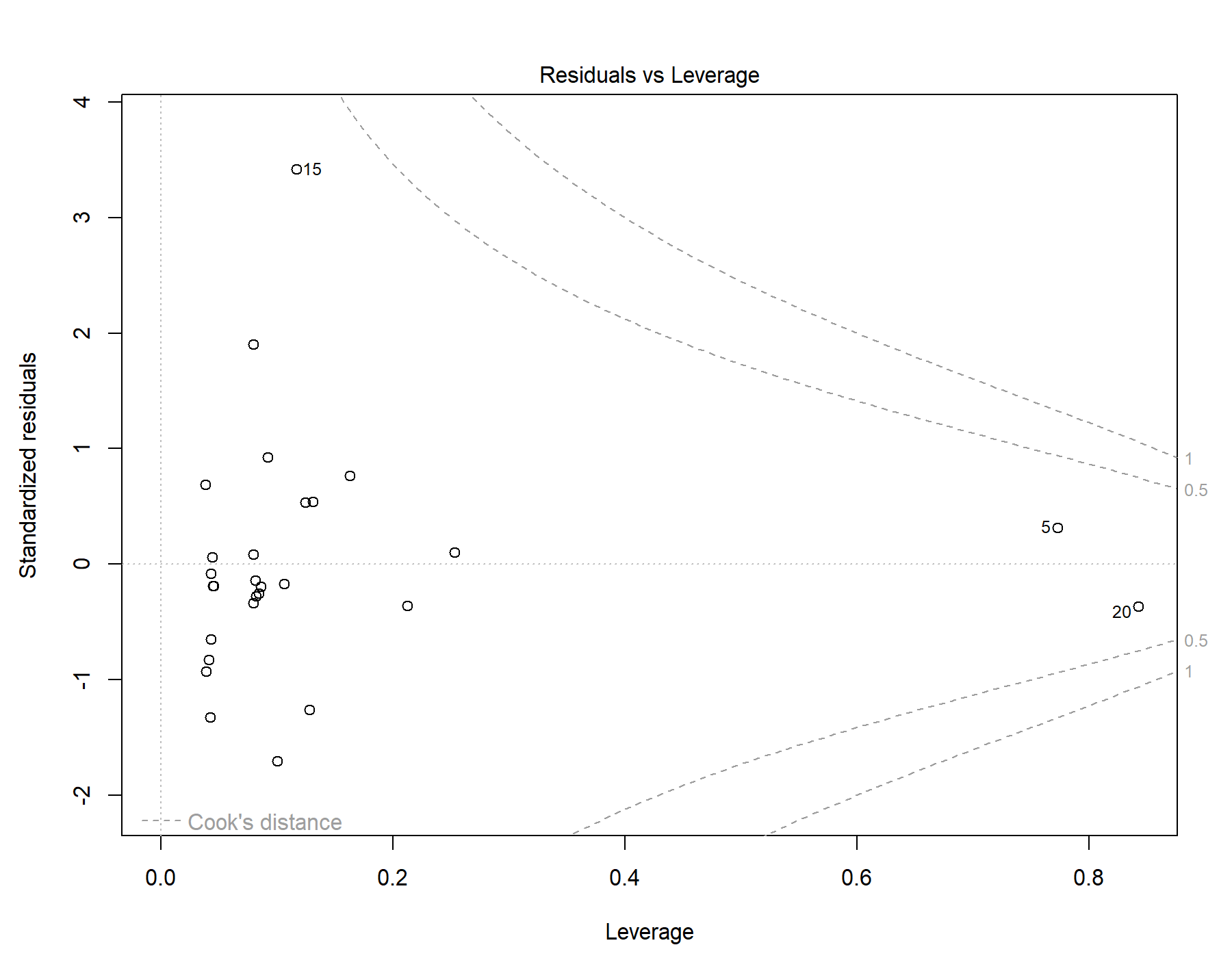

Supplemental Figure 98. Residuals vs leverage diagnostics for T1w+T2w WML segmentation. The 2 older female subjects are not visualized due to possessing leverage values approaching 1 in this restricted model, preventing the normal calculation of their standardized residuals.

### 48. Global | Old only | T1w-alone WML vol | RMSSD | Exploratory restricted interaction model

Model: T1w-alone WML vol ~ RMSSD + Age + Sex + Age *	RMSSD + Sex * RMSSD

#### 48.1. Standardized coefficients

| **Term** | **Estimate** | **Std. Error** | **t** | **p** | **β** |
| --- | --- | --- | --- | --- | --- |
| (Intercept) | 0.00131 | 0.00356 | 0.368 | .716 |  |
| RMSSD | 4.175e-04 | 2.467e-04 | 1.69 | .104 | 21.1 |
| Age | 4.931e-04 | 2.497e-04 | 1.97 | .060 | 0.428 |
| Sex | -0.00309 | 0.00190 | -1.63 | .116 | -0.738 |
| RMSSD:Age | 2.603e-06 | 9.078e-06 | 0.287 | .777 | 1.34 |
| RMSSD:Sex | -2.241e-04 | 1.130e-04 | -1.98 | .059 | -22.6 |

***Supplemental Table 151.*** *Standardized coefficients values for T1w-alone WML segmentation.*

#### 48.2. 95% confidence intervals

| **Term** | **Lower 95%** | **Upper 95%** |
| --- | --- | --- |
| (Intercept) | -0.00603 | 0.00865 |
| RMSSD | -9.164e-05 | 9.267e-04 |
| Age | -2.229e-05 | 0.00101 |
| Sex | -0.00701 | 8.214e-04 |
| RMSSD:Age | -1.613e-05 | 2.134e-05 |
| RMSSD:Sex | -4.573e-04 | 9.009e-06 |

***Supplemental Table 152.*** *95% confidence intervals values for T1w-alone WML segmentation.*

#### 48.3. Breusch-Pagan test

| **Test** | **Statistic** | **df** | **p** |
| --- | --- | --- | --- |
| Breusch-Pagan | 3.349 | 5 | .646 |

***Supplemental Table 153.*** *Breusch-Pagan test values for T1w-alone WML segmentation.*

#### 48.4. Normal Q-Q plot

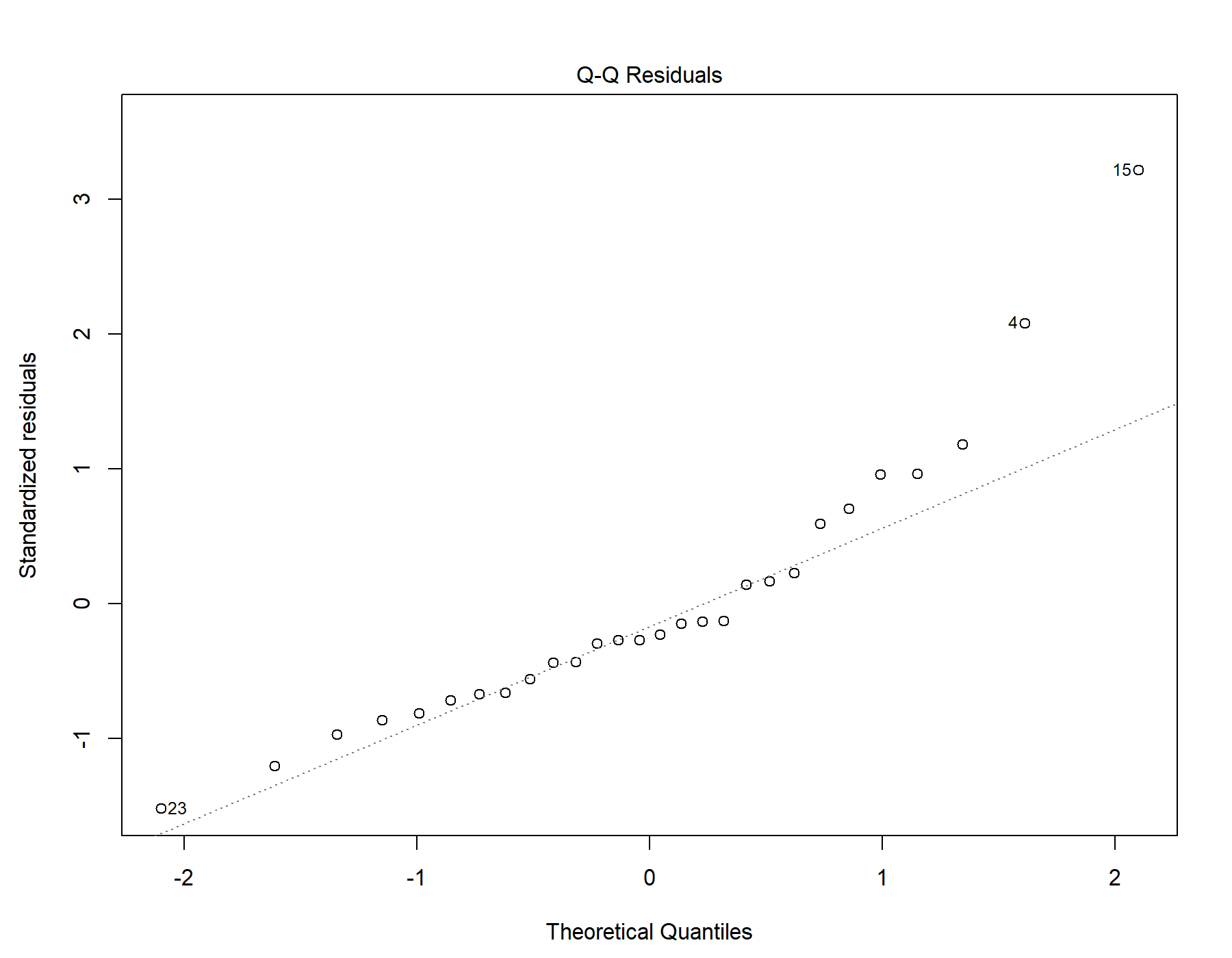

Supplemental Figure 99. Normal distribution quantile-quantile plot for T1w-alone WML segmentation.

#### 48.5. Residuals vs leverage

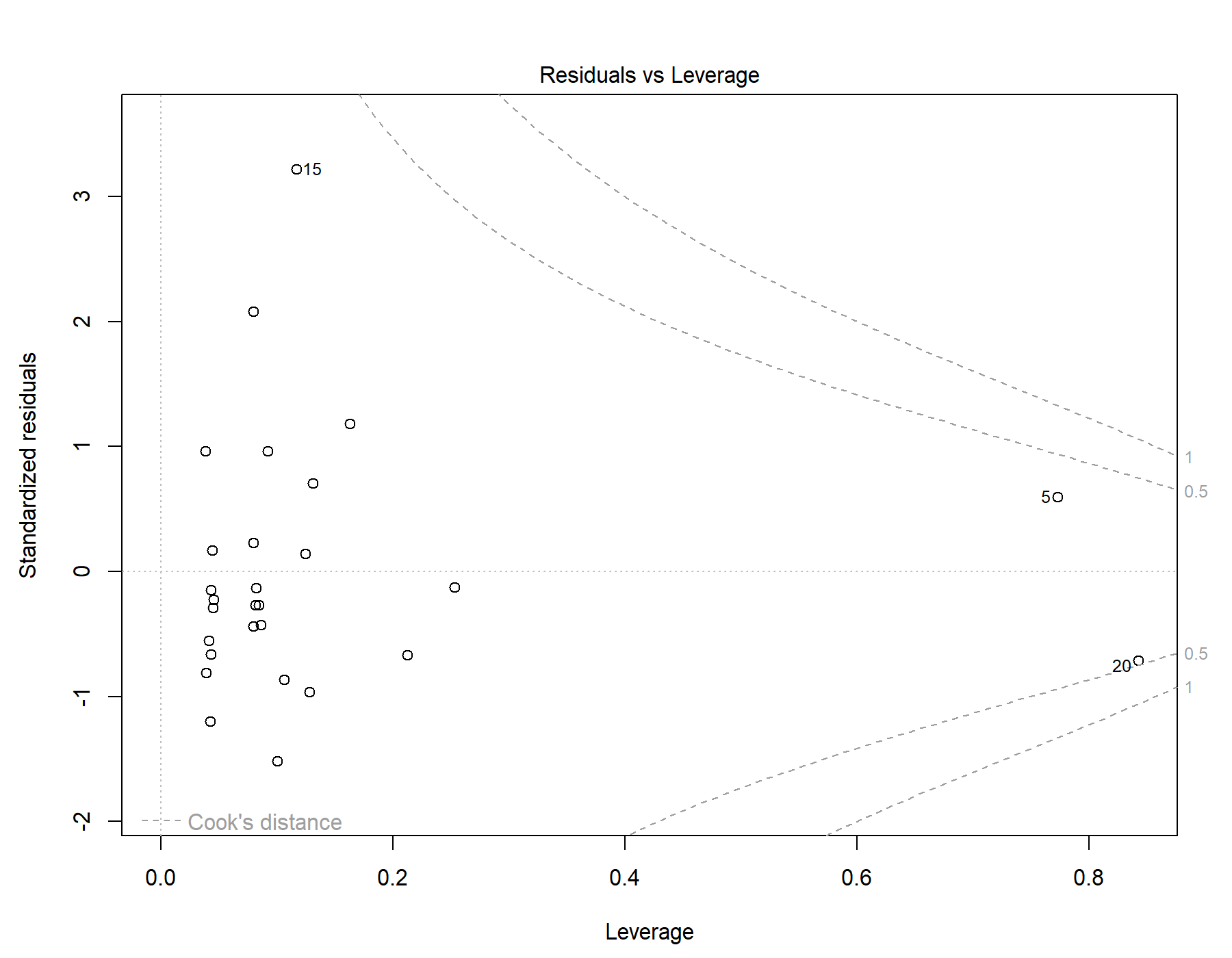

Supplemental Figure 100. Residuals vs leverage diagnostics for T1w-alone WML segmentation. The 2 older female subjects are not visualized due to possessing leverage values approaching 1 in this restricted model, preventing the normal calculation of their standardized residuals.

### 49. Global | Old only | T1w+FLAIR WML vol | SDNN.LF._log | Exploratory restricted interaction model

Model: T1w+FLAIR WML vol ~ SDNN.LF._log + Age + Sex + Age * SDNN.LF._log + Sex * SDNN.LF._log

#### 49.1. Standardized coefficients

| **Term** | **Estimate** | **Std. Error** | **t** | **p** | **β** |
| --- | --- | --- | --- | --- | --- |
| (Intercept) | -0.00533 | 0.00315 | -1.69 | .103 |  |
| SDNN.LF._log | -0.00213 | 0.00698 | -0.306 | .763 | -1.00 |
| Age | 5.425e-04 | 2.669e-04 | 2.03 | .053 | 0.412 |
| Sex | -8.805e-06 | 9.950e-04 | -0.00885 | .993 | -0.00184 |
| SDNN.LF._log:Age | -3.452e-04 | 6.121e-04 | -0.564 | .578 | -1.62 |
| SDNN.LF._log:Sex | 0.00262 | 0.00178 | 1.47 | .154 | 2.41 |

***Supplemental Table 154.*** *Standardized coefficients values for T1w+FLAIR WML segmentation.*

#### 49.2. 95% confidence intervals

| **Term** | **Lower 95%** | **Upper 95%** |
| --- | --- | --- |
| (Intercept) | -0.0118 | 0.00116 |
| SDNN.LF._log | -0.0165 | 0.0123 |
| Age | -8.303e-06 | 0.00109 |
| Sex | -0.00206 | 0.00204 |
| SDNN.LF._log:Age | -0.00161 | 9.181e-04 |
| SDNN.LF._log:Sex | -0.00106 | 0.00631 |

***Supplemental Table 155.*** *95% confidence intervals values for T1w+FLAIR WML segmentation.*

#### 49.3. Breusch-Pagan test

| **Test** | **Statistic** | **df** | **p** |
| --- | --- | --- | --- |
| Breusch-Pagan | 3.407 | 5 | .638 |

***Supplemental Table 156.*** *Breusch-Pagan test values for T1w+FLAIR WML segmentation.*

#### 49.4. Normal Q-Q plot

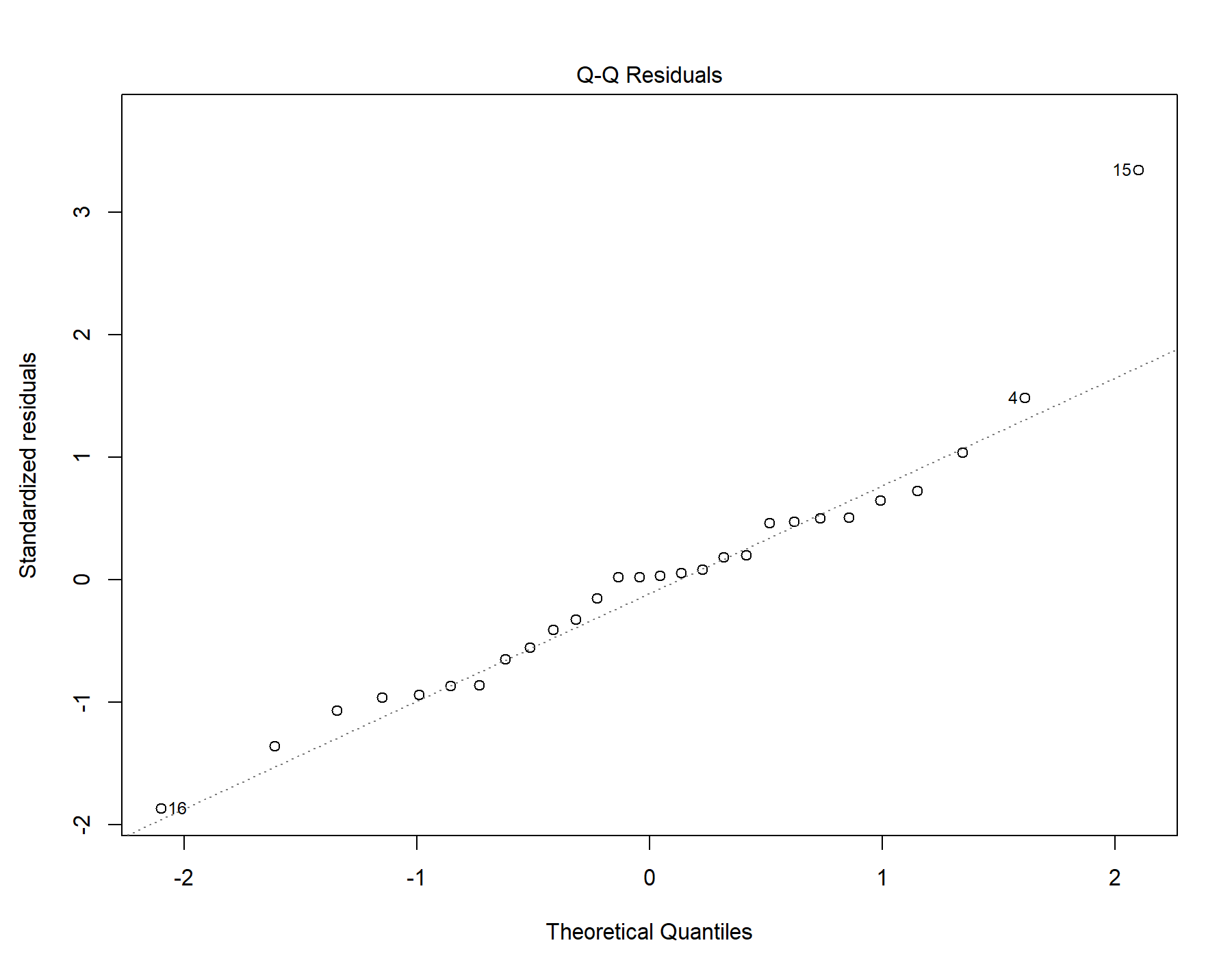

Supplemental Figure 101. Normal distribution quantile-quantile plot for T1w+FLAIR WML segmentation.

#### 49.5. Residuals vs leverage

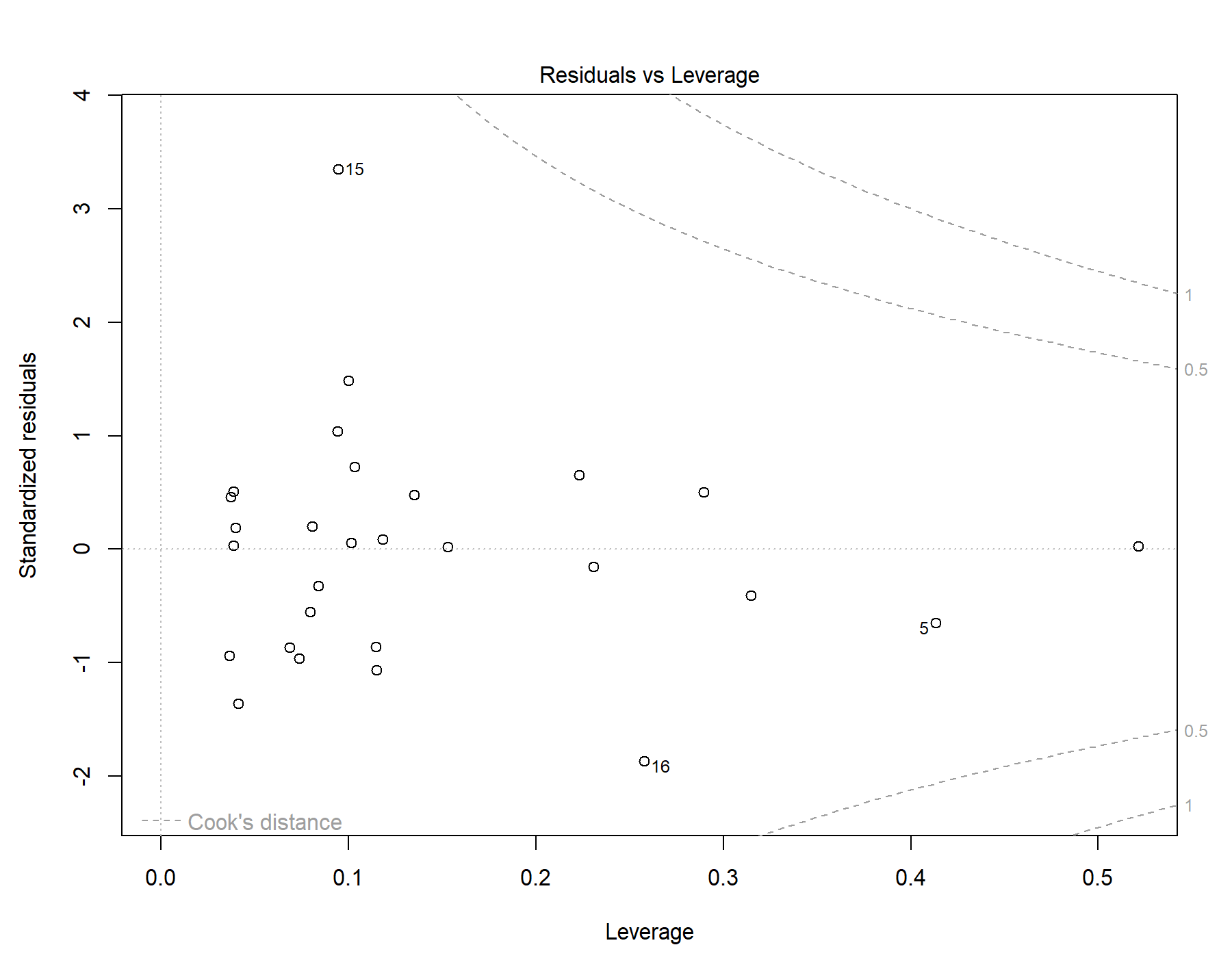

Supplemental Figure 102. Residuals vs leverage diagnostics for T1w+FLAIR WML segmentation. The 2 older female subjects are not visualized due to possessing leverage values approaching 1 in this restricted model, preventing the normal calculation of their standardized residuals.

### 50. Global | Old only | T1w+T2w WML vol | SDNN.LF._log | Exploratory restricted interaction model

Model: T1w+T2w WML vol ~ SDNN.LF._log + Age + Sex + Age * SDNN.LF._log + Sex * SDNN.LF._log

#### 50.1. Standardized coefficients

| **Term** | **Estimate** | **Std. Error** | **t** | **p** | **β** |
| --- | --- | --- | --- | --- | --- |
| (Intercept) | -0.00600 | 0.00332 | -1.81 | .083 |  |
| SDNN.LF._log | -0.00474 | 0.00736 | -0.645 | .525 | -2.09 |
| Age | 6.028e-04 | 2.813e-04 | 2.14 | .042 | 0.430 |
| Sex | 2.790e-05 | 0.00105 | 0.0266 | .979 | 0.00548 |
| SDNN.LF._log:Age | -2.351e-04 | 6.451e-04 | -0.364 | .719 | -1.03 |
| SDNN.LF._log:Sex | 0.00339 | 0.00188 | 1.80 | .084 | 2.92 |

***Supplemental Table 157.*** *Standardized coefficients values for T1w+T2w WML segmentation.*

#### 50.2. 95% confidence intervals

| **Term** | **Lower 95%** | **Upper 95%** |
| --- | --- | --- |
| (Intercept) | -0.0128 | 8.467e-04 |
| SDNN.LF._log | -0.0199 | 0.0104 |
| Age | 2.228e-05 | 0.00118 |
| Sex | -0.00214 | 0.00219 |
| SDNN.LF._log:Age | -0.00157 | 0.00110 |
| SDNN.LF._log:Sex | -4.885e-04 | 0.00728 |

***Supplemental Table 158.*** *95% confidence intervals values for T1w+T2w WML segmentation.*

#### 50.3. Breusch-Pagan test

| **Test** | **Statistic** | **df** | **p** |
| --- | --- | --- | --- |
| Breusch-Pagan | 2.387 | 5 | .793 |

***Supplemental Table 159.*** *Breusch-Pagan test values for T1w+T2w WML segmentation.*

#### 50.4. Normal Q-Q plot

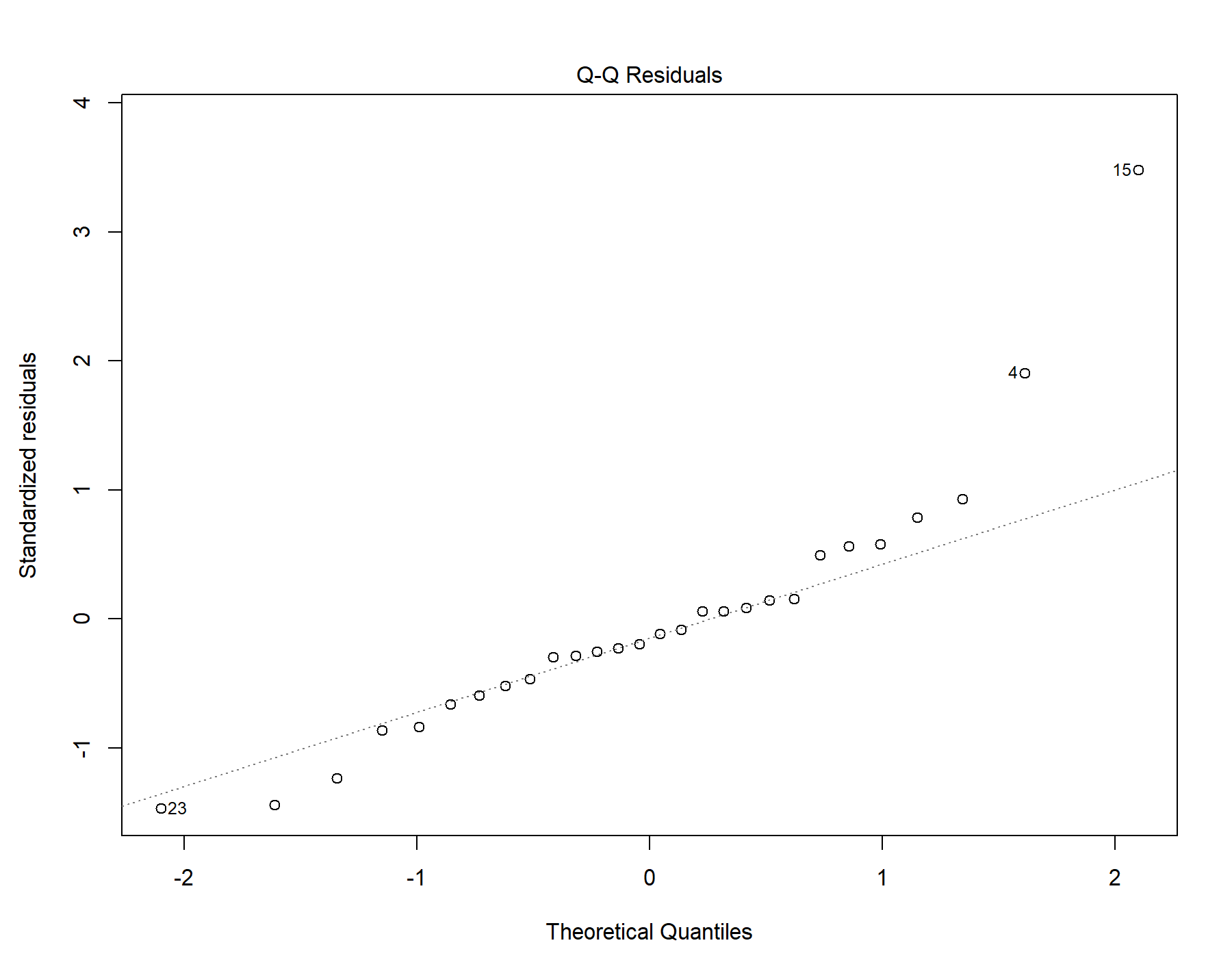

Supplemental Figure 103. Normal distribution quantile-quantile plot for T1w+T2w WML segmentation.

#### 50.5. Residuals vs leverage

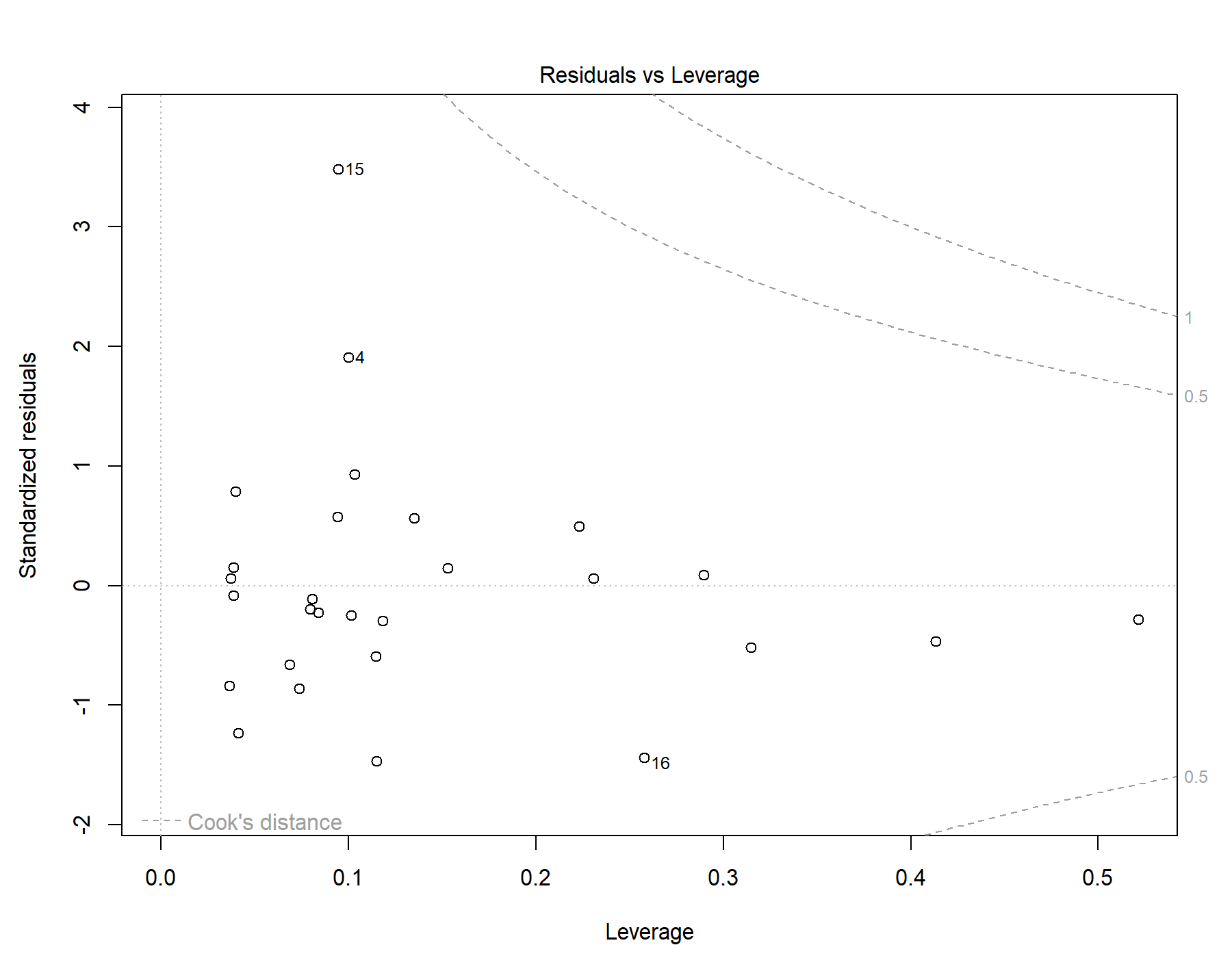

Supplemental Figure 104. Residuals vs leverage diagnostics for T1w+T2w WML segmentation. The 2 older female subjects are not visualized due to possessing leverage values approaching 1 in this restricted model, preventing the normal calculation of their standardized residuals.

### 51. Global | Old only | T1w-alone WML vol | SDNN.LF._log | Exploratory restricted interaction model

Model: T1w-alone WML vol ~ SDNN.LF._log + Age + Sex + Age * SDNN.LF._log + Sex * SDNN.LF._log

#### 51.1. Standardized coefficients

| **Term** | **Estimate** | **Std. Error** | **t** | **p** | **β** |
| --- | --- | --- | --- | --- | --- |
| (Intercept) | -0.00378 | 0.00282 | -1.34 | .193 |  |
| SDNN.LF._log | -0.00326 | 0.00625 | -0.521 | .607 | -1.75 |
| Age | 4.057e-04 | 2.390e-04 | 1.70 | .102 | 0.352 |
| Sex | -1.158e-04 | 8.909e-04 | -0.130 | .898 | -0.0276 |
| SDNN.LF._log:Age | -2.240e-04 | 5.480e-04 | -0.409 | .686 | -1.20 |
| SDNN.LF._log:Sex | 0.00261 | 0.00160 | 1.63 | .115 | 2.74 |

***Supplemental Table 160.*** *Standardized coefficients values for T1w-alone WML segmentation.*

#### 51.2. 95% confidence intervals

| **Term** | **Lower 95%** | **Upper 95%** |
| --- | --- | --- |
| (Intercept) | -0.00959 | 0.00204 |
| SDNN.LF._log | -0.0162 | 0.00964 |
| Age | -8.746e-05 | 8.989e-04 |
| Sex | -0.00195 | 0.00172 |
| SDNN.LF._log:Age | -0.00136 | 9.071e-04 |
| SDNN.LF._log:Sex | -6.881e-04 | 0.00591 |

***Supplemental Table 161.*** *95% confidence intervals values for T1w-alone WML segmentation.*

#### 51.3. Breusch-Pagan test

| **Test** | **Statistic** | **df** | **p** |
| --- | --- | --- | --- |
| Breusch-Pagan | 2.064 | 5 | .840 |

***Supplemental Table 162.*** *Breusch-Pagan test values for T1w-alone WML segmentation.*

#### 51.4. Normal Q-Q plot

Supplemental Figure 105. Normal distribution quantile-quantile plot for T1w-alone WML segmentation.

#### 51.5. Residuals vs leverage

Supplemental Figure 106. Residuals vs leverage diagnostics for T1w-alone WML segmentation. The 2 older female subjects are not visualized due to possessing leverage values approaching 1 in this restricted model, preventing the normal calculation of their standardized residuals.

### 52. Global | Old only | T1w+FLAIR WML vol | SDNN.HF._log | Exploratory restricted interaction model

Model: T1w+FLAIR WML vol ~ SDNN.HF._log + Age + Sex + Age * SDNN.HF._log + Sex * SDNN.HF._log

#### 52.1. Standardized coefficients

| **Term** | **Estimate** | **Std. Error** | **t** | **p** | **β** |
| --- | --- | --- | --- | --- | --- |
| (Intercept) | -0.00917 | 0.00387 | -2.37 | .026 |  |
| SDNN.HF._log | -0.0383 | 0.0246 | -1.55 | .133 | -16.2 |
| Age | 5.493e-04 | 2.637e-04 | 2.08 | .048 | 0.418 |
| Sex | 0.00188 | 0.00118 | 1.59 | .125 | 0.393 |
| SDNN.HF._log:Age | -2.540e-04 | 6.242e-04 | -0.407 | .688 | -1.07 |
| SDNN.HF._log:Sex | 0.0202 | 0.0122 | 1.66 | .109 | 17.1 |

***Supplemental Table 163.*** *Standardized coefficients values for T1w+FLAIR WML segmentation.*

#### 52.2. 95% confidence intervals

| **Term** | **Lower 95%** | **Upper 95%** |
| --- | --- | --- |
| (Intercept) | -0.0172 | -0.00118 |
| SDNN.HF._log | -0.0891 | 0.0126 |
| Age | 5.028e-06 | 0.00109 |
| Sex | -5.581e-04 | 0.00432 |
| SDNN.HF._log:Age | -0.00154 | 0.00103 |
| SDNN.HF._log:Sex | -0.00488 | 0.0453 |

***Supplemental Table 164.*** *95% confidence intervals values for T1w+FLAIR WML segmentation.*

#### 52.3. Breusch-Pagan test

| **Test** | **Statistic** | **df** | **p** |
| --- | --- | --- | --- |
| Breusch-Pagan | 3.353 | 5 | .646 |

***Supplemental Table 165.*** *Breusch-Pagan test values for T1w+FLAIR WML segmentation.*

#### 52.4. Normal Q-Q plot

Supplemental Figure 107. Normal distribution quantile-quantile plot for T1w+FLAIR WML segmentation.

#### 52.5. Residuals vs leverage

Supplemental Figure 108. Residuals vs leverage diagnostics for T1w+FLAIR WML segmentation. The 2 older female subjects are not visualized due to possessing leverage values approaching 1 in this restricted model, preventing the normal calculation of their standardized residuals.

### 53. Global | Old only | T1w+T2w WML vol | SDNN.HF._log | Exploratory restricted interaction model

Model: T1w+T2w WML vol ~ SDNN.HF._log + Age + Sex + Age * SDNN.HF._log + Sex * SDNN.HF._log

#### 53.1. Standardized coefficients

| **Term** | **Estimate** | **Std. Error** | **t** | **p** | **β** |
| --- | --- | --- | --- | --- | --- |
| (Intercept) | -0.0104 | 0.00405 | -2.57 | .017 |  |
| SDNN.HF._log | -0.0489 | 0.0258 | -1.90 | .070 | -19.4 |
| Age | 5.986e-04 | 2.756e-04 | 2.17 | .040 | 0.427 |
| Sex | 0.00226 | 0.00123 | 1.83 | .080 | 0.443 |
| SDNN.HF._log:Age | -2.133e-04 | 6.524e-04 | -0.327 | .747 | -0.847 |
| SDNN.HF._log:Sex | 0.0253 | 0.0127 | 1.99 | .058 | 20.1 |

***Supplemental Table 166.*** *Standardized coefficients values for T1w+T2w WML segmentation.*

#### 53.2. 95% confidence intervals

| **Term** | **Lower 95%** | **Upper 95%** |
| --- | --- | --- |
| (Intercept) | -0.0188 | -0.00206 |
| SDNN.HF._log | -0.102 | 0.00424 |
| Age | 2.976e-05 | 0.00117 |
| Sex | -2.904e-04 | 0.00480 |
| SDNN.HF._log:Age | -0.00156 | 0.00113 |
| SDNN.HF._log:Sex | -9.252e-04 | 0.0515 |

***Supplemental Table 167.*** *95% confidence intervals values for T1w+T2w WML segmentation.*

#### 53.3. Breusch-Pagan test

| **Test** | **Statistic** | **df** | **p** |
| --- | --- | --- | --- |
| Breusch-Pagan | 2.34 | 5 | .800 |

***Supplemental Table 168.*** *Breusch-Pagan test values for T1w+T2w WML segmentation.*

#### 53.4. Normal Q-Q plot

Supplemental Figure 109. Normal distribution quantile-quantile plot for T1w+T2w WML segmentation.

#### 53.5. Residuals vs leverage

Supplemental Figure 110. Residuals vs leverage diagnostics for T1w+T2w WML segmentation. The 2 older female subjects are not visualized due to possessing leverage values approaching 1 in this restricted model, preventing the normal calculation of their standardized residuals.

### 54. Global | Old only | T1w-alone WML vol | SDNN.HF._log | Exploratory restricted interaction model

Model: T1w-alone WML vol ~ SDNN.HF._log + Age + Sex + Age * SDNN.HF._log + Sex * SDNN.HF._log

#### 54.1. Standardized coefficients

| **Term** | **Estimate** | **Std. Error** | **t** | **p** | **β** |
| --- | --- | --- | --- | --- | --- |
| (Intercept) | -0.00738 | 0.00347 | -2.13 | .044 |  |
| SDNN.HF._log | -0.0391 | 0.0221 | -1.77 | .089 | -18.9 |
| Age | 4.110e-04 | 2.363e-04 | 1.74 | .095 | 0.357 |
| Sex | 0.00166 | 0.00106 | 1.57 | .129 | 0.397 |
| SDNN.HF._log:Age | -4.030e-05 | 5.592e-04 | -0.0721 | .943 | -0.195 |
| SDNN.HF._log:Sex | 0.0196 | 0.0109 | 1.80 | .085 | 18.9 |

***Supplemental Table 169.*** *Standardized coefficients values for T1w-alone WML segmentation.*

#### 54.2. 95% confidence intervals

| **Term** | **Lower 95%** | **Upper 95%** |
| --- | --- | --- |
| (Intercept) | -0.0145 | -2.206e-04 |
| SDNN.HF._log | -0.0847 | 0.00646 |
| Age | -7.661e-05 | 8.986e-04 |
| Sex | -5.198e-04 | 0.00385 |
| SDNN.HF._log:Age | -0.00119 | 0.00111 |
| SDNN.HF._log:Sex | -0.00289 | 0.0421 |

***Supplemental Table 170.*** *95% confidence intervals values for T1w-alone WML segmentation.*

#### 54.3. Breusch-Pagan test

| **Test** | **Statistic** | **df** | **p** |
| --- | --- | --- | --- |
| Breusch-Pagan | 2.186 | 5 | .823 |

***Supplemental Table 171.*** *Breusch-Pagan test values for T1w-alone WML segmentation.*

#### 54.4. Normal Q-Q plot

Supplemental Figure 111. Normal distribution quantile-quantile plot for T1w-alone WML segmentation.

#### 54.5. Residuals vs leverage

Supplemental Figure 112. Residuals vs leverage diagnostics for T1w-alone WML segmentation. The 2 older female subjects are not visualized due to possessing leverage values approaching 1 in this restricted model, preventing the normal calculation of their standardized residuals.

### 55. Periventricular | All subjects | T1w+FLAIR WML vol | RMSSD | Exploratory unadjusted model

Model: T1w+FLAIR WML vol ~ RMSSD

#### 55.1. Standardized coefficients

| **Term** | **Estimate** | **Std. Error** | **t** | **p** | **β** |
| --- | --- | --- | --- | --- | --- |
| (Intercept) | -6.135e-20 | 6.336e-05 | -9.682e-16 | 1.000 |  |
| RMSSD | -4.196e-06 | 1.527e-06 | -2.75 | .007 | -0.233 |

***Supplemental Table 172.*** *Standardized coefficients values for T1w+FLAIR WML segmentation.*

#### 55.2. 95% confidence intervals

| **Term** | **Lower 95%** | **Upper 95%** |
| --- | --- | --- |
| (Intercept) | -1.253e-04 | 1.253e-04 |
| RMSSD | -7.217e-06 | -1.175e-06 |

***Supplemental Table 173.*** *95% confidence intervals values for T1w+FLAIR WML segmentation.*

#### 55.3. Breusch-Pagan test

| **Test** | **Statistic** | **df** | **p** |
| --- | --- | --- | --- |
| Breusch-Pagan | 1.843 | 1 | .175 |

***Supplemental Table 174.*** *Breusch-Pagan test values for T1w+FLAIR WML segmentation.*

#### 55.4. Normal Q-Q plot

Supplemental Figure 113. Normal distribution quantile-quantile plot for T1w+FLAIR WML segmentation.

#### 55.5. Residuals vs leverage

Supplemental Figure 114. Residuals vs leverage diagnostics for T1w+FLAIR WML segmentation.

### 56. Periventricular | All subjects | T1w+T2w WML vol | RMSSD | Exploratory unadjusted model

Model: T1w+T2w WML vol ~ RMSSD

#### 56.1. Standardized coefficients

| **Term** | **Estimate** | **Std. Error** | **t** | **p** | **β** |
| --- | --- | --- | --- | --- | --- |
| (Intercept) | -6.054e-20 | 7.204e-05 | -8.403e-16 | 1.000 |  |
| RMSSD | -3.537e-06 | 1.736e-06 | -2.04 | .044 | -0.175 |

***Supplemental Table 175.*** *Standardized coefficients values for T1w+T2w WML segmentation.*

#### 56.2. 95% confidence intervals

| **Term** | **Lower 95%** | **Upper 95%** |
| --- | --- | --- |
| (Intercept) | -1.425e-04 | 1.425e-04 |
| RMSSD | -6.972e-06 | -1.023e-07 |

***Supplemental Table 176.*** *95% confidence intervals values for T1w+T2w WML segmentation.*

#### 56.3. Breusch-Pagan test

| **Test** | **Statistic** | **df** | **p** |
| --- | --- | --- | --- |
| Breusch-Pagan | 1.845 | 1 | .174 |

***Supplemental Table 177.*** *Breusch-Pagan test values for T1w+T2w WML segmentation.*

#### 56.4. Normal Q-Q plot

Supplemental Figure 115. Normal distribution quantile-quantile plot for T1w+T2w WML segmentation.

#### 56.5. Residuals vs leverage

Supplemental Figure 116. Residuals vs leverage diagnostics for T1w+T2w WML segmentation.

### 57. Periventricular | All subjects | T1w-alone WML vol | RMSSD | Exploratory unadjusted model

Model: T1w-alone WML vol ~ RMSSD

#### 57.1. Standardized coefficients

| **Term** | **Estimate** | **Std. Error** | **t** | **p** | **β** |
| --- | --- | --- | --- | --- | --- |
| (Intercept) | 6.170e-20 | 6.207e-05 | 9.941e-16 | 1.000 |  |
| RMSSD | -3.143e-06 | 1.496e-06 | -2.10 | .038 | -0.180 |

***Supplemental Table 178.*** *Standardized coefficients values for T1w-alone WML segmentation.*

#### 57.2. 95% confidence intervals

| **Term** | **Lower 95%** | **Upper 95%** |
| --- | --- | --- |
| (Intercept) | -1.228e-04 | 1.228e-04 |
| RMSSD | -6.102e-06 | -1.833e-07 |

***Supplemental Table 179.*** *95% confidence intervals values for T1w-alone WML segmentation.*

#### 57.3. Breusch-Pagan test

| **Test** | **Statistic** | **df** | **p** |
| --- | --- | --- | --- |
| Breusch-Pagan | 1.657 | 1 | .198 |

***Supplemental Table 180.*** *Breusch-Pagan test values for T1w-alone WML segmentation.*

#### 57.4. Normal Q-Q plot

Supplemental Figure 117. Normal distribution quantile-quantile plot for T1w-alone WML segmentation.

#### 57.5. Residuals vs leverage

Supplemental Figure 118. Residuals vs leverage diagnostics for T1w-alone WML segmentation.

### 58. Periventricular | All subjects | T1w+FLAIR WML vol | SDNN.LF._log | Exploratory unadjusted model

Model: T1w+FLAIR WML vol ~ SDNN.LF._log

#### 58.1. Standardized coefficients

| **Term** | **Estimate** | **Std. Error** | **t** | **p** | **β** |
| --- | --- | --- | --- | --- | --- |
| (Intercept) | -6.259e-20 | 6.360e-05 | -9.840e-16 | 1.000 |  |
| SDNN.LF._log | -4.100e-04 | 1.610e-04 | -2.55 | .012 | -0.216 |

***Supplemental Table 181.*** *Standardized coefficients values for T1w+FLAIR WML segmentation.*

#### 58.2. 95% confidence intervals

| **Term** | **Lower 95%** | **Upper 95%** |
| --- | --- | --- |
| (Intercept) | -1.258e-04 | 1.258e-04 |
| SDNN.LF._log | -7.284e-04 | -9.149e-05 |

***Supplemental Table 182.*** *95% confidence intervals values for T1w+FLAIR WML segmentation.*

#### 58.3. Breusch-Pagan test

| **Test** | **Statistic** | **df** | **p** |
| --- | --- | --- | --- |
| Breusch-Pagan | 1.355 | 1 | .244 |

***Supplemental Table 183.*** *Breusch-Pagan test values for T1w+FLAIR WML segmentation.*

#### 58.4. Normal Q-Q plot

Supplemental Figure 119. Normal distribution quantile-quantile plot for T1w+FLAIR WML segmentation.

#### 58.5. Residuals vs leverage

Supplemental Figure 120. Residuals vs leverage diagnostics for T1w+FLAIR WML segmentation.

### 59. Periventricular | All subjects | T1w+T2w WML vol | SDNN.LF._log | Exploratory unadjusted model

Model: T1w+T2w WML vol ~ SDNN.LF._log

#### 59.1. Standardized coefficients

| **Term** | **Estimate** | **Std. Error** | **t** | **p** | **β** |
| --- | --- | --- | --- | --- | --- |
| (Intercept) | -6.285e-20 | 7.168e-05 | -8.768e-16 | 1.000 |  |
| SDNN.LF._log | -4.268e-04 | 1.814e-04 | -2.35 | .020 | -0.201 |

***Supplemental Table 184.*** *Standardized coefficients values for T1w+T2w WML segmentation.*

#### 59.2. 95% confidence intervals

| **Term** | **Lower 95%** | **Upper 95%** |
| --- | --- | --- |
| (Intercept) | -1.418e-04 | 1.418e-04 |
| SDNN.LF._log | -7.857e-04 | -6.786e-05 |

***Supplemental Table 185.*** *95% confidence intervals values for T1w+T2w WML segmentation.*

#### 59.3. Breusch-Pagan test

| **Test** | **Statistic** | **df** | **p** |
| --- | --- | --- | --- |
| Breusch-Pagan | 1.11 | 1 | .292 |

***Supplemental Table 186.*** *Breusch-Pagan test values for T1w+T2w WML segmentation.*

#### 59.4. Normal Q-Q plot

Supplemental Figure 121. Normal distribution quantile-quantile plot for T1w+T2w WML segmentation.

#### 59.5. Residuals vs leverage

Supplemental Figure 122. Residuals vs leverage diagnostics for T1w+T2w WML segmentation.

### 60. Periventricular | All subjects | T1w-alone WML vol | SDNN.LF._log | Exploratory unadjusted model

Model: T1w-alone WML vol ~ SDNN.LF._log

#### 60.1. Standardized coefficients

| **Term** | **Estimate** | **Std. Error** | **t** | **p** | **β** |
| --- | --- | --- | --- | --- | --- |
| (Intercept) | 5.966e-20 | 6.174e-05 | 9.663e-16 | 1.000 |  |
| SDNN.LF._log | -3.788e-04 | 1.563e-04 | -2.42 | .017 | -0.206 |

***Supplemental Table 187.*** *Standardized coefficients values for T1w-alone WML segmentation.*

#### 60.2. 95% confidence intervals

| **Term** | **Lower 95%** | **Upper 95%** |
| --- | --- | --- |
| (Intercept) | -1.221e-04 | 1.221e-04 |
| SDNN.LF._log | -6.879e-04 | -6.960e-05 |

***Supplemental Table 188.*** *95% confidence intervals values for T1w-alone WML segmentation.*

#### 60.3. Breusch-Pagan test

| **Test** | **Statistic** | **df** | **p** |
| --- | --- | --- | --- |
| Breusch-Pagan | 1.221 | 1 | .269 |

***Supplemental Table 189.*** *Breusch-Pagan test values for T1w-alone WML segmentation.*

#### 60.4. Normal Q-Q plot

Supplemental Figure 123. Normal distribution quantile-quantile plot for T1w-alone WML segmentation.

#### 60.5. Residuals vs leverage

Supplemental Figure 124. Residuals vs leverage diagnostics for T1w-alone WML segmentation.

### 61. Periventricular | All subjects | T1w+FLAIR WML vol | SDNN.HF._log | Exploratory unadjusted model

Model: T1w+FLAIR WML vol ~ SDNN.HF._log

#### 61.1. Standardized coefficients

| **Term** | **Estimate** | **Std. Error** | **t** | **p** | **β** |
| --- | --- | --- | --- | --- | --- |
| (Intercept) | -1.093e-19 | 6.186e-05 | -1.767e-15 | 1.000 |  |
| SDNN.HF._log | -6.152e-04 | 1.621e-04 | -3.80 | <.001 | -0.314 |

***Supplemental Table 190.*** *Standardized coefficients values for T1w+FLAIR WML segmentation.*

#### 61.2. 95% confidence intervals

| **Term** | **Lower 95%** | **Upper 95%** |
| --- | --- | --- |
| (Intercept) | -1.224e-04 | 1.224e-04 |
| SDNN.HF._log | -9.358e-04 | -2.946e-04 |

***Supplemental Table 191.*** *95% confidence intervals values for T1w+FLAIR WML segmentation.*

#### 61.3. Breusch-Pagan test

| **Test** | **Statistic** | **df** | **p** |
| --- | --- | --- | --- |
| Breusch-Pagan | 1.184 | 1 | .277 |

***Supplemental Table 192.*** *Breusch-Pagan test values for T1w+FLAIR WML segmentation.*

#### 61.4. Normal Q-Q plot

Supplemental Figure 125. Normal distribution quantile-quantile plot for T1w+FLAIR WML segmentation.

#### 61.5. Residuals vs leverage

Supplemental Figure 126. Residuals vs leverage diagnostics for T1w+FLAIR WML segmentation.

### 62. Periventricular | All subjects | T1w+T2w WML vol | SDNN.HF._log | Exploratory unadjusted model

Model: T1w+T2w WML vol ~ SDNN.HF._log

#### 62.1. Standardized coefficients

| **Term** | **Estimate** | **Std. Error** | **t** | **p** | **β** |
| --- | --- | --- | --- | --- | --- |
| (Intercept) | -1.145e-19 | 6.965e-05 | -1.643e-15 | 1.000 |  |
| SDNN.HF._log | -6.749e-04 | 1.825e-04 | -3.70 | <.001 | -0.306 |

***Supplemental Table 193.*** *Standardized coefficients values for T1w+T2w WML segmentation.*

#### 62.2. 95% confidence intervals

| **Term** | **Lower 95%** | **Upper 95%** |
| --- | --- | --- |
| (Intercept) | -1.378e-04 | 1.378e-04 |
| SDNN.HF._log | -0.00104 | -3.139e-04 |

***Supplemental Table 194.*** *95% confidence intervals values for T1w+T2w WML segmentation.*

#### 62.3. Breusch-Pagan test

| **Test** | **Statistic** | **df** | **p** |
| --- | --- | --- | --- |
| Breusch-Pagan | 1.116 | 1 | .291 |

***Supplemental Table 195.*** *Breusch-Pagan test values for T1w+T2w WML segmentation.*

#### 62.4. Normal Q-Q plot

Supplemental Figure 127. Normal distribution quantile-quantile plot for T1w+T2w WML segmentation.

#### 62.5. Residuals vs leverage

Supplemental Figure 128. Residuals vs leverage diagnostics for T1w+T2w WML segmentation.

### 63. Periventricular | All subjects | T1w-alone WML vol | SDNN.HF._log | Exploratory unadjusted model

Model: T1w-alone WML vol ~ SDNN.HF._log

#### 63.1. Standardized coefficients

| **Term** | **Estimate** | **Std. Error** | **t** | **p** | **β** |
| --- | --- | --- | --- | --- | --- |
| (Intercept) | 1.577e-20 | 6.012e-05 | 2.624e-16 | 1.000 |  |
| SDNN.HF._log | -5.768e-04 | 1.575e-04 | -3.66 | <.001 | -0.304 |

***Supplemental Table 196.*** *Standardized coefficients values for T1w-alone WML segmentation.*

#### 63.2. 95% confidence intervals

| **Term** | **Lower 95%** | **Upper 95%** |
| --- | --- | --- |
| (Intercept) | -1.189e-04 | 1.189e-04 |
| SDNN.HF._log | -8.884e-04 | -2.652e-04 |

***Supplemental Table 197.*** *95% confidence intervals values for T1w-alone WML segmentation.*

#### 63.3. Breusch-Pagan test

| **Test** | **Statistic** | **df** | **p** |
| --- | --- | --- | --- |
| Breusch-Pagan | 0.97 | 1 | .325 |

***Supplemental Table 198.*** *Breusch-Pagan test values for T1w-alone WML segmentation.*

#### 63.4. Normal Q-Q plot

Supplemental Figure 129. Normal distribution quantile-quantile plot for T1w-alone WML segmentation.

#### 63.5. Residuals vs leverage

Supplemental Figure 130. Residuals vs leverage diagnostics for T1w-alone WML segmentation.

### 64. Periventricular | All subjects | T1w+FLAIR WML vol | RMSSD | Primary interaction model

Model: T1w+FLAIR WML vol ~ RMSSD + Age +	Sex + Age * RMSSD + Sex * RMSSD

#### 64.1. Standardized coefficients

| **Term** | **Estimate** | **Std. Error** | **t** | **p** | **β** |
| --- | --- | --- | --- | --- | --- |
| (Intercept) | -4.395e-04 | 1.828e-04 | -2.40 | .018 |  |
| RMSSD | 1.413e-06 | 5.429e-06 | 0.260 | .795 | 0.0783 |
| Age | 1.400e-04 | 1.453e-05 | 9.63 | <.001 | 0.658 |
| Sex | -3.805e-05 | 1.096e-04 | -0.347 | .729 | -0.0234 |
| RMSSD:Age | -5.127e-07 | 2.936e-07 | -1.75 | .083 | -0.181 |
| RMSSD:Sex | -3.489e-07 | 3.149e-06 | -0.111 | .912 | -0.0357 |

***Supplemental Table 199.*** *Standardized coefficients values for T1w+FLAIR WML segmentation.*

#### 64.2. 95% confidence intervals

| **Term** | **Lower 95%** | **Upper 95%** |
| --- | --- | --- |
| (Intercept) | -8.012e-04 | -7.779e-05 |
| RMSSD | -9.329e-06 | 1.216e-05 |
| Age | 1.112e-04 | 1.687e-04 |
| Sex | -2.549e-04 | 1.788e-04 |
| RMSSD:Age | -1.094e-06 | 6.822e-08 |
| RMSSD:Sex | -6.581e-06 | 5.883e-06 |

***Supplemental Table 200.*** *95% confidence intervals values for T1w+FLAIR WML segmentation.*

#### 64.3. Breusch-Pagan test

| **Test** | **Statistic** | **df** | **p** |
| --- | --- | --- | --- |
| Breusch-Pagan | 16.913 | 5 | .005 |

***Supplemental Table 201.*** *Breusch-Pagan test values for T1w+FLAIR WML segmentation.*

#### 64.4. Normal Q-Q plot

Supplemental Figure 131. Normal distribution quantile-quantile plot for T1w+FLAIR WML segmentation.

#### 64.5. Residuals vs leverage

Supplemental Figure 132. Residuals vs leverage diagnostics for T1w+FLAIR WML segmentation.

### 65. Periventricular | All subjects | T1w+T2w WML vol | RMSSD | Primary interaction model

Model: T1w+T2w WML vol ~ RMSSD + Age +	Sex + Age * RMSSD + Sex * RMSSD

#### 65.1. Standardized coefficients

| **Term** | **Estimate** | **Std. Error** | **t** | **p** | **β** |
| --- | --- | --- | --- | --- | --- |
| (Intercept) | -3.930e-04 | 2.017e-04 | -1.95 | .054 |  |
| RMSSD | 4.621e-06 | 5.992e-06 | 0.771 | .442 | 0.228 |
| Age | 1.674e-04 | 1.604e-05 | 10.4 | <.001 | 0.701 |
| Sex | -1.216e-04 | 1.209e-04 | -1.01 | .317 | -0.0667 |
| RMSSD:Age | -5.434e-07 | 3.240e-07 | -1.68 | .096 | -0.171 |
| RMSSD:Sex | -1.361e-06 | 3.476e-06 | -0.391 | .696 | -0.124 |

***Supplemental Table 202.*** *Standardized coefficients values for T1w+T2w WML segmentation.*

#### 65.2. 95% confidence intervals

| **Term** | **Lower 95%** | **Upper 95%** |
| --- | --- | --- |
| (Intercept) | -7.921e-04 | 6.222e-06 |
| RMSSD | -7.234e-06 | 1.648e-05 |
| Age | 1.357e-04 | 1.992e-04 |
| Sex | -3.609e-04 | 1.177e-04 |
| RMSSD:Age | -1.185e-06 | 9.768e-08 |
| RMSSD:Sex | -8.238e-06 | 5.517e-06 |

***Supplemental Table 203.*** *95% confidence intervals values for T1w+T2w WML segmentation.*

#### 65.3. Breusch-Pagan test

| **Test** | **Statistic** | **df** | **p** |
| --- | --- | --- | --- |
| Breusch-Pagan | 17.114 | 5 | .004 |

***Supplemental Table 204.*** *Breusch-Pagan test values for T1w+T2w WML segmentation.*

#### 65.4. Normal Q-Q plot

Supplemental Figure 133. Normal distribution quantile-quantile plot for T1w+T2w WML segmentation.

#### 65.5. Residuals vs leverage

Supplemental Figure 134. Residuals vs leverage diagnostics for T1w+T2w WML segmentation.

### 66. Periventricular | All subjects | T1w-alone WML vol | RMSSD | Primary interaction model

Model: T1w-alone WML vol ~ RMSSD + Age + Sex +	Age * RMSSD + Sex * RMSSD

#### 66.1. Standardized coefficients

| **Term** | **Estimate** | **Std. Error** | **t** | **p** | **β** |
| --- | --- | --- | --- | --- | --- |
| (Intercept) | -3.300e-04 | 1.751e-04 | -1.88 | .062 |  |
| RMSSD | 3.875e-06 | 5.202e-06 | 0.745 | .458 | 0.222 |
| Age | 1.437e-04 | 1.392e-05 | 10.3 | <.001 | 0.698 |
| Sex | -1.078e-04 | 1.050e-04 | -1.03 | .307 | -0.0685 |
| RMSSD:Age | -4.262e-07 | 2.813e-07 | -1.52 | .132 | -0.156 |
| RMSSD:Sex | -1.281e-06 | 3.018e-06 | -0.424 | .672 | -0.135 |

***Supplemental Table 205.*** *Standardized coefficients values for T1w-alone WML segmentation.*

#### 66.2. 95% confidence intervals

| **Term** | **Lower 95%** | **Upper 95%** |
| --- | --- | --- |
| (Intercept) | -6.765e-04 | 1.660e-05 |
| RMSSD | -6.418e-06 | 1.417e-05 |
| Age | 1.161e-04 | 1.712e-04 |
| Sex | -3.155e-04 | 9.998e-05 |
| RMSSD:Age | -9.828e-07 | 1.304e-07 |
| RMSSD:Sex | -7.252e-06 | 4.691e-06 |

***Supplemental Table 206.*** *95% confidence intervals values for T1w-alone WML segmentation.*

#### 66.3. Breusch-Pagan test

| **Test** | **Statistic** | **df** | **p** |
| --- | --- | --- | --- |
| Breusch-Pagan | 19.945 | 5 | .001 |

***Supplemental Table 207.*** *Breusch-Pagan test values for T1w-alone WML segmentation.*

#### 66.4. Normal Q-Q plot

Supplemental Figure 135. Normal distribution quantile-quantile plot for T1w-alone WML segmentation.

#### 66.5. Residuals vs leverage

Supplemental Figure 136. Residuals vs leverage diagnostics for T1w-alone WML segmentation.

### 67. Periventricular | All subjects | T1w+FLAIR WML vol | SDNN.LF._log | Primary interaction model

Model: T1w+FLAIR WML vol ~ SDNN.LF._log + Age + Sex + Age * SDNN.LF._log + Sex * SDNN.LF._log

#### 67.1. Standardized coefficients

| **Term** | **Estimate** | **Std. Error** | **t** | **p** | **β** |
| --- | --- | --- | --- | --- | --- |
| (Intercept) | -4.027e-04 | 1.786e-04 | -2.25 | .026 |  |
| SDNN.LF._log | -8.914e-04 | 5.594e-04 | -1.59 | .114 | -0.470 |
| Age | 1.438e-04 | 1.425e-05 | 10.1 | <.001 | 0.677 |
| Sex | -6.862e-05 | 1.067e-04 | -0.643 | .521 | -0.0422 |
| SDNN.LF._log:Age | -7.239e-05 | 2.995e-05 | -2.42 | .017 | -0.265 |
| SDNN.LF._log:Sex | 6.327e-04 | 3.234e-04 | 1.96 | .053 | 0.618 |

***Supplemental Table 208.*** *Standardized coefficients values for T1w+FLAIR WML segmentation.*

#### 67.2. 95% confidence intervals

| **Term** | **Lower 95%** | **Upper 95%** |
| --- | --- | --- |
| (Intercept) | -7.562e-04 | -4.929e-05 |
| SDNN.LF._log | -0.00200 | 2.155e-04 |
| Age | 1.156e-04 | 1.720e-04 |
| Sex | -2.798e-04 | 1.426e-04 |
| SDNN.LF._log:Age | -1.317e-04 | -1.313e-05 |
| SDNN.LF._log:Sex | -7.095e-06 | 0.00127 |

***Supplemental Table 209.*** *95% confidence intervals values for T1w+FLAIR WML segmentation.*

#### 67.3. Breusch-Pagan test

| **Test** | **Statistic** | **df** | **p** |
| --- | --- | --- | --- |
| Breusch-Pagan | 14.436 | 5 | .013 |

***Supplemental Table 210.*** *Breusch-Pagan test values for T1w+FLAIR WML segmentation.*

#### 67.4. Normal Q-Q plot

Supplemental Figure 137. Normal distribution quantile-quantile plot for T1w+FLAIR WML segmentation.

#### 67.5. Residuals vs leverage

Supplemental Figure 138. Residuals vs leverage diagnostics for T1w+FLAIR WML segmentation.

### 68. Periventricular | All subjects | T1w+T2w WML vol | SDNN.LF._log | Primary interaction model

Model: T1w+T2w WML vol ~ SDNN.LF._log + Age + Sex + Age * SDNN.LF._log + Sex * SDNN.LF._log

#### 68.1. Standardized coefficients

| **Term** | **Estimate** | **Std. Error** | **t** | **p** | **β** |
| --- | --- | --- | --- | --- | --- |
| (Intercept) | -3.900e-04 | 1.970e-04 | -1.98 | .050 |  |
| SDNN.LF._log | -6.276e-04 | 6.170e-04 | -1.02 | .311 | -0.295 |
| Age | 1.678e-04 | 1.571e-05 | 10.7 | <.001 | 0.703 |
| Sex | -1.275e-04 | 1.177e-04 | -1.08 | .281 | -0.0699 |
| SDNN.LF._log:Age | -8.119e-05 | 3.304e-05 | -2.46 | .015 | -0.265 |
| SDNN.LF._log:Sex | 5.275e-04 | 3.566e-04 | 1.48 | .142 | 0.459 |

***Supplemental Table 211.*** *Standardized coefficients values for T1w+T2w WML segmentation.*

#### 68.2. 95% confidence intervals

| **Term** | **Lower 95%** | **Upper 95%** |
| --- | --- | --- |
| (Intercept) | -7.798e-04 | -1.317e-07 |
| SDNN.LF._log | -0.00185 | 5.933e-04 |
| Age | 1.367e-04 | 1.989e-04 |
| Sex | -3.604e-04 | 1.055e-04 |
| SDNN.LF._log:Age | -1.466e-04 | -1.582e-05 |
| SDNN.LF._log:Sex | -1.782e-04 | 0.00123 |

***Supplemental Table 212.*** *95% confidence intervals values for T1w+T2w WML segmentation.*

#### 68.3. Breusch-Pagan test

| **Test** | **Statistic** | **df** | **p** |
| --- | --- | --- | --- |
| Breusch-Pagan | 12.823 | 5 | .025 |

***Supplemental Table 213.*** *Breusch-Pagan test values for T1w+T2w WML segmentation.*

#### 68.4. Normal Q-Q plot

Supplemental Figure 139. Normal distribution quantile-quantile plot for T1w+T2w WML segmentation.

#### 68.5. Residuals vs leverage

Supplemental Figure 140. Residuals vs leverage diagnostics for T1w+T2w WML segmentation.

### 69. Periventricular | All subjects | T1w-alone WML vol | SDNN.LF._log | Primary interaction model

Model: T1w-alone WML vol ~ SDNN.LF._log + Age + Sex + Age * SDNN.LF._log + Sex * SDNN.LF._log

#### 69.1. Standardized coefficients

| **Term** | **Estimate** | **Std. Error** | **t** | **p** | **β** |
| --- | --- | --- | --- | --- | --- |
| (Intercept) | -3.245e-04 | 1.705e-04 | -1.90 | .059 |  |
| SDNN.LF._log | -5.148e-04 | 5.340e-04 | -0.964 | .337 | -0.281 |
| Age | 1.437e-04 | 1.360e-05 | 10.6 | <.001 | 0.698 |
| Sex | -1.153e-04 | 1.019e-04 | -1.13 | .260 | -0.0733 |
| SDNN.LF._log:Age | -7.154e-05 | 2.859e-05 | -2.50 | .014 | -0.270 |
| SDNN.LF._log:Sex | 4.379e-04 | 3.087e-04 | 1.42 | .158 | 0.442 |

***Supplemental Table 214.*** *Standardized coefficients values for T1w-alone WML segmentation.*

#### 69.2. 95% confidence intervals

| **Term** | **Lower 95%** | **Upper 95%** |
| --- | --- | --- |
| (Intercept) | -6.619e-04 | 1.284e-05 |
| SDNN.LF._log | -0.00157 | 5.419e-04 |
| Age | 1.168e-04 | 1.706e-04 |
| Sex | -3.169e-04 | 8.633e-05 |
| SDNN.LF._log:Age | -1.281e-04 | -1.496e-05 |
| SDNN.LF._log:Sex | -1.729e-04 | 0.00105 |

***Supplemental Table 215.*** *95% confidence intervals values for T1w-alone WML segmentation.*

#### 69.3. Breusch-Pagan test

| **Test** | **Statistic** | **df** | **p** |
| --- | --- | --- | --- |
| Breusch-Pagan | 15.406 | 5 | .009 |

***Supplemental Table 216.*** *Breusch-Pagan test values for T1w-alone WML segmentation.*

#### 69.4. Normal Q-Q plot

Supplemental Figure 141. Normal distribution quantile-quantile plot for T1w-alone WML segmentation.

#### 69.5. Residuals vs leverage

Supplemental Figure 142. Residuals vs leverage diagnostics for T1w-alone WML segmentation.

### 70. Periventricular | All subjects | T1w+FLAIR WML vol | SDNN.HF._log | Primary interaction model

Model: T1w+FLAIR WML vol ~ SDNN.HF._log + Age + Sex + Age * SDNN.HF._log + Sex * SDNN.HF._log

#### 70.1. Standardized coefficients

| **Term** | **Estimate** | **Std. Error** | **t** | **p** | **β** |
| --- | --- | --- | --- | --- | --- |
| (Intercept) | -4.341e-04 | 1.816e-04 | -2.39 | .018 |  |
| SDNN.HF._log | -1.975e-04 | 5.873e-04 | -0.336 | .737 | -0.101 |
| Age | 1.367e-04 | 1.530e-05 | 8.93 | <.001 | 0.643 |
| Sex | -3.783e-05 | 1.095e-04 | -0.346 | .730 | -0.0233 |
| SDNN.HF._log:Age | -4.570e-05 | 3.388e-05 | -1.35 | .180 | -0.159 |
| SDNN.HF._log:Sex | 1.383e-04 | 3.526e-04 | 0.392 | .696 | 0.130 |

***Supplemental Table 217.*** *Standardized coefficients values for T1w+FLAIR WML segmentation.*

#### 70.2. 95% confidence intervals

| **Term** | **Lower 95%** | **Upper 95%** |
| --- | --- | --- |
| (Intercept) | -7.935e-04 | -7.474e-05 |
| SDNN.HF._log | -0.00136 | 9.646e-04 |
| Age | 1.064e-04 | 1.669e-04 |
| Sex | -2.545e-04 | 1.788e-04 |
| SDNN.HF._log:Age | -1.127e-04 | 2.133e-05 |
| SDNN.HF._log:Sex | -5.594e-04 | 8.360e-04 |

***Supplemental Table 218.*** *95% confidence intervals values for T1w+FLAIR WML segmentation.*

#### 70.3. Breusch-Pagan test

| **Test** | **Statistic** | **df** | **p** |
| --- | --- | --- | --- |
| Breusch-Pagan | 14.807 | 5 | .011 |

***Supplemental Table 219.*** *Breusch-Pagan test values for T1w+FLAIR WML segmentation.*

#### 70.4. Normal Q-Q plot

Supplemental Figure 143. Normal distribution quantile-quantile plot for T1w+FLAIR WML segmentation.

#### 70.5. Residuals vs leverage

Supplemental Figure 144. Residuals vs leverage diagnostics for T1w+FLAIR WML segmentation.

### 71. Periventricular | All subjects | T1w+T2w WML vol | SDNN.HF._log | Primary interaction model

Model: T1w+T2w WML vol ~ SDNN.HF._log + Age + Sex + Age * SDNN.HF._log + Sex * SDNN.HF._log

#### 71.1. Standardized coefficients

| **Term** | **Estimate** | **Std. Error** | **t** | **p** | **β** |
| --- | --- | --- | --- | --- | --- |
| (Intercept) | -4.125e-04 | 1.991e-04 | -2.07 | .040 |  |
| SDNN.HF._log | 2.917e-04 | 6.438e-04 | 0.453 | .651 | 0.132 |
| Age | 1.599e-04 | 1.677e-05 | 9.53 | <.001 | 0.670 |
| Sex | -9.890e-05 | 1.200e-04 | -0.824 | .412 | -0.0542 |
| SDNN.HF._log:Age | -4.892e-05 | 3.713e-05 | -1.32 | .190 | -0.152 |
| SDNN.HF._log:Sex | -1.165e-04 | 3.865e-04 | -0.301 | .764 | -0.0976 |

***Supplemental Table 220.*** *Standardized coefficients values for T1w+T2w WML segmentation.*

#### 71.2. 95% confidence intervals

| **Term** | **Lower 95%** | **Upper 95%** |
| --- | --- | --- |
| (Intercept) | -8.064e-04 | -1.853e-05 |
| SDNN.HF._log | -9.822e-04 | 0.00157 |
| Age | 1.267e-04 | 1.931e-04 |
| Sex | -3.364e-04 | 1.386e-04 |
| SDNN.HF._log:Age | -1.224e-04 | 2.456e-05 |
| SDNN.HF._log:Sex | -8.813e-04 | 6.483e-04 |

***Supplemental Table 221.*** *95% confidence intervals values for T1w+T2w WML segmentation.*

#### 71.3. Breusch-Pagan test

| **Test** | **Statistic** | **df** | **p** |
| --- | --- | --- | --- |
| Breusch-Pagan | 13.856 | 5 | .017 |

***Supplemental Table 222.*** *Breusch-Pagan test values for T1w+T2w WML segmentation.*

#### 71.4. Normal Q-Q plot

Supplemental Figure 145. Normal distribution quantile-quantile plot for T1w+T2w WML segmentation.

#### 71.5. Residuals vs leverage

Supplemental Figure 146. Residuals vs leverage diagnostics for T1w+T2w WML segmentation.

### 72. Periventricular | All subjects | T1w-alone WML vol | SDNN.HF._log | Primary interaction model

Model: T1w-alone WML vol ~ SDNN.HF._log + Age + Sex + Age * SDNN.HF._log + Sex * SDNN.HF._log

#### 72.1. Standardized coefficients

| **Term** | **Estimate** | **Std. Error** | **t** | **p** | **β** |
| --- | --- | --- | --- | --- | --- |
| (Intercept) | -3.451e-04 | 1.729e-04 | -2.0 | .048 |  |
| SDNN.HF._log | 1.695e-04 | 5.591e-04 | 0.303 | .762 | 0.0892 |
| Age | 1.379e-04 | 1.457e-05 | 9.46 | <.001 | 0.670 |
| Sex | -9.110e-05 | 1.042e-04 | -0.874 | .384 | -0.0579 |
| SDNN.HF._log:Age | -4.038e-05 | 3.225e-05 | -1.25 | .213 | -0.145 |
| SDNN.HF._log:Sex | -5.729e-05 | 3.357e-04 | -0.171 | .865 | -0.0556 |

***Supplemental Table 223.*** *Standardized coefficients values for T1w-alone WML segmentation.*

#### 72.2. 95% confidence intervals

| **Term** | **Lower 95%** | **Upper 95%** |
| --- | --- | --- |
| (Intercept) | -6.872e-04 | -3.035e-06 |
| SDNN.HF._log | -9.368e-04 | 0.00128 |
| Age | 1.090e-04 | 1.667e-04 |
| Sex | -2.974e-04 | 1.152e-04 |
| SDNN.HF._log:Age | -1.042e-04 | 2.342e-05 |
| SDNN.HF._log:Sex | -7.215e-04 | 6.069e-04 |

***Supplemental Table 224.*** *95% confidence intervals values for T1w-alone WML segmentation.*

#### 72.3. Breusch-Pagan test

| **Test** | **Statistic** | **df** | **p** |
| --- | --- | --- | --- |
| Breusch-Pagan | 16.903 | 5 | .005 |

***Supplemental Table 225.*** *Breusch-Pagan test values for T1w-alone WML segmentation.*

#### 72.4. Normal Q-Q plot

Supplemental Figure 147. Normal distribution quantile-quantile plot for T1w-alone WML segmentation.

#### 72.5. Residuals vs leverage

Supplemental Figure 148. Residuals vs leverage diagnostics for T1w-alone WML segmentation.

### 73. Periventricular | Female only | T1w+FLAIR WML vol | RMSSD | Exploratory restricted interaction model

Model: T1w+FLAIR WML vol ~ RMSSD + Age +	Age * RMSSD

#### 73.1. Standardized coefficients

| **Term** | **Estimate** | **Std. Error** | **t** | **p** | **β** |
| --- | --- | --- | --- | --- | --- |
| (Intercept) | -4.168e-04 | 1.292e-04 | -3.23 | .003 |  |
| RMSSD | -8.374e-06 | 3.444e-06 | -2.43 | .020 | -0.693 |
| Age | 2.771e-04 | 8.144e-05 | 3.40 | .002 | 1.32 |
| RMSSD:Age | 7.831e-06 | 2.973e-06 | 2.63 | .012 | 1.32 |

***Supplemental Table 226.*** *Standardized coefficients values for T1w+FLAIR WML segmentation.*

#### 73.2. 95% confidence intervals

| **Term** | **Lower 95%** | **Upper 95%** |
| --- | --- | --- |
| (Intercept) | -6.786e-04 | -1.551e-04 |
| RMSSD | -1.535e-05 | -1.395e-06 |
| Age | 1.121e-04 | 4.421e-04 |
| RMSSD:Age | 1.808e-06 | 1.385e-05 |

***Supplemental Table 227.*** *95% confidence intervals values for T1w+FLAIR WML segmentation.*

#### 73.3. Breusch-Pagan test

| **Test** | **Statistic** | **df** | **p** |
| --- | --- | --- | --- |
| Breusch-Pagan | 1.865 | 3 | .601 |

***Supplemental Table 228.*** *Breusch-Pagan test values for T1w+FLAIR WML segmentation.*

#### 73.4. Normal Q-Q plot

Supplemental Figure 149. Normal distribution quantile-quantile plot for T1w+FLAIR WML segmentation.

#### 73.5. Residuals vs leverage

Supplemental Figure 150. Residuals vs leverage diagnostics for T1w+FLAIR WML segmentation.

### 74. Periventricular | Female only | T1w+T2w WML vol | RMSSD | Exploratory restricted interaction model

Model: T1w+T2w WML vol ~ RMSSD + Age +	Age * RMSSD

#### 74.1. Standardized coefficients

| **Term** | **Estimate** | **Std. Error** | **t** | **p** | **β** |
| --- | --- | --- | --- | --- | --- |
| (Intercept) | -5.134e-04 | 1.607e-04 | -3.20 | .003 |  |
| RMSSD | -8.367e-06 | 4.283e-06 | -1.95 | .058 | -0.564 |
| Age | 3.425e-04 | 1.013e-04 | 3.38 | .002 | 1.32 |
| RMSSD:Age | 9.805e-06 | 3.697e-06 | 2.65 | .012 | 1.35 |

***Supplemental Table 229.*** *Standardized coefficients values for T1w+T2w WML segmentation.*

#### 74.2. 95% confidence intervals

| **Term** | **Lower 95%** | **Upper 95%** |
| --- | --- | --- |
| (Intercept) | -8.389e-04 | -1.879e-04 |
| RMSSD | -1.705e-05 | 3.118e-07 |
| Age | 1.373e-04 | 5.477e-04 |
| RMSSD:Age | 2.314e-06 | 1.730e-05 |

***Supplemental Table 230.*** *95% confidence intervals values for T1w+T2w WML segmentation.*

#### 74.3. Breusch-Pagan test

| **Test** | **Statistic** | **df** | **p** |
| --- | --- | --- | --- |
| Breusch-Pagan | 4.038 | 3 | .257 |

***Supplemental Table 231.*** *Breusch-Pagan test values for T1w+T2w WML segmentation.*

#### 74.4. Normal Q-Q plot

Supplemental Figure 151. Normal distribution quantile-quantile plot for T1w+T2w WML segmentation.

#### 74.5. Residuals vs leverage

Supplemental Figure 152. Residuals vs leverage diagnostics for T1w+T2w WML segmentation.

### 75. Periventricular | Female only | T1w-alone WML vol | RMSSD | Exploratory restricted interaction model

Model: T1w-alone WML vol ~ RMSSD + Age + Age *	RMSSD

#### 75.1. Standardized coefficients

| **Term** | **Estimate** | **Std. Error** | **t** | **p** | **β** |
| --- | --- | --- | --- | --- | --- |
| (Intercept) | -4.311e-04 | 1.570e-04 | -2.75 | .009 |  |
| RMSSD | -6.551e-06 | 4.186e-06 | -1.57 | .126 | -0.466 |
| Age | 2.838e-04 | 9.898e-05 | 2.87 | .007 | 1.16 |
| RMSSD:Age | 7.738e-06 | 3.613e-06 | 2.14 | .039 | 1.12 |

***Supplemental Table 232.*** *Standardized coefficients values for T1w-alone WML segmentation.*

#### 75.2. 95% confidence intervals

| **Term** | **Lower 95%** | **Upper 95%** |
| --- | --- | --- |
| (Intercept) | -7.492e-04 | -1.129e-04 |
| RMSSD | -1.503e-05 | 1.930e-06 |
| Age | 8.322e-05 | 4.843e-04 |
| RMSSD:Age | 4.168e-07 | 1.506e-05 |

***Supplemental Table 233.*** *95% confidence intervals values for T1w-alone WML segmentation.*

#### 75.3. Breusch-Pagan test

| **Test** | **Statistic** | **df** | **p** |
| --- | --- | --- | --- |
| Breusch-Pagan | 3.362 | 3 | .339 |

***Supplemental Table 234.*** *Breusch-Pagan test values for T1w-alone WML segmentation.*

#### 75.4. Normal Q-Q plot

Supplemental Figure 153. Normal distribution quantile-quantile plot for T1w-alone WML segmentation.

#### 75.5. Residuals vs leverage

Supplemental Figure 154. Residuals vs leverage diagnostics for T1w-alone WML segmentation.

### 76. Periventricular | Female only | T1w+FLAIR WML vol | SDNN.LF._log | Exploratory restricted interaction model

Model: T1w+FLAIR WML vol ~ SDNN.LF._log + Age +	Age * SDNN.LF._log

#### 76.1. Standardized coefficients

| **Term** | **Estimate** | **Std. Error** | **t** | **p** | **β** |
| --- | --- | --- | --- | --- | --- |
| (Intercept) | -1.769e-04 | 7.620e-05 | -2.32 | .026 |  |
| SDNN.LF._log | -1.350e-04 | 2.215e-04 | -0.610 | .546 | -0.106 |
| Age | 8.928e-05 | 2.853e-05 | 3.13 | .003 | 0.424 |
| SDNN.LF._log:Age | -1.337e-04 | 6.071e-05 | -2.20 | .034 | -0.387 |

***Supplemental Table 235.*** *Standardized coefficients values for T1w+FLAIR WML segmentation.*

#### 76.2. 95% confidence intervals

| **Term** | **Lower 95%** | **Upper 95%** |
| --- | --- | --- |
| (Intercept) | -3.313e-04 | -2.253e-05 |
| SDNN.LF._log | -5.837e-04 | 3.137e-04 |
| Age | 3.147e-05 | 1.471e-04 |
| SDNN.LF._log:Age | -2.567e-04 | -1.065e-05 |

***Supplemental Table 236.*** *95% confidence intervals values for T1w+FLAIR WML segmentation.*

#### 76.3. Breusch-Pagan test

| **Test** | **Statistic** | **df** | **p** |
| --- | --- | --- | --- |
| Breusch-Pagan | 5.302 | 3 | .151 |

***Supplemental Table 237.*** *Breusch-Pagan test values for T1w+FLAIR WML segmentation.*

#### 76.4. Normal Q-Q plot

Supplemental Figure 155. Normal distribution quantile-quantile plot for T1w+FLAIR WML segmentation.

#### 76.5. Residuals vs leverage

Supplemental Figure 156. Residuals vs leverage diagnostics for T1w+FLAIR WML segmentation.

### 77. Periventricular | Female only | T1w+T2w WML vol | SDNN.LF._log | Exploratory restricted interaction model

Model: T1w+T2w WML vol ~ SDNN.LF._log + Age +	Age * SDNN.LF._log

#### 77.1. Standardized coefficients

| **Term** | **Estimate** | **Std. Error** | **t** | **p** | **β** |
| --- | --- | --- | --- | --- | --- |
| (Intercept) | -2.222e-04 | 9.284e-05 | -2.39 | .022 |  |
| SDNN.LF._log | 2.956e-04 | 2.698e-04 | 1.10 | .280 | 0.190 |
| Age | 1.104e-04 | 3.476e-05 | 3.18 | .003 | 0.427 |
| SDNN.LF._log:Age | -2.589e-04 | 7.398e-05 | -3.50 | .001 | -0.610 |

***Supplemental Table 238.*** *Standardized coefficients values for T1w+T2w WML segmentation.*

#### 77.2. 95% confidence intervals

| **Term** | **Lower 95%** | **Upper 95%** |
| --- | --- | --- |
| (Intercept) | -4.103e-04 | -3.404e-05 |
| SDNN.LF._log | -2.511e-04 | 8.424e-04 |
| Age | 3.994e-05 | 1.808e-04 |
| SDNN.LF._log:Age | -4.088e-04 | -1.091e-04 |

***Supplemental Table 239.*** *95% confidence intervals values for T1w+T2w WML segmentation.*

#### 77.3. Breusch-Pagan test

| **Test** | **Statistic** | **df** | **p** |
| --- | --- | --- | --- |
| Breusch-Pagan | 5.439 | 3 | .142 |

***Supplemental Table 240.*** *Breusch-Pagan test values for T1w+T2w WML segmentation.*

#### 77.4. Normal Q-Q plot

Supplemental Figure 157. Normal distribution quantile-quantile plot for T1w+T2w WML segmentation.

#### 77.5. Residuals vs leverage

Supplemental Figure 158. Residuals vs leverage diagnostics for T1w+T2w WML segmentation.

### 78. Periventricular | Female only | T1w-alone WML vol | SDNN.LF._log | Exploratory restricted interaction model

Model: T1w-alone WML vol ~ SDNN.LF._log + Age +	Age * SDNN.LF._log

#### 78.1. Standardized coefficients

| **Term** | **Estimate** | **Std. Error** | **t** | **p** | **β** |
| --- | --- | --- | --- | --- | --- |
| (Intercept) | -2.083e-04 | 8.967e-05 | -2.32 | .026 |  |
| SDNN.LF._log | 2.976e-04 | 2.606e-04 | 1.14 | .261 | 0.202 |
| Age | 1.036e-04 | 3.357e-05 | 3.09 | .004 | 0.423 |
| SDNN.LF._log:Age | -2.375e-04 | 7.145e-05 | -3.32 | .002 | -0.591 |

***Supplemental Table 241.*** *Standardized coefficients values for T1w-alone WML segmentation.*

#### 78.2. 95% confidence intervals

| **Term** | **Lower 95%** | **Upper 95%** |
| --- | --- | --- |
| (Intercept) | -3.900e-04 | -2.664e-05 |
| SDNN.LF._log | -2.305e-04 | 8.256e-04 |
| Age | 3.558e-05 | 1.716e-04 |
| SDNN.LF._log:Age | -3.823e-04 | -9.273e-05 |

***Supplemental Table 242.*** *95% confidence intervals values for T1w-alone WML segmentation.*

#### 78.3. Breusch-Pagan test

| **Test** | **Statistic** | **df** | **p** |
| --- | --- | --- | --- |
| Breusch-Pagan | 2.66 | 3 | .447 |

***Supplemental Table 243.*** *Breusch-Pagan test values for T1w-alone WML segmentation.*

#### 78.4. Normal Q-Q plot

Supplemental Figure 159. Normal distribution quantile-quantile plot for T1w-alone WML segmentation.

#### 78.5. Residuals vs leverage

Supplemental Figure 160. Residuals vs leverage diagnostics for T1w-alone WML segmentation.

### 79. Periventricular | Female only | T1w+FLAIR WML vol | SDNN.HF._log | Exploratory restricted interaction model

Model: T1w+FLAIR WML vol ~ SDNN.HF._log + Age +	Age * SDNN.HF._log

#### 79.1. Standardized coefficients

| **Term** | **Estimate** | **Std. Error** | **t** | **p** | **β** |
| --- | --- | --- | --- | --- | --- |
| (Intercept) | 7.376e-05 | 1.121e-04 | 0.658 | .515 |  |
| SDNN.HF._log | 5.113e-04 | 3.331e-04 | 1.53 | .133 | 0.397 |
| Age | -8.834e-05 | 6.719e-05 | -1.31 | .197 | -0.419 |
| SDNN.HF._log:Age | -6.700e-04 | 2.533e-04 | -2.65 | .012 | -1.12 |

***Supplemental Table 244.*** *Standardized coefficients values for T1w+FLAIR WML segmentation.*

#### 79.2. 95% confidence intervals

| **Term** | **Lower 95%** | **Upper 95%** |
| --- | --- | --- |
| (Intercept) | -1.534e-04 | 3.010e-04 |
| SDNN.HF._log | -1.637e-04 | 0.00119 |
| Age | -2.245e-04 | 4.781e-05 |
| SDNN.HF._log:Age | -0.00118 | -1.569e-04 |

***Supplemental Table 245.*** *95% confidence intervals values for T1w+FLAIR WML segmentation.*

#### 79.3. Breusch-Pagan test

| **Test** | **Statistic** | **df** | **p** |
| --- | --- | --- | --- |
| Breusch-Pagan | 5.393 | 3 | .145 |

***Supplemental Table 246.*** *Breusch-Pagan test values for T1w+FLAIR WML segmentation.*

#### 79.4. Normal Q-Q plot

Supplemental Figure 161. Normal distribution quantile-quantile plot for T1w+FLAIR WML segmentation.

#### 79.5. Residuals vs leverage

Supplemental Figure 162. Residuals vs leverage diagnostics for T1w+FLAIR WML segmentation.

### 80. Periventricular | Female only | T1w+T2w WML vol | SDNN.HF._log | Exploratory restricted interaction model

Model: T1w+T2w WML vol ~ SDNN.HF._log + Age +	Age * SDNN.HF._log

#### 80.1. Standardized coefficients

| **Term** | **Estimate** | **Std. Error** | **t** | **p** | **β** |
| --- | --- | --- | --- | --- | --- |
| (Intercept) | 1.236e-04 | 1.402e-04 | 0.882 | .384 |  |
| SDNN.HF._log | 0.00101 | 4.164e-04 | 2.43 | .020 | 0.640 |
| Age | -1.332e-04 | 8.399e-05 | -1.59 | .121 | -0.515 |
| SDNN.HF._log:Age | -9.249e-04 | 3.166e-04 | -2.92 | .006 | -1.26 |

***Supplemental Table 247.*** *Standardized coefficients values for T1w+T2w WML segmentation.*

#### 80.2. 95% confidence intervals

| **Term** | **Lower 95%** | **Upper 95%** |
| --- | --- | --- |
| (Intercept) | -1.604e-04 | 4.076e-04 |
| SDNN.HF._log | 1.666e-04 | 0.00185 |
| Age | -3.034e-04 | 3.696e-05 |
| SDNN.HF._log:Age | -0.00157 | -2.835e-04 |

***Supplemental Table 248.*** *95% confidence intervals values for T1w+T2w WML segmentation.*

#### 80.3. Breusch-Pagan test

| **Test** | **Statistic** | **df** | **p** |
| --- | --- | --- | --- |
| Breusch-Pagan | 2.541 | 3 | .468 |

***Supplemental Table 249.*** *Breusch-Pagan test values for T1w+T2w WML segmentation.*

#### 80.4. Normal Q-Q plot

Supplemental Figure 163. Normal distribution quantile-quantile plot for T1w+T2w WML segmentation.

#### 80.5. Residuals vs leverage

Supplemental Figure 164. Residuals vs leverage diagnostics for T1w+T2w WML segmentation.

### 81. Periventricular | Female only | T1w-alone WML vol | SDNN.HF._log | Exploratory restricted interaction model

Model: T1w-alone WML vol ~ SDNN.HF._log + Age +	Age * SDNN.HF._log

#### 81.1. Standardized coefficients

| **Term** | **Estimate** | **Std. Error** | **t** | **p** | **β** |
| --- | --- | --- | --- | --- | --- |
| (Intercept) | 1.248e-04 | 1.324e-04 | 0.942 | .352 |  |
| SDNN.HF._log | 9.464e-04 | 3.934e-04 | 2.41 | .021 | 0.632 |
| Age | -1.313e-04 | 7.936e-05 | -1.65 | .106 | -0.536 |
| SDNN.HF._log:Age | -8.908e-04 | 2.991e-04 | -2.98 | .005 | -1.28 |

***Supplemental Table 250.*** *Standardized coefficients values for T1w-alone WML segmentation.*

#### 81.2. 95% confidence intervals

| **Term** | **Lower 95%** | **Upper 95%** |
| --- | --- | --- |
| (Intercept) | -1.436e-04 | 3.931e-04 |
| SDNN.HF._log | 1.492e-04 | 0.00174 |
| Age | -2.921e-04 | 2.950e-05 |
| SDNN.HF._log:Age | -0.00150 | -2.847e-04 |

***Supplemental Table 251.*** *95% confidence intervals values for T1w-alone WML segmentation.*

#### 81.3. Breusch-Pagan test

| **Test** | **Statistic** | **df** | **p** |
| --- | --- | --- | --- |
| Breusch-Pagan | 0.739 | 3 | .864 |

***Supplemental Table 252.*** *Breusch-Pagan test values for T1w-alone WML segmentation.*

#### 81.4. Normal Q-Q plot

Supplemental Figure 165. Normal distribution quantile-quantile plot for T1w-alone WML segmentation.

#### 81.5. Residuals vs leverage

Supplemental Figure 166. Residuals vs leverage diagnostics for T1w-alone WML segmentation.

### 82. Periventricular | Male only | T1w+FLAIR WML vol | RMSSD | Exploratory restricted interaction model

Model: T1w+FLAIR WML vol ~ RMSSD + Age +	Age * RMSSD

#### 82.1. Standardized coefficients

| **Term** | **Estimate** | **Std. Error** | **t** | **p** | **β** |
| --- | --- | --- | --- | --- | --- |
| (Intercept) | -6.415e-04 | 9.381e-05 | -6.84 | <.001 |  |
| RMSSD | 1.667e-06 | 2.333e-06 | 0.715 | .477 | 0.0880 |
| Age | 1.466e-04 | 1.667e-05 | 8.79 | <.001 | 0.667 |
| RMSSD:Age | -6.495e-07 | 3.269e-07 | -1.99 | .050 | -0.243 |

***Supplemental Table 253.*** *Standardized coefficients values for T1w+FLAIR WML segmentation.*

#### 82.2. 95% confidence intervals

| **Term** | **Lower 95%** | **Upper 95%** |
| --- | --- | --- |
| (Intercept) | -8.279e-04 | -4.551e-04 |
| RMSSD | -2.968e-06 | 6.302e-06 |
| Age | 1.135e-04 | 1.797e-04 |
| RMSSD:Age | -1.299e-06 | 1.160e-10 |

***Supplemental Table 254.*** *95% confidence intervals values for T1w+FLAIR WML segmentation.*

#### 82.3. Breusch-Pagan test

| **Test** | **Statistic** | **df** | **p** |
| --- | --- | --- | --- |
| Breusch-Pagan | 11.215 | 3 | .011 |

***Supplemental Table 255.*** *Breusch-Pagan test values for T1w+FLAIR WML segmentation.*

#### 82.4. Normal Q-Q plot

Supplemental Figure 167. Normal distribution quantile-quantile plot for T1w+FLAIR WML segmentation.

#### 82.5. Residuals vs leverage

Supplemental Figure 168. Residuals vs leverage diagnostics for T1w+FLAIR WML segmentation.

### 83. Periventricular | Male only | T1w+T2w WML vol | RMSSD | Exploratory restricted interaction model

Model: T1w+T2w WML vol ~ RMSSD + Age +	Age * RMSSD

#### 83.1. Standardized coefficients

| **Term** | **Estimate** | **Std. Error** | **t** | **p** | **β** |
| --- | --- | --- | --- | --- | --- |
| (Intercept) | -7.641e-04 | 1.008e-04 | -7.58 | <.001 |  |
| RMSSD | 3.027e-06 | 2.508e-06 | 1.21 | .231 | 0.143 |
| Age | 1.753e-04 | 1.792e-05 | 9.78 | <.001 | 0.714 |
| RMSSD:Age | -7.059e-07 | 3.514e-07 | -2.01 | .048 | -0.236 |

***Supplemental Table 256.*** *Standardized coefficients values for T1w+T2w WML segmentation.*

#### 83.2. 95% confidence intervals

| **Term** | **Lower 95%** | **Upper 95%** |
| --- | --- | --- |
| (Intercept) | -9.644e-04 | -5.637e-04 |
| RMSSD | -1.956e-06 | 8.009e-06 |
| Age | 1.397e-04 | 2.109e-04 |
| RMSSD:Age | -1.404e-06 | -7.581e-09 |

***Supplemental Table 257.*** *95% confidence intervals values for T1w+T2w WML segmentation.*

#### 83.3. Breusch-Pagan test

| **Test** | **Statistic** | **df** | **p** |
| --- | --- | --- | --- |
| Breusch-Pagan | 10.667 | 3 | .014 |

***Supplemental Table 258.*** *Breusch-Pagan test values for T1w+T2w WML segmentation.*

#### 83.4. Normal Q-Q plot

Supplemental Figure 169. Normal distribution quantile-quantile plot for T1w+T2w WML segmentation.

#### 83.5. Residuals vs leverage

Supplemental Figure 170. Residuals vs leverage diagnostics for T1w+T2w WML segmentation.

### 84. Periventricular | Male only | T1w-alone WML vol | RMSSD | Exploratory restricted interaction model

Model: T1w-alone WML vol ~ RMSSD + Age + Age *	RMSSD

#### 84.1. Standardized coefficients

| **Term** | **Estimate** | **Std. Error** | **t** | **p** | **β** |
| --- | --- | --- | --- | --- | --- |
| (Intercept) | -6.502e-04 | 8.592e-05 | -7.57 | <.001 |  |
| RMSSD | 2.183e-06 | 2.137e-06 | 1.02 | .310 | 0.121 |
| Age | 1.497e-04 | 1.527e-05 | 9.81 | <.001 | 0.715 |
| RMSSD:Age | -5.518e-07 | 2.994e-07 | -1.84 | .069 | -0.217 |

***Supplemental Table 259.*** *Standardized coefficients values for T1w-alone WML segmentation.*

#### 84.2. 95% confidence intervals

| **Term** | **Lower 95%** | **Upper 95%** |
| --- | --- | --- |
| (Intercept) | -8.209e-04 | -4.795e-04 |
| RMSSD | -2.062e-06 | 6.429e-06 |
| Age | 1.194e-04 | 1.800e-04 |
| RMSSD:Age | -1.147e-06 | 4.311e-08 |

***Supplemental Table 260.*** *95% confidence intervals values for T1w-alone WML segmentation.*

#### 84.3. Breusch-Pagan test

| **Test** | **Statistic** | **df** | **p** |
| --- | --- | --- | --- |
| Breusch-Pagan | 13.262 | 3 | .004 |

***Supplemental Table 261.*** *Breusch-Pagan test values for T1w-alone WML segmentation.*

#### 84.4. Normal Q-Q plot

Supplemental Figure 171. Normal distribution quantile-quantile plot for T1w-alone WML segmentation.

#### 84.5. Residuals vs leverage

Supplemental Figure 172. Residuals vs leverage diagnostics for T1w-alone WML segmentation.

### 85. Periventricular | Male only | T1w+FLAIR WML vol | SDNN.LF._log | Exploratory restricted interaction model

Model: T1w+FLAIR WML vol ~ SDNN.LF._log + Age +	Age * SDNN.LF._log

#### 85.1. Standardized coefficients

| **Term** | **Estimate** | **Std. Error** | **t** | **p** | **β** |
| --- | --- | --- | --- | --- | --- |
| (Intercept) | -6.564e-04 | 9.481e-05 | -6.92 | <.001 |  |
| SDNN.LF._log | 3.210e-04 | 2.652e-04 | 1.21 | .229 | 0.162 |
| Age | 1.513e-04 | 1.681e-05 | 9.00 | <.001 | 0.688 |
| SDNN.LF._log:Age | -6.154e-05 | 3.538e-05 | -1.74 | .085 | -0.232 |

***Supplemental Table 262.*** *Standardized coefficients values for T1w+FLAIR WML segmentation.*

#### 85.2. 95% confidence intervals

| **Term** | **Lower 95%** | **Upper 95%** |
| --- | --- | --- |
| (Intercept) | -8.448e-04 | -4.680e-04 |
| SDNN.LF._log | -2.059e-04 | 8.479e-04 |
| Age | 1.179e-04 | 1.847e-04 |
| SDNN.LF._log:Age | -1.318e-04 | 8.761e-06 |

***Supplemental Table 263.*** *95% confidence intervals values for T1w+FLAIR WML segmentation.*

#### 85.3. Breusch-Pagan test

| **Test** | **Statistic** | **df** | **p** |
| --- | --- | --- | --- |
| Breusch-Pagan | 10.186 | 3 | .017 |

***Supplemental Table 264.*** *Breusch-Pagan test values for T1w+FLAIR WML segmentation.*

#### 85.4. Normal Q-Q plot

Supplemental Figure 173. Normal distribution quantile-quantile plot for T1w+FLAIR WML segmentation.

#### 85.5. Residuals vs leverage

Supplemental Figure 174. Residuals vs leverage diagnostics for T1w+FLAIR WML segmentation.

### 86. Periventricular | Male only | T1w+T2w WML vol | SDNN.LF._log | Exploratory restricted interaction model

Model: T1w+T2w WML vol ~ SDNN.LF._log + Age +	Age * SDNN.LF._log

#### 86.1. Standardized coefficients

| **Term** | **Estimate** | **Std. Error** | **t** | **p** | **β** |
| --- | --- | --- | --- | --- | --- |
| (Intercept) | -7.631e-04 | 1.016e-04 | -7.51 | <.001 |  |
| SDNN.LF._log | 2.967e-04 | 2.841e-04 | 1.04 | .299 | 0.134 |
| Age | 1.772e-04 | 1.801e-05 | 9.84 | <.001 | 0.722 |
| SDNN.LF._log:Age | -5.714e-05 | 3.790e-05 | -1.51 | .135 | -0.193 |

***Supplemental Table 265.*** *Standardized coefficients values for T1w+T2w WML segmentation.*

#### 86.2. 95% confidence intervals

| **Term** | **Lower 95%** | **Upper 95%** |
| --- | --- | --- |
| (Intercept) | -9.649e-04 | -5.613e-04 |
| SDNN.LF._log | -2.678e-04 | 8.611e-04 |
| Age | 1.414e-04 | 2.130e-04 |
| SDNN.LF._log:Age | -1.324e-04 | 1.816e-05 |

***Supplemental Table 266.*** *95% confidence intervals values for T1w+T2w WML segmentation.*

#### 86.3. Breusch-Pagan test

| **Test** | **Statistic** | **df** | **p** |
| --- | --- | --- | --- |
| Breusch-Pagan | 8.767 | 3 | .033 |

***Supplemental Table 267.*** *Breusch-Pagan test values for T1w+T2w WML segmentation.*

#### 86.4. Normal Q-Q plot

Supplemental Figure 175. Normal distribution quantile-quantile plot for T1w+T2w WML segmentation.

#### 86.5. Residuals vs leverage

Supplemental Figure 176. Residuals vs leverage diagnostics for T1w+T2w WML segmentation.

### 87. Periventricular | Male only | T1w-alone WML vol | SDNN.LF._log | Exploratory restricted interaction model

Model: T1w-alone WML vol ~ SDNN.LF._log + Age +	Age * SDNN.LF._log

#### 87.1. Standardized coefficients

| **Term** | **Estimate** | **Std. Error** | **t** | **p** | **β** |
| --- | --- | --- | --- | --- | --- |
| (Intercept) | -6.509e-04 | 8.616e-05 | -7.55 | <.001 |  |
| SDNN.LF._log | 2.420e-04 | 2.410e-04 | 1.00 | .318 | 0.128 |
| Age | 1.510e-04 | 1.528e-05 | 9.89 | <.001 | 0.721 |
| SDNN.LF._log:Age | -5.010e-05 | 3.215e-05 | -1.56 | .123 | -0.198 |

***Supplemental Table 268.*** *Standardized coefficients values for T1w-alone WML segmentation.*

#### 87.2. 95% confidence intervals

| **Term** | **Lower 95%** | **Upper 95%** |
| --- | --- | --- |
| (Intercept) | -8.221e-04 | -4.797e-04 |
| SDNN.LF._log | -2.368e-04 | 7.209e-04 |
| Age | 1.207e-04 | 1.814e-04 |
| SDNN.LF._log:Age | -1.140e-04 | 1.378e-05 |

***Supplemental Table 269.*** *95% confidence intervals values for T1w-alone WML segmentation.*

#### 87.3. Breusch-Pagan test

| **Test** | **Statistic** | **df** | **p** |
| --- | --- | --- | --- |
| Breusch-Pagan | 11.172 | 3 | .011 |

***Supplemental Table 270.*** *Breusch-Pagan test values for T1w-alone WML segmentation.*

#### 87.4. Normal Q-Q plot

Supplemental Figure 177. Normal distribution quantile-quantile plot for T1w-alone WML segmentation.

#### 87.5. Residuals vs leverage

Supplemental Figure 178. Residuals vs leverage diagnostics for T1w-alone WML segmentation.

### 88. Periventricular | Male only | T1w+FLAIR WML vol | SDNN.HF._log | Exploratory restricted interaction model

Model: T1w+FLAIR WML vol ~ SDNN.HF._log + Age +	Age * SDNN.HF._log

#### 88.1. Standardized coefficients

| **Term** | **Estimate** | **Std. Error** | **t** | **p** | **β** |
| --- | --- | --- | --- | --- | --- |
| (Intercept) | -6.369e-04 | 9.845e-05 | -6.47 | <.001 |  |
| SDNN.HF._log | 1.367e-04 | 2.878e-04 | 0.475 | .636 | 0.0664 |
| Age | 1.452e-04 | 1.772e-05 | 8.19 | <.001 | 0.661 |
| SDNN.HF._log:Age | -4.993e-05 | 3.784e-05 | -1.32 | .190 | -0.183 |

***Supplemental Table 271.*** *Standardized coefficients values for T1w+FLAIR WML segmentation.*

#### 88.2. 95% confidence intervals

| **Term** | **Lower 95%** | **Upper 95%** |
| --- | --- | --- |
| (Intercept) | -8.325e-04 | -4.413e-04 |
| SDNN.HF._log | -4.351e-04 | 7.086e-04 |
| Age | 1.100e-04 | 1.804e-04 |
| SDNN.HF._log:Age | -1.251e-04 | 2.527e-05 |

***Supplemental Table 272.*** *95% confidence intervals values for T1w+FLAIR WML segmentation.*

#### 88.3. Breusch-Pagan test

| **Test** | **Statistic** | **df** | **p** |
| --- | --- | --- | --- |
| Breusch-Pagan | 9.803 | 3 | .020 |

***Supplemental Table 273.*** *Breusch-Pagan test values for T1w+FLAIR WML segmentation.*

#### 88.4. Normal Q-Q plot

Supplemental Figure 179. Normal distribution quantile-quantile plot for T1w+FLAIR WML segmentation.

#### 88.5. Residuals vs leverage

Supplemental Figure 180. Residuals vs leverage diagnostics for T1w+FLAIR WML segmentation.

### 89. Periventricular | Male only | T1w+T2w WML vol | SDNN.HF._log | Exploratory restricted interaction model

Model: T1w+T2w WML vol ~ SDNN.HF._log + Age +	Age * SDNN.HF._log

#### 89.1. Standardized coefficients

| **Term** | **Estimate** | **Std. Error** | **t** | **p** | **β** |
| --- | --- | --- | --- | --- | --- |
| (Intercept) | -7.395e-04 | 1.049e-04 | -7.05 | <.001 |  |
| SDNN.HF._log | 1.141e-04 | 3.065e-04 | 0.372 | .710 | 0.0496 |
| Age | 1.694e-04 | 1.887e-05 | 8.98 | <.001 | 0.690 |
| SDNN.HF._log:Age | -5.224e-05 | 4.031e-05 | -1.30 | .198 | -0.171 |

***Supplemental Table 274.*** *Standardized coefficients values for T1w+T2w WML segmentation.*

#### 89.2. 95% confidence intervals

| **Term** | **Lower 95%** | **Upper 95%** |
| --- | --- | --- |
| (Intercept) | -9.478e-04 | -5.311e-04 |
| SDNN.HF._log | -4.949e-04 | 7.232e-04 |
| Age | 1.319e-04 | 2.069e-04 |
| SDNN.HF._log:Age | -1.323e-04 | 2.785e-05 |

***Supplemental Table 275.*** *95% confidence intervals values for T1w+T2w WML segmentation.*

#### 89.3. Breusch-Pagan test

| **Test** | **Statistic** | **df** | **p** |
| --- | --- | --- | --- |
| Breusch-Pagan | 8.53 | 3 | .036 |

***Supplemental Table 276.*** *Breusch-Pagan test values for T1w+T2w WML segmentation.*

#### 89.4. Normal Q-Q plot

Supplemental Figure 181. Normal distribution quantile-quantile plot for T1w+T2w WML segmentation.

#### 89.5. Residuals vs leverage

Supplemental Figure 182. Residuals vs leverage diagnostics for T1w+T2w WML segmentation.

### 90. Periventricular | Male only | T1w-alone WML vol | SDNN.HF._log | Exploratory restricted interaction model

Model: T1w-alone WML vol ~ SDNN.HF._log + Age +	Age * SDNN.HF._log

#### 90.1. Standardized coefficients

| **Term** | **Estimate** | **Std. Error** | **t** | **p** | **β** |
| --- | --- | --- | --- | --- | --- |
| (Intercept) | -6.330e-04 | 8.927e-05 | -7.09 | <.001 |  |
| SDNN.HF._log | 9.143e-05 | 2.609e-04 | 0.350 | .727 | 0.0466 |
| Age | 1.454e-04 | 1.606e-05 | 9.05 | <.001 | 0.695 |
| SDNN.HF._log:Age | -4.185e-05 | 3.431e-05 | -1.22 | .226 | -0.161 |

***Supplemental Table 277.*** *Standardized coefficients values for T1w-alone WML segmentation.*

#### 90.2. 95% confidence intervals

| **Term** | **Lower 95%** | **Upper 95%** |
| --- | --- | --- |
| (Intercept) | -8.104e-04 | -4.556e-04 |
| SDNN.HF._log | -4.271e-04 | 6.099e-04 |
| Age | 1.135e-04 | 1.773e-04 |
| SDNN.HF._log:Age | -1.100e-04 | 2.633e-05 |

***Supplemental Table 278.*** *95% confidence intervals values for T1w-alone WML segmentation.*

#### 90.3. Breusch-Pagan test

| **Test** | **Statistic** | **df** | **p** |
| --- | --- | --- | --- |
| Breusch-Pagan | 11.071 | 3 | .011 |

***Supplemental Table 279.*** *Breusch-Pagan test values for T1w-alone WML segmentation.*

#### 90.4. Normal Q-Q plot

Supplemental Figure 183. Normal distribution quantile-quantile plot for T1w-alone WML segmentation.

#### 90.5. Residuals vs leverage

Supplemental Figure 184. Residuals vs leverage diagnostics for T1w-alone WML segmentation.

### 91. Periventricular | Young only | T1w+FLAIR WML vol | RMSSD | Exploratory restricted interaction model

Model: T1w+FLAIR WML vol ~ RMSSD + Age +	Sex + Age * RMSSD + Sex * RMSSD

#### 91.1. Standardized coefficients

| **Term** | **Estimate** | **Std. Error** | **t** | **p** | **β** |
| --- | --- | --- | --- | --- | --- |
| (Intercept) | -5.469e-05 | 1.165e-04 | -0.469 | .640 |  |
| RMSSD | -1.808e-07 | 3.262e-06 | -0.0554 | .956 | -0.0225 |
| Age | 7.724e-05 | 4.460e-05 | 1.73 | .086 | 0.183 |
| Sex | -5.297e-05 | 6.216e-05 | -0.852 | .396 | -0.0877 |
| RMSSD:Age | -1.706e-06 | 1.596e-06 | -1.07 | .288 | -0.307 |
| RMSSD:Sex | 1.408e-06 | 1.771e-06 | 0.795 | .429 | 0.303 |

***Supplemental Table 280.*** *Standardized coefficients values for T1w+FLAIR WML segmentation.*

#### 91.2. 95% confidence intervals

| **Term** | **Lower 95%** | **Upper 95%** |
| --- | --- | --- |
| (Intercept) | -2.859e-04 | 1.765e-04 |
| RMSSD | -6.655e-06 | 6.293e-06 |
| Age | -1.127e-05 | 1.657e-04 |
| Sex | -1.763e-04 | 7.039e-05 |
| RMSSD:Age | -4.873e-06 | 1.461e-06 |
| RMSSD:Sex | -2.107e-06 | 4.923e-06 |

***Supplemental Table 281.*** *95% confidence intervals values for T1w+FLAIR WML segmentation.*

#### 91.3. Breusch-Pagan test

| **Test** | **Statistic** | **df** | **p** |
| --- | --- | --- | --- |
| Breusch-Pagan | 5.621 | 5 | .345 |

***Supplemental Table 282.*** *Breusch-Pagan test values for T1w+FLAIR WML segmentation.*

#### 91.4. Normal Q-Q plot

Supplemental Figure 185. Normal distribution quantile-quantile plot for T1w+FLAIR WML segmentation.

#### 91.5. Residuals vs leverage

Supplemental Figure 186. Residuals vs leverage diagnostics for T1w+FLAIR WML segmentation.

### 92. Periventricular | Young only | T1w+T2w WML vol | RMSSD | Exploratory restricted interaction model

Model: T1w+T2w WML vol ~ RMSSD + Age +	Sex + Age * RMSSD + Sex * RMSSD

#### 92.1. Standardized coefficients

| **Term** | **Estimate** | **Std. Error** | **t** | **p** | **β** |
| --- | --- | --- | --- | --- | --- |
| (Intercept) | 7.990e-05 | 1.321e-04 | 0.605 | .547 |  |
| RMSSD | 2.715e-06 | 3.698e-06 | 0.734 | .465 | 0.296 |
| Age | 7.980e-05 | 5.056e-05 | 1.58 | .118 | 0.165 |
| Sex | -1.374e-04 | 7.047e-05 | -1.95 | .054 | -0.199 |
| RMSSD:Age | -1.896e-06 | 1.809e-06 | -1.05 | .297 | -0.299 |
| RMSSD:Sex | 6.560e-07 | 2.008e-06 | 0.327 | .745 | 0.124 |

***Supplemental Table 283.*** *Standardized coefficients values for T1w+T2w WML segmentation.*

#### 92.2. 95% confidence intervals

| **Term** | **Lower 95%** | **Upper 95%** |
| --- | --- | --- |
| (Intercept) | -1.822e-04 | 3.420e-04 |
| RMSSD | -4.624e-06 | 1.005e-05 |
| Age | -2.053e-05 | 1.801e-04 |
| Sex | -2.773e-04 | 2.417e-06 |
| RMSSD:Age | -5.486e-06 | 1.694e-06 |
| RMSSD:Sex | -3.328e-06 | 4.641e-06 |

***Supplemental Table 284.*** *95% confidence intervals values for T1w+T2w WML segmentation.*

#### 92.3. Breusch-Pagan test

| **Test** | **Statistic** | **df** | **p** |
| --- | --- | --- | --- |
| Breusch-Pagan | 5.259 | 5 | .385 |

***Supplemental Table 285.*** *Breusch-Pagan test values for T1w+T2w WML segmentation.*

#### 92.4. Normal Q-Q plot

Supplemental Figure 187. Normal distribution quantile-quantile plot for T1w+T2w WML segmentation.

#### 92.5. Residuals vs leverage

Supplemental Figure 188. Residuals vs leverage diagnostics for T1w+T2w WML segmentation.

### 93. Periventricular | Young only | T1w-alone WML vol | RMSSD | Exploratory restricted interaction model

Model: T1w-alone WML vol ~ RMSSD + Age + Sex +	Age * RMSSD + Sex * RMSSD

#### 93.1. Standardized coefficients

| **Term** | **Estimate** | **Std. Error** | **t** | **p** | **β** |
| --- | --- | --- | --- | --- | --- |
| (Intercept) | 8.647e-05 | 1.210e-04 | 0.714 | .477 |  |
| RMSSD | 2.405e-06 | 3.389e-06 | 0.710 | .480 | 0.289 |
| Age | 5.374e-05 | 4.633e-05 | 1.16 | .249 | 0.123 |
| Sex | -1.134e-04 | 6.458e-05 | -1.76 | .082 | -0.181 |
| RMSSD:Age | -1.573e-06 | 1.658e-06 | -0.949 | .345 | -0.274 |
| RMSSD:Sex | 3.314e-07 | 1.840e-06 | 0.180 | .857 | 0.0690 |

***Supplemental Table 286.*** *Standardized coefficients values for T1w-alone WML segmentation.*

#### 93.2. 95% confidence intervals

| **Term** | **Lower 95%** | **Upper 95%** |
| --- | --- | --- |
| (Intercept) | -1.537e-04 | 3.267e-04 |
| RMSSD | -4.320e-06 | 9.131e-06 |
| Age | -3.820e-05 | 1.457e-04 |
| Sex | -2.415e-04 | 1.476e-05 |
| RMSSD:Age | -4.863e-06 | 1.717e-06 |
| RMSSD:Sex | -3.320e-06 | 3.983e-06 |

***Supplemental Table 287.*** *95% confidence intervals values for T1w-alone WML segmentation.*

#### 93.3. Breusch-Pagan test

| **Test** | **Statistic** | **df** | **p** |
| --- | --- | --- | --- |
| Breusch-Pagan | 4.307 | 5 | .506 |

***Supplemental Table 288.*** *Breusch-Pagan test values for T1w-alone WML segmentation.*

#### 93.4. Normal Q-Q plot

Supplemental Figure 189. Normal distribution quantile-quantile plot for T1w-alone WML segmentation.

#### 93.5. Residuals vs leverage

Supplemental Figure 190. Residuals vs leverage diagnostics for T1w-alone WML segmentation.

### 94. Periventricular | Young only | T1w+FLAIR WML vol | SDNN.LF._log | Exploratory restricted interaction model

Model: T1w+FLAIR WML vol ~ SDNN.LF._log + Age + Sex + Age * SDNN.LF._log + Sex * SDNN.LF._log

#### 94.1. Standardized coefficients

| **Term** | **Estimate** | **Std. Error** | **t** | **p** | **β** |
| --- | --- | --- | --- | --- | --- |
| (Intercept) | -5.199e-05 | 1.131e-04 | -0.460 | .647 |  |
| SDNN.LF._log | -7.760e-04 | 3.551e-04 | -2.19 | .031 | -0.852 |
| Age | 9.768e-05 | 4.257e-05 | 2.29 | .024 | 0.231 |
| Sex | -6.638e-05 | 5.818e-05 | -1.14 | .257 | -0.110 |
| SDNN.LF._log:Age | 5.641e-05 | 1.581e-04 | 0.357 | .722 | 0.0964 |
| SDNN.LF._log:Sex | 4.274e-04 | 1.933e-04 | 2.21 | .029 | 0.819 |

***Supplemental Table 289.*** *Standardized coefficients values for T1w+FLAIR WML segmentation.*

#### 94.2. 95% confidence intervals

| **Term** | **Lower 95%** | **Upper 95%** |
| --- | --- | --- |
| (Intercept) | -2.765e-04 | 1.725e-04 |
| SDNN.LF._log | -0.00148 | -7.130e-05 |
| Age | 1.321e-05 | 1.822e-04 |
| Sex | -1.818e-04 | 4.908e-05 |
| SDNN.LF._log:Age | -2.574e-04 | 3.703e-04 |
| SDNN.LF._log:Sex | 4.380e-05 | 8.110e-04 |

***Supplemental Table 290.*** *95% confidence intervals values for T1w+FLAIR WML segmentation.*

#### 94.3. Breusch-Pagan test

| **Test** | **Statistic** | **df** | **p** |
| --- | --- | --- | --- |
| Breusch-Pagan | 18.083 | 5 | .003 |

***Supplemental Table 291.*** *Breusch-Pagan test values for T1w+FLAIR WML segmentation.*

#### 94.4. Normal Q-Q plot

Supplemental Figure 191. Normal distribution quantile-quantile plot for T1w+FLAIR WML segmentation.

#### 94.5. Residuals vs leverage

Supplemental Figure 192. Residuals vs leverage diagnostics for T1w+FLAIR WML segmentation.

### 95. Periventricular | Young only | T1w+T2w WML vol | SDNN.LF._log | Exploratory restricted interaction model

Model: T1w+T2w WML vol ~ SDNN.LF._log + Age + Sex + Age * SDNN.LF._log + Sex * SDNN.LF._log

#### 95.1. Standardized coefficients

| **Term** | **Estimate** | **Std. Error** | **t** | **p** | **β** |
| --- | --- | --- | --- | --- | --- |
| (Intercept) | 5.662e-05 | 1.314e-04 | 0.431 | .667 |  |
| SDNN.LF._log | -3.559e-04 | 4.125e-04 | -0.863 | .390 | -0.342 |
| Age | 9.997e-05 | 4.945e-05 | 2.02 | .046 | 0.207 |
| Sex | -1.300e-04 | 6.758e-05 | -1.92 | .057 | -0.188 |
| SDNN.LF._log:Age | 2.044e-04 | 1.837e-04 | 1.11 | .269 | 0.305 |
| SDNN.LF._log:Sex | 9.761e-05 | 2.245e-04 | 0.435 | .665 | 0.164 |

***Supplemental Table 292.*** *Standardized coefficients values for T1w+T2w WML segmentation.*

#### 95.2. 95% confidence intervals

| **Term** | **Lower 95%** | **Upper 95%** |
| --- | --- | --- |
| (Intercept) | -2.041e-04 | 3.174e-04 |
| SDNN.LF._log | -0.00117 | 4.626e-04 |
| Age | 1.843e-06 | 1.981e-04 |
| Sex | -2.641e-04 | 4.107e-06 |
| SDNN.LF._log:Age | -1.602e-04 | 5.689e-04 |
| SDNN.LF._log:Sex | -3.480e-04 | 5.432e-04 |

***Supplemental Table 293.*** *95% confidence intervals values for T1w+T2w WML segmentation.*

#### 95.3. Breusch-Pagan test

| **Test** | **Statistic** | **df** | **p** |
| --- | --- | --- | --- |
| Breusch-Pagan | 9.345 | 5 | .096 |

***Supplemental Table 294.*** *Breusch-Pagan test values for T1w+T2w WML segmentation.*

#### 95.4. Normal Q-Q plot

Supplemental Figure 193. Normal distribution quantile-quantile plot for T1w+T2w WML segmentation.

#### 95.5. Residuals vs leverage

Supplemental Figure 194. Residuals vs leverage diagnostics for T1w+T2w WML segmentation.

### 96. Periventricular | Young only | T1w-alone WML vol | SDNN.LF._log | Exploratory restricted interaction model

Model: T1w-alone WML vol ~ SDNN.LF._log + Age + Sex + Age * SDNN.LF._log + Sex * SDNN.LF._log

#### 96.1. Standardized coefficients

| **Term** | **Estimate** | **Std. Error** | **t** | **p** | **β** |
| --- | --- | --- | --- | --- | --- |
| (Intercept) | 6.833e-05 | 1.203e-04 | 0.568 | .571 |  |
| SDNN.LF._log | -2.382e-04 | 3.777e-04 | -0.631 | .530 | -0.253 |
| Age | 7.085e-05 | 4.528e-05 | 1.56 | .121 | 0.162 |
| Sex | -1.096e-04 | 6.189e-05 | -1.77 | .080 | -0.175 |
| SDNN.LF._log:Age | 1.407e-04 | 1.682e-04 | 0.836 | .405 | 0.232 |
| SDNN.LF._log:Sex | 6.849e-05 | 2.056e-04 | 0.333 | .740 | 0.127 |

***Supplemental Table 295.*** *Standardized coefficients values for T1w-alone WML segmentation.*

#### 96.2. 95% confidence intervals

| **Term** | **Lower 95%** | **Upper 95%** |
| --- | --- | --- |
| (Intercept) | -1.705e-04 | 3.071e-04 |
| SDNN.LF._log | -9.878e-04 | 5.114e-04 |
| Age | -1.902e-05 | 1.607e-04 |
| Sex | -2.324e-04 | 1.321e-05 |
| SDNN.LF._log:Age | -1.932e-04 | 4.745e-04 |
| SDNN.LF._log:Sex | -3.396e-04 | 4.766e-04 |

***Supplemental Table 296.*** *95% confidence intervals values for T1w-alone WML segmentation.*

#### 96.3. Breusch-Pagan test

| **Test** | **Statistic** | **df** | **p** |
| --- | --- | --- | --- |
| Breusch-Pagan | 7.412 | 5 | .192 |

***Supplemental Table 297.*** *Breusch-Pagan test values for T1w-alone WML segmentation.*

#### 96.4. Normal Q-Q plot

Supplemental Figure 195. Normal distribution quantile-quantile plot for T1w-alone WML segmentation.

#### 96.5. Residuals vs leverage

Supplemental Figure 196. Residuals vs leverage diagnostics for T1w-alone WML segmentation.

### 97. Periventricular | Young only | T1w+FLAIR WML vol | SDNN.HF._log | Exploratory restricted interaction model

Model: T1w+FLAIR WML vol ~ SDNN.HF._log + Age + Sex + Age * SDNN.HF._log + Sex * SDNN.HF._log

#### 97.1. Standardized coefficients

| **Term** | **Estimate** | **Std. Error** | **t** | **p** | **β** |
| --- | --- | --- | --- | --- | --- |
| (Intercept) | -4.087e-05 | 1.114e-04 | -0.367 | .715 |  |
| SDNN.HF._log | 6.989e-05 | 3.450e-04 | 0.203 | .840 | 0.0742 |
| Age | 4.654e-05 | 4.336e-05 | 1.07 | .286 | 0.110 |
| Sex | -4.105e-05 | 5.840e-05 | -0.703 | .484 | -0.0679 |
| SDNN.HF._log:Age | -4.933e-04 | 1.758e-04 | -2.81 | .006 | -0.775 |
| SDNN.HF._log:Sex | 3.107e-04 | 1.921e-04 | 1.62 | .109 | 0.557 |

***Supplemental Table 298.*** *Standardized coefficients values for T1w+FLAIR WML segmentation.*

#### 97.2. 95% confidence intervals

| **Term** | **Lower 95%** | **Upper 95%** |
| --- | --- | --- |
| (Intercept) | -2.620e-04 | 1.803e-04 |
| SDNN.HF._log | -6.148e-04 | 7.545e-04 |
| Age | -3.951e-05 | 1.326e-04 |
| Sex | -1.569e-04 | 7.484e-05 |
| SDNN.HF._log:Age | -8.423e-04 | -1.444e-04 |
| SDNN.HF._log:Sex | -7.058e-05 | 6.919e-04 |

***Supplemental Table 299.*** *95% confidence intervals values for T1w+FLAIR WML segmentation.*

#### 97.3. Breusch-Pagan test

| **Test** | **Statistic** | **df** | **p** |
| --- | --- | --- | --- |
| Breusch-Pagan | 9.061 | 5 | .107 |

***Supplemental Table 300.*** *Breusch-Pagan test values for T1w+FLAIR WML segmentation.*

#### 97.4. Normal Q-Q plot

Supplemental Figure 197. Normal distribution quantile-quantile plot for T1w+FLAIR WML segmentation.

#### 97.5. Residuals vs leverage

Supplemental Figure 198. Residuals vs leverage diagnostics for T1w+FLAIR WML segmentation.

### 98. Periventricular | Young only | T1w+T2w WML vol | SDNN.HF._log | Exploratory restricted interaction model

Model: T1w+T2w WML vol ~ SDNN.HF._log + Age + Sex + Age * SDNN.HF._log + Sex * SDNN.HF._log

#### 98.1. Standardized coefficients

| **Term** | **Estimate** | **Std. Error** | **t** | **p** | **β** |
| --- | --- | --- | --- | --- | --- |
| (Intercept) | 9.297e-05 | 1.315e-04 | 0.707 | .481 |  |
| SDNN.HF._log | 4.554e-04 | 4.071e-04 | 1.12 | .266 | 0.423 |
| Age | 5.097e-05 | 5.117e-05 | 0.996 | .322 | 0.106 |
| Sex | -1.187e-04 | 6.891e-05 | -1.72 | .088 | -0.172 |
| SDNN.HF._log:Age | -3.346e-04 | 2.075e-04 | -1.61 | .110 | -0.460 |
| SDNN.HF._log:Sex | -1.314e-05 | 2.267e-04 | -0.0580 | .954 | -0.0206 |

***Supplemental Table 301.*** *Standardized coefficients values for T1w+T2w WML segmentation.*

#### 98.2. 95% confidence intervals

| **Term** | **Lower 95%** | **Upper 95%** |
| --- | --- | --- |
| (Intercept) | -1.680e-04 | 3.539e-04 |
| SDNN.HF._log | -3.524e-04 | 0.00126 |
| Age | -5.057e-05 | 1.525e-04 |
| Sex | -2.555e-04 | 1.801e-05 |
| SDNN.HF._log:Age | -7.463e-04 | 7.718e-05 |
| SDNN.HF._log:Sex | -4.630e-04 | 4.367e-04 |

***Supplemental Table 302.*** *95% confidence intervals values for T1w+T2w WML segmentation.*

#### 98.3. Breusch-Pagan test

| **Test** | **Statistic** | **df** | **p** |
| --- | --- | --- | --- |
| Breusch-Pagan | 6.015 | 5 | .305 |

***Supplemental Table 303.*** *Breusch-Pagan test values for T1w+T2w WML segmentation.*

#### 98.4. Normal Q-Q plot

Supplemental Figure 199. Normal distribution quantile-quantile plot for T1w+T2w WML segmentation.

#### 98.5. Residuals vs leverage

Supplemental Figure 200. Residuals vs leverage diagnostics for T1w+T2w WML segmentation.

### 99. Periventricular | Young only | T1w-alone WML vol | SDNN.HF._log | Exploratory restricted interaction model

Model: T1w-alone WML vol ~ SDNN.HF._log + Age + Sex + Age * SDNN.HF._log + Sex * SDNN.HF._log

#### 99.1. Standardized coefficients

| **Term** | **Estimate** | **Std. Error** | **t** | **p** | **β** |
| --- | --- | --- | --- | --- | --- |
| (Intercept) | 9.308e-05 | 1.208e-04 | 0.771 | .443 |  |
| SDNN.HF._log | 2.618e-04 | 3.740e-04 | 0.700 | .486 | 0.269 |
| Age | 3.760e-05 | 4.701e-05 | 0.800 | .426 | 0.0860 |
| Sex | -1.012e-04 | 6.331e-05 | -1.60 | .113 | -0.162 |
| SDNN.HF._log:Age | -1.979e-04 | 1.906e-04 | -1.04 | .302 | -0.300 |
| SDNN.HF._log:Sex | -9.657e-06 | 2.083e-04 | -0.0464 | .963 | -0.0167 |

***Supplemental Table 304.*** *Standardized coefficients values for T1w-alone WML segmentation.*

#### 99.2. 95% confidence intervals

| **Term** | **Lower 95%** | **Upper 95%** |
| --- | --- | --- |
| (Intercept) | -1.466e-04 | 3.328e-04 |
| SDNN.HF._log | -4.804e-04 | 0.00100 |
| Age | -5.569e-05 | 1.309e-04 |
| Sex | -2.268e-04 | 2.446e-05 |
| SDNN.HF._log:Age | -5.762e-04 | 1.804e-04 |
| SDNN.HF._log:Sex | -4.230e-04 | 4.037e-04 |

***Supplemental Table 305.*** *95% confidence intervals values for T1w-alone WML segmentation.*

#### 99.3. Breusch-Pagan test

| **Test** | **Statistic** | **df** | **p** |
| --- | --- | --- | --- |
| Breusch-Pagan | 4.649 | 5 | .460 |

***Supplemental Table 306.*** *Breusch-Pagan test values for T1w-alone WML segmentation.*

#### 99.4. Normal Q-Q plot

Supplemental Figure 201. Normal distribution quantile-quantile plot for T1w-alone WML segmentation.

#### 99.5. Residuals vs leverage

Supplemental Figure 202. Residuals vs leverage diagnostics for T1w-alone WML segmentation.

### 100. Periventricular | Old only | T1w+FLAIR WML vol | RMSSD | Exploratory restricted interaction model

Model: T1w+FLAIR WML vol ~ RMSSD + Age +	Sex + Age * RMSSD + Sex * RMSSD

#### 100.1. Standardized coefficients

| **Term** | **Estimate** | **Std. Error** | **t** | **p** | **β** |
| --- | --- | --- | --- | --- | --- |
| (Intercept) | 9.037e-05 | 0.00353 | 0.0256 | .980 |  |
| RMSSD | 3.579e-04 | 2.448e-04 | 1.46 | .157 | 18.1 |
| Age | 5.188e-04 | 2.478e-04 | 2.09 | .047 | 0.449 |
| Sex | -0.00261 | 0.00188 | -1.39 | .178 | -0.621 |
| RMSSD:Age | 2.902e-06 | 9.010e-06 | 0.322 | .750 | 1.49 |
| RMSSD:Sex | -1.964e-04 | 1.121e-04 | -1.75 | .093 | -19.8 |

***Supplemental Table 307.*** *Standardized coefficients values for T1w+FLAIR WML segmentation.*

#### 100.2. 95% confidence intervals

| **Term** | **Lower 95%** | **Upper 95%** |
| --- | --- | --- |
| (Intercept) | -0.00720 | 0.00738 |
| RMSSD | -1.474e-04 | 8.633e-04 |
| Age | 7.276e-06 | 0.00103 |
| Sex | -0.00649 | 0.00127 |
| RMSSD:Age | -1.569e-05 | 2.150e-05 |
| RMSSD:Sex | -4.278e-04 | 3.499e-05 |

***Supplemental Table 308.*** *95% confidence intervals values for T1w+FLAIR WML segmentation.*

#### 100.3. Breusch-Pagan test

| **Test** | **Statistic** | **df** | **p** |
| --- | --- | --- | --- |
| Breusch-Pagan | 4.881 | 5 | .431 |

***Supplemental Table 309.*** *Breusch-Pagan test values for T1w+FLAIR WML segmentation.*

#### 100.4. Normal Q-Q plot

Supplemental Figure 203. Normal distribution quantile-quantile plot for T1w+FLAIR WML segmentation.

#### 100.5. Residuals vs leverage

Supplemental Figure 204. Residuals vs leverage diagnostics for T1w+FLAIR WML segmentation. The 2 older female subjects are not visualized due to possessing leverage values approaching 1 in this restricted model, preventing the normal calculation of their standardized residuals.

### 101. Periventricular | Old only | T1w+T2w WML vol | RMSSD | Exploratory restricted interaction model

Model: T1w+T2w WML vol ~ RMSSD + Age +	Sex + Age * RMSSD + Sex * RMSSD

#### 101.1. Standardized coefficients

| **Term** | **Estimate** | **Std. Error** | **t** | **p** | **β** |
| --- | --- | --- | --- | --- | --- |
| (Intercept) | 0.00185 | 0.00383 | 0.482 | .634 |  |
| RMSSD | 4.931e-04 | 2.655e-04 | 1.86 | .076 | 23.2 |
| Age | 5.078e-04 | 2.688e-04 | 1.89 | .071 | 0.409 |
| Sex | -0.00343 | 0.00204 | -1.68 | .106 | -0.759 |
| RMSSD:Age | 1.093e-07 | 9.771e-06 | 0.0112 | .991 | 0.0521 |
| RMSSD:Sex | -2.494e-04 | 1.216e-04 | -2.05 | .051 | -23.4 |

***Supplemental Table 310.*** *Standardized coefficients values for T1w+T2w WML segmentation.*

#### 101.2. 95% confidence intervals

| **Term** | **Lower 95%** | **Upper 95%** |
| --- | --- | --- |
| (Intercept) | -0.00606 | 0.00975 |
| RMSSD | -5.489e-05 | 0.00104 |
| Age | -4.687e-05 | 0.00106 |
| Sex | -0.00764 | 7.854e-04 |
| RMSSD:Age | -2.006e-05 | 2.028e-05 |
| RMSSD:Sex | -5.004e-04 | 1.555e-06 |

***Supplemental Table 311.*** *95% confidence intervals values for T1w+T2w WML segmentation.*

#### 101.3. Breusch-Pagan test

| **Test** | **Statistic** | **df** | **p** |
| --- | --- | --- | --- |
| Breusch-Pagan | 3.906 | 5 | .563 |

***Supplemental Table 312.*** *Breusch-Pagan test values for T1w+T2w WML segmentation.*

#### 101.4. Normal Q-Q plot

Supplemental Figure 205. Normal distribution quantile-quantile plot for T1w+T2w WML segmentation.

#### 101.5. Residuals vs leverage

Supplemental Figure 206. Residuals vs leverage diagnostics for T1w+T2w WML segmentation. The 2 older female subjects are not visualized due to possessing leverage values approaching 1 in this restricted model, preventing the normal calculation of their standardized residuals.

### 102. Periventricular | Old only | T1w-alone WML vol | RMSSD | Exploratory restricted interaction model

Model: T1w-alone WML vol ~ RMSSD + Age + Sex +	Age * RMSSD + Sex * RMSSD

#### 102.1. Standardized coefficients

| **Term** | **Estimate** | **Std. Error** | **t** | **p** | **β** |
| --- | --- | --- | --- | --- | --- |
| (Intercept) | 0.00169 | 0.00329 | 0.513 | .613 |  |
| RMSSD | 3.669e-04 | 2.282e-04 | 1.61 | .121 | 20.4 |
| Age | 4.179e-04 | 2.309e-04 | 1.81 | .083 | 0.398 |
| Sex | -0.00292 | 0.00175 | -1.66 | .109 | -0.763 |
| RMSSD:Age | 3.567e-06 | 8.396e-06 | 0.425 | .675 | 2.01 |
| RMSSD:Sex | -2.034e-04 | 1.045e-04 | -1.95 | .063 | -22.5 |

***Supplemental Table 313.*** *Standardized coefficients values for T1w-alone WML segmentation.*

#### 102.2. 95% confidence intervals

| **Term** | **Lower 95%** | **Upper 95%** |
| --- | --- | --- |
| (Intercept) | -0.00510 | 0.00848 |
| RMSSD | -1.040e-04 | 8.378e-04 |
| Age | -5.874e-05 | 8.945e-04 |
| Sex | -0.00654 | 7.024e-04 |
| RMSSD:Age | -1.376e-05 | 2.089e-05 |
| RMSSD:Sex | -4.191e-04 | 1.219e-05 |

***Supplemental Table 314.*** *95% confidence intervals values for T1w-alone WML segmentation.*

#### 102.3. Breusch-Pagan test

| **Test** | **Statistic** | **df** | **p** |
| --- | --- | --- | --- |
| Breusch-Pagan | 4.144 | 5 | .529 |

***Supplemental Table 315.*** *Breusch-Pagan test values for T1w-alone WML segmentation.*

#### 102.4. Normal Q-Q plot

Supplemental Figure 207. Normal distribution quantile-quantile plot for T1w-alone WML segmentation.

#### 102.5. Residuals vs leverage

Supplemental Figure 208. Residuals vs leverage diagnostics for T1w-alone WML segmentation. The 2 older female subjects are not visualized due to possessing leverage values approaching 1 in this restricted model, preventing the normal calculation of their standardized residuals.

### 103. Periventricular | Old only | T1w+FLAIR WML vol | SDNN.LF._log | Exploratory restricted interaction model

Model: T1w+FLAIR WML vol ~ SDNN.LF._log + Age + Sex + Age * SDNN.LF._log + Sex * SDNN.LF._log

#### 103.1. Standardized coefficients

| **Term** | **Estimate** | **Std. Error** | **t** | **p** | **β** |
| --- | --- | --- | --- | --- | --- |
| (Intercept) | -0.00396 | 0.00284 | -1.39 | .177 |  |
| SDNN.LF._log | -0.00189 | 0.00631 | -0.300 | .767 | -1.01 |
| Age | 4.162e-04 | 2.411e-04 | 1.73 | .097 | 0.361 |
| Sex | -7.510e-05 | 8.990e-04 | -0.0835 | .934 | -0.0179 |
| SDNN.LF._log:Age | -2.805e-04 | 5.530e-04 | -0.507 | .617 | -1.50 |
| SDNN.LF._log:Sex | 0.00222 | 0.00161 | 1.37 | .182 | 2.32 |

***Supplemental Table 316.*** *Standardized coefficients values for T1w+FLAIR WML segmentation.*

#### 103.2. 95% confidence intervals

| **Term** | **Lower 95%** | **Upper 95%** |
| --- | --- | --- |
| (Intercept) | -0.00983 | 0.00191 |
| SDNN.LF._log | -0.0149 | 0.0111 |
| Age | -8.143e-05 | 9.139e-04 |
| Sex | -0.00193 | 0.00178 |
| SDNN.LF._log:Age | -0.00142 | 8.608e-04 |
| SDNN.LF._log:Sex | -0.00111 | 0.00554 |

***Supplemental Table 317.*** *95% confidence intervals values for T1w+FLAIR WML segmentation.*

#### 103.3. Breusch-Pagan test

| **Test** | **Statistic** | **df** | **p** |
| --- | --- | --- | --- |
| Breusch-Pagan | 3.364 | 5 | .644 |

***Supplemental Table 318.*** *Breusch-Pagan test values for T1w+FLAIR WML segmentation.*

#### 103.4. Normal Q-Q plot

Supplemental Figure 209. Normal distribution quantile-quantile plot for T1w+FLAIR WML segmentation.

#### 103.5. Residuals vs leverage

Supplemental Figure 210. Residuals vs leverage diagnostics for T1w+FLAIR WML segmentation. The 2 older female subjects are not visualized due to possessing leverage values approaching 1 in this restricted model, preventing the normal calculation of their standardized residuals.

### 104. Periventricular | Old only | T1w+T2w WML vol | SDNN.LF._log | Exploratory restricted interaction model

Model: T1w+T2w WML vol ~ SDNN.LF._log + Age + Sex + Age * SDNN.LF._log + Sex * SDNN.LF._log

#### 104.1. Standardized coefficients

| **Term** | **Estimate** | **Std. Error** | **t** | **p** | **β** |
| --- | --- | --- | --- | --- | --- |
| (Intercept) | -0.00433 | 0.00304 | -1.42 | .168 |  |
| SDNN.LF._log | -0.00474 | 0.00674 | -0.704 | .488 | -2.36 |
| Age | 4.504e-04 | 2.578e-04 | 1.75 | .093 | 0.363 |
| Sex | -5.925e-05 | 9.611e-04 | -0.0616 | .951 | -0.0131 |
| SDNN.LF._log:Age | -1.408e-04 | 5.912e-04 | -0.238 | .814 | -0.699 |
| SDNN.LF._log:Sex | 0.00295 | 0.00172 | 1.71 | .100 | 2.87 |

***Supplemental Table 319.*** *Standardized coefficients values for T1w+T2w WML segmentation.*

#### 104.2. 95% confidence intervals

| **Term** | **Lower 95%** | **Upper 95%** |
| --- | --- | --- |
| (Intercept) | -0.0106 | 0.00195 |
| SDNN.LF._log | -0.0187 | 0.00917 |
| Age | -8.166e-05 | 9.824e-04 |
| Sex | -0.00204 | 0.00192 |
| SDNN.LF._log:Age | -0.00136 | 0.00108 |
| SDNN.LF._log:Sex | -6.059e-04 | 0.00651 |

***Supplemental Table 320.*** *95% confidence intervals values for T1w+T2w WML segmentation.*

#### 104.3. Breusch-Pagan test

| **Test** | **Statistic** | **df** | **p** |
| --- | --- | --- | --- |
| Breusch-Pagan | 2.459 | 5 | .783 |

***Supplemental Table 321.*** *Breusch-Pagan test values for T1w+T2w WML segmentation.*

#### 104.4. Normal Q-Q plot

Supplemental Figure 211. Normal distribution quantile-quantile plot for T1w+T2w WML segmentation.

#### 104.5. Residuals vs leverage

Supplemental Figure 212. Residuals vs leverage diagnostics for T1w+T2w WML segmentation. The 2 older female subjects are not visualized due to possessing leverage values approaching 1 in this restricted model, preventing the normal calculation of their standardized residuals.

### 105. Periventricular | Old only | T1w-alone WML vol | SDNN.LF._log | Exploratory restricted interaction model

Model: T1w-alone WML vol ~ SDNN.LF._log + Age + Sex + Age * SDNN.LF._log + Sex * SDNN.LF._log

#### 105.1. Standardized coefficients

| **Term** | **Estimate** | **Std. Error** | **t** | **p** | **β** |
| --- | --- | --- | --- | --- | --- |
| (Intercept) | -0.00294 | 0.00261 | -1.13 | .271 |  |
| SDNN.LF._log | -0.00339 | 0.00578 | -0.587 | .563 | -2.0 |
| Age | 3.315e-04 | 2.210e-04 | 1.50 | .147 | 0.316 |
| Sex | -1.700e-04 | 8.240e-04 | -0.206 | .838 | -0.0445 |
| SDNN.LF._log:Age | -1.624e-04 | 5.069e-04 | -0.320 | .751 | -0.952 |
| SDNN.LF._log:Sex | 0.00239 | 0.00148 | 1.61 | .120 | 2.74 |

***Supplemental Table 322.*** *Standardized coefficients values for T1w-alone WML segmentation.*

#### 105.2. 95% confidence intervals

| **Term** | **Lower 95%** | **Upper 95%** |
| --- | --- | --- |
| (Intercept) | -0.00832 | 0.00244 |
| SDNN.LF._log | -0.0153 | 0.00854 |
| Age | -1.246e-04 | 7.877e-04 |
| Sex | -0.00187 | 0.00153 |
| SDNN.LF._log:Age | -0.00121 | 8.838e-04 |
| SDNN.LF._log:Sex | -6.645e-04 | 0.00544 |

***Supplemental Table 323.*** *95% confidence intervals values for T1w-alone WML segmentation.*

#### 105.3. Breusch-Pagan test

| **Test** | **Statistic** | **df** | **p** |
| --- | --- | --- | --- |
| Breusch-Pagan | 2.707 | 5 | .745 |

***Supplemental Table 324.*** *Breusch-Pagan test values for T1w-alone WML segmentation.*

#### 105.4. Normal Q-Q plot

Supplemental Figure 213. Normal distribution quantile-quantile plot for T1w-alone WML segmentation.

#### 105.5. Residuals vs leverage

Supplemental Figure 214. Residuals vs leverage diagnostics for T1w-alone WML segmentation. The 2 older female subjects are not visualized due to possessing leverage values approaching 1 in this restricted model, preventing the normal calculation of their standardized residuals.

### 106. Periventricular | Old only | T1w+FLAIR WML vol | SDNN.HF._log | Exploratory restricted interaction model

Model: T1w+FLAIR WML vol ~ SDNN.HF._log + Age + Sex + Age * SDNN.HF._log + Sex * SDNN.HF._log

#### 106.1. Standardized coefficients

| **Term** | **Estimate** | **Std. Error** | **t** | **p** | **β** |
| --- | --- | --- | --- | --- | --- |
| (Intercept) | -0.00721 | 0.00351 | -2.05 | .051 |  |
| SDNN.HF._log | -0.0328 | 0.0224 | -1.47 | .155 | -15.8 |
| Age | 4.255e-04 | 2.393e-04 | 1.78 | .088 | 0.369 |
| Sex | 0.00151 | 0.00107 | 1.41 | .172 | 0.360 |
| SDNN.HF._log:Age | -1.284e-04 | 5.664e-04 | -0.227 | .823 | -0.619 |
| SDNN.HF._log:Sex | 0.0169 | 0.0110 | 1.53 | .138 | 16.3 |

***Supplemental Table 325.*** *Standardized coefficients values for T1w+FLAIR WML segmentation.*

#### 106.2. 95% confidence intervals

| **Term** | **Lower 95%** | **Upper 95%** |
| --- | --- | --- |
| (Intercept) | -0.0145 | 3.675e-05 |
| SDNN.HF._log | -0.0790 | 0.0133 |
| Age | -6.843e-05 | 9.194e-04 |
| Sex | -7.023e-04 | 0.00372 |
| SDNN.HF._log:Age | -0.00130 | 0.00104 |
| SDNN.HF._log:Sex | -0.00586 | 0.0397 |

***Supplemental Table 326.*** *95% confidence intervals values for T1w+FLAIR WML segmentation.*

#### 106.3. Breusch-Pagan test

| **Test** | **Statistic** | **df** | **p** |
| --- | --- | --- | --- |
| Breusch-Pagan | 3.446 | 5 | .632 |

***Supplemental Table 327.*** *Breusch-Pagan test values for T1w+FLAIR WML segmentation.*

#### 106.4. Normal Q-Q plot

Supplemental Figure 215. Normal distribution quantile-quantile plot for T1w+FLAIR WML segmentation.

#### 106.5. Residuals vs leverage

Supplemental Figure 216. Residuals vs leverage diagnostics for T1w+FLAIR WML segmentation. The 2 older female subjects are not visualized due to possessing leverage values approaching 1 in this restricted model, preventing the normal calculation of their standardized residuals.

### 107. Periventricular | Old only | T1w+T2w WML vol | SDNN.HF._log | Exploratory restricted interaction model

Model: T1w+T2w WML vol ~ SDNN.HF._log + Age + Sex + Age * SDNN.HF._log + Sex * SDNN.HF._log

#### 107.1. Standardized coefficients

| **Term** | **Estimate** | **Std. Error** | **t** | **p** | **β** |
| --- | --- | --- | --- | --- | --- |
| (Intercept) | -0.00810 | 0.00373 | -2.17 | .040 |  |
| SDNN.HF._log | -0.0437 | 0.0237 | -1.84 | .078 | -19.6 |
| Age | 4.487e-04 | 2.539e-04 | 1.77 | .090 | 0.362 |
| Sex | 0.00184 | 0.00114 | 1.62 | .119 | 0.407 |
| SDNN.HF._log:Age | -2.318e-05 | 6.011e-04 | -0.0386 | .970 | -0.104 |
| SDNN.HF._log:Sex | 0.0218 | 0.0117 | 1.86 | .075 | 19.5 |

***Supplemental Table 328.*** *Standardized coefficients values for T1w+T2w WML segmentation.*

#### 107.2. 95% confidence intervals

| **Term** | **Lower 95%** | **Upper 95%** |
| --- | --- | --- |
| (Intercept) | -0.0158 | -4.075e-04 |
| SDNN.HF._log | -0.0927 | 0.00528 |
| Age | -7.536e-05 | 9.728e-04 |
| Sex | -5.078e-04 | 0.00419 |
| SDNN.HF._log:Age | -0.00126 | 0.00122 |
| SDNN.HF._log:Sex | -0.00237 | 0.0459 |

***Supplemental Table 329.*** *95% confidence intervals values for T1w+T2w WML segmentation.*

#### 107.3. Breusch-Pagan test

| **Test** | **Statistic** | **df** | **p** |
| --- | --- | --- | --- |
| Breusch-Pagan | 2.526 | 5 | .773 |

***Supplemental Table 330.*** *Breusch-Pagan test values for T1w+T2w WML segmentation.*

#### 107.4. Normal Q-Q plot

Supplemental Figure 217. Normal distribution quantile-quantile plot for T1w+T2w WML segmentation.

#### 107.5. Residuals vs leverage

Supplemental Figure 218. Residuals vs leverage diagnostics for T1w+T2w WML segmentation. The 2 older female subjects are not visualized due to possessing leverage values approaching 1 in this restricted model, preventing the normal calculation of their standardized residuals.

### 108. Periventricular | Old only | T1w-alone WML vol | SDNN.HF._log | Exploratory restricted interaction model

Model: T1w-alone WML vol ~ SDNN.HF._log + Age + Sex + Age * SDNN.HF._log + Sex * SDNN.HF._log

#### 108.1. Standardized coefficients

| **Term** | **Estimate** | **Std. Error** | **t** | **p** | **β** |
| --- | --- | --- | --- | --- | --- |
| (Intercept) | -0.00618 | 0.00321 | -1.93 | .066 |  |
| SDNN.HF._log | -0.0364 | 0.0204 | -1.78 | .088 | -19.2 |
| Age | 3.366e-04 | 2.186e-04 | 1.54 | .137 | 0.320 |
| Sex | 0.00143 | 9.791e-04 | 1.46 | .157 | 0.374 |
| SDNN.HF._log:Age | 4.590e-05 | 5.176e-04 | 0.0887 | .930 | 0.243 |
| SDNN.HF._log:Sex | 0.0178 | 0.0101 | 1.77 | .090 | 18.8 |

***Supplemental Table 331.*** *Standardized coefficients values for T1w-alone WML segmentation.*

#### 108.2. 95% confidence intervals

| **Term** | **Lower 95%** | **Upper 95%** |
| --- | --- | --- |
| (Intercept) | -0.0128 | 4.452e-04 |
| SDNN.HF._log | -0.0785 | 0.00580 |
| Age | -1.147e-04 | 7.879e-04 |
| Sex | -5.907e-04 | 0.00345 |
| SDNN.HF._log:Age | -0.00102 | 0.00111 |
| SDNN.HF._log:Sex | -0.00299 | 0.0386 |

***Supplemental Table 332.*** *95% confidence intervals values for T1w-alone WML segmentation.*

#### 108.3. Breusch-Pagan test

| **Test** | **Statistic** | **df** | **p** |
| --- | --- | --- | --- |
| Breusch-Pagan | 2.882 | 5 | .718 |

***Supplemental Table 333.*** *Breusch-Pagan test values for T1w-alone WML segmentation.*

#### 108.4. Normal Q-Q plot

Supplemental Figure 219. Normal distribution quantile-quantile plot for T1w-alone WML segmentation.

#### 108.5. Residuals vs leverage

Supplemental Figure 220. Residuals vs leverage diagnostics for T1w-alone WML segmentation. The 2 older female subjects are not visualized due to possessing leverage values approaching 1 in this restricted model, preventing the normal calculation of their standardized residuals.
